# Choices to landscapes: Mechanisms of animal movement scale to landscape patterns of space use

**DOI:** 10.1101/2024.02.28.582548

**Authors:** Will Rogers, Scott Yanco, Walter Jetz

## Abstract

Understanding the geographic distributions of animals is central to ecological inquiry and conservation planning. Movement-based habitat selection models offer a powerful tool for identifying preferred environmental attributes, yet applying these models to predict animal geographic distributions faces methodological and computational challenges. Here, we present a framework that integrates habitat selection and movement behaviors to generate landscape-scale space use predictions. Through simulations and empirical data, we demonstrate that combining local selection and movement dynamics yields highly accurate emergent spatial distribution predictions. Our framework outperforms occurrence-based frameworks across individual, population, and regional scales. By explicitly addressing the role of movement constraints and selection patterns in heterogeneous environments, our framework bridges animal movement and spatial distribution modeling in a scalable manner. This approach offers a new paradigm to link organism-environment interactions from individual space use to habitat connectivity and population distributions relevant to policy and conservation.

## Introduction

Understanding how organisms interact with their environments is foundational to ecological inquiry, revealing critical insights into spatiotemporal patterns of biodiversity (Buckley et al. 2011; Woodin et al. 2013; Evans et al. 2015), population dynamics (Bozinovic et al. 2011; Madin et al. 2012), and genetic connectivity (Cushman & Lewis 2010; Fletcher et al. 2022). Beyond theory, accurately linking where organisms occur with the environmental conditions they select is essential for conservation planning and ecological forecasting (Urban 2015). Models based on spatial observation data, which make scalable predictions by correlating species occurrences with environmental conditions, often average over the underlying ecological mechanisms (Elith & Leathwick 2009). While they provide broad applicability, these models risk failure under novel conditions, especially as biotic interactions shift with exposure to changing climates and land use (Pearson & Dawson 2003). Mechanistic models, in contrast, aim to identify concrete environmental relationships driving species occurrence (Kearney & Porter 2009; Buckley et al. 2011). However, mechanistic processes are challenging to integrate into spatial predictions, such models may fall short of offering actionable insights and, worse, may even fail compared to pattern-based models. The recent explosion and projected growth of individual-based tracking data (Kays & Wikelski 2023) amplifies this challenge amidst a suite of new opportunities for biodiversity change monitoring (Jetz et al. 2022; Davidson et al. 2025), underscoring the need for approaches that can translate fine-scale behavioral data into population-or species-level spatial predictions (Nathan et al. 2008; Hooten et al. 2017). Capturing these interactions requires more than presence data; it demands a nuanced understanding of how animal behaviors like movement and habitat selection shape observed patterns within complex, fragmented landscapes (Schick et al. 2008; Matthiopoulos et al. 2020). Resolving how to incorporate mechanistic insights of animal movement into scalable, predictive models of space use is critical as we confront the twin global crises of environmental change and biodiversity loss (Powers & Jetz 2019).

Inferring relationships between animals and their environment has traditionally relied on observations of occurrence patterns over some set of environmental contexts which vary geographically (Boyce & McDonald 1999; Boyce et al. 2002), typically by using environmental dimensions to contrast observations from locations where organisms are putatively absent (Mackey & Lindenmayer 2001; Johnson et al. 2006; Moorcroft & Barnett 2008). For example, species distribution models (SDMs) compare the patterns of presences as compared to possible null distributions over large geographic extents (Elith et al. 2011; Merow et al. 2014). Resource selection functions (RSFs) – a methodologically similar framework to SDMs (Fieberg et al. 2021; Northrup et al. 2022) – use logistic regression to model the probability of individual occurrences within some domain of predefined availability, like individual home ranges (Boyce & McDonald 1999; Manly et al. 2007). Both SDMs and RSFs are occurrence-based frameworks that capture information about multiple biological mechanisms leading to species or individual occurrences while accounting for elements of the observation process itself (Keating & Cherry 2004; Phillips et al. 2009; Sastre & Lobo 2009; Wisz & Guisan 2009; Newbold et al. 2010). Notably, SDMs and RSFs assume that all environmental conditions within the landscape, an animal’s home range, or other selection domains were instantaneously available for that animal to select, irrespective of the geographic configuration of these environments or the sequence of such choices made by the organism.

Movement constraints and habitat arrangement limit an organism’s access to certain environmental conditions within its geographic range (Fig. 1A), as isolated patches of preferred habitat may be unreachable and, therefore, unused (Soberon & Peterson 2005; Matthiopoulos et al. 2020; Van Moorter et al. 2023). These limitations shape spatial occurrence patterns, especially in heterogeneous or movement-restricted landscapes (Fig. 1B) (Barnett & Moorcroft 2008; Signer et al. 2017). This is an important problem overlooked by SDMs and RSFs, which unwittingly average over movement and dispersal as a driver of selection. Step-selection functions (SSFs) address this deficiency by using GPS data to model selection based on quasi-realistic, stepwise availability conditioned by movement constraints and even estimating those constraints concomitantly (Avgar et al. 2016). However, SSFs lack a natural method to directly predict geographic distributions because they infer preference conditioned on each step, functionally limiting spatial predictions to comparisons against an (often arbitrarily) chosen baseline. This relativized projection of SSFs also ignores the movement constraints inherent in model parametrization and risks misrepresenting actual use across a landscape (Signer et al. 2017). Solutions such as SSF-based agent-based simulations (Signer et al. 2017; Potts & Borger 2023) or Markov chain Monte Carlo adaptations of movement frameworks (Michelot et al. 2019) provide compelling frameworks for projecting local movement mechanisms, but these methods are computationally intensive and lack guaranteed convergence. Multi-stage frameworks, which separately model resource selection (location quality) and step selection (connectivity), offer scalable alternatives but require complex discretization of movement and selection mechanisms (Van Moorter et al. 2023). Bridging movement constraints and environmental selection in geographic projections remains critical to linking fine-scale organism-environment interactions to broader spatial distributions for ecological and conservation applications (Schick et al. 2008; Matthiopoulos 2022).

To address this problem, we introduce an approach that leverages SSFs to model concurrent movement and selection processes, enabling geographically explicit predictions of habitat use that (i) mechanistically capture organism-environment interactions and (ii) improve accuracy over occurrence-based frameworks by incorporating movement constraints. Specifically, we fit SSFs to estimate local-scale movement and retrieve stable-state distributions (SSDs) through eigendecomposition within an ergodic framework. We then compare our combined framework to occurrence-based (RSF) and selection-only (SSF) models to evaluate predictive accuracy across varying animal selection patterns, landscape heterogeneity, and three empirical cases of increasingly out-of-sample prediction.

## Methods

We introduce a way to isolate expected population-level space use from individually fit SSFs, capitalizing on locally estimated selection and movement patterns. In this framework, local movement probabilities are defined through both fine-scale habitat selection and distance-weighted movement. These local movement patterns are used to generate population-level space use through eigendecomposition of movement probabilities, generating a steady-state distribution of probability of use for the landscape.

### Landscapes as transition matrices

We first considered a landscape, G, made up of n cells along with environmental data (Fig. 2A). We treated the occupancy of these cells as discrete states, between which individuals might transition according to rates described by a stochastic transition matrix, A, of n rows and columns. We defined each value in this matrix, Ai,j, as the probability of moving from one cell (Gi) to another (Gj) in the landscape. For an animal in Gi, the sum of all transition probabilities to all other cells in G (including Gi) must sum to one:

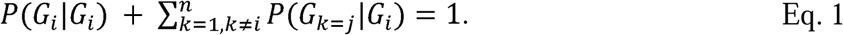

In this way, Matrix A contains the directional probabilities of movement between all cells in G based on beginning in any given cell, Gi, parametrized by later movement modeling and creating a row-stochastic matrix. The inverse of this matrix, A-1, defines a column-stochastic matrix with a first principal eigenvector (associated with an eigenvalue of one) defining the probabilities of the stable-state distribution across G (Fletcher et al. 2019).

### SSFs can parametrize transition matrices

To gather these transition probabilities, we used predicted relative risk informed by fitted SSFs, a quantity derived by comparing baseline conditions (at Gi) to a set of available conditions (G) and often described as a relative strength of selection (Avgar et al. 2017). Relative risks are not equivalent to probabilities but can be kernelized when choices are finite. To do this, we divide Eq. 1 by the probability of moving from Gi to Gi:

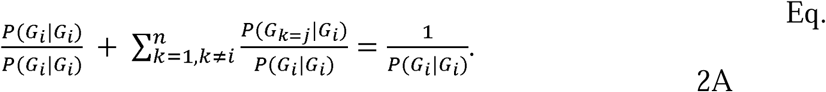

Therefore, Eq. 2A shows that the sum of relative risks for moving to all cells in G is equal to the inverse probability of remaining in the current cell. Or, simplified, the inverse of the sum of all relative selection strengths is equal to the probability of remaining in Gi (Eq. 2B):

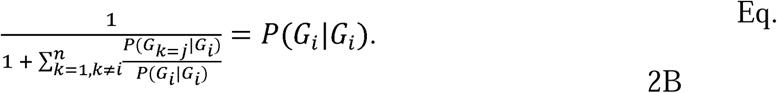

More importantly, Eq. 2B now describes how to link a limited set of relative selection strengths to the probabilities of remaining in place. Therefore, we could solve for the probabilities of moving to all comparison cells by multiplying the relative selection strength returned by the SSF predictions by the probability of staying in place.

### Extension to Integrated Step Selection Functions

SSFs can also explicitly incorporate behavioral elements of movement within habitat selection models. Integrated SSFs (iSSFs) include terms to capture movement kernels, resulting in a distributional description of the likely movement distances around a focal position (Avgar et al. 2016). Functionally, this movement kernel can force the probability of selection for very long-distance movements to zero even when habitat conditions are otherwise suitable. In this way, iSSF models can define local neighborhoods of functionally available habitat for animal movement. By incorporating distance into SSF models, we can confidently compare Gi to cells in G within a biologically informed limit on neighborhood distances (herein the 95th percentile of observed step distances, Fig. 2G). This substantially reduces the information contained in the transition matrix A, treating it as a sparse matrix containing only transitions to cells within a reasonable biological limit of movement (Fig. 2H) and enabling stable-state distribution calculations (Fig. 2K). We then compared SSD predictions to pattern-only (RSF, Figs. 2B, 2D, 2I) and selection-only predictions (SSF, Figs. 2F, 2J).

### Simulation

We stimulated animal movements based on an SSF framework proposed by Potts and Borger (2023). We generated random environmental variables by simulating four environmental variables (100 cells x 100 cells), each with 10 different scales of spatially autocorrelated Gaussian random fields (Hanson et al. 2024) (S1 Figs. 1-2). We also specified theoretical animals with 10 different magnitudes of resource selection across the four environmental variables (β1-4, S1 Table 1). For each set of spatial autocorrelation scales (landscape heterogeneity) and animal selection strength, we multiplied selection coefficients (β1-4) by respective environmental surfaces (1-4) to generate a selection surface (S1 Fig. 3). We repeated this process, creating five random landscapes per level of heterogeneity. We used these selection surfaces, a step length constraint, and a tendency to maintain step angles across steps to simulate animal movements (See S1 for more details). Two sets of movements were generated per simulation: a focal set of 10,000 steps used in modeling (S1 Fig. 4) and a non-focal set of 500 random animals each for 1,000 steps (500,000 total) for validation. We then compared kernelized selection and simulated space use across levels of landscape heterogeneity and animal selection strength (Fig. 1C-E).

For each of the 500 simulations, we converted the focal 10,000 steps into movement tracks, jittered locations slightly to create a positive distribution of step lengths, and sampled 100 random steps per observed step using the “amt” package (Signer et al. 2019). Based on observed step lengths, we sampled random steps from a uniform distribution with a minimum of 0 and a maximum 25% larger than the maximum observed step length. This increased random sampling of step distances both close and far from the animal mover. We fit SSFs by treating the step-terminus values of the four input environmental layers as predictors and including step length and log-step length as predictors using the “amt” package (S1 Table 2) (Signer et al. 2019). We used the fitted SSF models to parametrize A and generate stable-state distributions for each simulation (S1 Fig. 8). We then generated opposing selection-only predictions by using SSFs to predict the selection probability for all cells in the landscape using a random cell in the landscape as a baseline (S1 Fig. 9). Finally, we fit RSFs using the “amt” package, sampling 10 random points per occurrence within the minimum convex polygon of use locations and treating the four input rasters as covariates (S1 Table 2) (Signer et al. 2019). We then used these RSFs to generate opposing spatial predictions to SSD and SSF for each of the 500 simulations (S1 Fig.10).

To isolate the effect of landscape heterogeneity and animal selection patterns on inferences about animal selection (S1 Figs. 11-12) and spatial predictions (S1 Figs. 12-25), we primarily used Shannon entropy, relative entropy (or Kullback–Leibler divergence), and patterns of cumulative use probability. Shannon entropy, defined as 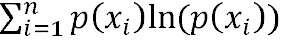 where p(xi) is the probability of use of a cell in the landscape, is a measure of heterogeneity in probability distributions. As Shannon entropy decreases, probability distribution outcomes become more certain; for landscapes, this means that animal space use is heterogeneously concentrated in relatively few locations. Relative entropy, or Kullback–Leibler divergence, was measured as 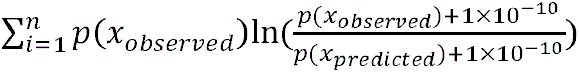 where p(xobserved) is the probability of use of a cell in the landscape in the validation 500,000 steps and p(xpredicted) is the probability of use of a cell in the landscape in a spatial prediction method. The constant 1X 10-10 ensures that sites which are unvisited in validation movements are defined. As relative entropy increases, the worse that a predicted probability distribution approximates the observed (or true) probability distribution. Cumulative use probability was calculated by ordering probability distributions from highest to lowest relative probability of use and then calculating the cumulative sum. For comparisons, we specifically quantified the total area of cells required to generate a cumulative probability of use of0.5, or the minimum area in which an animal is expected to use 50% of the time. Further description of the simulation and more extensive methods are provided in the Supplement (S1).

### Case Study #1 – Intra-individual prediction

To demonstrate the predictive utility of SSD for individual space use, we tested the predictive accuracy of SSD vs. SSF and RSF using a red deer (*Cervus elaphus*) dataset for a single individual included in the “amt” package (Signer *et al*. 2019), using distance to forest and forest cover (calculated from a binary forest raster provided in the “amt” package, aggregated to a resolution of 125m by 125m, S2 Table 1). We separated the first (chronologically) 75% of observed data as an in-sample training dataset and retained the latter 25% of observed data as an out-of-sample testing dataset (S2 Fig. 1). As above, we sampled random steps from a wider uniform distribution with 100 random points per observed step. We included an interaction between distance to forest and forest cover in models along with step length and log-step length in SSF models (S2 Table 2). We generated the SSD as above using a neighborhood defined by the 95^th^ percentile of observed step lengths (S2 Fig. 2). We then generated the RSF and SSF probabilities as in simulations using the same predictor structures (S2 Table 2, S2 Figs. 3-4).

**Figure 1:**
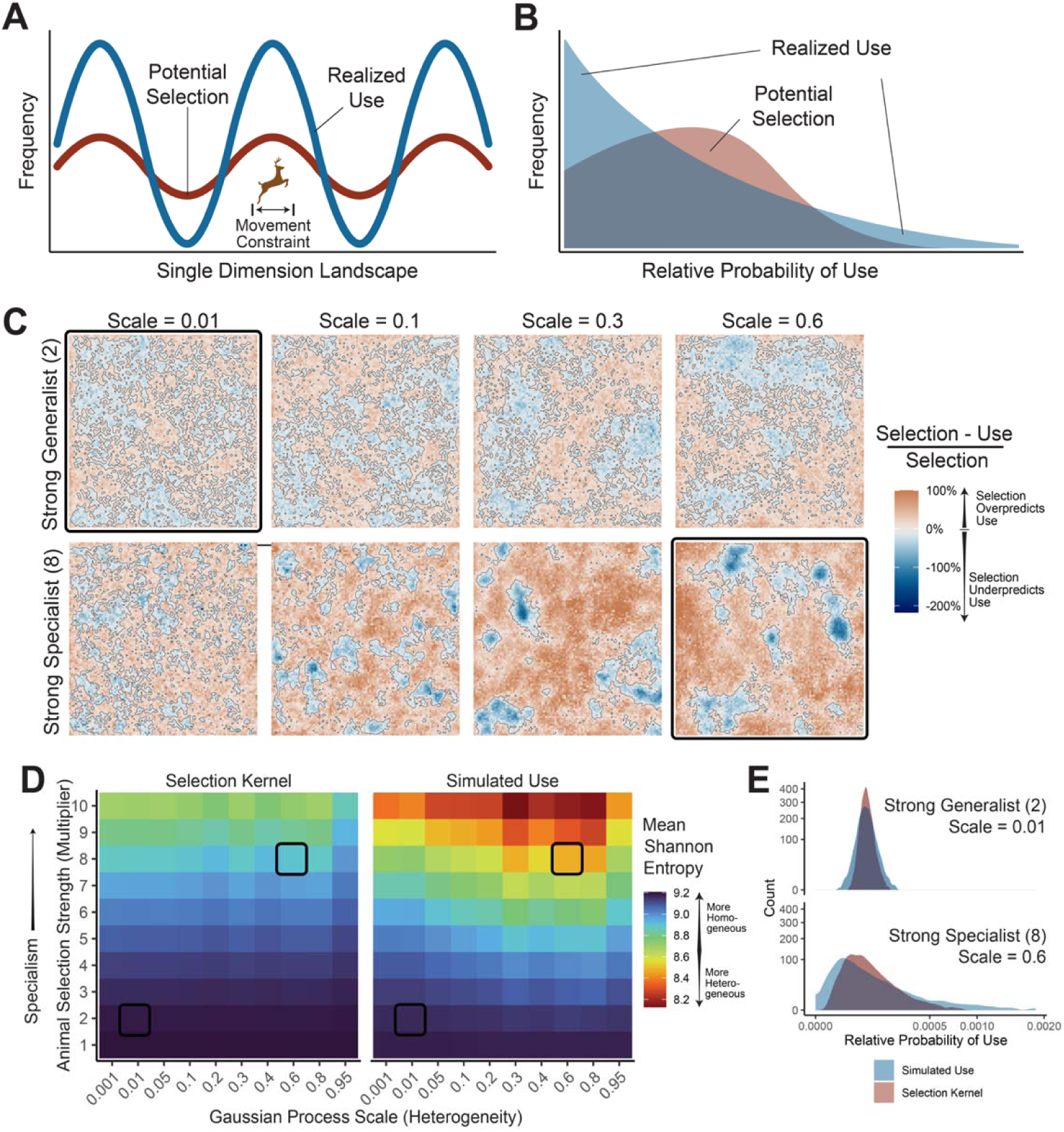
Habitat preferences do not equal realized use in simulations. (**A**) The theoretical frequency of use of a single-dimensional environment based on the preference (potential use, red) vs. the realized frequency of use given movement constraints in a heterogeneous habitat (blue). (**B**) Corresponding depictions of the frequency of use of locations in the landscape without (red) and with movement constraints (blue). (**C**) The effect of landscape heterogeneity (Gaussian random fields scale) and habitat selection strength multiplier (2 – weak; 8 – strong) on the relative difference between habitat selection and true probability of use in a single set of simulations, calculated as the relative selection kernel minus true use probability kernel divided by the relative selection kernel. Differences in surfaces are red (where realized selection overpredicts potential use probability) and blue (where realized selection underpredicts potential use probability) and contours highlight where transitions occur. (**D**) The mean Shannon entropy across 5 simulation iterations per level of selection strength multiplier (y-axis) and landscape heterogeneity (x-axis) for the relative selection kernel and the true probability of use. (**E**) For two sets of simulations (with specialist selection [multiplier = 8] and high heterogeneity [scale = 0.6] or generalist selection [multiplier = 2] and low heterogeneity [scale = 0.01]; highlighted by black squares in **C** and **D**), density plots of relative probability of use based on selection kernels (red) and actual probability of use in simulations (blue).

**Figure 2:**
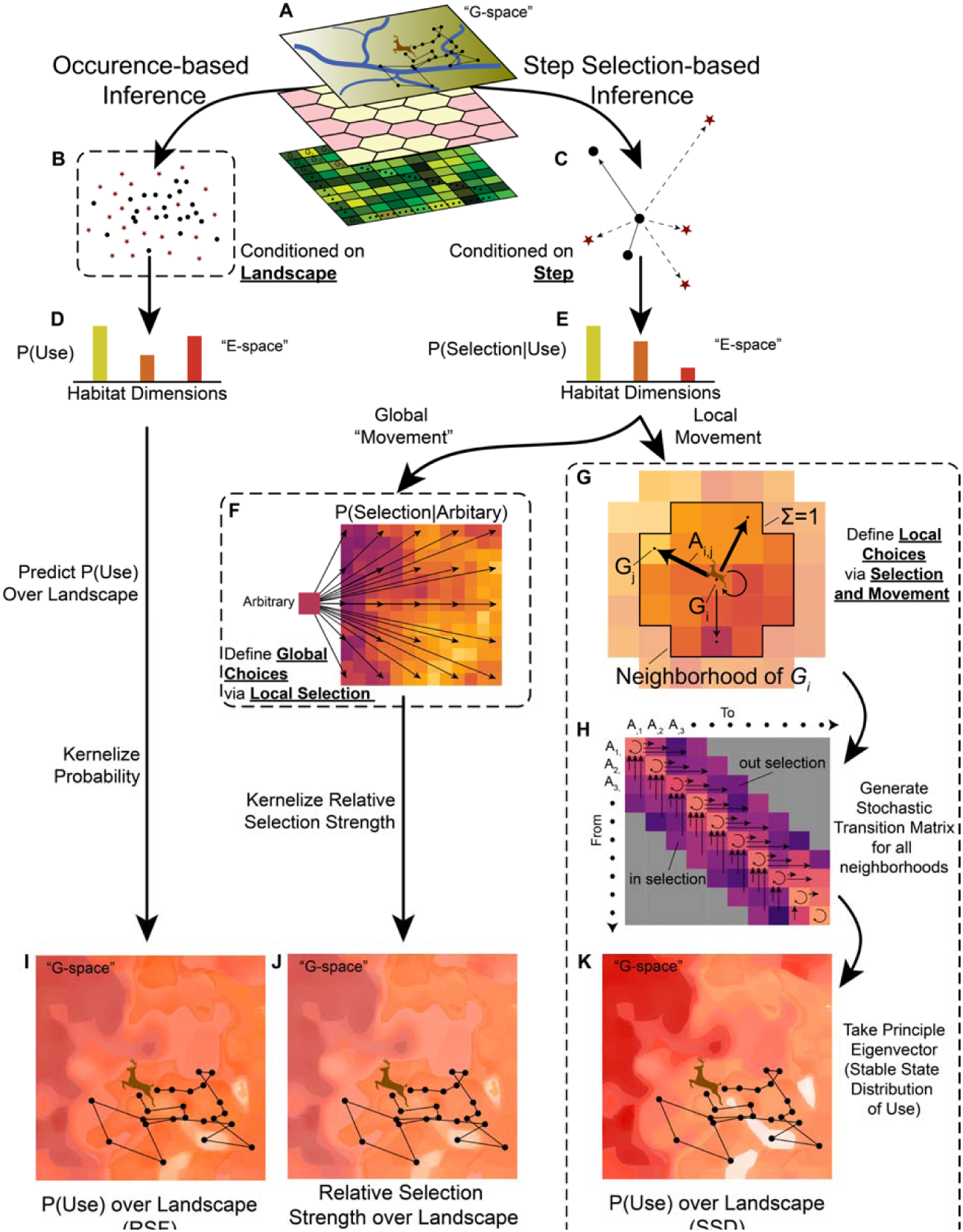
Movement data-based landscape predictions. Converting animal movements into spatial predictions of use can occur in three main ways. First, (**A**) animal movements are compared to environmental contexts in landscapes made of discrete cells, *G*. (**B**) Animal locations can then be placed into use vs. available frameworks conditional on the landscape, suitable for resource selection analyses (RSFs). (**C**) Alternatively, tracking data can be converted into use vs. available frameworks conditional on steps, suitable for step selection analyses (SSFs). (**D**) RSFs predict the probability of use over a landscape, which can be kernelized to return (**I**) probabilities of use over *G*. (**E**) SSFs predict the relative probability between comparisons within a context, a distinct metric from the probability of use. (**F**) It is common to make an arbitrary comparison between some either observed or unobserved environmental contexts in *G* to all possible contexts in *G*. (**J**) These spatial relative risks are then kernelized, representing something akin to RSF output, though mathematically inequivalent. (**G**) Harnessing the ability of SSFs to parametrize local movement patterns, we can algebraically solve the probability of moving within a set of local environmental contexts (*G_j_*) by defining that the relative selection strength for a focal cell (*G_i_*) compared to itself must equal 1. (**H**) Aggregating patterns of in– and out-selection over all neighborhoods of all *G_i_* resolves a sparse stochastic matrix for all reasonable movements. (**K**) Solving for the principal eigenvector of the stochastic matrix returns the stable state distribution – the probability of ending in some *G_i_* given all possible starts over the landscape, *G*.

**Figure 3:**
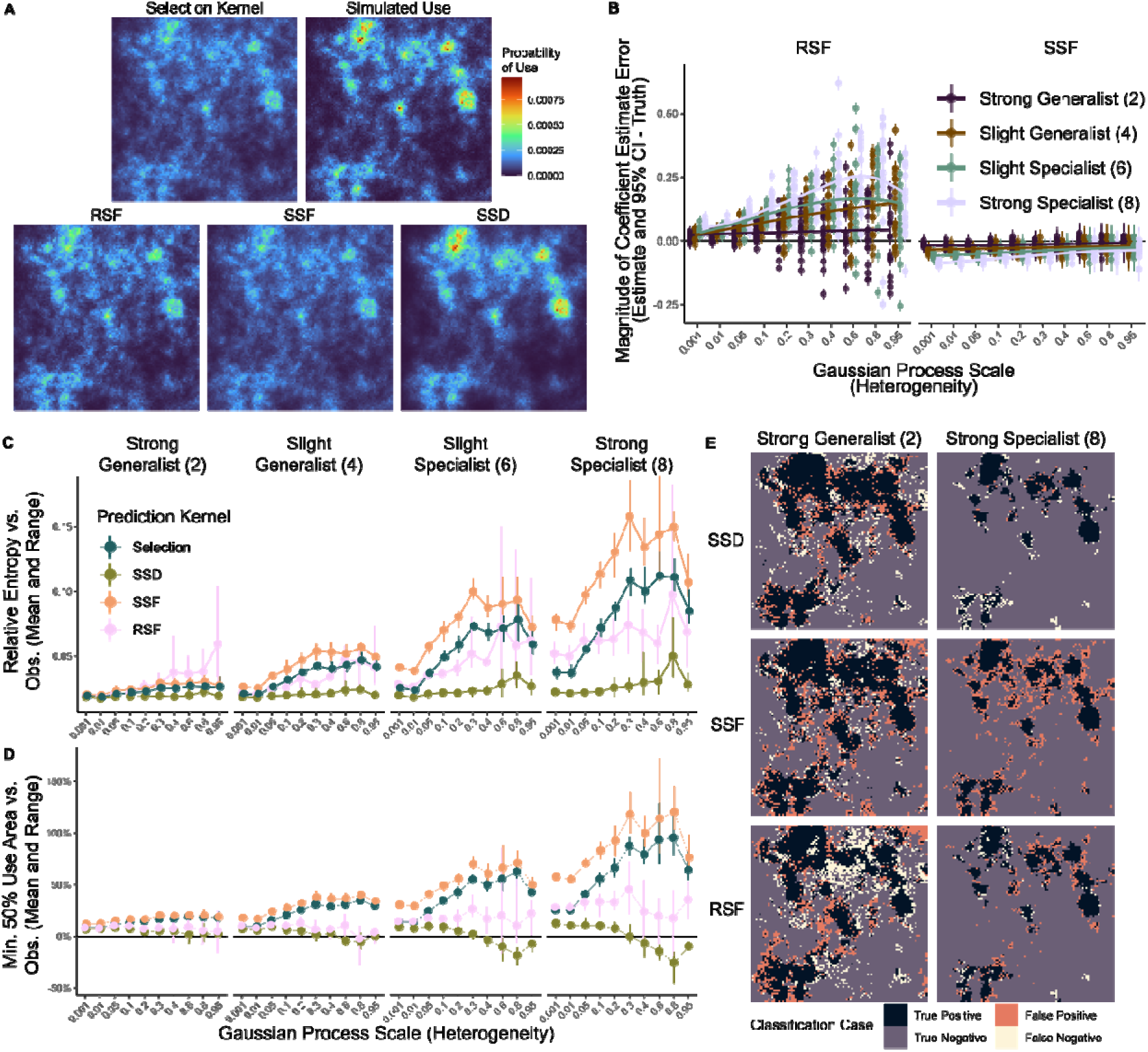
Predictions incorporating movement mechanisms recover simulated patterns of use. Within simulations, we generated (**A**) a selection kernel (the product of selection strengths for four randomly generated environmental variables), simulated use probability from 500,000 steps of animals with constraints on step length and turn angle, and predictions generated from a focal set of 10,000 steps for resource selection function (RSF), kernelized step-selection function (SSF) relative selection strength, and the stable state distribution (SSD). (**B**) Five sets of simulations with different levels of Gaussian random fields scale (x-axis) and strengths of animal selection (multipliers, color scale) revealed differences between the estimated selection coefficient for each environmental variable and the true value of the selection coefficient based on RSF or SSF frameworks. We converted errors and 95% confidence intervals (CIs) such that directions of comparison are aligned (e.g., positive differences imply more extreme inferences, irrespective of positive or negative selection). We summarized the performance of predictions relative to simulated use via (**C**) relative entropy and (**D**) top 50% use area of known selection kernels and RSF, SSF, and SSD predictions vs. simulated use across varying levels of heterogeneity (x-axis) and animal selection strengths (horizontal facet). (**E**) SSD, SSF, and RSF resulted in distinct predictions about animal space use, highlighted by classification (fill color) of habitat into top 50% use area vs. simulated use for RSF, SSF, and SSD predictions (vertical facet) and different levels of selection strength (horizontal facet).

**Figure 4:**
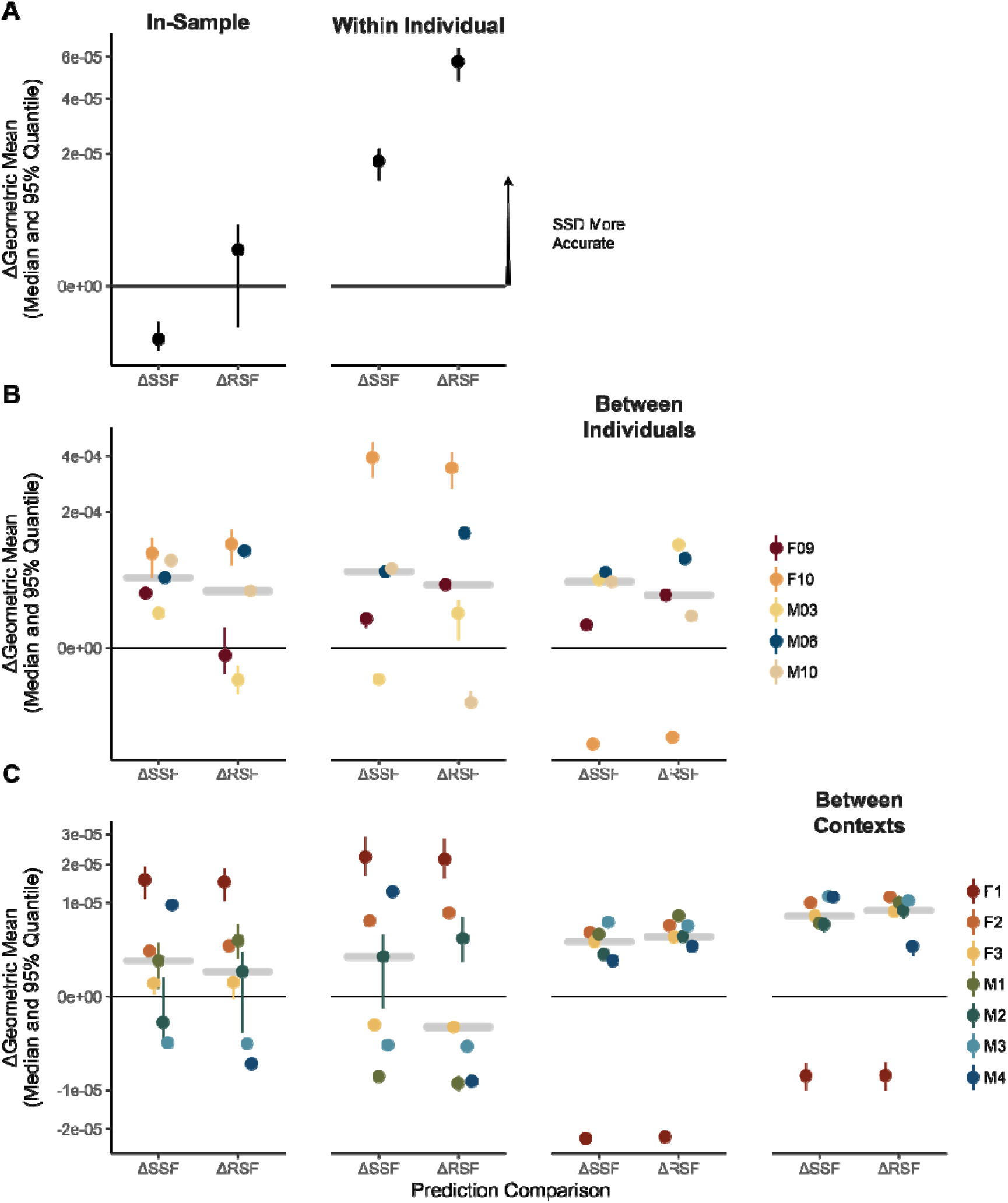
Predictions incorporating movement mechanisms better predict out-of-sample animal movements. (**A-C**) Median and 95% quantiles in 100 bootstraps of differences between stable state distribution (SSD) and resource selection function (RSF) or step-selection function (SSF) predictions in geometric mean of predicted probability of use at observed use sites. Bootstraps were performed by resampling modeled and out-of-sample data with replacement for (**A**) red deer (*Cervus elaphus*), (**B**) roe deer (*Capreolus capreolus*), and (**C**) fisher (*Pekania pennanti*) datasets. Validations were performed in four sets of comparisons, we used the first (chronologically) 75% of observed locations for individuals to predict on the modeled data (“In-Sample”), the next (chronologically) 25% of observed locations per individual (“Within Individual”), all observed locations for the other individuals in a dataset within the same geographic context (“Between Individuals”), and all observed locations for an individual in a completely different environmental context than the individuals modeled (“Between Contexts”). For visualization, the median difference for each set of comparisons is highlighted by a horizontal grey bar.

We then compared the heterogeneity and accuracy of each prediction. To quantify heterogeneity, we calculated the Shannon entropy of each predicted probability distribution. In this context, lower Shannon entropy describes predictions that the intensity of animal use is concentrated in fewer locations. To quantify accuracy, we used two main comparison heuristics. First, following Boyce *et al*. (2002), binned prediction surfaces into 10 bins of equal area and then calculated the Spearman rank correlation between observed use intensity and bin order.

Second, as all prediction kernels summed to one, we calculated the geometric mean of predicted use across use locations to discriminate more accurate predictions. Spearman rank correlations are common in resource selection validation (Morris *et al*. 2016), but they are sensitive to discrete prediction surfaces and our simulations showed that Spearman rank correlations may be biased to select homogeneous prediction surfaces (S1 Fig. 23). The geometric mean of predicted use, in contrast, was better able to accurately recover the difference in performance between predictions and was generally more conservative (S1 Fig. 24). To ensure that comparisons were not sensitive to the underlying out-of-sample datasets, we bootstrapped the observed set of GPS points (sampling with replacement) 100 times, generating 100 estimates of Spearman rank correlations and mean log-probability of use per predicted surface. We calculated the difference between SSD and RSF or SSF predictions to isolate comparisons. We report geometric mean results in the main text and Spearman rank comparisons in S2.

### Case Study #2 – Inter-individual predictions

To demonstrate the predictive utility of SSD across individuals, we used a roe deer (*Capreolus capreolus*) dataset for five individuals in Northern Italy (Cagnacci *et al*. 2011) to assemble the same SSD, SSF, and RSF predictions. We resampled tracks to 4 hours (the median interval in GPS transmissions), and as above, we isolated the first 75% of individual movements as in-sample, and the next 25% as out-of-sample testing datasets (S2 Fig. 5). Because the dataset contained migratory movements, we decided to model only summer movements of deer (defined as Julian day 120-290, S2 Fig. 5). We included square-root distance to forest (calculated from 2006 CORINE landcover classes of conifer, mixed, and broadleaf forests) and slope (calculated from STRM elevation models, aggregated at a scale of 129m by 185.2m) as covariates in our models and predictions (S2 Fig. 6) with step length and log-step length, while the RSF included the square-root of distance to forest and slope (S2 Table 2). We generated SSD (S2 Fig. 7), SSF (S2 Fig. 8), and RSF predictions as above (S2 Fig. 9). We performed out-of-sample evaluations as above with 100 bootstrapped geometric mean predicted use and Spearman rank correlations between surface deciles and observed use intensity in 25% out-of-sample for individuals. Additionally, we used individual-level models (fit using 75% of individual data) to predict the distributions of all other individuals’ spatial use intensity in the dataset and assessed accuracy using the same bootstrapping method (referred to as “between individual” predictions).

**Figure 5:**
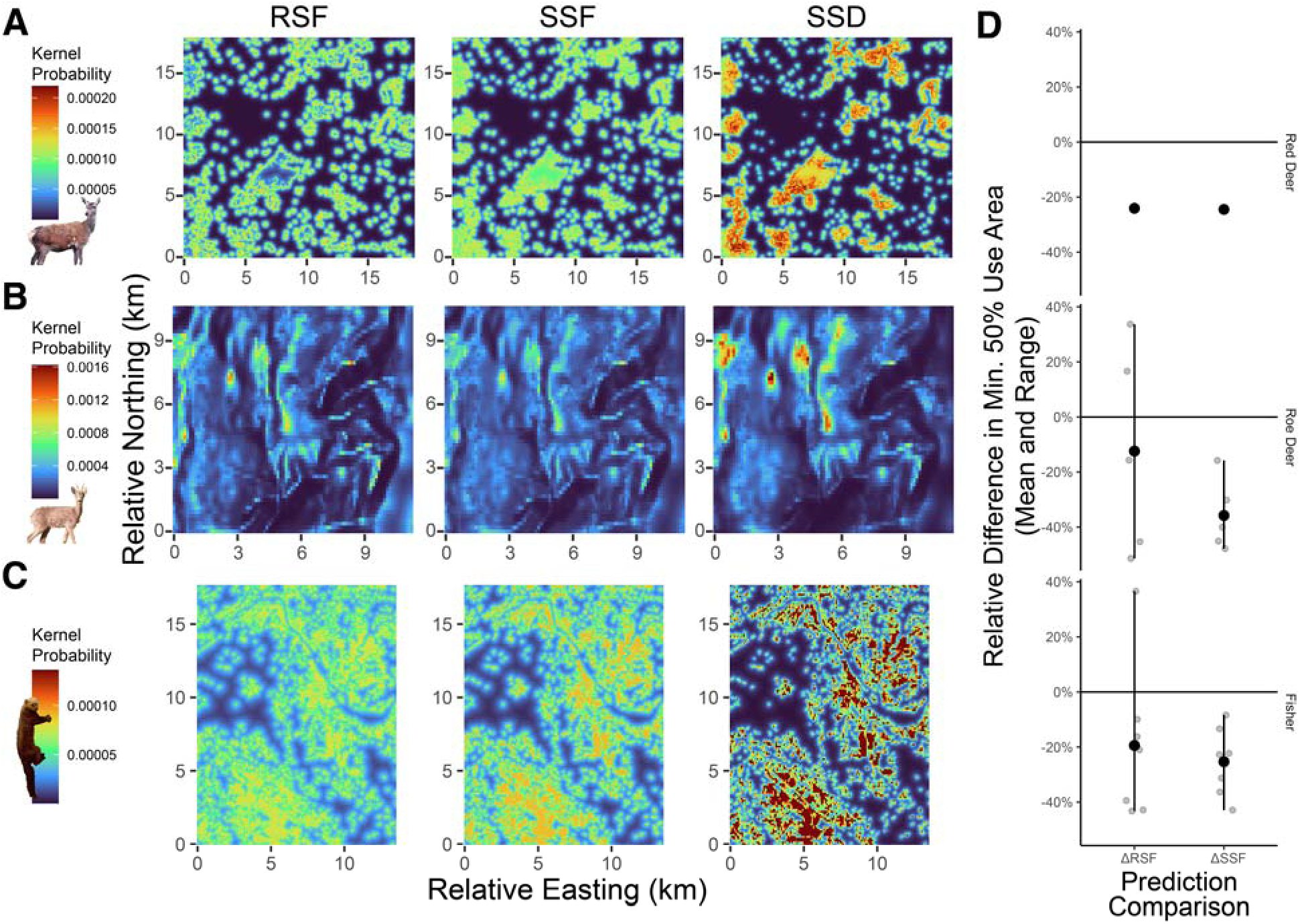
Predictions incorporating movement mechanisms are more spatially concentrated than those based on selection alone. (**A**) Spatial predictions of “use” (as a kernel over the landscape) for a red deer, (**B**) summer use of a single roe deer (“M10”), (**C**) and a single fisher (“M2”) based on output from a resource selection function (RSF), a step-selection function (SSF), and the stable state distribution (SSD) from local movement patterns estimated from the SSF. (**D**) For each case study, the relative difference in minimum area containing 50% of predicted use for SSD vs. RSF (RSF-SSD/RSF) and SSF predictions (SSF-SSD/SSF) are plotted for individuals. For visualization, the mean and range of these relative differences are overlaid on individual estimates. These results highlight that irrespective of accuracy, incorporating movement constraints in SSD always generates more spatially concentrated predictions than SSF, and SSD tends to also be more spatially concentrated than most RSF estimates, meaning that less area may be truly usable for animals after incorporating movement constraints.

### Case Study #3 – Prediction to novel contexts

Lastly, to explore the predictive capacity of the SSD method to novel landscapes, we used a dataset of eight fisher (*Pekania pennanti*) from Northern New York, USA (S2 Fig. 10) (LaPoint *et al*. 2013) to generate SSD, RSF, and SSF probabilities considering a landscape with square-root distance to forests as well as step length and log-step length (S2 Tables 1-2). Seven fisher inhabited a similar environmental extent with a mix of forested and human landscapes (approximately 10km by 15km, S2 Fig. 11), whereas one fisher occupied a completely different landscape approximately 30km away from the others and with far less human infrastructure and population density (S2 Fig. 15). We regarded the seven local fisher as focal individuals for modeling, and we retained the non-local fisher as a unique opportunity to evaluate individual-level predictive performance across environmental contexts. We calculated distance to forest using the National Landcover Database from 2006 to identify land classes which were wetland and deciduous, mixed, and evergreen forest. We dissolved the resulting raster to generate a distance to forest using QGIS (v. 3.22) and GDAL proximity analysis for computational efficiency before aggregating spatially to 91.2m by 123.2m resolution. Because individuals in the fisher dataset had different GPS fix-rates, we resampled tracks to the median fix rate for each animal (range: 2-15 minutes). We performed the same out-of-sample evaluations within individuals (75/25 split) and between individual comparisons (one individual predicting the distribution of all others) for all seven focal fishers using SSD (S2 Fig. 12), SSF (S2 Fig. 13), and RSF surfaces (S2 Fig. 14). We then compared each of the seven focal fisher movement models to predict the spatial distribution of the non-local fisher in the novel environmental context, again using 100 bootstraps of the test dataset to assess the accuracy of SSD (S2 Fig. 16), SSF (S2 Fig. 17), and RSF surfaces (S2 Fig. 18).

### Supplements and code

Description of the method and simulations (S1) and empirical examples (S2) are provided as supplements. The code used to generate these results is available online (https://github.com/will-rogers/SSD_SSF.git) and is provided in a compiled document as a supplement to the main text (S3). All code will be archived on Zenodo upon acceptance.

## Results

We assessed the new framework of stable-state distributions generated from local selection and movement patterns (SSD) in its ability to predict the distribution of organisms across landscapes. To demonstrate its performance to existing approaches we compared predictions to those from occurrence-based (RSF) and selection-alone (SSF) frameworks. To do so, we first isolated the effect of landscape heterogeneity and animal selection strengths in simulations. In three subsequent empirical examples, we showed how SSD, RSF, and SSF predictions vary in ability to predict in– and out-of-sample animal space use.

### Simulation

We simulated animal movement in 100 unique combinations of landscape heterogeneity and animal selection strength in five replicates. We estimated selection coefficients under RSF and SSF frameworks, and compared the predictive performance of RSF, SSF, and SSD predictions to simulated long-term space use. We found that animal selection strength created stark disparities between known patterns of selection and observed patterns of use, particularly at high levels of landscape heterogeneity (Figs. 1C). In more spatially autocorrelated and heterogeneous landscapes, the Shannon entropy of selection was largely unaffected (as autocorrelation affects the spatial structure, not the distribution, of selection values, Fig. 1D).

However, increasing animal selection strength strongly influenced the entropy of selection surfaces (Fig. 1D). Importantly, as selection strength and landscape heterogeneity increased, observed use patterns had much lower entropy (more spatially concentrated) than expected by selection alone (Figs. 1D-E). Comparisons of inferential methods (RSF vs. SSF) revealed that RSF-estimated selection coefficients were significantly biased both as individual selection strength increased but particularly as landscape heterogeneity increased, whereas SSF inferences appeared to generally decrease in bias as landscape heterogeneity increased (Fig. 1B, S1 Figs. 11-12). When comparing prediction surfaces, we found that SSD and, to a lesser degree, RSF predictions better matched heterogeneity in patterns of observed use (S1 Figs. 13-15).

Uniformly, SSD was the best predictor of observed space use, distantly followed by RSF and SSF predictions based on relative entropy, and this pattern was amplified at moderate-to-high landscape heterogeneity and increasing animal selection strengths (Fig. 3C, S1 Figs. 18). Finally, we found that all prediction surfaces overestimated the core space use (minimum 50% use area) of animals as compared to observed use, particularly at low levels of habitat heterogeneity; however, SSD delivered the most spatially concentrated predictions and was consistently the closest estimate of core space use as compared to observed use (Fig. 3D-E, S1 Figs. 16-17). Conversely, for heterogeneous landscapes with highly selective animals, selection-only predictions (either SSF or the true selection kernel) overestimated core 50% use area by more than 125-150% (Fig. 3D-E, S1 Figs. 16-17).

### Case Studies

As in simulations, SSD tended to produce more spatially concentrated predictions than SSF and RSF for the red deer (*n=*1), roe deer (*n=*5), and fisher (*n=*7) datasets (S2 Figs. 1-18). Based on the geometric mean of predicted use, SSD was a better predictor of the modeled first 75% of data per individual than SSF for 10 of 13 animals or RSF for nine of 13 animals (“in-sample”, Fig. 4). This persisted to predictions for the next 25% of out-of-sample animal movements, with SSD favored over SSF for nine of 13 animals and favored over RSF for eight of 13 animals (“within individual”, Fig. 4). For interindividual out-of-sample predictions, SSD was favored over SSF and RSF for 10 of 12 animals (“between individuals”, Fig. 4). Similarly, out-of-sample predictions in a different environmental context favored SSD over SSF and RSF for six of seven individuals (“between contexts”, Fig. 4). Similar patterns were observed in comparisons using Spearman rank correlations (S2 Fig. 19). SSD predictions were more spatially concentrated than SSF and RSF predictions for 100% and 77% of animals, respectively (Fig. 5A-C). Importantly, this difference between SSD and predictions lacking movement constraints translated to substantive reductions in predicted core suitable habitat (based on minimum 50% predicted cumulative use area): SSD predictions estimated 20-40% less core habitat than SSF and 15-25% less core habitat than RSF.

## Discussion

Predicting animal distributions is crucial for ecological insights, conservation, and global policy. Expanding individual-level movement data offers new opportunities for biodiversity monitoring (Jetz *et al*. 2022; Kays & Wikelski 2023; Davidson *et al*. 2025) but also challenges in scaling movement patterns to landscape-level space use. For instance, mechanistic movement-based models like SSFs provide better estimates of habitat selection than pattern-based models like RSFs (Fig. 3B). However, SSFs fail to scale local selection patterns to predict landscape-level space use, resulting in paradoxically poor spatial predictions compared to RSFs (Fig. 3C). We showed that incorporating habitat selection and movement constraints in an ergodic framework (SSD) provides better predictions of animal space use than SSF or RSF in simulations (Fig. 3D) and in most empirical examples – from future movements of modeled animals to those of other animals in analogous or novel environments (Fig. 4A-C). Additionally, by including movement constraints, we demonstrated that SSF and RSF predictions can overestimate usable habitat by 15-40% (Fig. 5D). This work highlights the importance of incorporating both habitat selection and movement constraints to derive landscape-scale insights from individual-level data (Matthiopoulos *et al*. 2020).

Our approach builds on previous frameworks in connectivity modeling (Fletcher *et al*. 2019) as well as excellent theoretical (Barnett & Moorcroft 2008; Matthiopoulos *et al*. 2020; Potts & Borger 2023) and empirical addresses of movement modeling predictions (Signer *et al*. 2017; Michelot *et al*. 2019; Van Moorter *et al*. 2023; Forrest *et al*. 2024). For example, transition matrices underlie *circuitscape* (McRae *et al*. 2008) and more recent movement frameworks (Fletcher *et al*. 2022). Similarly, previous authors have shown that SSF models can be used to iteratively generate spatial distributions via agent-based frameworks (Michelot *et al*. 2019; Potts & Borger 2023; Signer *et al*. 2024). Our innovation lies in using SSFs to parametrize transition matrices with animal-estimated movement probabilities and in making eigendecomposition computationally feasible. Our ergodic framework provides a convenient way to effectively generate infinite simulations of SSF outputs, resulting in a predicted use definition which is aligned with traditional occurrence frameworks.

Our SSD approach offers immediate benefits for conservation, movement diagnostics, and multi-scale ecological inference. For conservation planning, SSD-based predictions outperform RSF and SSF estimates of space use. Accurate distribution predictions are crucial for conservation (Wiens *et al*. 2009; Evans *et al*. 2015; Araújo *et al*. 2019). By empirically capturing movement constraints, SSDs address shortcomings in existing approaches, providing more accurate depictions of space use. SSDs are also more conservative, indicating SSF and RSF models likely overestimate usable habitat. As a result, our framework provides a mechanistic alternative to occurrence-based approaches like SDMs and RSFs.

In connectivity modeling, SSDs provide an empirical method to generate transition matrices using SSFs, where other methods rely on transformations of predicted use surfaces. We expect this approach to result in more biologically realistic connectivity predictions, with implications for identifying conservation priorities, movement corridors, and disease spread in wildlife epidemiology. Further, SSD’s ability to account for animal traits in underlying movement modeling can enhance spatial prioritization efforts tailored to specific ages, sexes, or behavioral phenotypes (e.g., migratory).

As a diagnostic of animal movement modeling, SSD-based approaches offer the ability to spatially visualize SSF model predictions, otherwise only available through simulation from fitted SSFs (Potts & Borger 2023; Signer *et al*. 2024). Diagnosing model parametrization issues remains a large challenge in animal movement modeling (Fieberg *et al*. 2018; Signer *et al*. 2024). SSDs offer computationally efficient ways to map use probabilities, facilitating checks for missing variables or overfitting. These maps provide insights into the biological effects of habitat selection or movement constraints, currently available only via relative risk comparison (Avgar *et al*. 2017). SSD may not be feasible in all scenarios, but they add a valuable tool to refine movement models.

Finally, we envision SSDs will provide richer ecological insights into animal movement data. For example, SSD approaches can effectively create null models of animal space use based only on habitat selection and movement. By comparing observed space use to these simplistic realizations of movement models, we may reveal how factors like social interactions, learning, predators, parasites, or competitors shape individual ecology. Similarly, hierarchically organized movement data (e.g., individual, population, species) and inferential methods (e.g., Muff *et al*. 2019) can harness SSD frameworks to address cross-population or cross-species predictions at large spatial scales. This may provide insights into how individual heterogeneity, local adaptation, or species differences affect space use.

Future work based on this framework should consider expanded spatial scales of prediction and extensions to variable movement and selection dynamics. We considered landscapes with 4,000-22,000 cells, which ran in <5 minutes on a personal computer. Larger-scale predictions may require optimizing cell neighborhood size to balance computational speed and accuracy. Similarly, our framework currently omits turn angles due to computational constraints. Transition matrices model movements between states (cells) but not directional angles, complicating Markovian processes in SSFs. Though step distance often correlates with turn angle, this simplification warrants further exploration. Additionally, we estimated movement kernels using step length and log-step length, sampling unused steps uniformly. Alternative sampling distributions can improve habitat selection estimates but require correcting predictions for the underlying sampling distributions (Fieberg *et al*. 2021; Signer *et al*. 2024). Finally, our framework assumes static behavioral states and environmental conditions. Incorporating behavioral or environmental variability may require simulation frameworks, such as Forrest *et al*. (2024). Nonetheless, power iteration and similar matrix approaches could address temporally dynamic behaviors and landscapes.

In a changing world, animal-environment interactions are also likely to shift, challenging pattern-based models whose projections may falter when decoupled from mechanisms. Mechanistic models, though adaptable, lose accuracy if generating mechanisms are omitted. Our findings suggest combining movement and selection data in spatial projections can bridge these gaps, enhancing the utility of individual movement data for ecological and conservation applications.

## Acknowledgments

We thank Johannes Signer, Francesca Cagnacci, and Scott LaPoint for their generosity in providing the red deer, roe deer, and fisher datasets (respectively) as open-access resources for the scientific community, and for their permission to use those datasets in our study. We also thank John Fieberg for initial discussion and members of the Jetz and Ezenwa labs for their generous feedback on various stages of the project. WR acknowledges funding from the Gruber Foundation. SY acknowledges funding from the Max Planck – Yale Center for Biodiversity Movement and Global Change.

