## Supplement for "Choices to landscapes: Mechanisms of animal movement scale to landscape patterns of space use"

**Authors and affiliations:**

**Table of Contents:**

**S1: Simulation Appendix**

**S1:** Simulation methodology and investigation (pgs. 3-7)

**S1 Table 1:** Animal selection strength coefficients (pg. 8)

**S1 Table 2:** Model pseudocode for the simulation (pg. 9)

**S1 Fig. 1:** Example randomly generated rasters with Gaussian scale (pg. 10)

**S1 Fig. 2:** Example of four simulation rasters used per iteration (pg. 11)

**S1 Fig. 3:** Simulated environmental suitability (pg. 12)

**S1 Fig. 4:** Simulation tracks over environmental suitability (pg. 13)

**S1 Fig. 5:** Visualizing the neighborhood of *G_i_* (pg. 14)

**S1 Fig. 6:** Visualizing the neighborhood of *G_i_* at a proximate scale (pg. 15)

**S1 Fig. 7:** Comparison of selection and stable-state kernels from simulation example (pg. 16)

**S1 Fig. 8:** Comparison of true selection and step-selection-inferred selection kernels from simulation example (pg. 17)

**S1 Fig. 9:** Comparison of true selection and resource selection-inferred probability of use kernels from simulation example (pg. 18)

**S1 Fig. 10:** Proportion of selection coefficients contained in estimate CI by model type, heterogeneity, and animal selection strength (pg. 19)

**S1 Fig. 11:** Absolute error for selection coefficients by model type, heterogeneity, and animal selection strength (pg. 20)

**S1 Fig. 12:** Landscape predictions (SSD, SSF, RSF) vs true selection and observed use kernels (pg. 21)

**S1 Fig. 13:** Variation in observed probability of use vs. known selection kernels as a function of selection strength and landscape heterogeneity (pg. 22)

**S1 Fig. 14:** Mean and range of Shannon entropy for kernels across five simulations (pg. 23)

**S1 Fig. 15:** Landscape classification based on probability of use (pg. 24)

**S1 Fig. 16:** Total minimal area to reach a sum probability of use of 0.5 vs. simulated use across 5 simulations (pg. 25)

**S1 Fig. 17:** Relative entropy of predictions vs. simulated use across 5 simulations (pg. 26)

**S1 Fig. 18:** Root mean squared error of predictions vs. simulated use across 5 simulations (pg. 27)

**S1 Fig. 19:** Accuracy of predictions in classifying top simulated use area across 5 simulations (pg. 28)

**S1 Fig. 20:** Classification cases of prediction kernels vs. simulated use (pg. 29)

**S1 Fig. 21:** Pearson correlation between prediction and observed use kernels (pg. 30)

**S1 Fig. 22:** Spearman correlation between prediction and observed use kernels (pg. 31)

**S1 Fig. 23:** Mean log-predicted use across observed kernels (pg. 32)

**S1 Fig. 24:** Observed use kernels vs. predicted use kernels across all simulations (pg. 33)

**S2: Main Text SI Tables and Figures**

**S2 Table 1:** Data sources (pg. 34)

**S2 Table 2:** Model pseudocode for case studies (pg. 35)

**S2 Fig. 1:** Distance to forest and movement in the red deer (*Cervus elaphus*) case study (pg. 36)

**S2 Fig. 2:** Stable state distribution (SSD) surface for the single individual in the red deer case study (pg. 37)

**S2 Fig. 3:** Exponentiated step-selection function (SSF) surface for the single individual in the red deer case study, kernelized to sum to one (pg. 38)

**S2 Fig. 4:** Probability of use surface from a resource selection function (RSF) for the single individual in the red deer case study, kernelized to sum to one (pg. 39)

**S2 Fig. 5:** Individual roe deer (*Capreolus capreolus*) summer movement (pg. 40)

**S2 Fig. 6:** Visualization of roe deer covariates, in-sample, and out-of-sample data (pg. 41)

**S2 Fig. 7:** SSD predictions for individual roe deer (pg. 42)

**S2 Fig. 8:** SSF predictions for individual roe deer (pg. 43)

**S2 Fig. 9:** RSF predictions for individual roe deer (pg. 44)

**S2 Fig. 10:** Individual Fisher (*Pekania pennanti*) movement (pg. 45)

**S2 Fig. 11:** Visualization of focal fisher covariates, in-sample, and out-of-sample data (pg. 46)

**S2 Fig. 12:** SSD predictions for focal fisher in the focal landscape (pg. 47)

**S2 Fig. 13:** SSF predictions for focal fisher in the focal landscape (pg. 48)

**S2 Fig. 14:** RSF predictions for focal fisher in the focal landscape (pg. 49)

**S2 Fig. 15:** Visualization of non-focal fisher covariates and out-of-sample data (pg. 50)

**S2 Fig. 16:** SSD predictions for focal fisher in the non-focal landscape (pg. 51)

**S2 Fig. 17:** SSF predictions for focal fisher in the non-focal landscape (pg. 52)

**S2 Fig. 18:** RSF predictions for focal fisher in the non-focal landscape (pg. 53)

**S2 Fig. 19:** Spearman rank comparisons for all case studies (pg. 54)

**References** (pg. 55)

**S1: Simulation methodology and investigation**

*Simulation*

We tested how landscape heterogeneity and the strength of animal selection for resources impacted realized distributions of animals within simulations. To simulate animal movements, we generated various 100 x 100 rasters with random variation with standard deviations of 0.1 and Gaussian random field scales of 0.001, 0.01, 0.05, 0.1, 0.2, 0.3, 0.4, 0.6, 0.8, and 0.95 using the function *simulate_data* from the prioritizr package (Hanson et al. 2024). As the scale term increases, the simulated surface becomes more and more heterogenous (S1 Fig 1). For each simulation, we generated four simulation surfaces (e.g., S1 Fig 2), representing different environmentally-distributed variables an individual may select upon. We then specified ten animals with similar direction, but different magnitudes, of selection for each of the four environmental variables. The first animal was given selection coefficients (*β*’s) of 0.05, -0.05, 0.05, and -0.05 for each of the four landscape variables. Animals 2-10 increased linearly in their strength of selection, such that animal 10 had *β*’s 0.5, -0.5, 0.5, and -0.5. The specific values of selection coefficients are provided in S1 Table 1. Selection coefficients were then multiplied against respective simulation rasters, and rasters were added together to generate a raster of theoretical selection (S1 Fig 3).

Then, we used a movement simulation framework from Potts and Borger (2023) to generate theoretical animal movements on the randomly generated landscapes. We defined all simulations given movement limitation (*λ* in Potts and Borger’s implementation) of 1, meaning that locations further away were less likely to be moved to. We also defined a turn angle consistency (*κ* in Potts and Borger’s implementation) of 1, meaning that successive steps were more likely to maintain consistent direction. Though our method does not explicitly address turn angles, we chose include turn angle consistency in simulations for possibly greater realism. In this movement framework, an animal was placed on an initial cell in the landscape. We determined start locations by randomly selecting a location from the selection kernel over the generated landscape. Then, successive steps are made by calculating the distance between cells to generate selection decay as a function of distance (based on *λ*), turn angle consistency relative to past movements (based on *κ*), and the selection for raster cells based on the environmental dimensions and individual animal selection *β*’s. Readers interested in the mechanics and excellent extensions of this simulation framework are directed to Potts and Borger (2023).

Each animal (1-10) in each of the 10 different landscapes with varying degrees of heterogeneity was allowed to generate simulated movements for a total of 10,000 steps. As out-of-sample comparisons, we then simulated 500,000 steps with random restarts every 1,000 steps. These random restarts were initialized on the landscape based on the selection kernel. This effectively generated hundreds of “individuals” exploring different parts of simulated landscapes than our focal individual, potentially starting in or reaching different patches than were reachable by the focal individual given movement constraints. We repeated this process five times per combination of animal and landscape heterogeneity using newly generated landscapes in each iteration. In total, this generated 500 simulations of selection strength and landscape heterogeneity, with 10,000 steps in-sample (S1 Fig 4) and 50,000 steps out-of-sample. Out of sample steps were then aggregated per raster cell (*n* = 10,000) to generate an estimate of the long-term probability of use across the landscape. We also retained the selection kernel based on known animal selection strengths and randomly generated landscapes as a comparator.

*Fitting models and making predictions*

We then prepared the data for step-selection functions (SSFs, conditional logistic regression) and resource selection analyses (RSFs, logistic regression) by generating random “unused” points. We converted the dataset of 10,000 steps into a movement track, jittered locations very slightly (with uniform random noise -0.001 to 0.001 in both x and y dimensions) to create a continuous distribution of step lengths, and sampled 40 random steps per observed step using the “amt” package (Signer et al. 2019). We converted the observed distribution of step lengths into a wider uniform distribution; specifically, we decreased the minimum to 0 and increased the maximum to be 25% larger than the maximum observed step length. This increased random sampling of step distances that were both very close and very far based on the animal mover. Accordingly, we fit an SSF model treating the step-terminus values of the four input rasters as predictors in the SSF, and we included step length and log-step length as a predictor using the “amt” package (S1 Table 2) (Signer et al. 2019). For simplicity, in all analyses below, we did not model turn angle in all SSF models. We used the fitted step-selection model to parametrize the transition matrix, *A*, from instances of selection with a movement neighborhood (e.g., S1 Figs 5-6). We then generated a vector of stable-state distributions (SSDs) as described in the main text (e.g., S1 Fig 7). We also generated a landscape prediction based only on the SSF model, using a kernel of exponentiated selection strengths as a comparator (e.g., S1 Fig 8). For RSF models, we generated random points with a minimum convex polygon of use locations with 10 random “unused” points per use location. We then fit a logistic regression predicting use based on an additive combination of all for environmental layers using the “amt” package (S1 Table 2) (Signer et al. 2019), and generated predictions of probability of use across the randomly generated landscapes (e.g., S1 Fig 9).

*Inference comparisons*

For both SSF and RSF models, we were interested in how well the two modeling frameworks captured true patterns of individual selection across various levels of heterogeneity. From models, we generated the mean estimated selection for (or against) environmental layers and the 95% confidence around these estimates. We then compared (1) whether the true *β* selection coefficient was contained in this interval and (2) the absolute error between the estimated and true *β*’s. We found that, as environmental heterogeneity increased (determined by the Gaussian random fields scale term), the number of true *β*’s contained within estimated 95% CIs tended to decrease (S2 Fig 10). However, the effect of heterogeneity on bias in model-inferred selection appeared to be greater for RSF models than for SSF models (S1 Fig 10). The strength of individual resource selection (determined by the magnitude of true *β*’s) appeared to shape bias differently that landscape heterogeneity, with RSF models decreasing greatly in their ability to capture true selection patterns as individual selection strength magnitude increased (S1 Fig 10). Oppositely, SSF models, particularly in landscapes with greater heterogeneity, tended to decrease in bias as individual selection strength magnitude increased (S1 Fig 10). When investigating the absolute error (e.g., adjusted for direction of true *β*’s), we found concordant patterns: under RSF models, bias increased both as a function of individual selection strength and as a function of landscape heterogeneity (S1 Fig 11). Under SSF models, landscape heterogeneity only slightly increased overall variation in *β* estimates, and bias, particularly for specialist individuals, tended to decrease as a function of both heterogeneity and individual selection strength (S1 Fig 11).

*Prediction comparisons*

Next, we tested the ability of SSDs, SSFs, and RSFs to accurately predict animal space use over a range of landscape heterogeneity and animal selection strengths. For comparisons, we tested these surfaces against the observed probability of use based on out of sample movements as well as the true selection kernels generated only from the *β*’s of individual animals and simulated landscapes. Visually, SSD, SSF, and RSF predictions followed similar patterns as both the observed use and true selection-only kernels; however, the intensity of these distributions varied widely across different prediction frameworks, animal selection strengths, and landscape heterogeneities (S1 Fig 12). In general, the observed probability of use tended to have higher maxima in areas of high probability of use and lower minima in areas of lower probability of use as compared to the true selection-only kernels. In line with these visual inferences, the median prediction probability of use was uniformly lower for the observed use than the selection kernel, and the variation in predictions was uniformly higher for observed use as compared to selection kernels (S1 Fig 13). As individuals exerted stronger selection strengths and as the environment became more heterogeneous, observed use patterns became more heterogeneous (S1 Fig 13).

To quantify these patterns of heterogeneity, we investigated the Shannon entropy of individual predictions, calculated as: $\sum_{i=1}^{n} p\left( x_{i} \right)\ln(p\left( x_{i} \right))$. As Shannon entropy decreases, probability kernels imply more certain outcomes. In this context, lower Shannon entropy describes predictions that the intensity of animal use was concentrated in fewer locations. As animals became choosier and as landscapes became more heterogeneous, the Shannon entropy of prediction kernels tended to decrease. The observed probability of use was uniformly more heterogeneous than either the true selection kernel or the SSF-inferred selection kernel (S1 Fig 14). For landscapes that are less heterogenous, the observed probability of use was also always more heterogenous than RSF-inferred or SSD probability of use (S1 Fig 14). However, as landscape heterogeneity increases, both SSD and RSF predictions more closely align with the observed use heterogeneity. In some instances, the SSD was more heterogeneous than the observed use (S1 Fig 14).

We also examined the outcomes of predictions in the context of usable spatial extents. To do this we calculated the cumulative distribution of probability of use, ordered from highest probabilities to lowest. Arbitrarily, we then identified all raster cells that were contained in the first set whose members summed to equal to or less than 0.5. This defines the minimum number of locations on the landscape where the probability of use sums to 50% (S1 Fig 15). When aggregated, it was clear that the minimal area in the top half of the probability kernel was often grossly overestimated by the true selection kernel or SSF-inferred selection kernel (S1 Fig 16). In cases of strong selection by animals in high heterogeneity landscapes, it was evident that these selection-only methods overestimated the top-use area by up to 150%. RSF-inferred top-use area was often intermediate, such that RSF almost always overestimated area but not the extent of SSF (S1 Fig 16). Interestingly, it appeared that animal choosiness, not heterogeneity, was the chief determinant of area overestimation (S1 Fig 16). Finally, in low heterogeneity landscapes, SSD uniformly overpredicted top area use; however, SSD was the closest estimate to observed area use in landscapes with moderate to high heterogeneity (S1 Fig 16).

We then examined the accuracy of SSD, SSF, and RSF predictions compared against the observed probability of use. To compare these observed kernels against observed use, we used three main heuristics: (1) relative entropy, or Kullback–Leibler divergence, measured as $\sum_{i=1}^{n} p\left( x_{observed} \right)\ln(\frac{{p(x}_{observed})+{1\times10}^{-10}}{{p(x}_{predicted})+{1\times10}^{-10}})$ to ensure that unused sites were included in calculations (S1 Fig 17); (2) root-mean squared error (RMSE, S1 Fig 18); and (3) top use area classification accuracy, defined as the minimal area whose sum probability of use was equal to or less than 0.5 (S1 Figs 19-20). Overall, these comparisons revealed largely concordant findings: in cases where landscape heterogeneity was low, SSD tended to perform equally as well or worse than SSF or RSF in predicting observed use. As landscape heterogeneity increased, however, SSD tended to outperform SSF and RSF predictions (S1 Figs 17-19). As animal selection strength increased, predictions tended to become more dissimilar from observed probability use under relative entropy and RMSE (S1 Figs 17-18), though not for classification accuracy (S1 Fig 19). Moreover, individual selection strength appeared to have relatively little effect on the tradeoff between SSD and SSF or RSF (S1 Figs 17-19).

*Prediction comparisons with imperfect validation data*

Finally, we sought to better understand the accuracy of predictions when the true kernel of use probability was unknown or partially measured. In simulations, we allowed simulations to run for 1,000 iterations 500 times, but this does not necessarily mean that any of those animal sampling processes reached a true steady-state equilibrium. We felt that this provided a balance of total landscape sampling (though many quasi-independent sampling processes), and we also believed this to more faithfully describe the types of validation data potentially available in real-world scenarios. To explore relationships between kernel predictions and imperfect validation data, there are many possible comparison metrics. First, we could simply assume that the observed use intensity of cells should be linearly related to the prediction kernels, specifying that observed use intensity should be directly correlated to predictions. In such cases, Pearson correlation coefficients may provide a reasonable comparison tool (S1 Fig 21). Second, we might assume that observed use intensity is an imperfect sampling of real use, such that use intensity is related to predicted use, but not linearly so. This concept underlies common validation frameworks stemming from Boyce et al. (2002) calculated using Spearman rank correlations between binned prediction surfaces (either in equal area or equal intervals) and observed use intensity within bins across a number of cross validations. To capture this concept of nonlinearity between observed use intensity and prediction kernels, we simply calculate Spearman rank between these distributions (S1 Fig 22). Finally, as all prediction kernels summed to one, one way to compare prediction surfaces would be to compare the predicted probability of use as each use location: use locations classified by higher use probability between prediction surfaces may then discriminate more accurate predictions. Given the heterogeneity of surfaces, we found that it was necessary to use the geometric mean of predicted probability of use. In effect, we calculated the weighted mean log-probability of use across prediction surfaces for all use locations and then exponentiated this average back to the probability scale (S1 Fig 23).

Comparisons using Pearson, Spearman, and geometric mean of predicted use appeared to be more conservative regarding differences between performance of different predictions (S1 Figs 21-23) than comparison heuristics which assume that probability of use is known (e.g., relative entropy, RMSE S1 Figs 21-23). Importantly, the geometric mean probability of use appeared to be the most conservative of the comparison heuristics (S1 Figs 23). SSD predictions were nearly always favored over SSF or RSF predictions, and SSF was favored over RSF at very low levels of animal selection strength and higher levels of landscape heterogeneity (S1 Figs 21-23). Spearman rank correlations appeared to slightly favor SSF predictions relative to RSF, perhaps because of a slight to favor predictions with lower levels of heterogeneity though rank-based, rather than linear, relationships. This pattern was confirmed by plotting observed use intensity against prediction kernels: SSF tended to have a less variable relationship with observed use, but the relationship was biased off a one-to-one relationship (S1 Fig 24). SSD tended to have more variation in the predicted surface, but had a less-biased relationship with observed use, particularly for high levels of landscape heterogeneity or animal selection strength (S1 Fig 24).

**S1 Table 1: Animal selection strength coefficients**

| **Animal** | ***β_1_*** | ***β_2_*** | ***β_3_*** | ***β_4_*** |
| --- | --- | --- | --- | --- |
| **1** | 0.05 | -0.05 | 0.05 | -0.05 |
| **2** | 0.1 | -0.1 | 0.1 | -0.1 |
| **3** | 0.15 | -0.15 | 0.15 | -0.15 |
| **4** | 0.2 | -0.2 | 0.2 | -0.2 |
| **5** | 0.25 | 0.25 | 0.25 | -0.25 |
| **6** | 0.3 | -0.3 | 0.3 | -0.3 |
| **7** | 0.35 | -0.35 | 0.35 | -0.35 |
| **8** | 0.4 | -0.4 | 0.4 | -0.4 |
| **9** | 0.45 | -0.45 | 0.45 | -0.45 |
| **10** | 0.5 | -0.5 | 0.5 | -0.5 |

*β coefficients denote selection direction and magnitude for four independently simulated landscape variables for ten different animals within simulations*

**S1 Table 2: Model pseudocode for the simulation**

| Dataset | Model pseudocode (SSF) | Model pseudocode (RSF) |
| --- | --- | --- |
| Simulation | z.1 + z.2 + z.3 + z.4 + poly(sl_,2) + strata(step_id) | z.1 + z.2 + z.3 + z.4 |

*z.1, z.2, z.3, and z.4 are randomly generated rasters (100 x 100 units)*

*sl_ is step length in (arbitrary) simulation raster units*

*step_id is the unique identifier of an observed step and the 40 randomly sampled points for that same step*

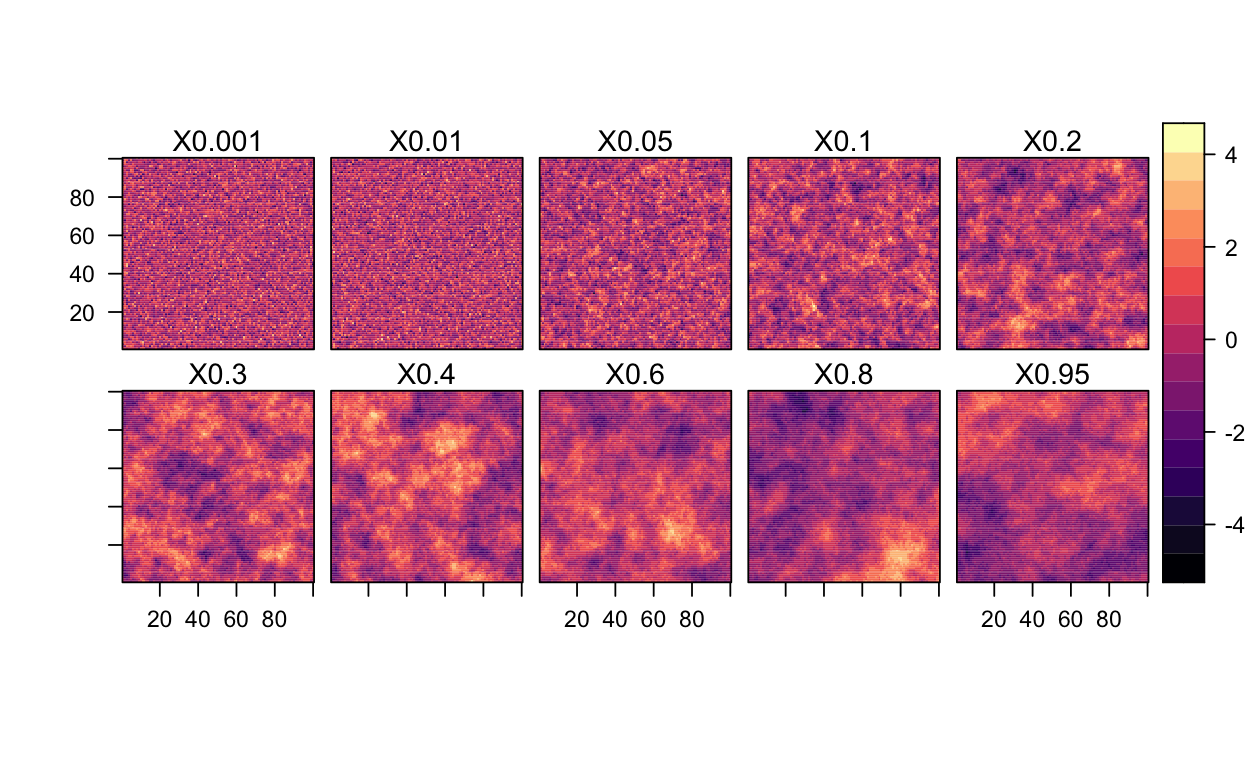

**S1 Fig 1: Example randomly generated rasters with Gaussian scale from 0.001 to 0.95**

These example rasters are parametrized with Gaussian random field scales used in the simulation, labeled by X followed by the scale value. Here, these rasters are visualized in their raw form before being combined by a set of random coefficients into a surface of suitability.

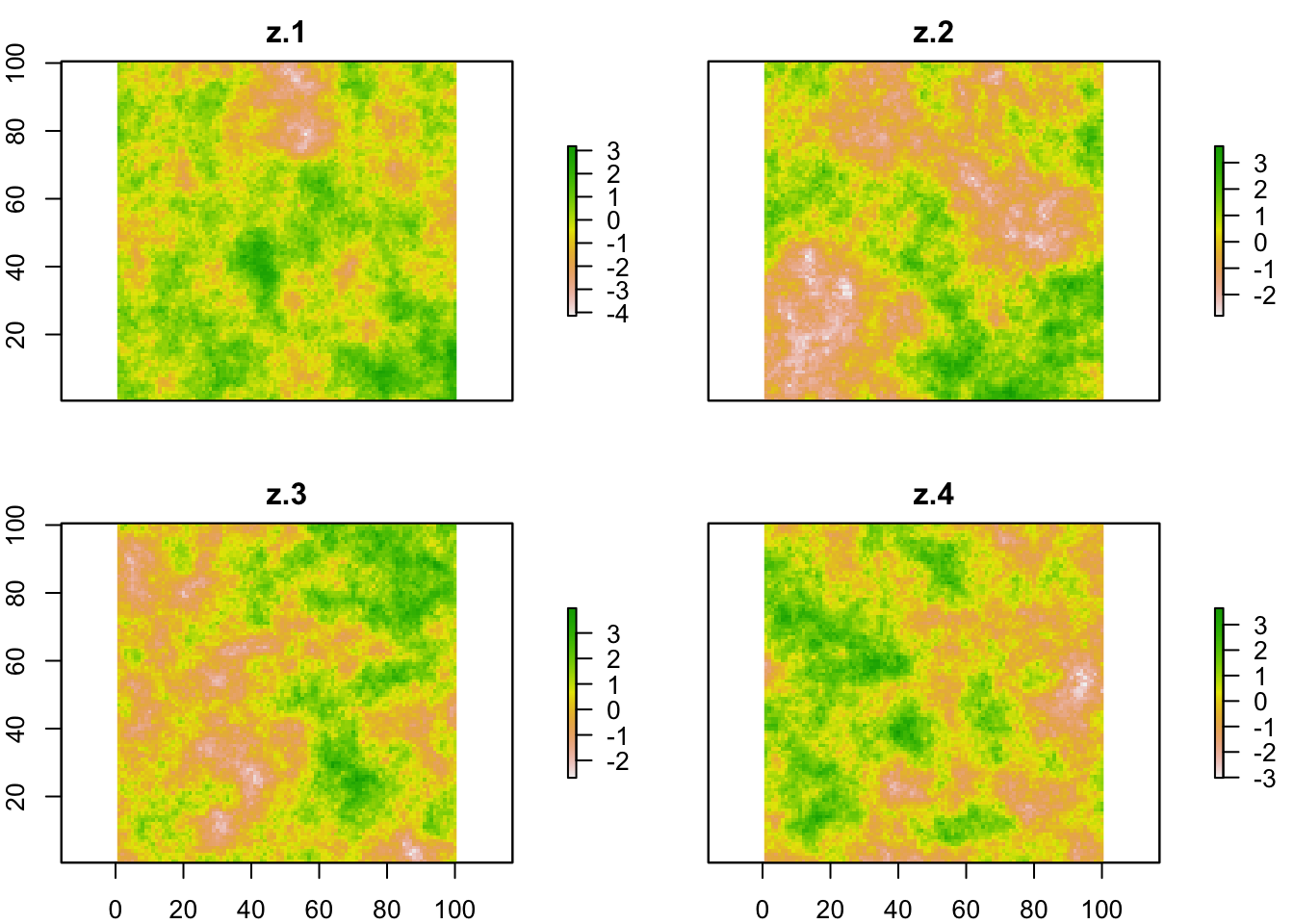

**S1 Fig 2: Example of four simulation rasters used per iteration**

The four rasters used in the simulation, labeled z.1-z.4. Here, these rasters are visualized in their raw form (log-scale), before being combined by a set of random coefficients into a surface of suitability.

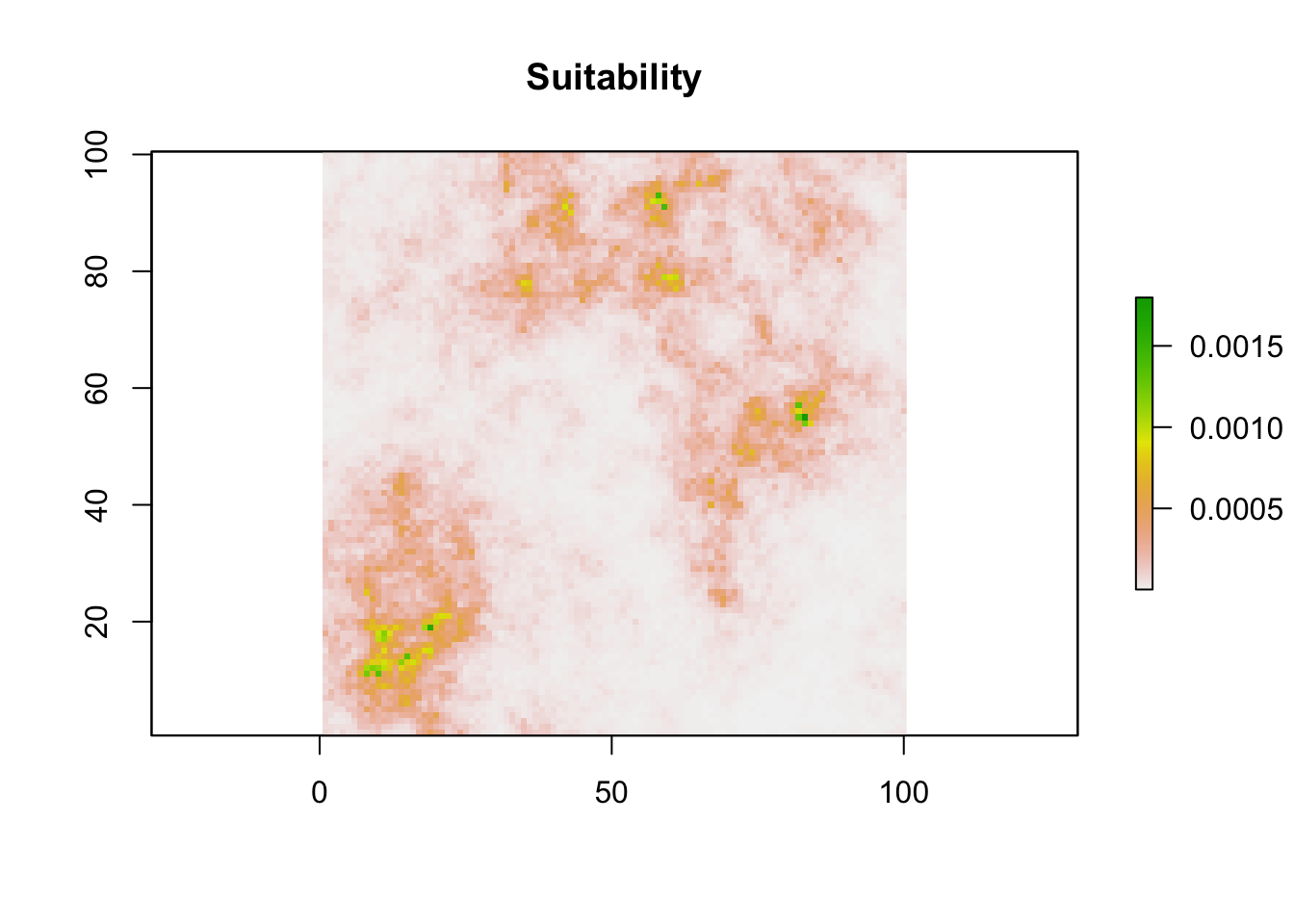

**S1 Fig 3: Simulated environmental suitability**

Habitat suitability on “probability” scale, combining the landscape rasters (S1 Fig 1) using randomly drawn coefficients (e.g., -0.3844678, -0.4846550, 0.1046449, -0.8872337) prior to exponentiating the surface and calculating the kernel of the surface.

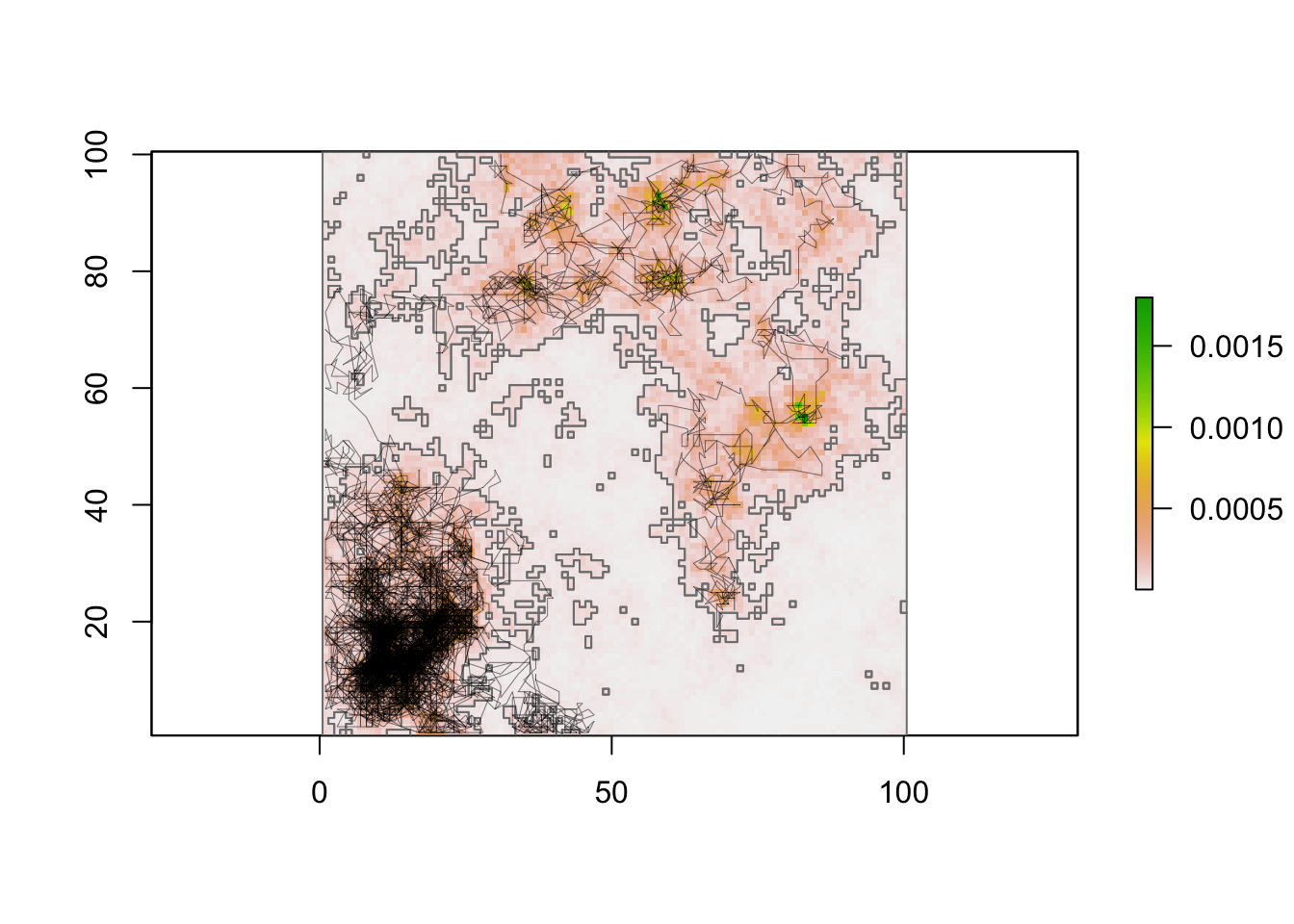

**S1 Fig 4: Simulation tracks over environmental suitability**

10,000 simulated steps based on suitability surface and an exponential decay in probability based on step length and increase in probability of with similar turn angles.

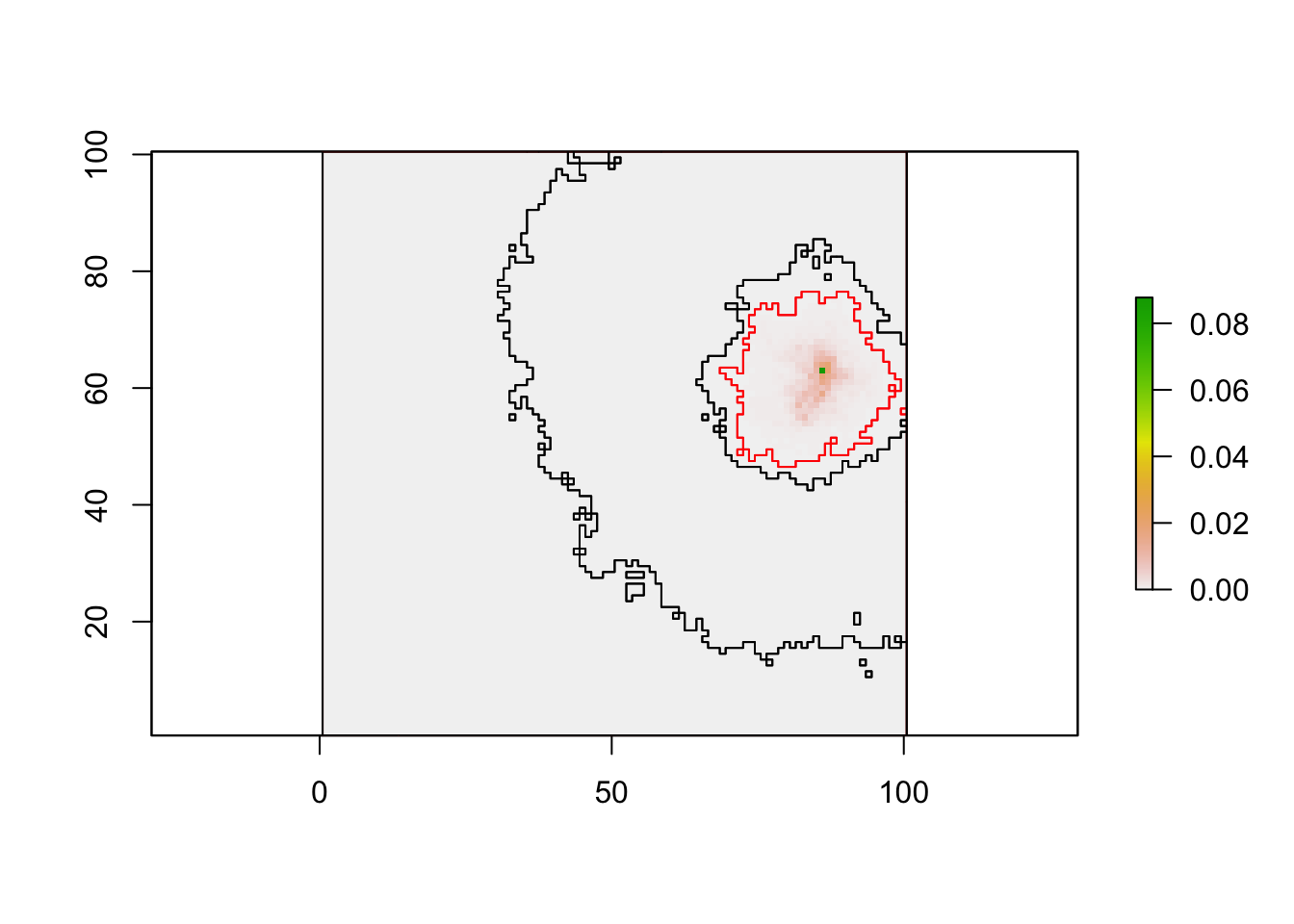

**S1 Fig 6: Visualizing the neighborhood of *G_i_***

Random context (*G_i_*) in black circle, with local movement probabilities across the entire surface. Within the red polygon are all locations with a movement probability greater than 0.0001. Within the smaller of the black polygons are all locations with greater probabilities of movement than 90% of the surface. Within the larger black polygon are all locations with greater probabilities of movement than 50% of the surface. This visualization clearly demonstrates how the movement kernel restricts “possible” movements, enabling the use of sparse matrices in latter computational steps.

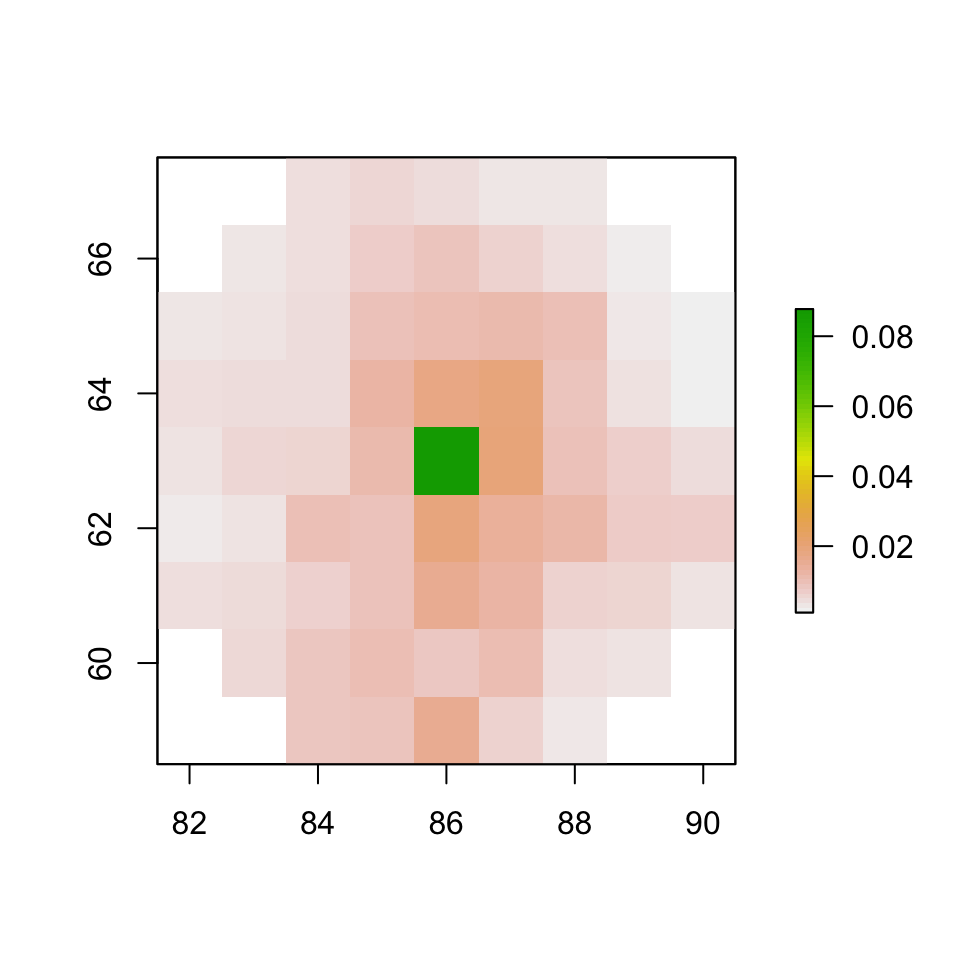

**S1 Fig 7: Visualizing the neighborhood of *G_i_*** **at a proximate scale**

Zooming in to a given context (*G_i_*) and the possible movement choices in its neighborhood (95% percentile of observed step lengths). There was a clear down-weight of remaining in the same cell, but also a clear down-weight of moving very far. Habitat heterogeneity around the center, however, shows variation in patterns of predicted movement within reasonable movement contexts.

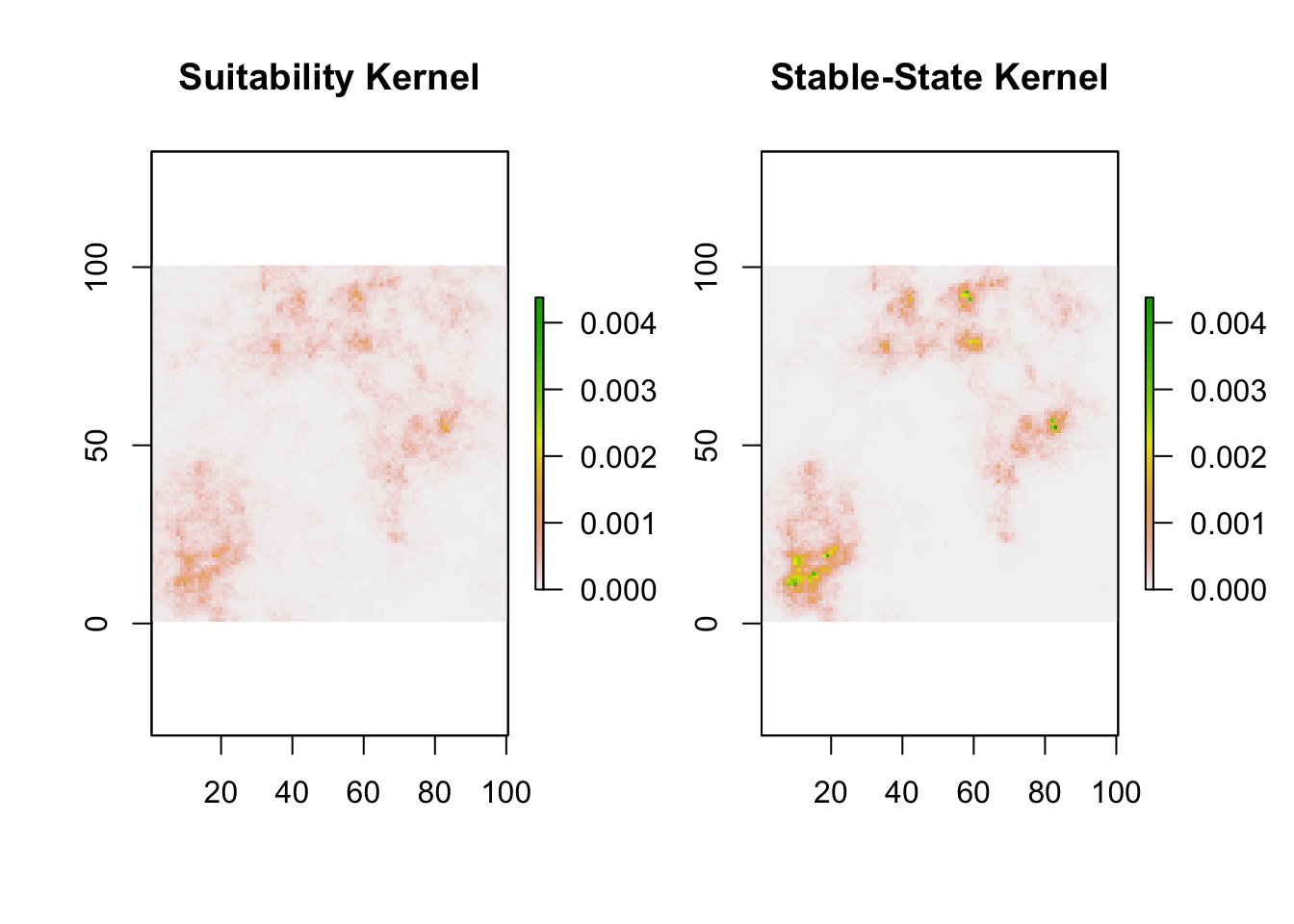

**S1 Fig 8: Comparison of selection and stable-state kernels from simulation example**

Stable state distribution (SSD) vs true selection kernel plotting. The comparison here uses the same color scale for both predictions, highlighting their departure from each other. Both kernels distributions are qualitatively similar, but their intensity was very different.

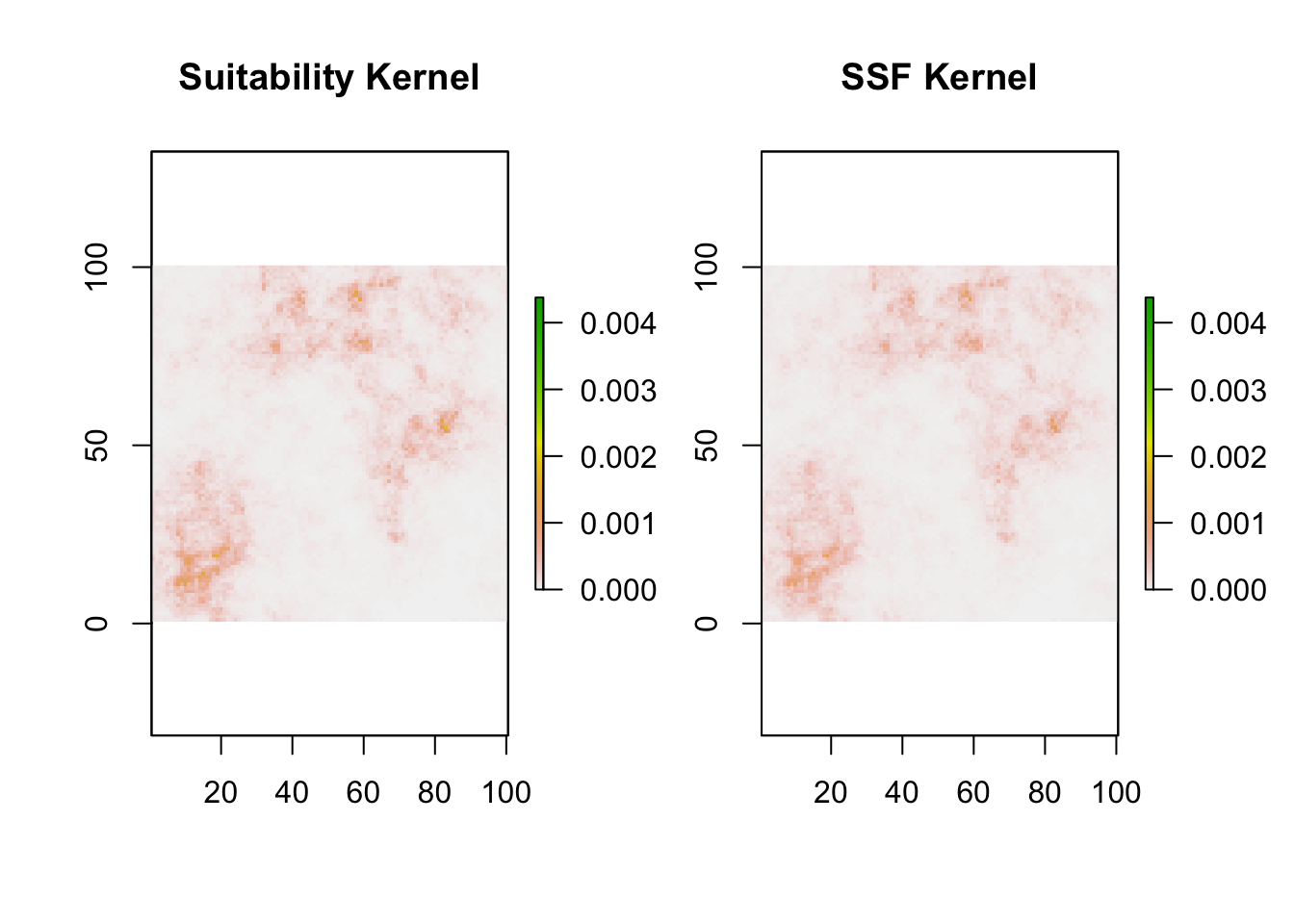

**S1 Fig 9: Comparison of true selection and step-selection-inferred selection kernels from simulation example**

Step selection function (SSF) vs true selection kernel plotting. The comparison here uses the same color scale for both predictions, highlighting their concordance. Both kernels distributions are qualitatively similar, and their intensity was also similar. In fact, these surfaces are nearly perfectly related (as the SSF was nearly perfectly recovering true selection patterns).

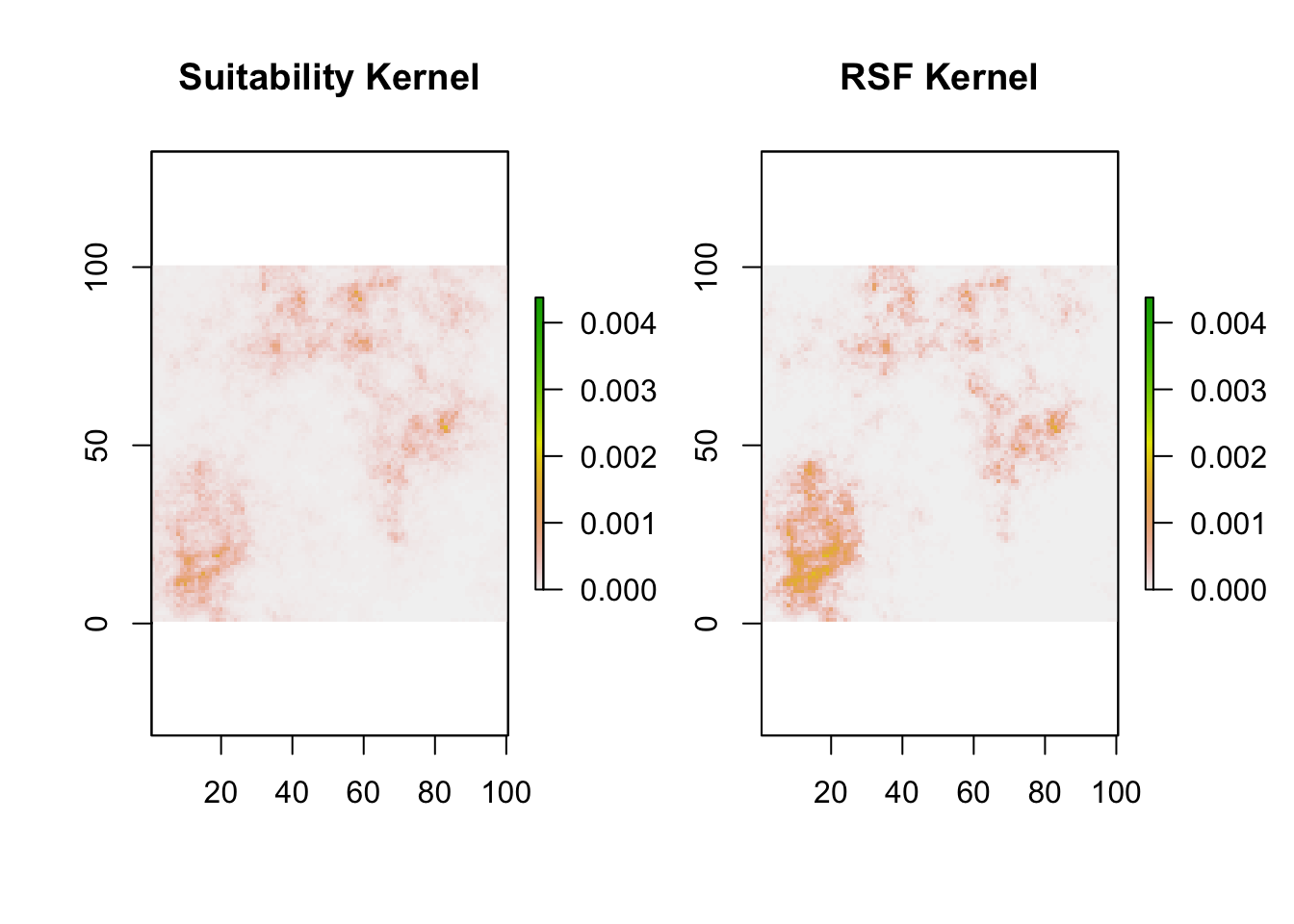

**S1 Fig 10: Comparison of true selection and resource selection-inferred probability of use kernels from simulation example**

Resource selection function (RSF) vs true selection kernel plotting. The comparison here uses the same color scale for both predictions. Both kernels distributions are qualitatively similar, and their intensity was somewhat similar.

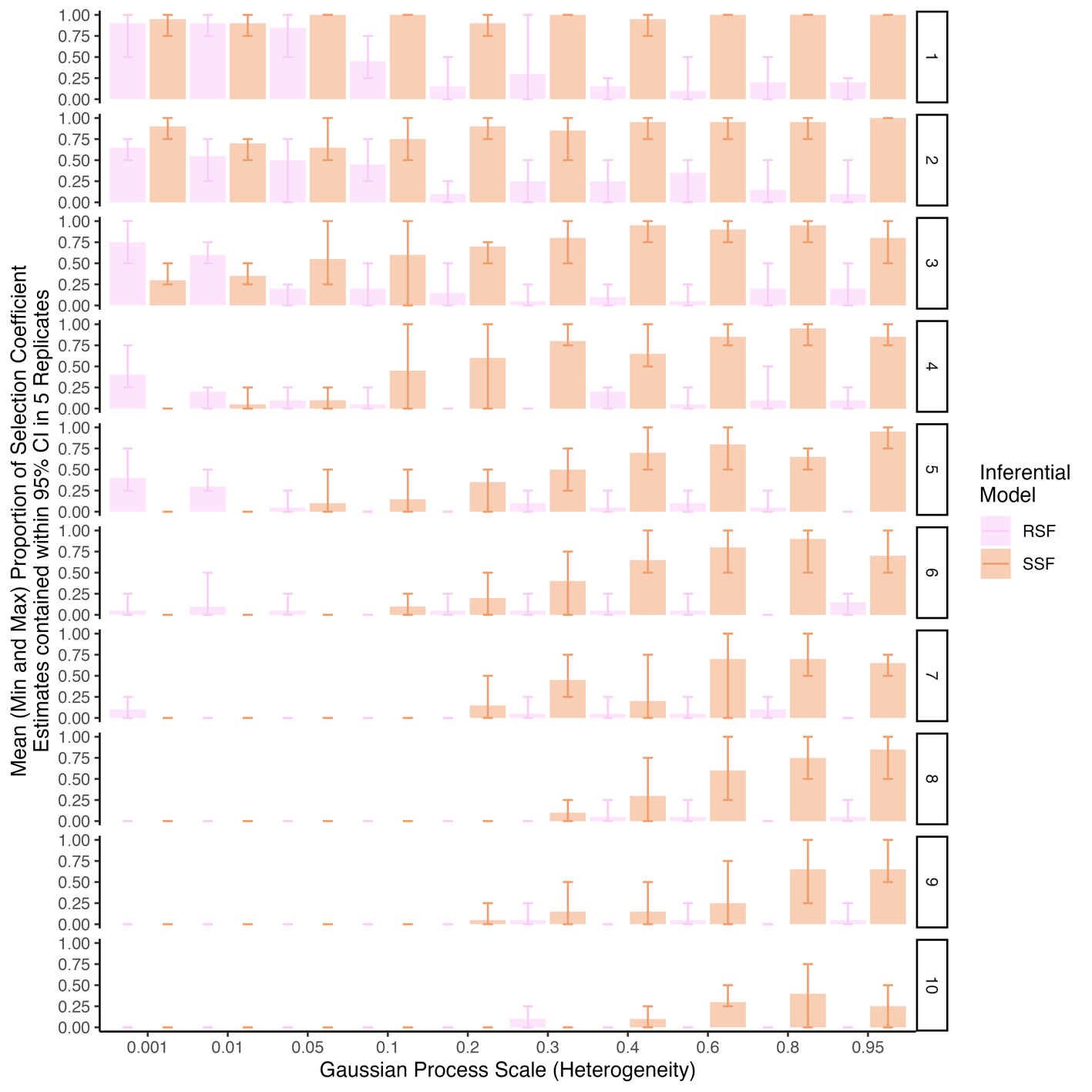

**S1 Fig 11: Proportion of selection coefficients contained in estimate CI by model type, heterogeneity, and animal selection strength**

In simulations (*n* = 500), animals with increasing selection strengths (*n* = 10) were allowed to move on landscapes with different heterogeneity (*n* = 10) in five independent iterations. From these movements, SSFs and RSFs were fit. The number of true selection coefficients contained in 95% confidence intervals (CIs) we tallied. The mean, min, and max proportion of coefficients contained are plotted by inferential model type.

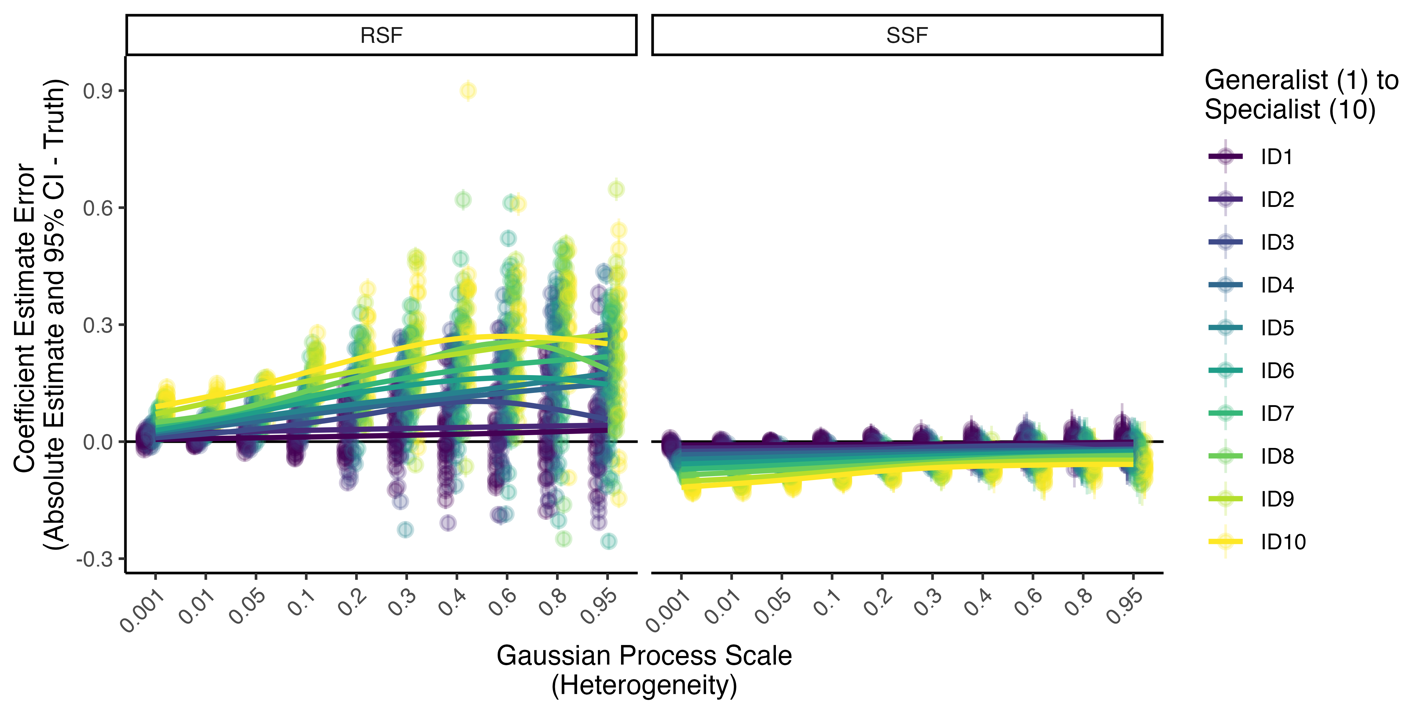

**S1 Fig 12: Absolute error for selection coefficients by model type, heterogeneity, and animal selection strength**

True selection coefficients were subtracted from SSF and RSF estimates and 95% CIs of selection coefficients after converting the direction of all selection coefficients to be in the same direction. This absolute error therefore describes more extreme (positive) and less extreme or reverse directionality (negative) of model inferred selection vs. truth. SSF inferences suffer less from overall changes in animal selection strength or heterogeneity than RSF inferences.

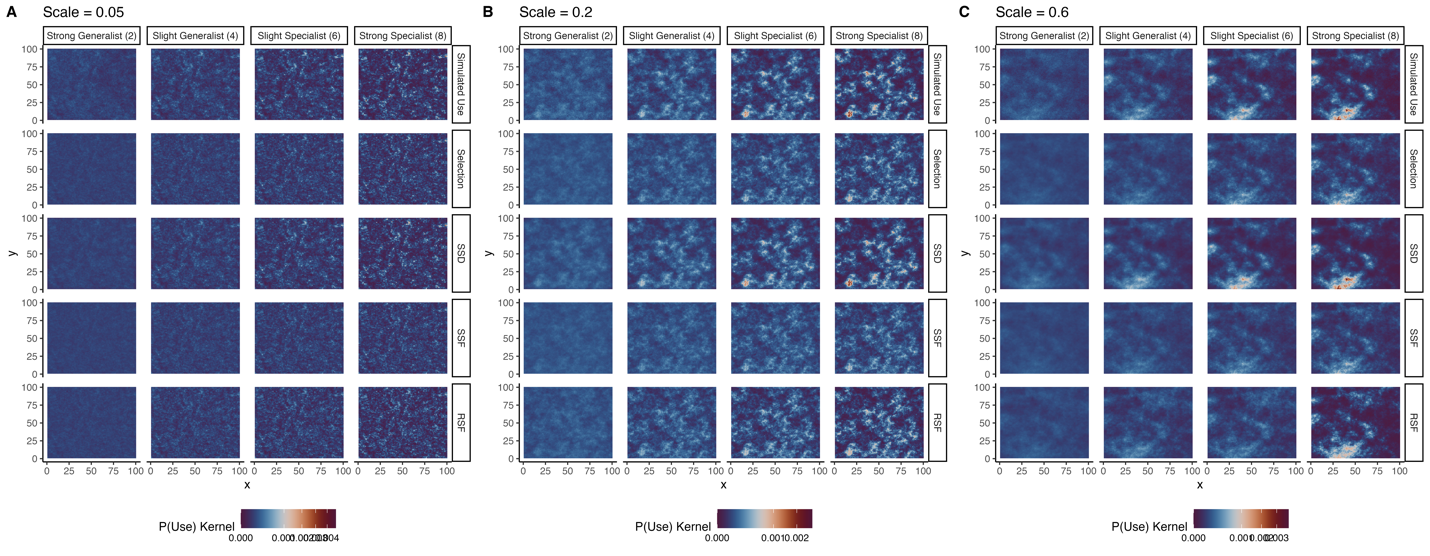

**S1 Fig 13: Landscape predictions (SSD, SSF, RSF) vs true selection and observed use kernels**

In simulations, long-term observed use (in 500,000 steps restarting every 1,000 steps) was tallied per raster cell to generate observed probability or intensity of use. In **A**, **B**, and **C**, a single realization of observed simulated use, the known selection kernel specified in simulations, and the three model predictions (vertical facets) are visualized at three different levels of landscape heterogeneity (0.04, 0.2, 0.6, respectively) for four different animals with increasing selection strength (2, 4, 6, 8, horizontal facets). In general, observed use and SSD predictions seem to visually be most similar, while the known selection kernel and SSF seem to be most similar. Importantly, the observed and true selection kernel are not similar in intensity, particularly at higher levels of landscape heterogeneity and animal selection strength. RSF predictions, generally, appear to fall most frequently between observed and selection kernels.

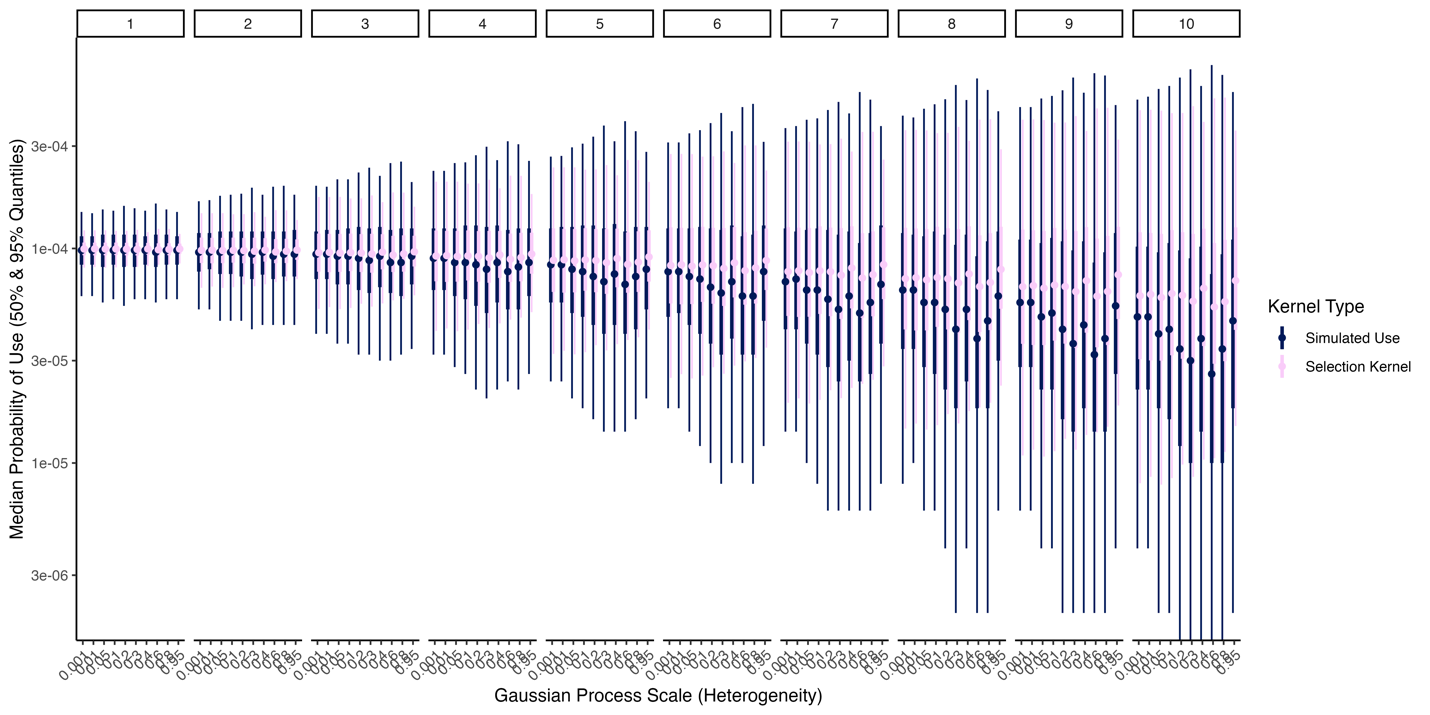

**S1 Fig 14: Variation in observed probability of use vs. known selection kernels as a function of selection strength and landscape heterogeneity**

Median and inner 50% and 95% quantiles were drawn from kernels of observed use and known selection across a single simulation per set of conditions. The simulation landscape has 10,000 cells, so at random the median probability of use should be 1x10^-4^. As animal selection strength increases, the variation in both the known selection kernel and the simulated use increase and the median probability use of a pixel in the landscape decreases dramatically. This increase in variation and decrease in median use was far greater for simulated use rather than known selection. Similarly, as landscape heterogeneity increases, observed use becomes more variable, while the selection kernel remains at roughly the same level of heterogeneity.

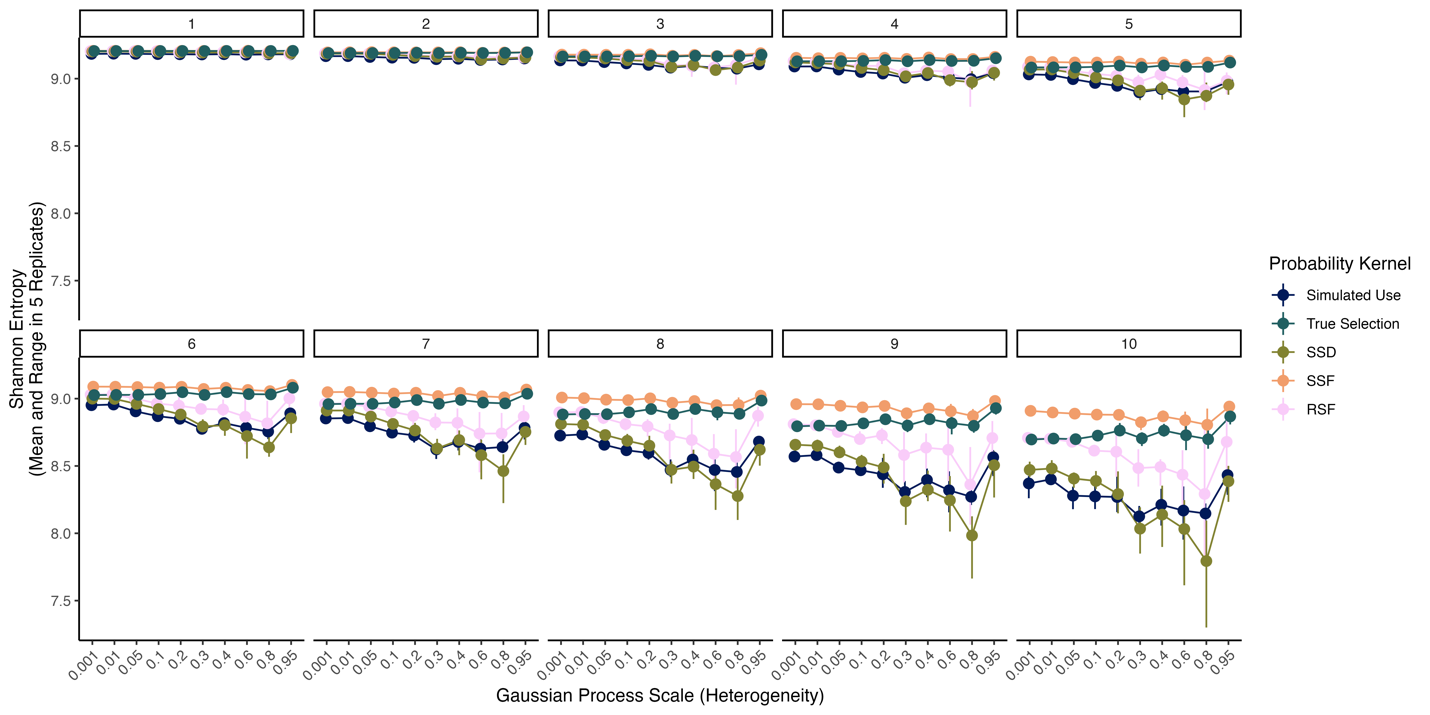

**S1 Fig 15: Mean and range of Shannon entropy for kernels across five simulations**

Shannon entropy was a measure of certainty in probability distributions, and thus describes heterogeneity in predictions of use across measures. SSD tends to more closely resemble heterogeneity in simulated use, particularly at higher levels of landscape heterogeneity.

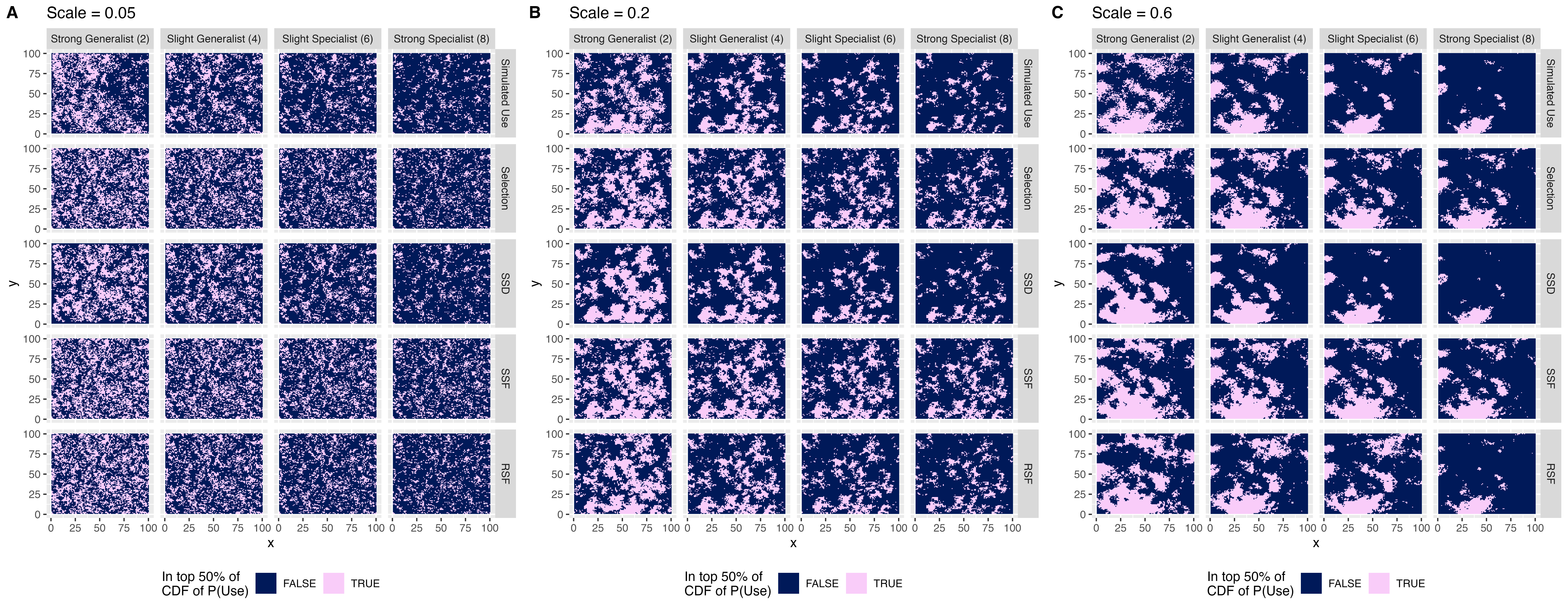

**S1 Fig 16: Landscape classification based on probability of use**

Prediction kernels were ordered and then cumulatively summed to determine whether they were in the top 0.5 of the prbability kernel – defining the minimal area required to reach a sum probability of use of 0.5. These are then plotted following S1 Fig 13 at three representative landscape scales (0.05, 0.2, 0.6) for four different animal selection profiles (2, 4, 6, 8) for all prediction, simulation, or known kernels for a single simulation per set of conditions. There are clear disparities in the distribution and patterns of top habitats across methods, with variation, particularly around the periphery of patches.

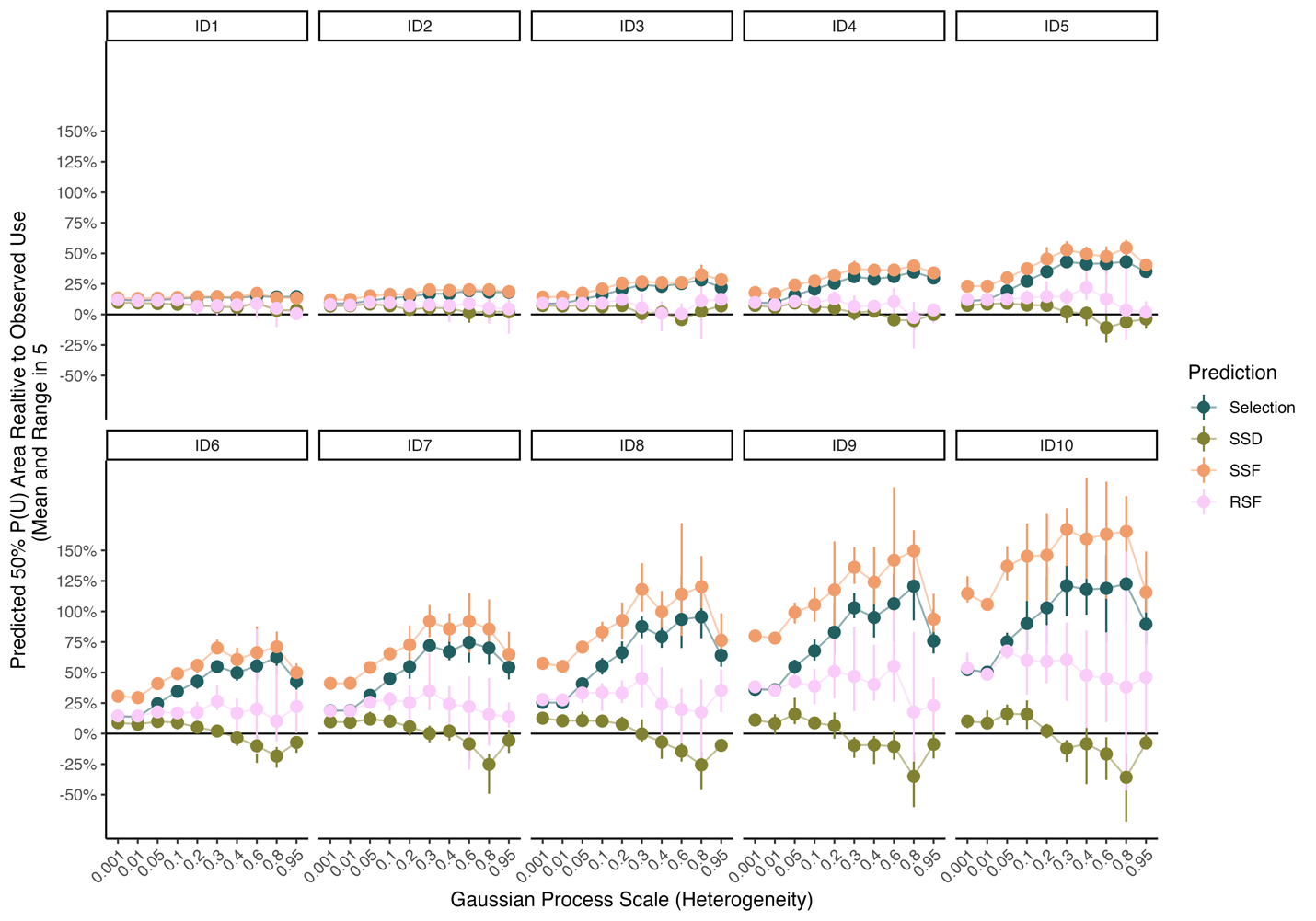

**S1 Fig 17: Total minimal area to reach a sum probability of use of 0.5 vs. simulated use across 5 simulations**

Ordering prediction and known selection kernels, we calculated the minimal are needed to total a probability of use equal to 0.5 and compared this to observed use kernels across five simulations generating a mean and range of area estimates as a proportion of observed area estimates. In all cases of low heterogeneity, predictions tended to overestimate area by 10%. However, SSD predictions are the only predictions that remain low in bias with increasing animal selection strengths (1-10). However, in more heterogeneous landscapes, SSD (and to a lesser degree RSF) much better recapitulated area-based definitions of top habitat. In contrast, relying on the selection kernel or SSF-inferred kernel led to stark overestimations of over 150%.

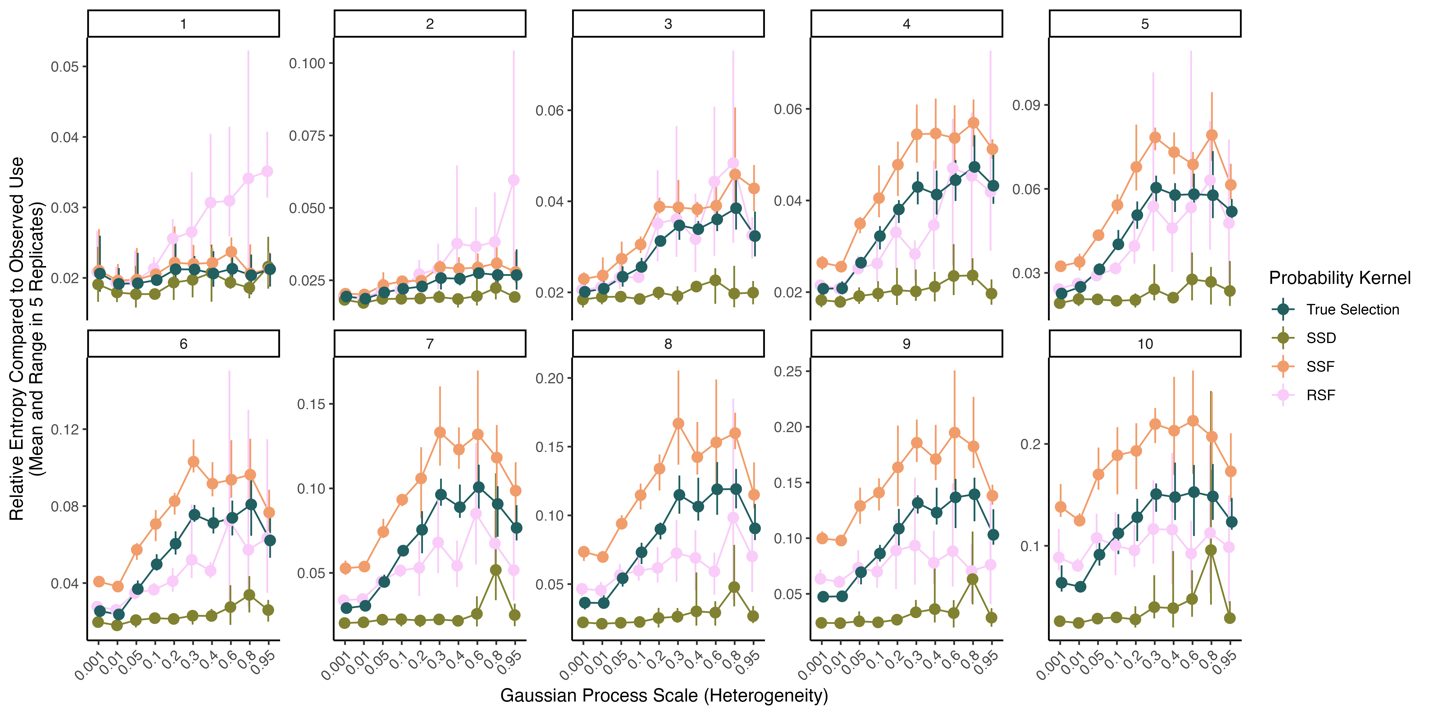

**S1 Fig 18: Relative entropy of predictions vs. simulated use across 5 simulations**

Across five simulations per set of conditions, we treated the observed use kernels as “truth” and the predictions and selection kernels as approximations of truth. We then calculated the relative entropy between these predictions and truth, generating a mean and range of relative entropies across all prediction or known selection kernels. Uniformly, SSD was more similar to the observed use distribution than were SSF or RSF predictions or the true selection kernel. SSF predictions were more similar to observed use in scenarios with weak animal selection strengths, but RSF was more similar to observed use distributions as selection strength and habitat heterogeneity increased.

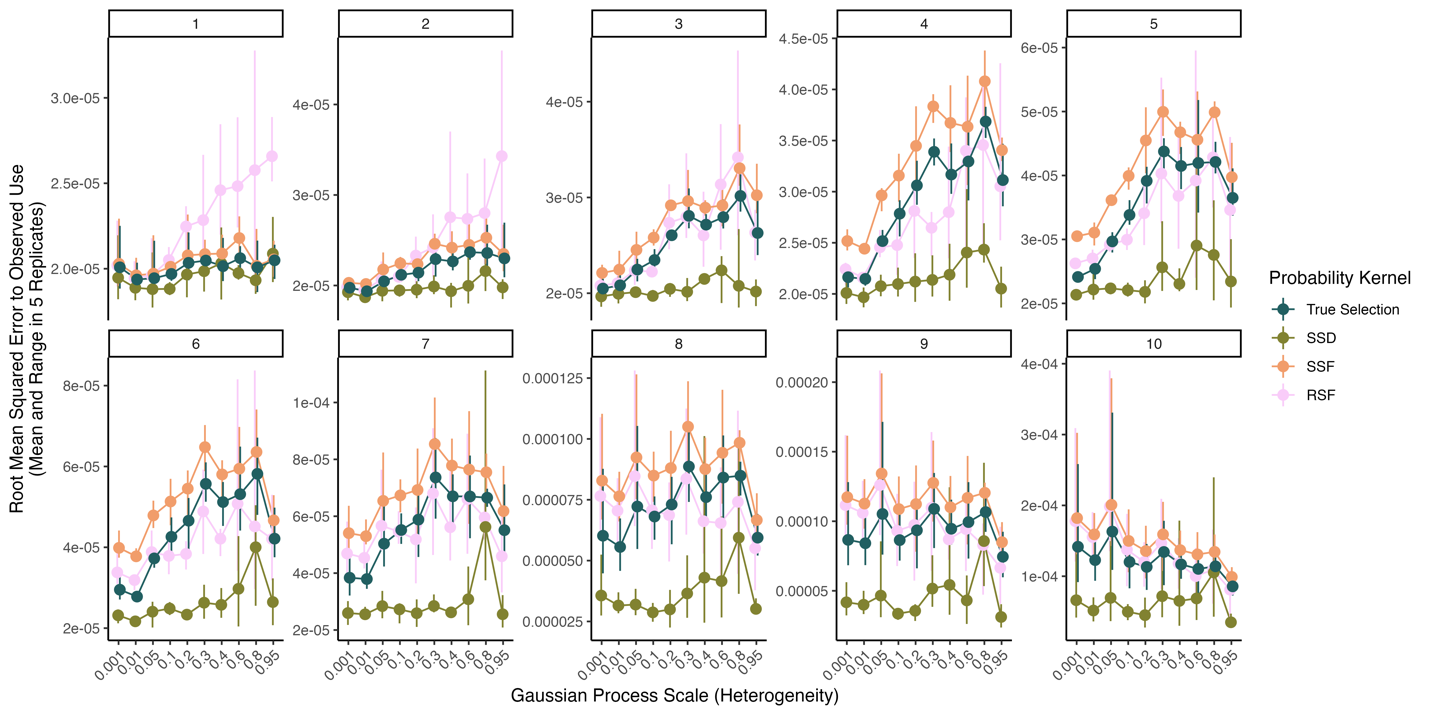

**S1 Fig 19: Root mean squared error of predictions vs. simulated use across 5 simulations**

Across five simulations per set of conditions, we calculated the root mean squared error (RMSE) between kernel predictions and observed use, generating a mean and range of RMSE across all prediction or known selection kernels. Uniformly, SSD was more similar to the observed use distribution than were SSF or RSF predictions or the true selection kernel. SSF predictions were more similar to observed use in scenarios with weak animal selection strengths, but RSF was more similar to observed use distributions as selection strength and habitat heterogeneity increased.

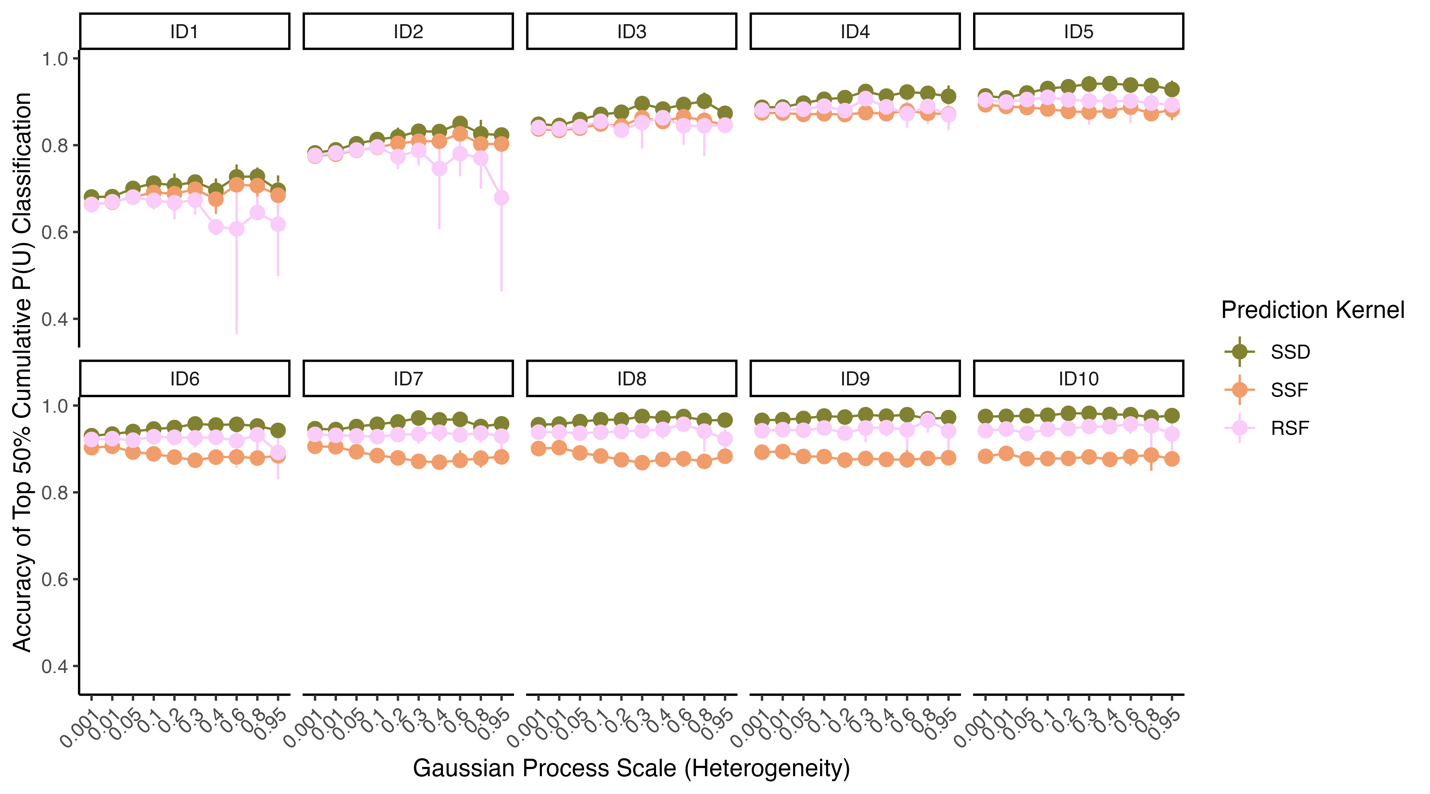

**S1 Fig 20: Accuracy of predictions in classifying top simulated use area across 5 simulations**

Across five simulations per set of conditions, we calculated the minimal area necessary to sum to total probability of use of 0.5. We then determined which raster cells in the landscape were contained in this upper distribution and compared the accuracy of defining top area use spatially vs. the observed use kernel. We generated a mean and range of accuracy across all prediction kernels. Uniformly, SSD was more accurate in classifying top use areas thank either SSF or RSF. At low levels of animal selection strength SSF was more accurate than RSF, but this pattern quickly reversed with stronger selection strength or more heterogenous habitats.

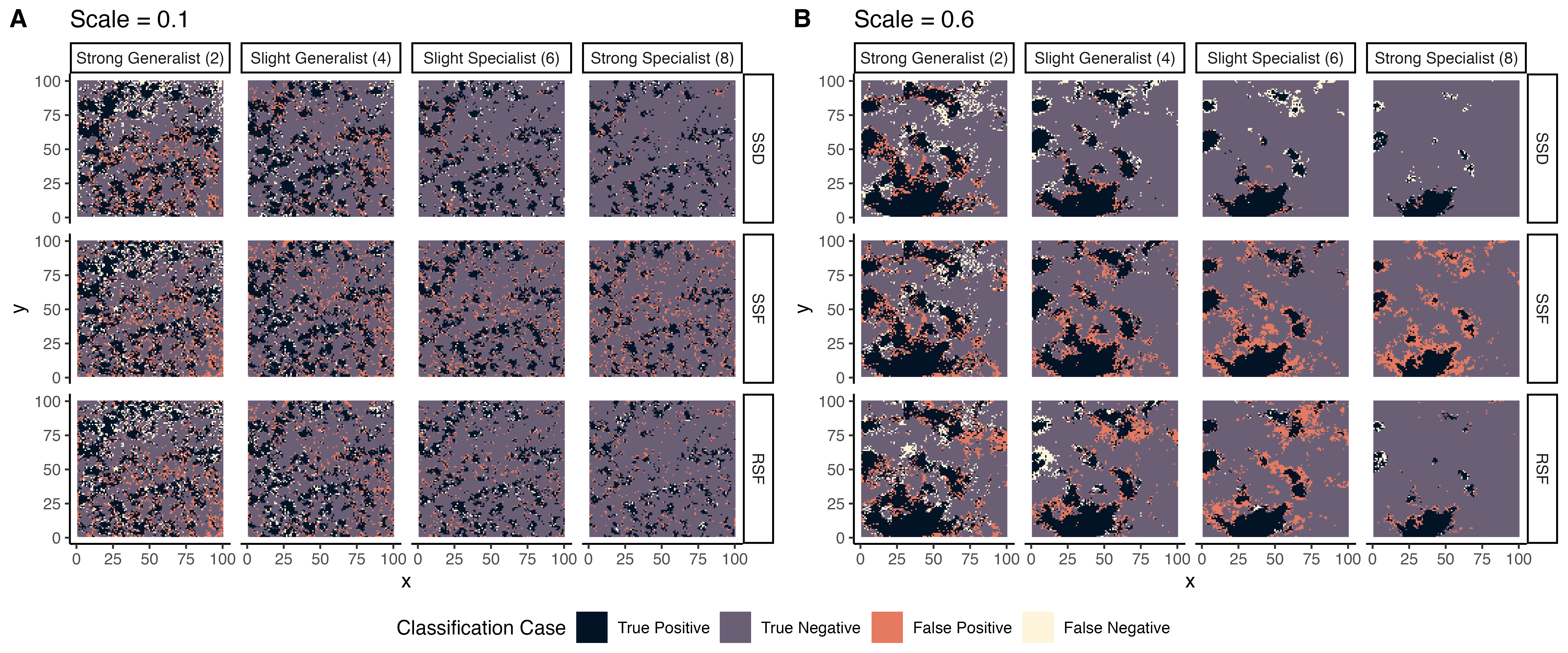

**S1 Fig 21: Classification cases of prediction kernels vs. simulated use**

For a single iteration per condition, we calculated the minimal area necessary to sum to total probability of use of 0.5. We then determined which raster cells in the landscape were contained in this upper distribution and compared the accuracy of defining top area use spatially vs. the observed use kernel. Raster cells were then attributed: “True Positive” meaning both the prediction and observed use kernels matched in attribution of top habitat; “True Negative” meaning both the prediction and observed use kernels matched in attribution of bottom habitat; “False Positive” meaning the prediction classified the cell as top habitat, while observed use classified as bottom habitat; and, “False Negative” meaning the prediction classified the cell as bottom habitat, while observed use classified as top habitat. These are then visualized for two landscapes with 0.1 or 0.6 as scale for heterogeneity for four types of animal selection profiles (2, 4, 6, 8) for SSD, SSF, and RSF predictions. There was important variation across conditions, but, in general, it appeared that SSD at high habitat heterogeneity tends to outperform SSF and RSF.

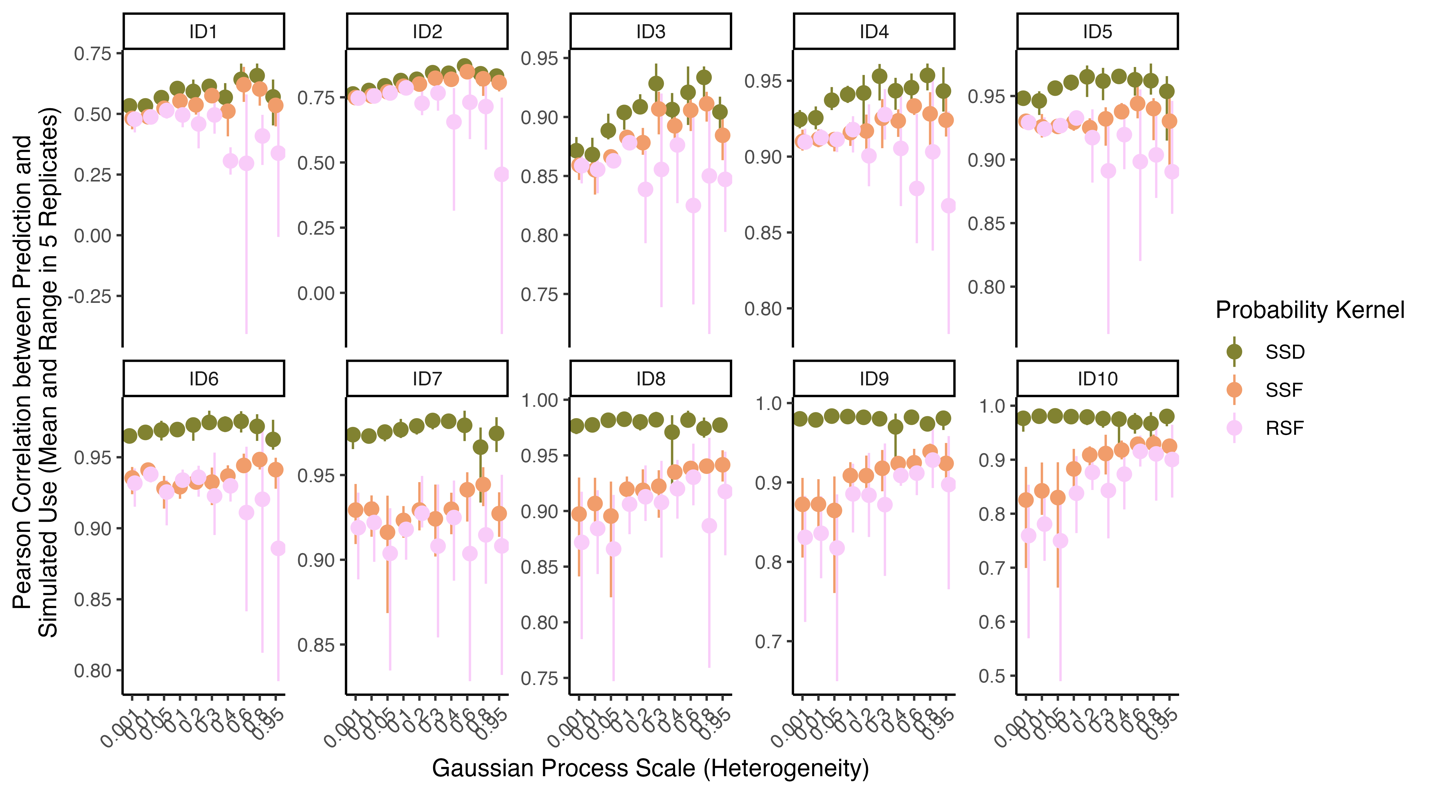

**S1 Fig 22: Pearson correlation between prediction and observed use kernels**

In five simulations per set of conditions, the Pearson correlation between simulated use kernels and SSD, SSF, and RSF kernels was calculated, generating a mean and range per set. In general, SSD was always favored, and SSF was favored over RSF at low selection strength and higher landscape heterogeneity.

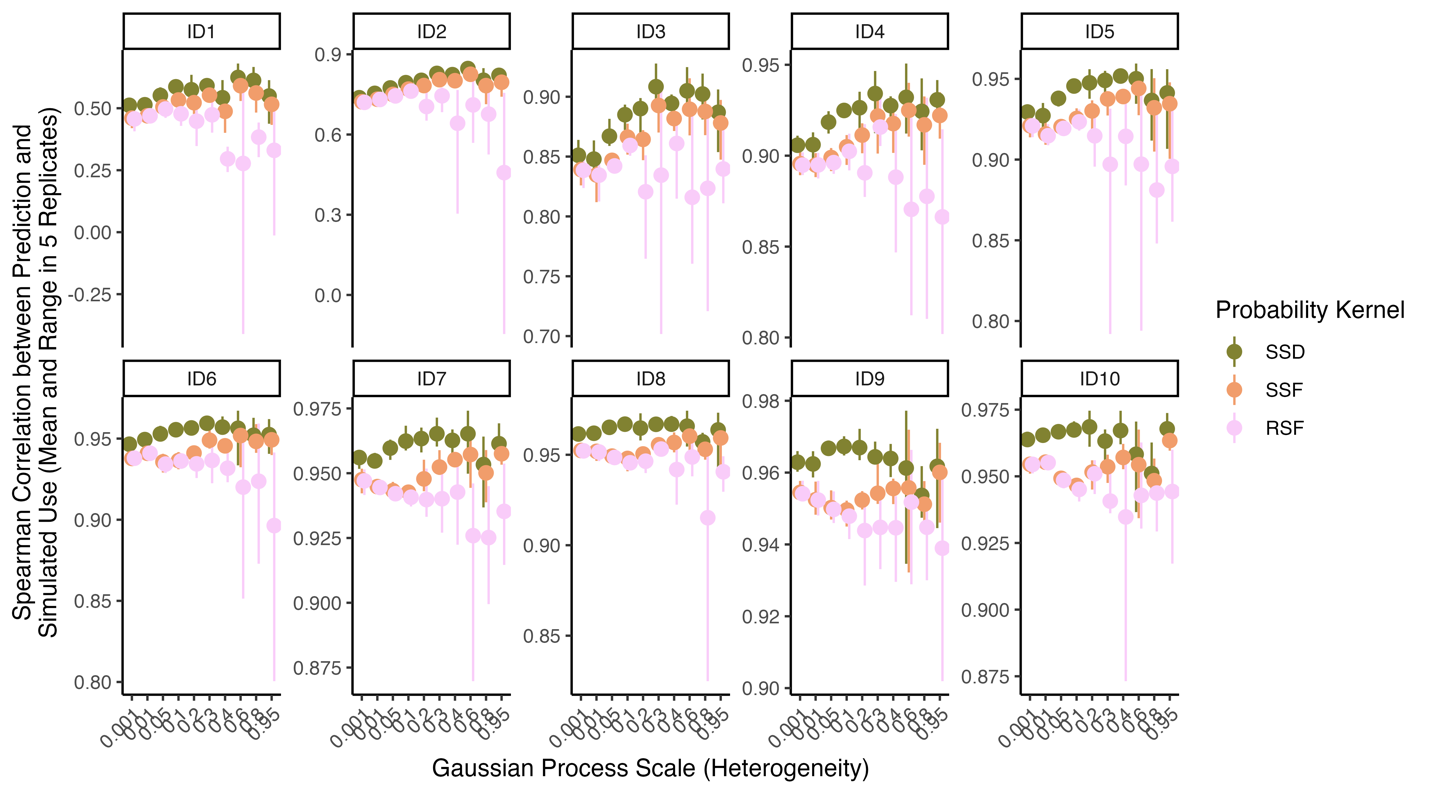

**S1 Fig 23: Spearman correlation between prediction and observed use kernels**

In five simulations per set of conditions, the Spearman correlation between simulated use kernels and SSD, SSF, and RSF kernels was calculated, generating a mean and range per set. In general, SSD was always favored, and SSF was favored over RSF at low selection strength and higher landscape heterogeneity.

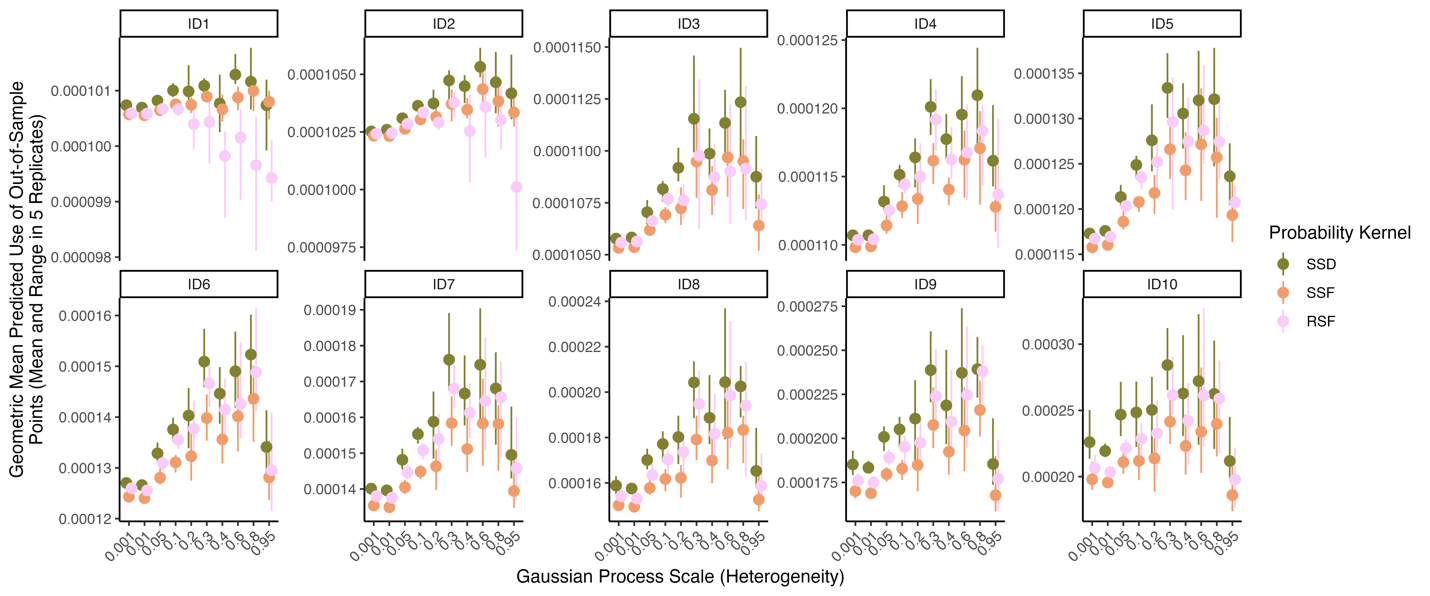

**S1 Fig 24: Geometric mean of predicted use across observed kernels**

In five simulations per set of conditions, we calculated the mean log-predicted use for SSD, SSF, and RSF predictions at every use location in simulated movements. This generated an estimate of use probability, weighted by observed use patterns, which was summarized as a mean and range for each set of conditions. In general, SSD was always favored, and SSF was favored over RSF at low selection strength and higher landscape heterogeneity. However, RSF was favored over SSF as selection strength increases.

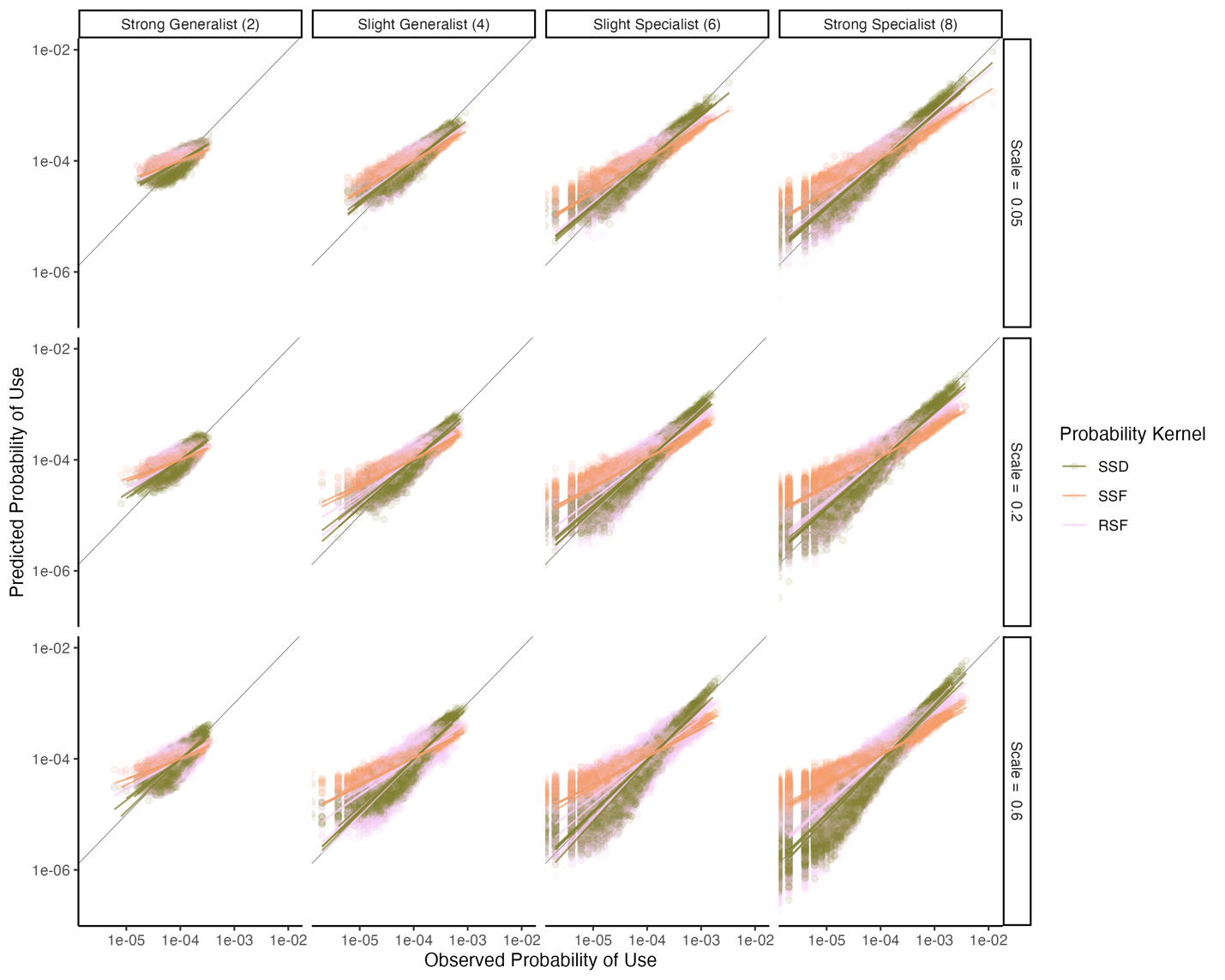

**S1 Fig 25: Observed use kernels vs. predicted use kernels across all simulations**

In five simulations per set of conditions, we visualized the observed probability of use vs that predicted probability of use under SSD, SSF, and RSF predictions. Points denote individual raster cells across all predictions, and lines denote linear relationships between observed and prediction kernels for each iteration. These relationships are visualized for four different animal selection profiles (2, 4, 6, 8) and three different scales of landscape heterogeneity (0.05, 0.2, 0.6).

**S2 Table 1: Data sources**

| Name | Source | Citation / identifier |
| --- | --- | --- |
| Red Deer track | “amt” package | Signer et al. (2019) |
| Roe Deer tracks (n = 5) | MoveBank: study9480191 | Cagnacci et al. (2011) |
| Fisher (n = 8) | MoveBank: study6925808 | (LaPoint et al. 2013) |
| sh_forest | “amt” package | Signer et al. (2019) |
| STRM 30m | https://srtm.csi.cgiar.org/srtmdata/ |  |
| Corine Land Cover (CLC) 2006, Version 2020_20u1 100m | http://land.copernicus.eu/pan-european/corine-land-cover/clc-2006/view |  |
| SRTM1 N42 W074 V3 | https://www.usgs.gov/centers/eros/science/usgs-eros-archive-digital-elevation-shuttle-radar-topography-mission-srtm-1 |  |
| NLCD 2011 Land Cover (CONUS) | https://www.mrlc.gov/data/nlcd-2011-land-cover-conus |  |
| US Population Density (2010) | https://www.sciencebase.gov/catalog/item/  57753ebee4b07dd077c70868 | Falcone (2016) |

**S2 Table 2: Model pseudocode for the case studies**

| Dataset | Model pseudocode (SSF) | Model pseudocode (RSF) |
| --- | --- | --- |
| Red Deer (Northern Germany) | dist.forest*forest + sl_ + log(sl_) | dist.forest*forest |
| Roe Deer (Northern Italy) | sqrt(ForestD) + Slope + sl_ + log(sl_) | sqrt(ForestD) + Slope |
| Fisher (NY, USA) | sqrt(DistForest) + sl_ + log(sl_) | sqrt(DistForest) |

*sl_ is step length in dataset-relevant units*

*step_id is the unique identifier of an observed step and the 40 randomly sampled points for that same context*

*dist.forest is the calculated distance to forest in the Germany Red Deer dataset based on a binary forest raster (sh_forest)*

*forest is the proportion of forest cover in the Germany Red Deer dataset based on a binary forest raster (sh_forest) aggregated by a factor of 5*

*ForestD is the calculated distance to forest in the Italy Roe Deer dataset based on a binary forest raster (composed of conifer, mixed, and broadleaf forest)*

*Slope is the calculated slope based on a digital elevation model*

*DistForest is the is the calculated distance to forest or wetland in the Fisher dataset based on a binary forest raster*

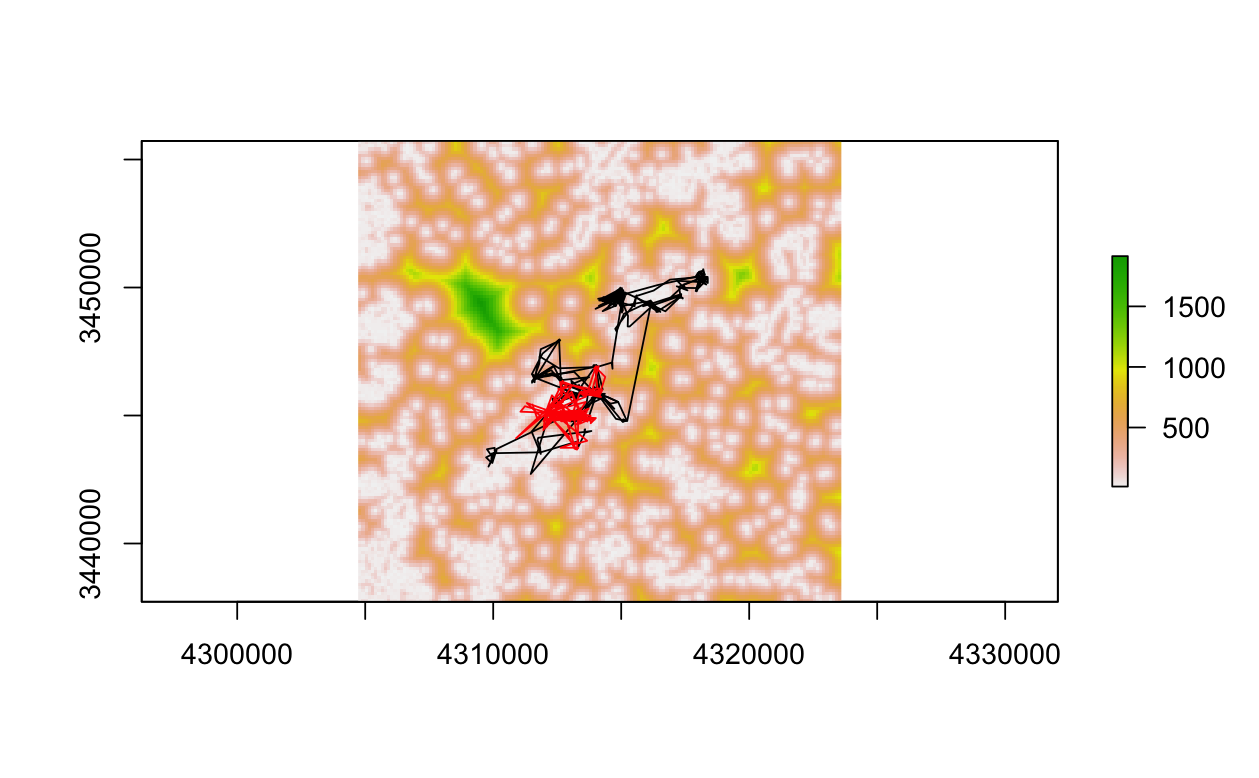

**S2 Fig 1: Distance to forest and movement in the red deer (*Cervus elaphus*) case study**

Distance to forest and in-sample movements (black) and out-of-sample movements (red) used in the red deer case study.

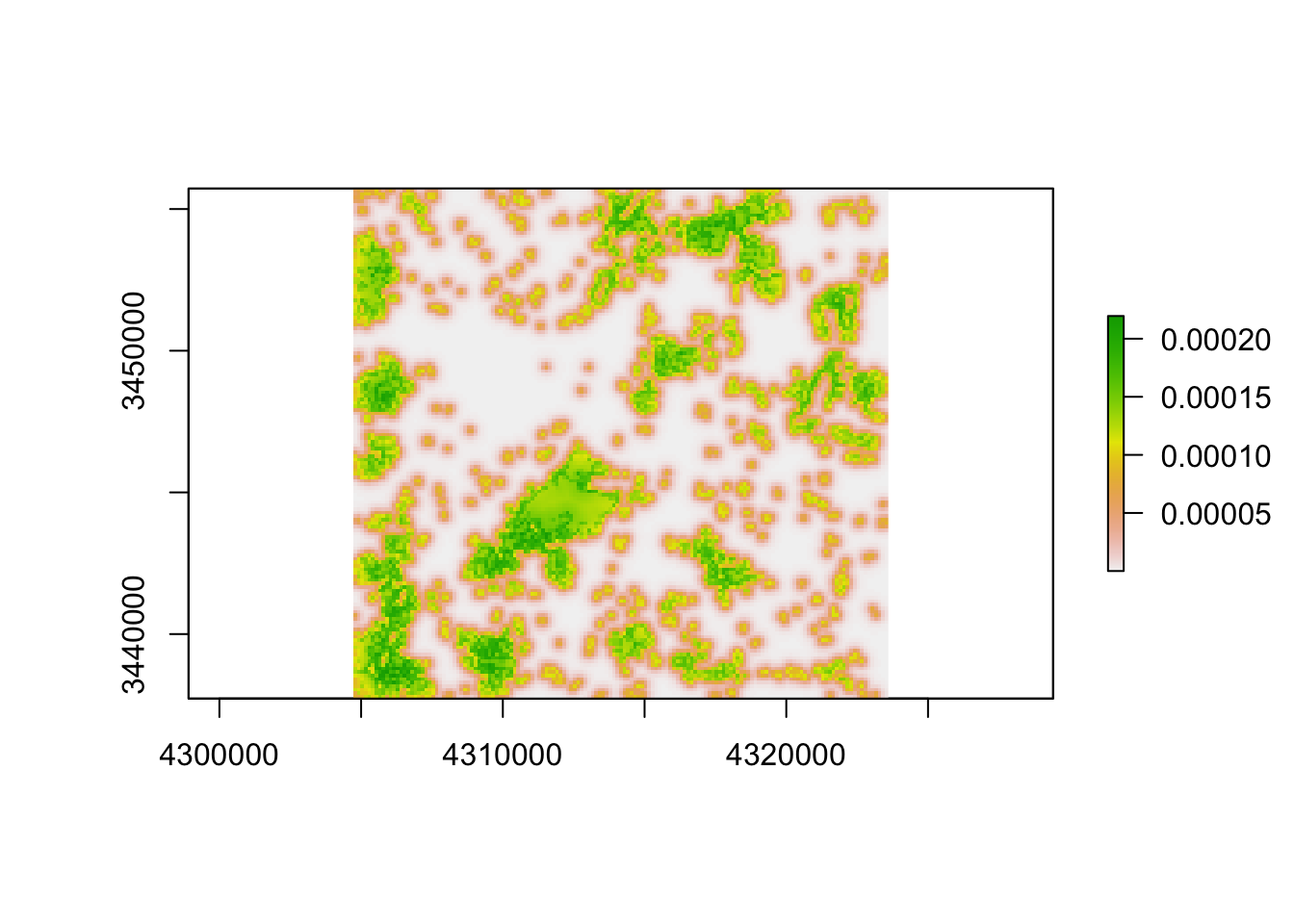

**S2 Fig 2: Stable state distribution (SSD) surface for the single individual in the red deer case study**

This figure was visualized in the main text but provided an individual color scale here to provide the viewer a more detailed view of the individual prediction.

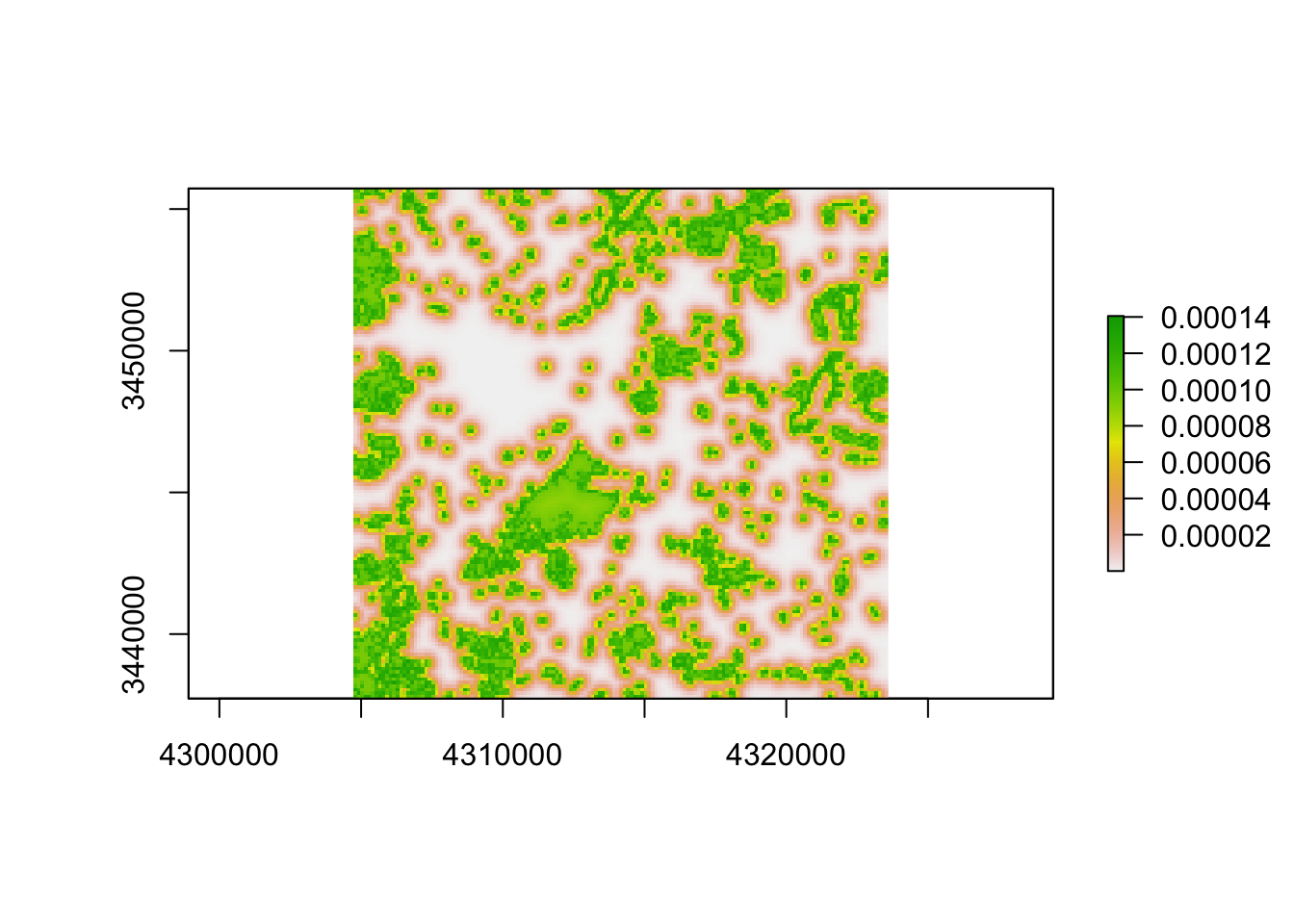

**S2 Fig 3: Exponentiated step-selection function (SSF) surface for the single individual in the red deer case study, kernelized to sum to one**

This figure was visualized in the main text, but provided an individual color scale here to provide the viewer a more detailed view of the individual prediction.

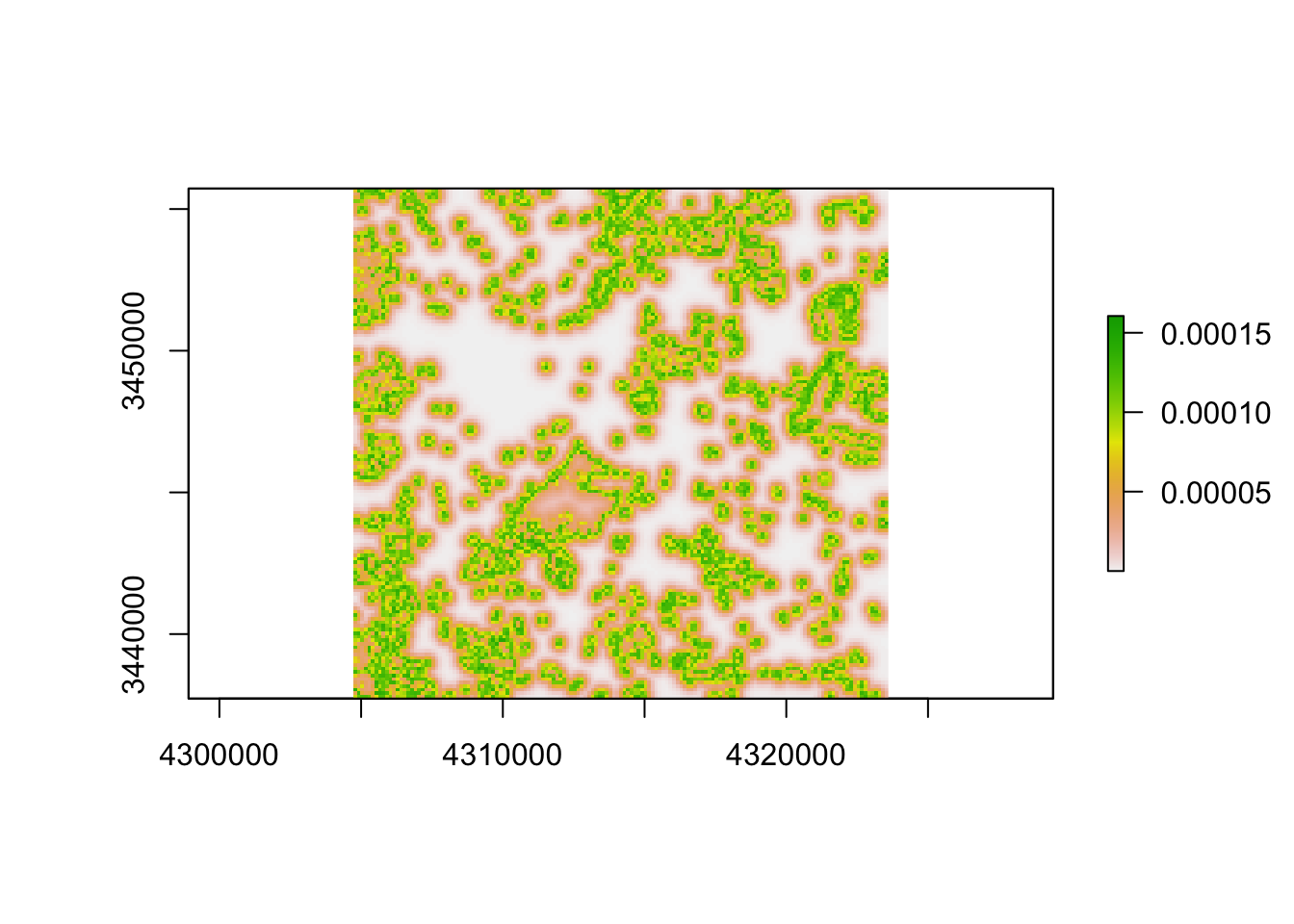

**S2 Fig 4: Probability of use surface from a resource selection function (RSF) for the single individual in the red deer case study, kernelized to sum to one**

This figure was visualized in the main text, but provided an individual color scale here to provide the viewer a more detailed view of the individual prediction.

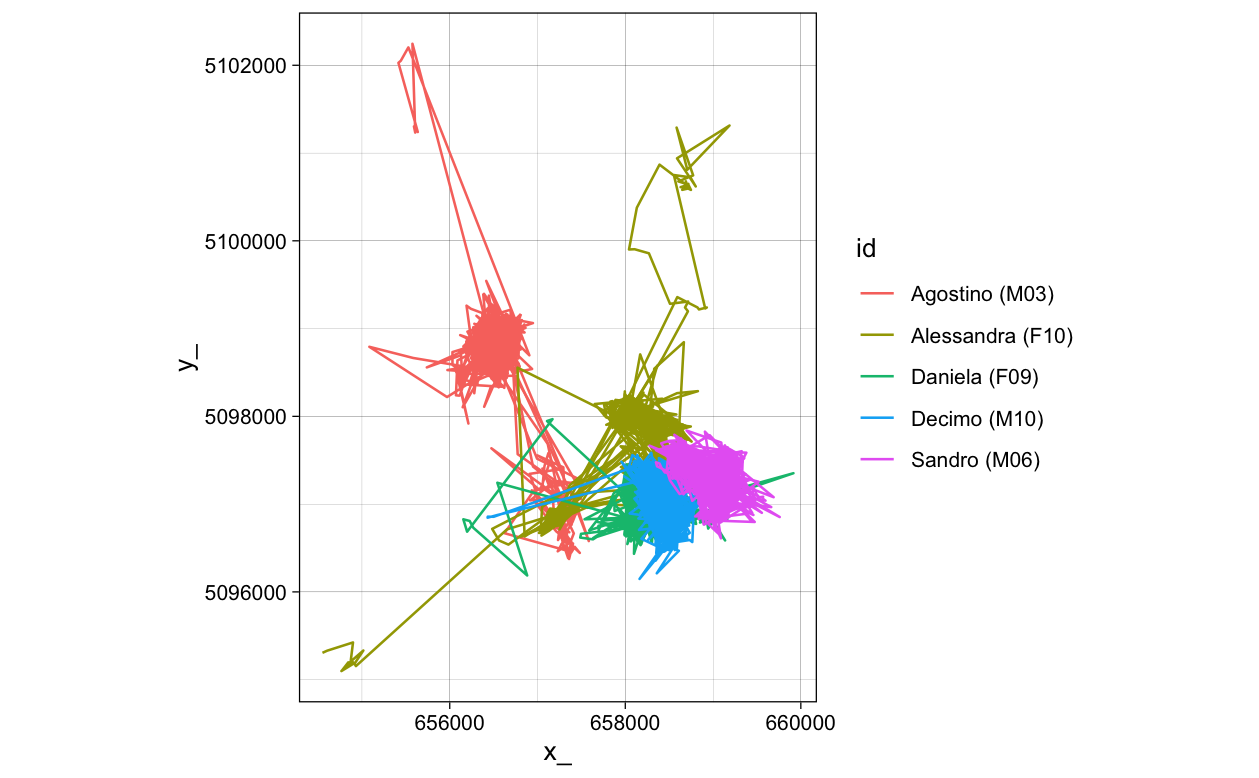

**S2 Fig 5: Individual roe deer (*Capreolus capreolus*) summer movement**

Summer roe deer movements were trimmed to Julian days 120-290 for use in the roe deer case study.

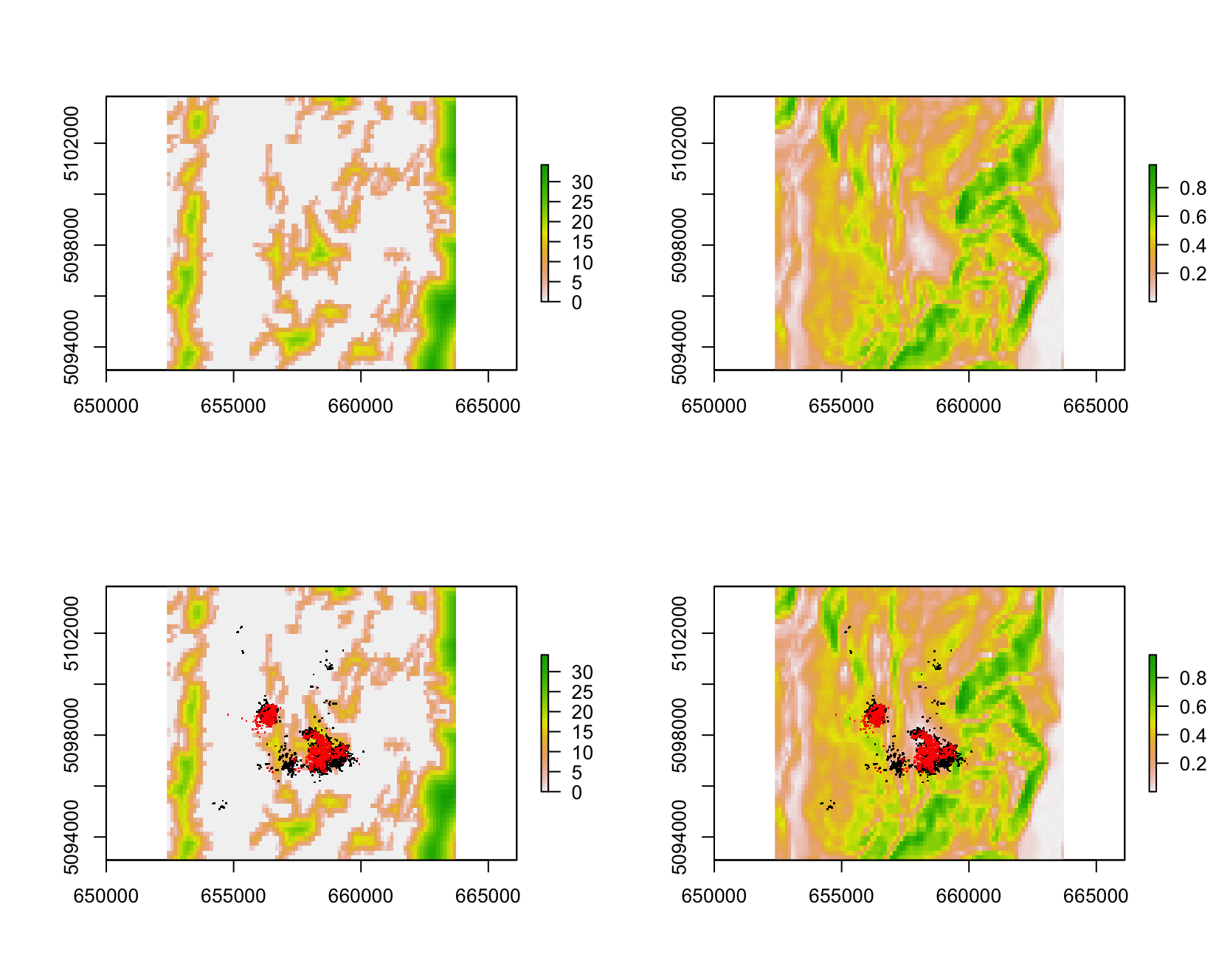

**S2 Fig 6: Visualization of roe deer covariates, in-sample, and out-of-sample data**

Square-root of distance to forest (in map units, left) and slope (in radians, right) used in the roe deer case study with training (black) and testing points (red) for each individual overlaid in the lower panel.

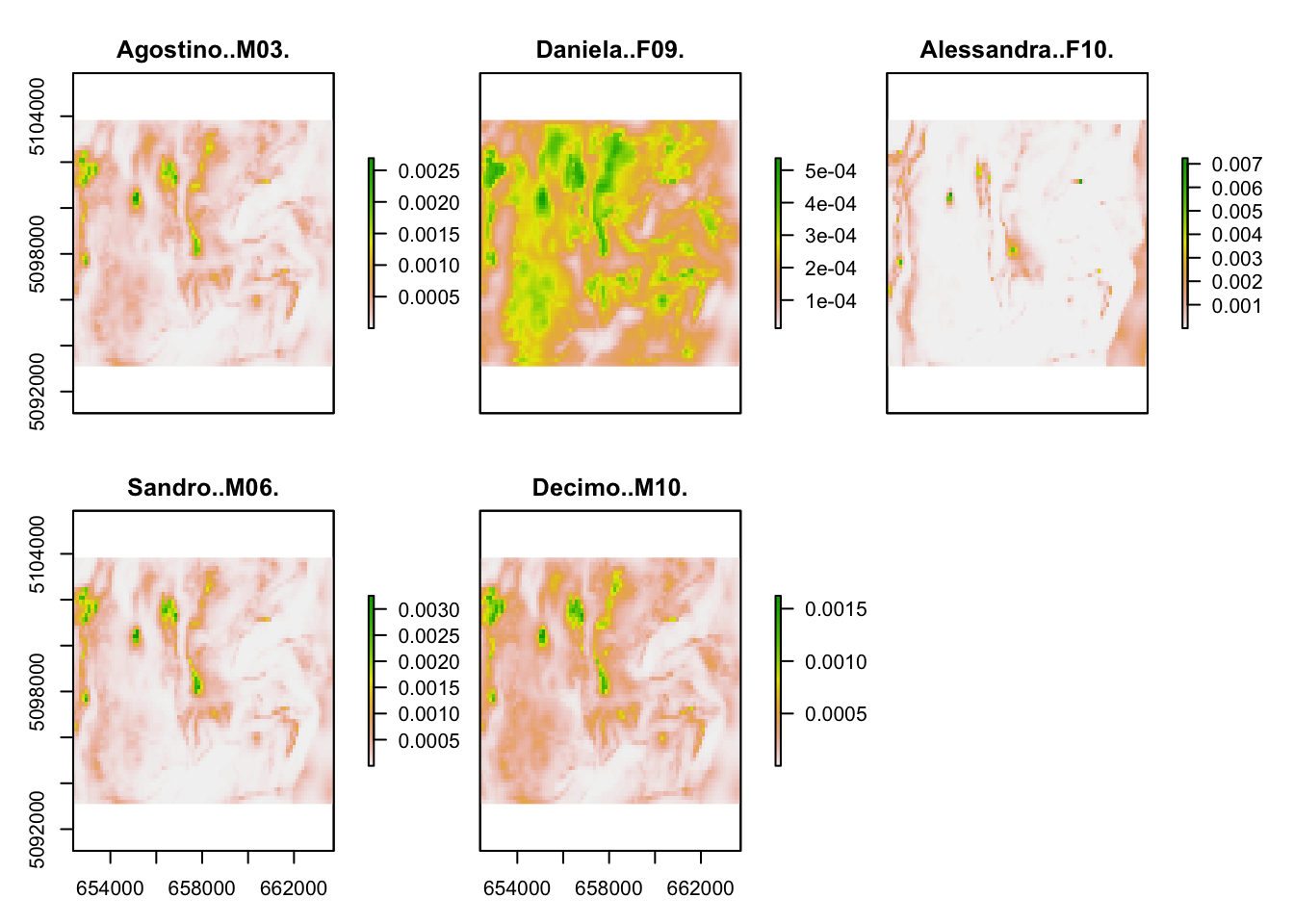

**S2 Fig 7: SSD predictions for individual roe deer**

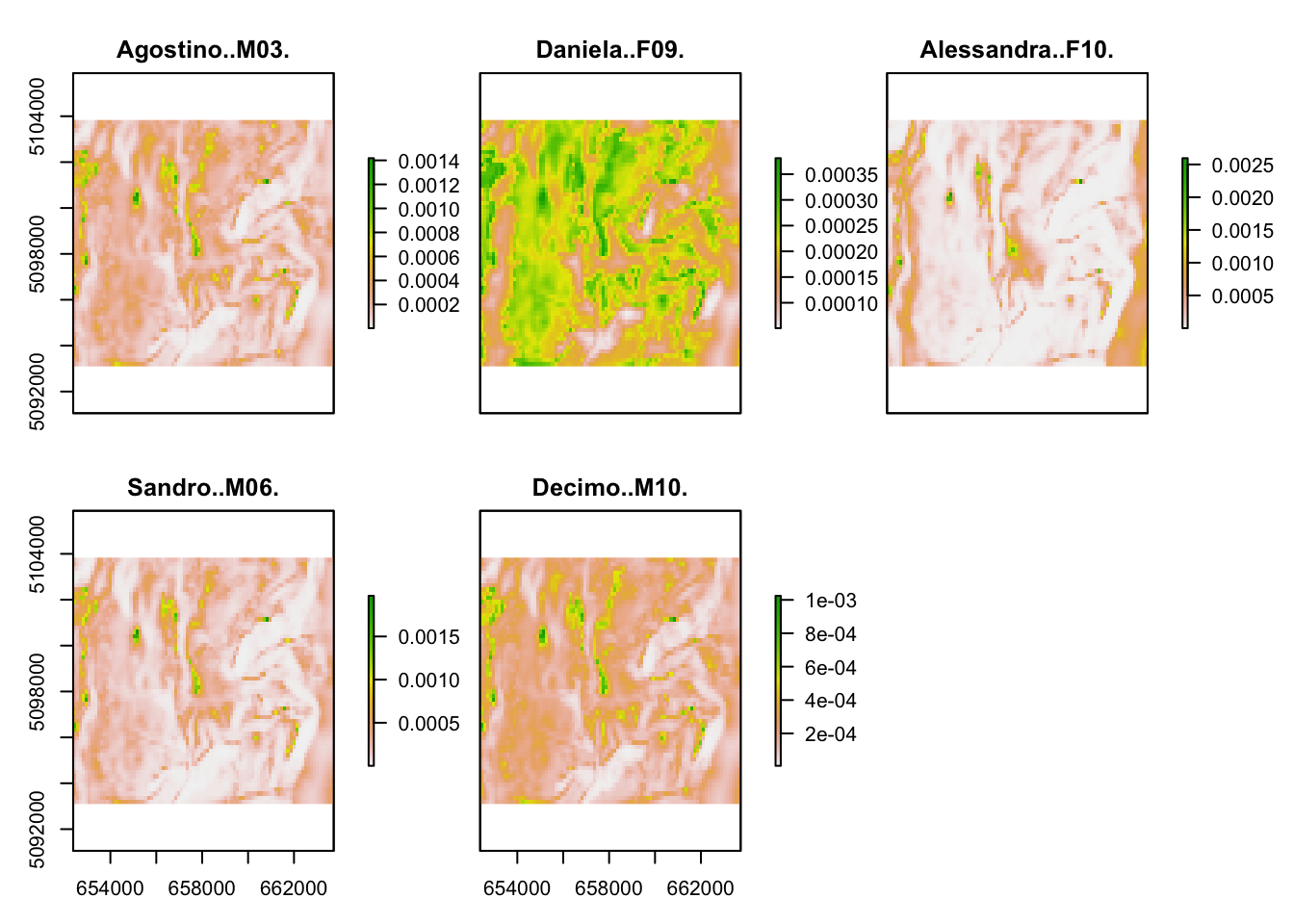

**S2 Fig 8: SSF predictions for individual roe deer**

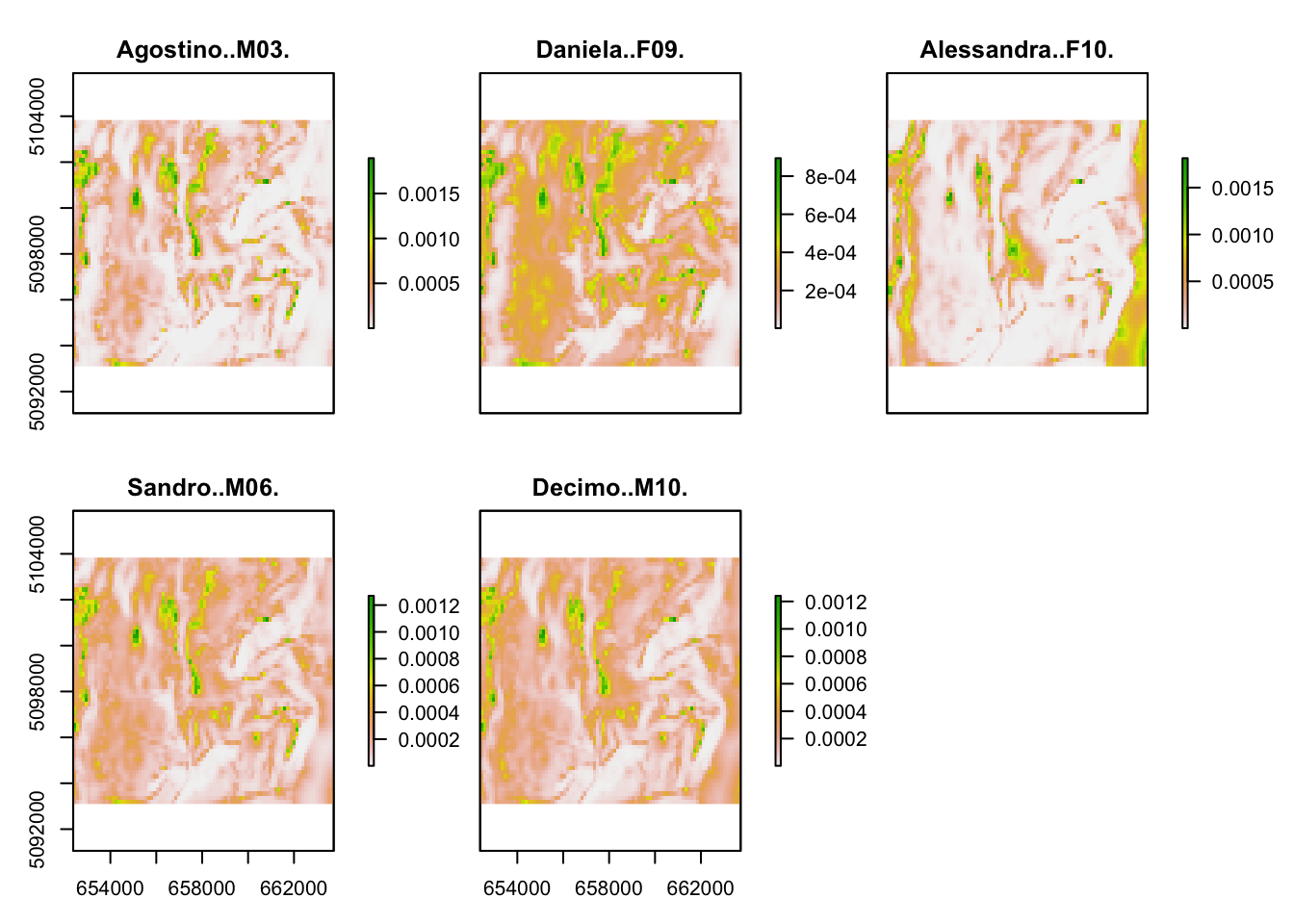

**S2 Fig 9: RSF predictions for individual roe deer**

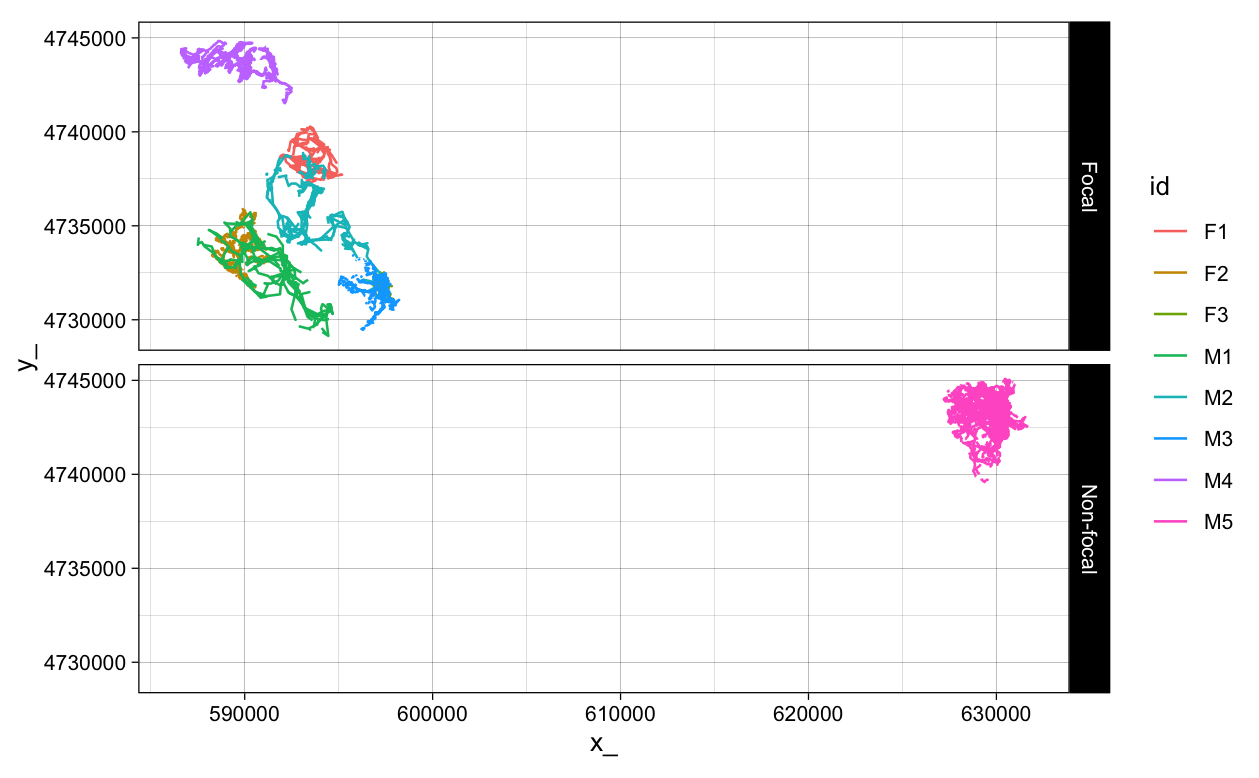

**S2 Fig 10: Individual Fisher (*Pekania pennanti*) movement**

Fisher movements (*n* = 8) used in the fisher case study, though only the “focal” seven individuals were modeled. The “non-focal” individual was used as a testing point for model predictions for the focal individuals.

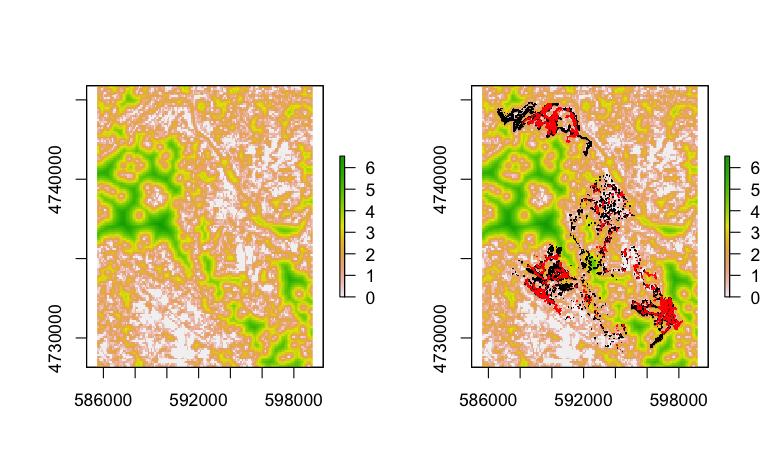

**S2 Fig 11: Visualization of focal fisher covariates, in-sample, and out-of-sample data**

Square-root of distance to forest underlying the local fisher movements with training (black) and testing points (red) for each individual overlaid in the lower panel. Note the extreme differences in scale from the non-focal landscape.

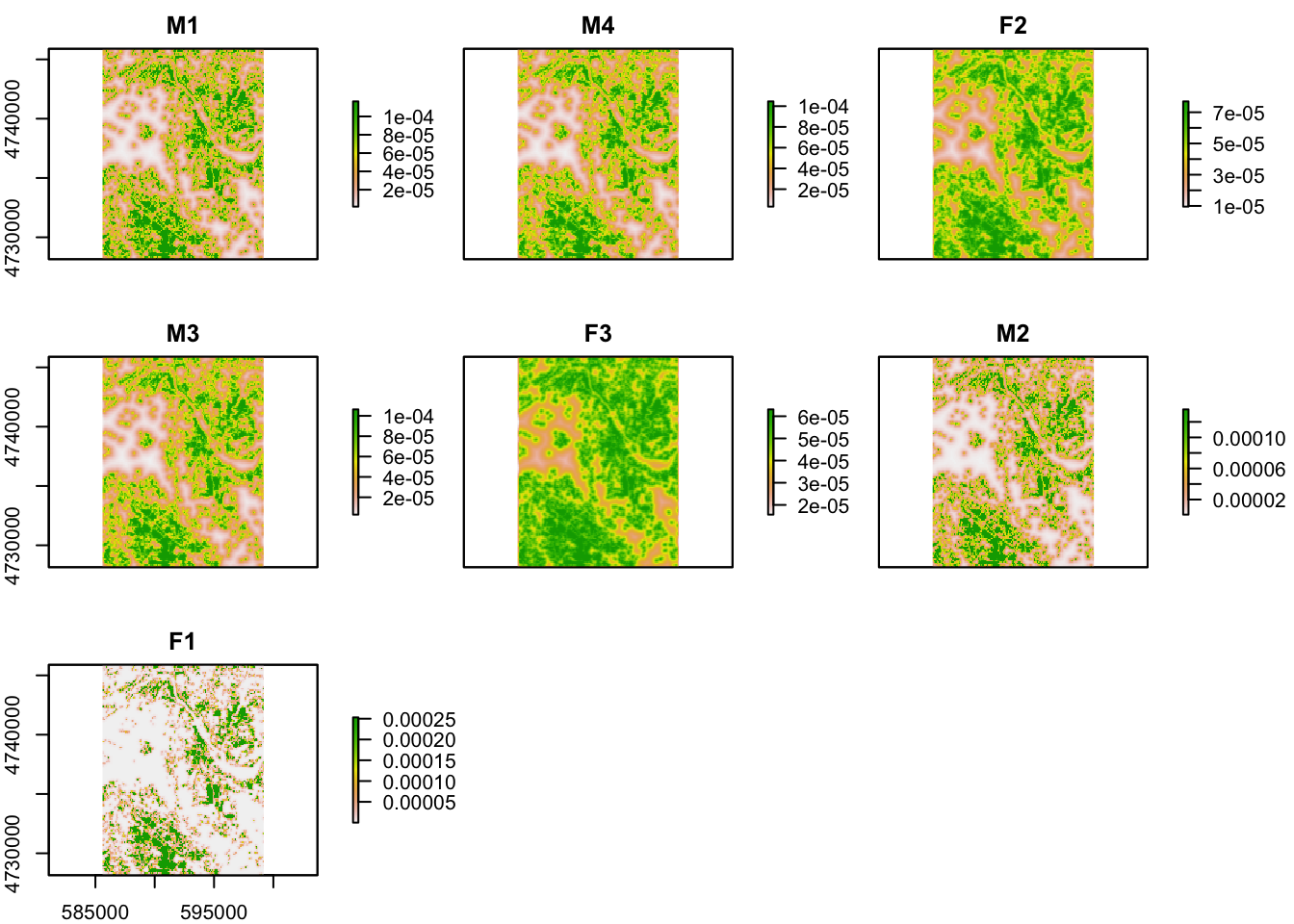

**S2 Fig 12: SSD predictions for focal fisher in the focal landscape**

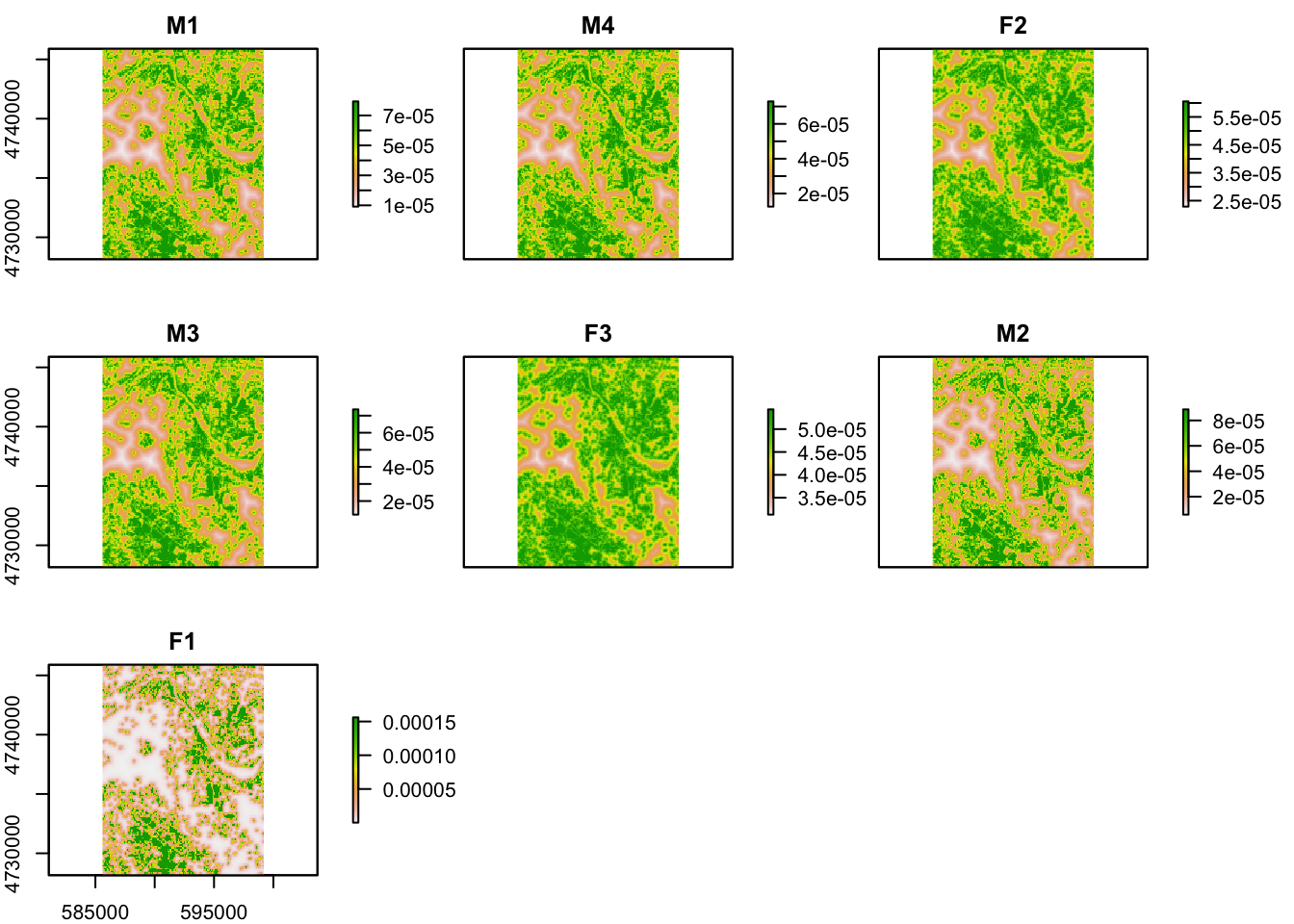

**S2 Fig 13: SSF predictions for focal fisher in the focal landscape**

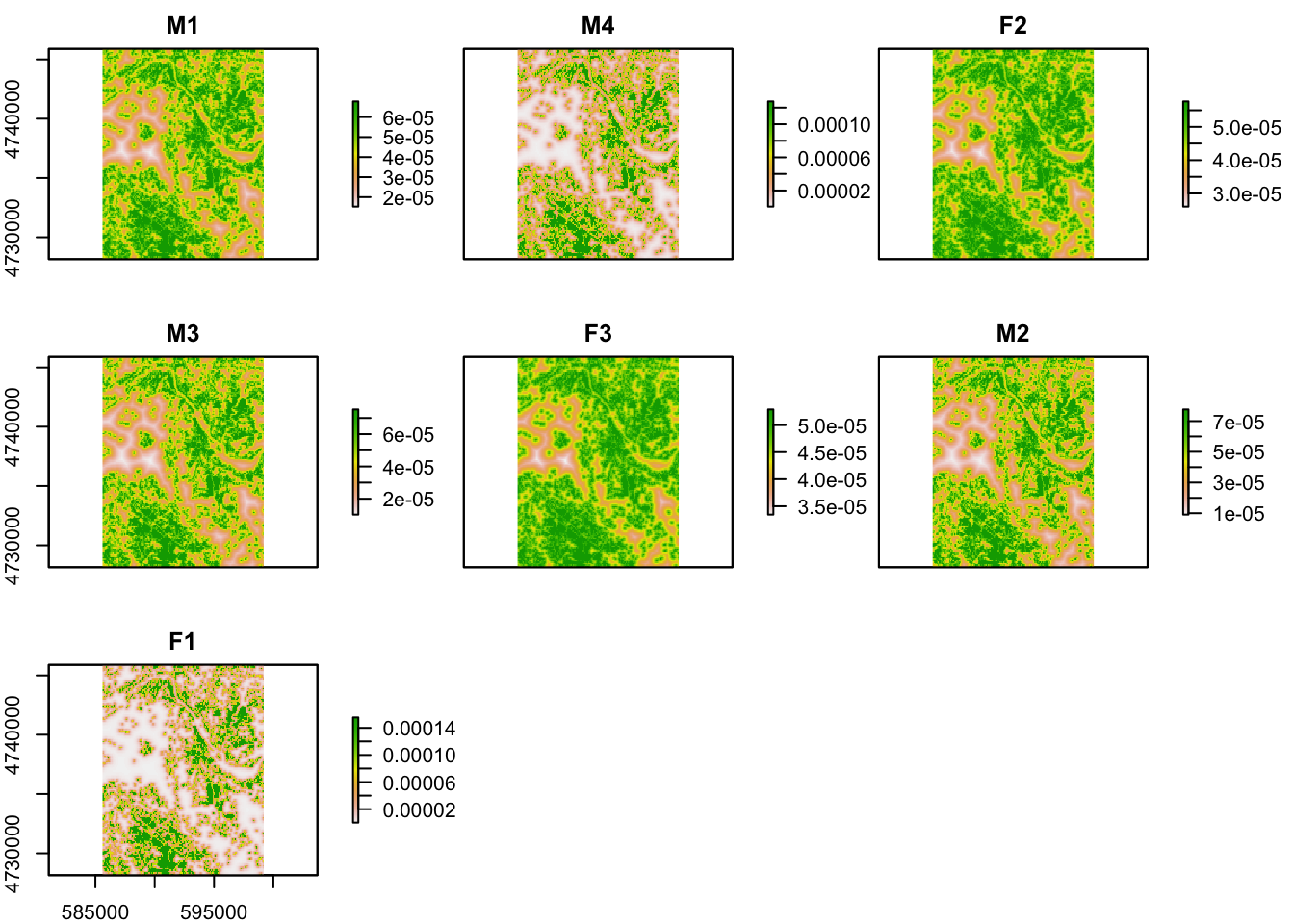

**S2 Fig 14: RSF predictions for focal fisher in the focal landscape**

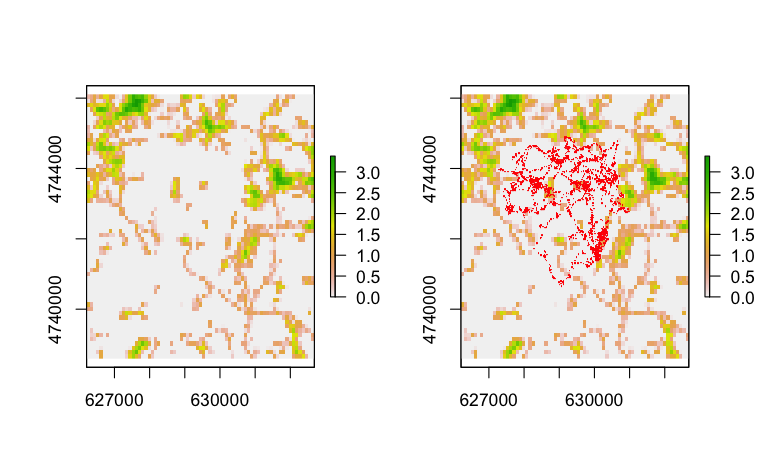

**S2 Fig 15: Visualization of non-focal fisher covariates and out-of-sample data**

Square-root of distance to forest or wetland (in map units, left) and square-root of human population density (right) underlying the non-local or non-focal fisher movements with testing points (red) for the out-of-sample individual overlaid in the lower panel. Note the extreme differences in scale from the focal landscape.

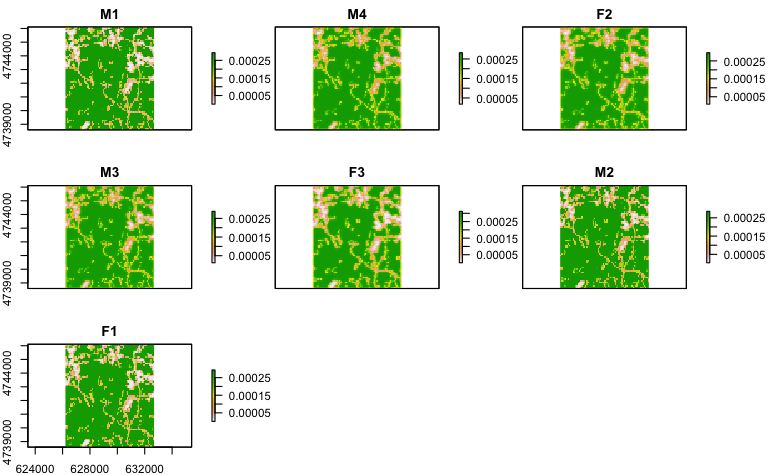

**S2 Fig 16: SSD predictions for focal fisher in the non-focal landscape**

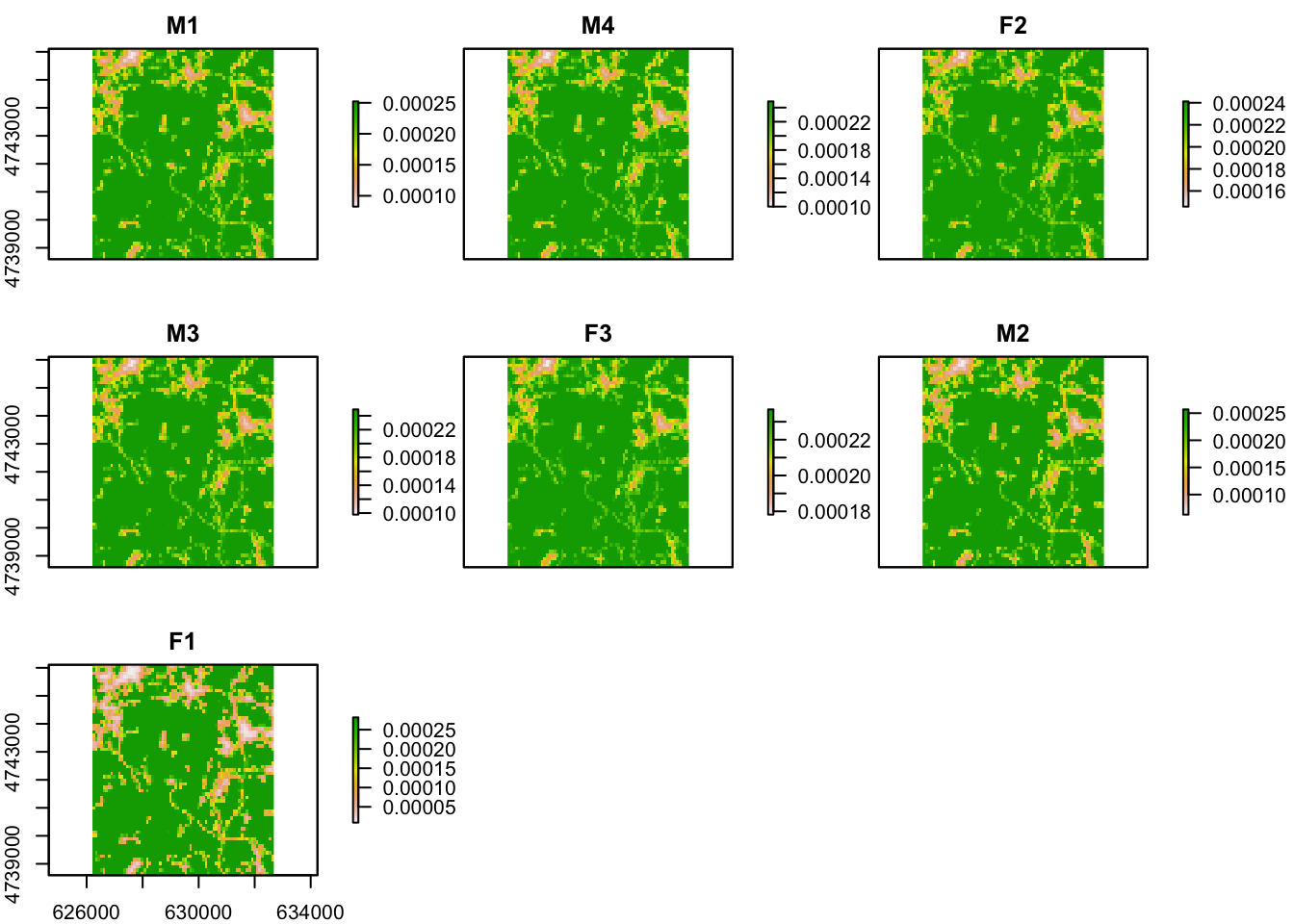

**S2 Fig 17: SSF predictions for focal fisher in the non-focal landscape**

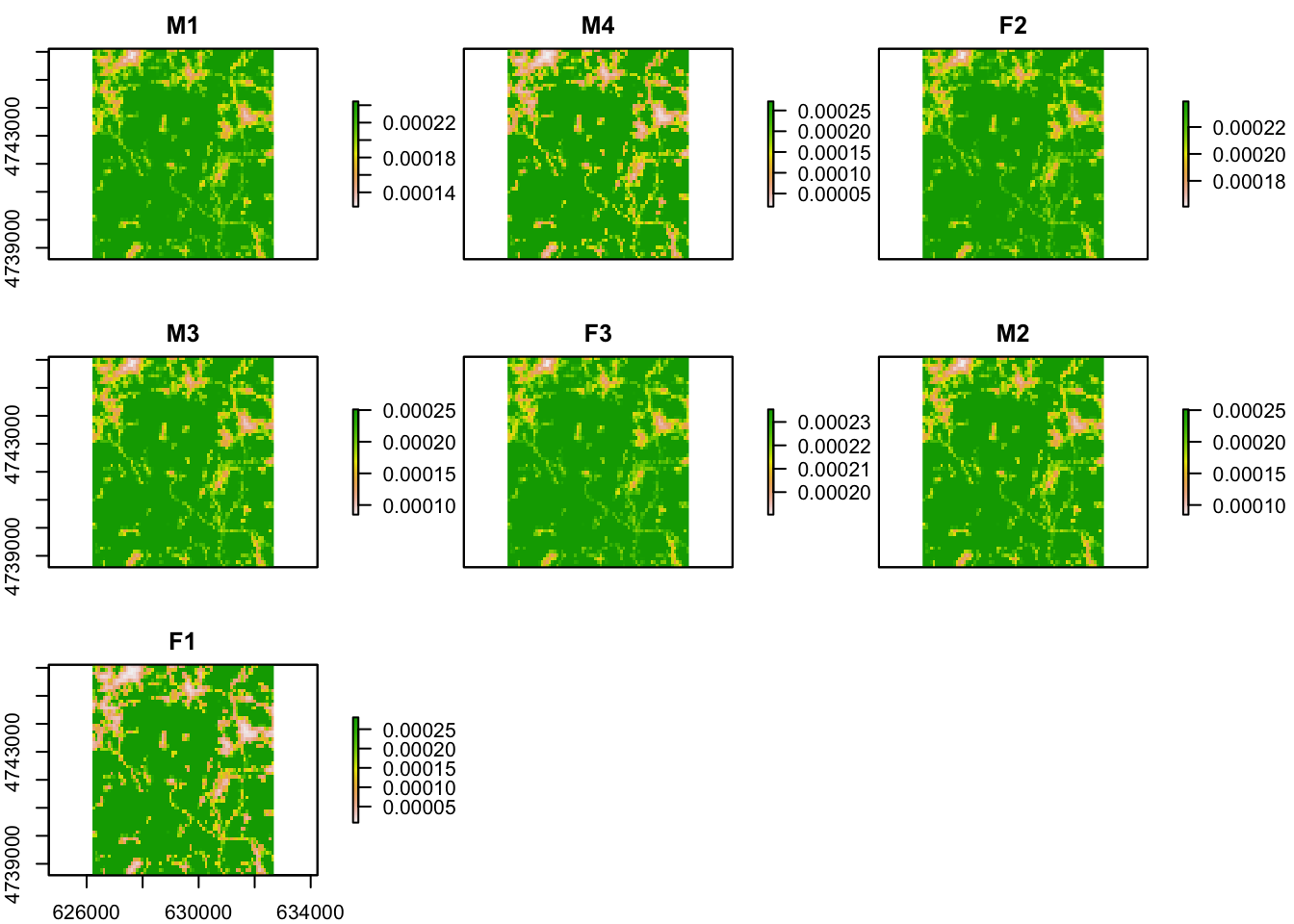

**S2 Fig 18: RSF predictions for focal fisher in the non-focal landscape**

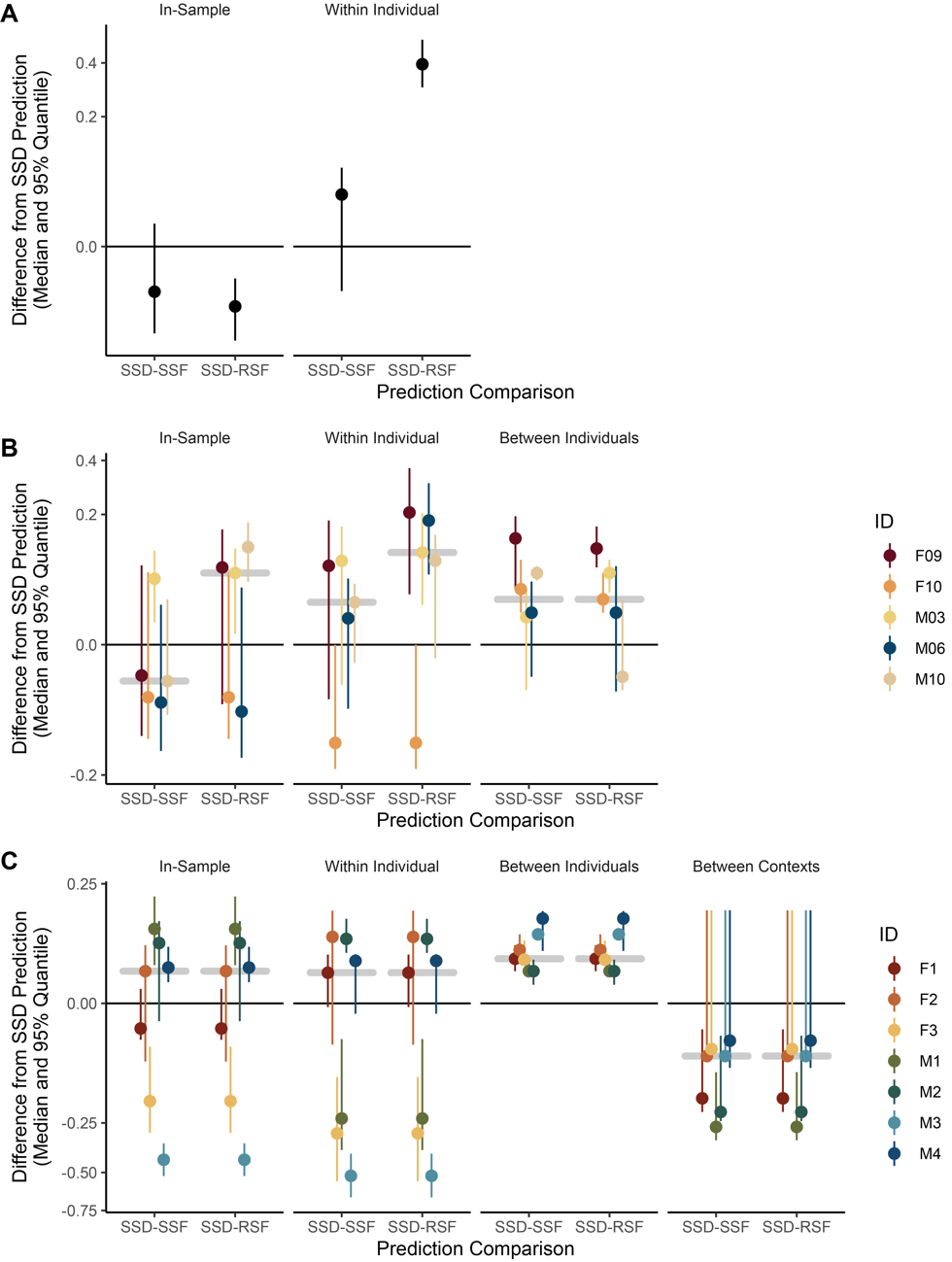

**S2 Fig. 19: Spearman rank comparisons for all case studies**

(**A-C**) Median and 95% quantiles in 100 bootstraps of differences between stable state distribution (SSD) and resource selection function (RSF) or step-selection function (SSF) predictions in Spearman rank correlation of predicted probability of use vs intensity of use. Bootstraps were performed by resampling modeled and out-of-sample data with replacement for (**A**) red deer (*Cervus elaphus*), (**B**) roe deer (*Capreolus capreolus*), and (**C**) fisher (*Pekania pennanti*) datasets. Validations were performed in four sets of comparisons, we used the first (chronologically) 75% of observed locations for individuals to predict on the modeled data (“In-Sample”), the next (chronologically) 25% of observed locations per individual (“Within Individual”), all observed locations for the other individuals in a dataset within the same geographic context (“Between Individuals”), and all observed locations for an individual in a completely different environmental context than the individuals modeled (“Between Contexts”). For visualization, the median difference for each set of comparisons was highlighted by a horizontal grey bar. Note that, for **C**, SSF and RSF predictions were not continuous enough to enable 10 unique quantiles; instead, “In-Sample”, “Within Individual”, and “Between Individuals” SSF and RSF comparisons rely on nine bins, and “In-Sample” relies on four bins.
