## Supplementary material for "Choices to landscapes: Mechanisms of animal movement scale to landscape patterns of space use": S3: PaperCompilation.html

S3: Paper Code Compilation


### S3: Paper Code Compilation

#### 2025-03-04

```
set.seed(100)
```

### 1 Overview

#### 1.1 Dependencies

```
# devtools::install_github("jmsigner/amt", force = T) #error in sapply for bursts, fixed in june 2024
library(tidyverse)
```

```
## Warning: package 'lubridate' was built under R version 4.3.3
```

```
## ── Attaching core tidyverse packages ──────────────────────── tidyverse 2.0.0 ──
## ✔ dplyr     1.1.4     ✔ readr     2.1.5
## ✔ forcats   1.0.0     ✔ stringr   1.5.1
## ✔ ggplot2   3.5.1     ✔ tibble    3.2.1
## ✔ lubridate 1.9.4     ✔ tidyr     1.3.1
## ✔ purrr     1.0.2     
## ── Conflicts ────────────────────────────────────────── tidyverse_conflicts() ──
## ✖ dplyr::filter() masks stats::filter()
## ✖ dplyr::lag()    masks stats::lag()
## ℹ Use the conflicted package (<http://conflicted.r-lib.org/>) to force all conflicts to become errors
```

```
library(lubridate)
library(amt)
```

```
## 
## Attaching package: 'amt'
## 
## The following object is masked from 'package:stats':
## 
##     filter
```

```
library(raster)
```

```
## Warning: package 'raster' was built under R version 4.3.3
```

```
## Loading required package: sp
## 
## Attaching package: 'sp'
## 
## The following object is masked from 'package:amt':
## 
##     bbox
## 
## 
## Attaching package: 'raster'
## 
## The following object is masked from 'package:amt':
## 
##     select
## 
## The following object is masked from 'package:dplyr':
## 
##     select
```

```
library(parallel)
library(data.table)
```

```
## Warning: package 'data.table' was built under R version 4.3.3
```

```
## 
## Attaching package: 'data.table'
## 
## The following object is masked from 'package:raster':
## 
##     shift
## 
## The following objects are masked from 'package:lubridate':
## 
##     hour, isoweek, mday, minute, month, quarter, second, wday, week,
##     yday, year
## 
## The following objects are masked from 'package:dplyr':
## 
##     between, first, last
## 
## The following object is masked from 'package:purrr':
## 
##     transpose
```

```
library(Matrix)
```

```
## 
## Attaching package: 'Matrix'
## 
## The following objects are masked from 'package:tidyr':
## 
##     expand, pack, unpack
```

```
library(progress)
library(pbapply)
library(pbmcapply)
library(igraph)
```

```
## 
## Attaching package: 'igraph'
## 
## The following object is masked from 'package:raster':
## 
##     union
## 
## The following objects are masked from 'package:lubridate':
## 
##     %--%, union
## 
## The following objects are masked from 'package:dplyr':
## 
##     as_data_frame, groups, union
## 
## The following objects are masked from 'package:purrr':
## 
##     compose, simplify
## 
## The following object is masked from 'package:tidyr':
## 
##     crossing
## 
## The following object is masked from 'package:tibble':
## 
##     as_data_frame
## 
## The following objects are masked from 'package:stats':
## 
##     decompose, spectrum
## 
## The following object is masked from 'package:base':
## 
##     union
```

```
library(rARPACK)
library(ggpubr)
```

```
## 
## Attaching package: 'ggpubr'
## 
## The following object is masked from 'package:raster':
## 
##     rotate
```

```
library(prioritizr)
```

```
## Warning: package 'prioritizr' was built under R version 4.3.3
```

```
library(conover.test)
library(scico)
library(rasterVis)
```

```
## Loading required package: lattice
```

```
library(scales)
```

```
## 
## Attaching package: 'scales'
## 
## The following object is masked from 'package:purrr':
## 
##     discard
## 
## The following object is masked from 'package:readr':
## 
##     col_factor
```

```
library(latex2exp)
```

#### 1.2 Create the surface prediction

To side-step issues where rasters are of different resolution, we can
create our own custom grid for predictions that match the most confined
extents of the rasters. We also might want to specify the resolution of
that underlying raster. If that raster is in utm (as defined below), we
can specify x and y cell sizes

```
create_mock_surface <- function(raster.list, multiple.extents = F, resolution = list(x = 100, y = 100)){
  
  # If rasters are all of the same extent, take the extent
  if(multiple.extents == F){
    xmin <- extent(raster.list)[1]
    xmax <- extent(raster.list)[2]
    ymin <- extent(raster.list)[3]
    ymax <- extent(raster.list)[4]
  }
  
  # If rasters are of different extent, take the overlap extent
  if(multiple.extents == T){
    xmin <- max(unlist(lapply(raster.list, function(x) {extent(x)[1]})))
    xmax <- max(unlist(lapply(raster.list, function(x) {extent(x)[2]})))
    ymin <- max(unlist(lapply(raster.list, function(x) {extent(x)[3]})))
    ymax <- max(unlist(lapply(raster.list, function(x) {extent(x)[4]})))
  }
  
  # Create new raster 
  mock.surface <- raster(
    ncol=(xmax-xmin)/resolution$x, # raster automatically rounds, total cols
    nrow=(ymax-ymin)/resolution$y, # raster automatically rounds, total rows
    xmn=xmin, # min x exent
    xmx=xmax, # max x exent
    ymn=ymin, # min y exent
    ymx=ymax, # max y exent
    crs = crs(raster.list[[1]])) # take CRS from first raster, requires that rasters match in CRS
  
  values(mock.surface) <- 1:(ncol(mock.surface)*nrow(mock.surface)) # not important, just for visualization
  
  return(mock.surface)
}
```

#### 1.3 Check model input

We should check that a model is correctly specified, before throwing
errors down the line.

```
check_ssf <- function(ssf.obj) {
  inherits(ssf.obj, c("fit_clogit")) | inherits(ssf.obj, c("gam")) # not broad enough, basically just from amt at the moment
  
}
```

#### 1.4 Find reasonable step distances for neighborhoods down the line

We should create a reasonable null step, and we can do so by using
the estimated unif distribution from amt. However, we could just as
easily take quantiles from the distribution of step lengths we observe,
instead.

```
step_distance <- function(ssf.obj, quantile) {
  if(!check_ssf(ssf.obj)) stop("Check that SSF model is valid")
  
  if(ssf.obj$sl_$name == "gamma") {
    step <- qgamma(quantile, # user specified quantile
                 shape = ssf.obj$sl_$params$shape, # estimated from amt
                 scale = ssf.obj$sl_$params$scale) # estimated from amt
  }
  
  if(ssf.obj$sl_$name == "exp") {
    step <- qexp(quantile, # user specified quantile
                 rate = ssf.obj$sl_$params$rate) # estimated from amt
  }
  
  if(ssf.obj$sl_$name == "unif") {
    step <- quantile(ssf.obj$model$model$sl_[which(as.character(ssf.obj$model$model[,1])=="1")], 0.95) # estimated from amt
  }
  
  return(step)
}
```

#### 1.5 Get data from prediction surface

Now that we have a valid SSF object and a prediction surface, we need
to find our prediction cells of interest. Lets get our raster data.

```
get_cells <- function(ssf.obj, mock.surface, raster, accessory.x.preds = NULL){
  if(!check_ssf(ssf.obj)) stop("Check that SSF model is valid")
  
  pred.xy <- raster::coordinates(mock.surface) # get coordinates from grid we created
  
  predict.data <- data.frame(cbind(pred.xy, raster::extract(raster, pred.xy, df=TRUE))) # makes raster values a data frame
  
  predict.data$step_id_unique = ssf.obj$model$xlevels$`strata(step_id_)`[1] # fix the strata to something reasonable
  
  if(!is.null(accessory.x.preds)) {
    predict.data <- cbind(predict.data, accessory.x.preds) # this adds extraneous x values that are not in matrix
  }
  
  cells <- nrow(pred.xy) # number of cells
  
  predict.data$cellnr <- 1:cells # assign cell numbers, redundant of ID
  
  return(predict.data)
}
```

#### 1.6 Check possible prediction surfaces

Not all combinations of parameter space are valid, and we often are
missing raster data at edges

```
get_cell_data <- function(ssf.obj, pred.data){
  if(!check_ssf(ssf.obj)) stop("Check that SSF model is valid")
  
  cells <- nrow(pred.data) # number of cells
  
  sample <- pred.data %>% 
    drop_na() %>% 
    sample_n(1)
  
  pred.data$sl_ <- sqrt((sample$x - pred.data$x)^2 + (sample$y - pred.data$y)^2) 
  sample$sl_ <- 0
  
  pred.data$x2_ <- pred.data$x
  pred.data$y2_ <- pred.data$y
  
  sample$x2_ <- sample$x
  sample$y2_ <- sample$y
  
  # relying on amt step-selection models, we can predict log-RSS
  # there are better ways to predict, but this is simple
  
  if("gam" %in% class(ssf.obj)) {
    log.rss <- log(c(predict(ssf.obj, newdata = pred.data))/ c(predict(ssf.obj, newdata = sample)))
    full.raster.data <- data.frame(pred.data, lRSS = log.rss)
  } else {
    
    log.rss <- amt::log_rss(ssf.obj, # the model
                          pred.data, # the raster data (including missing values)
                          sample,  # a row of the raster data (excluding missing values)
                          ci = NA) 
    
    if(c(Inf) %in% abs(log.rss$df$log_rss)) {
      pred.data$sl_[which(pred.data$sl_ == 0)] <- 0.01
      sample$sl_[which(sample$sl_ == 0)] <- 0.01
      log.rss <- amt::log_rss(ssf.obj, # the model
                          pred.data, # the raster data (including missing values)
                          sample,  # a row of the raster data (excluding missing values)
                          ci = NA)
    }
    
    full.raster.data <- data.frame(pred.data, lRSS = log.rss$df$log_rss)
    
  }
   # bind predictions to the original data
  
  return(full.raster.data)
  
}
```

#### 1.7 We need a quick way to find values for comparison

We need to find neighbors of cells in our matrix. This is easy, but
could require n^3 comparisons. So many comparisons are computationally
inefficient. One thing we can rely on is that all cells in a raster (or
matrix) can be indexed. Additionally, distances between cells in a
raster are repetitive. In general, there are only ~0.2% unique pairwise
distances between cell indices. We can find these unique distances
because we know the difference in index of the comparisons, the column
identify of the minimum index, and the observed distance. This allows us
to have a table that we can call repetitively instead of recalculating
millions of distances.

```
neighbor_lookup <- function(mock.surface, cell.data, cell.data.list = NULL){
  cols <- mock.surface@ncols # columns in prediction
  rows <- mock.surface@nrows # rows in prediction
  cells <- cols*rows # number of cells
  index <- 1:cells # all index values in our prediction raster
  
  # create a matrix for each column in the first row and its comparisons to distance to all other cells 
  
  # if(is.null(cell.data.list)){
  #   print("Splitting cell.data into list")
  #   cell.data.list. <- split(cell.data, cell.data$cellnr) # split the prediction data into row-wise lists to use lapply
  # }
  # 
  # if(!is.null(cell.data.list)){
  #   print("Using inputted list of cell data")
  #   cell.data.list. <- cell.data.list # split the prediction data into row-wise lists to use lapply
  # }
    
  # this is a progress bar we can use in a for loop
  pb <- progress_bar$new(format = "(:spin) [:bar] :percent [Elapsed time: :elapsedfull || Estimated time remaining: :eta]", total = cols, complete = "=", incomplete = "-", current = ">", clear = FALSE, width = 100)
  
  for(i in 1:cols) { # step through columns
    dist <- pointDistance(cell.data[i,c("x","y")], # choose the first row cell by column (raster indices are row-wise)
                            cell.data[i:cells,c("x","y")], # choose all other cells
                            lonlat = F) # we are using UTM 
    if(i == 1) mat.dist <- matrix(dist, ncol = 1)
    if(i > 1) mat.dist <- cbind(mat.dist, c(dist, rep(NA, i-1))) # for each additional column, there are i-1 comparisons that are repeated (unnecessary)
    pb$tick() # for progress bar
  }
  
  return(mat.dist)
}
```

#### 1.8 Find neighbors within distance

Now that we have our call-up table, we can use it to generate
neighbors. This is one of the most taxing (computationally) steps of the
whole process.

```
neighbor_finder <- function(ssf.obj = m2, cell.data, neighbors.found, quantile = 0.95, cell.data.list = NULL, distance.override = NULL){
  
  if(is.null(distance.override)) neighborhood.distance <- step_distance(ssf.obj, quantile) # take the X% step distance as your neighborhood 
  
  if(!is.null(distance.override)) neighborhood.distance <- distance.override
  
  cols <- ncol(neighbors.found) # columns of our call-up table
  differences <- nrow(neighbors.found) # number of differences in index values 
  
  print("Creating neighbor comparisons")
  vector <- c(neighbors.found) # convert the neighbors to a vector we can index later, this is the purpose of all those NA's earlier based on i-1 unique distances
  
  print("Finding valid comparisons")
  valid <- vector < neighborhood.distance # T/F whether those neighborhood distances are less than our threshold
  
  if(is.null(cell.data.list)){
    print("Splitting cell.data into list")
    cell.data.list. <- split(cell.data, cell.data$cellnr) # split the prediction data into row-wise lists to use lapply
  }
  
  if(!is.null(cell.data.list)){
    print("Using inputted list of cell data")
    cell.data.list. <- cell.data.list # split the prediction data into row-wise lists to use lapply
  }
  
  print("Running comparisons")
  neighbor.mat <- pblapply(cell.data.list., function(x){ # step through each row (see list split above)
    focal <- as.numeric(x$cellnr) # value of row cell number 
    delta <- abs(focal - cell.data$cellnr) # difference in row cell number vs all others
    index <- ifelse(focal < cell.data$cellnr, focal, cell.data$cellnr) # report the minimum cell index (based on our call-up structure)
    
    index.col <- index%%cols # use the remainder function to get the column number (see below, we have to force zeros to the column number because the remainder of the final column is zero)
    
    df <- data.frame(difference = delta + 1, # we have to add one because differences of 0 are stored in row 1, differences of 1 in row 2, etc. 
                     col = ifelse(index.col == 0, cols, index.col), # forcing remainders of zero the number of columns
                     cell.nr = cell.data$cellnr) # just tracking the cell number we are comparing against for use later
    
    # filter the data frame based on whether our call-up values are less than the neighborhood
    df <- df %>% 
      filter(valid[difference+((col-1)*differences)]) 
    
    # filter the data set to unique rows and columns for call-up (to accommodate memory issues)
    df.distinct <- df %>% distinct(difference, col) 
    
    # find the unique distances (trims time down)
    df.distinct$distances <- vector[df.distinct$difference + ((df.distinct$col - 1)*differences)]
    
    # throw the unique distances back to the full data set 
    df <- merge(df, df.distinct, by = c("difference","col"))
    
    # package into a nice data frame for export
    data.frame(row = focal, column = df$cell.nr, distance = df$distances)
  })
  
  # bind the output list
  neighbors <- rbindlist(neighbor.mat)
  
  # create a sparse matrix based on the focal id, alternate id, and distance
  sparse.neighbors <- sparseMatrix(i = neighbors$row, j = neighbors$column, x = neighbors$distance)
  
  # return both the matrix and the unbound list of neighbor cells
  return(list(matrix = sparse.neighbors, by.cell = neighbor.mat))
}
```

#### 1.9 We need to split data into focal and neighbor groups

Now that we have data divided so that we know the neighbors of a
focal cell, we can split the data into a format that is conducive to SSF
predictions. We split these into two data frames, `.given`
for the focal cell and `.for` for the neighboring cells.

```
compile_ssf_comparisons <- function(sparse.neighbors, cell.data) {
  
  # this is why the export of the neighbors as individual lists was important
  ssf.comparisons <- lapply(sparse.neighbors$by.cell, function(x){ 
    baseline <- cell.data[x$column[which(x$distance == 0)],] # baseline will have a distance of zero (focal)
    
    baseline$sl_ <- 0 # create a variable "step" that records this zero distance
    
    alternate <- cell.data[x$column,] # grab all the other cell.data for neighboring cells (including focal cell)
    
    alternate$sl_ <- x$distance # force distance to this new variable step
    
    list(.given = baseline, .for = alternate) # return a list of focal and neigboring cell data
    
  })
  
  return(ssf.comparisons)
}
```

#### 1.10 Now we need to predict our surface

Using our fit movement model and the metadata for our prediction
surface, we can estimate the relative risk of selecting the focal cell
and all other cells in the neighborhood. We can show that all
probabilities of “choosing” a cell in the prediction surface must sum to
one, thus the probability of selecting the focal cell is the inverse of
he sum of all relative probabilites. From log-RSS, we just exponentiate,
and take the inverse sum. To find the probability of choosing all cells,
we just multiply the probability of selecting the focal cell against all
relative risks! Easy! This tells us the probability of selecting each of
those cells given the comparison to the focal cell, so not the out-right
probability of selection, but it gets us closer.

```
predict_ssf_comparisons <- function(ssf.obj, ssf.comparisons = ssf.comparisons) {
  
  print("Estimating probability surface")
  
  ## this is straight from amt - i just dont want to recalculate the uncenter term every exposure because it saves about 90% of the time of this function
    uncenter <- sum(coef(ssf.obj$model) * ssf.obj$model$means, na.rm=TRUE)
  
  prediction.list <- pbmclapply(ssf.comparisons, function(x){ # step through the list of SSF objects
    x1_dummy <- x$.for
    x2_dummy <- x$.given
    
    x1_dummy$step_id_ = ssf.obj$model$model$`strata(step_id_)`[1]
    x2_dummy$step_id_ = ssf.obj$model$model$`strata(step_id_)`[1]
    
    x1_dummy$sl_[which(x1_dummy$sl_ == 0)] <- 0.001
    x2_dummy$sl_[which(x2_dummy$sl_ == 0)] <- 0.001
    
    #Calculate y_x
    pred_x1 <- predict(ssf.obj$model, newdata = x1_dummy, type = "lp", reference = "sample",
                    se.fit = F)
    pred_x2 <- predict(ssf.obj$model, newdata = x2_dummy, type = "lp", reference = "sample",
                    se.fit = F)
    
    y_x1 <- pred_x1 + uncenter
    y_x2 <- pred_x2 + uncenter
  
    log_rss <- unname(y_x1 - y_x2)
    x$.for$Prob <- exp(log_rss)*(1/sum(exp(log_rss))) # exponentiate and multiply against relative risk
    
    x$.for # return the data frame with probabilities
  })
  
  print("Compiling probability surface")
  
  for(i in 1:length(prediction.list)){
    prediction.list[[i]]$focal.cell <- i # specify the focal cell for each comparison
  } 
  
  print("Making sparse matrix for transitions")
  bound <- rbindlist(prediction.list) # bind all data frames 
  
  # use indexing to make a massive sparse matrix quickly 
  # Sparse.Matrix.lrss <- sparseMatrix(bound$focal.cell, bound$cellnr, x = bound$log_rss)
  # Sparse.Matrix.rss <- sparseMatrix(bound$focal.cell, bound$cellnr, x = bound$rss)
  Sparse.Matrix <- sparseMatrix(bound$focal.cell, bound$cellnr, x = bound$Prob)
  # Sparse.Matrix.l <- sparseMatrix(bound$focal.cell, bound$cellnr, x = bound$Prob.l) 
  # Sparse.Matrix.h <- sparseMatrix(bound$focal.cell, bound$cellnr, x = bound$Prob.h) 
  
  # return the prediction list and sparse matrix
  return(list(prob.surface = prediction.list, 
              # lrss.matrix = Sparse.Matrix.lrss, rss.matrix = Sparse.Matrix.rss, 
              prob.matrix = Sparse.Matrix
              # , sparse.matrix = Sparse.Matrix, sparse.matrix.l = Sparse.Matrix.l, sparse.matrix.h = Sparse.Matrix.h
              ))  
}
```

#### 1.11 Creating a function to bootstrap GPS locations to compare intensity vs predicted use.

```
compare_spearman <- function(data = ssf1.train, rstack = deer1.rstack, bootstrap = 100, bins = 10){
  rasters <- names(rstack) # layers of predicted surface
  intersected <- data %>% 
    extract_covariates(rstack) %>% 
    drop_na() # pull out all the predictions per point
  
  predictions <- values(rstack) # grab the matrix of predicted surfaces
  
  interval <- apply(predictions, 2, function(x) {
    quantile(x, seq(0, 1, length.out = bins + 1), na.rm = T)
    # seq(min(x), max(x), length.out = bins + 1)
  }) # generate centiles of the predicted surfaces
  
  
  unique.bins <- apply(interval, 2, function(x) length(unique(x)))<(bins + 1)
  
  if(T %in% unique.bins){
    print("unique bins less than requested bins, decreasing bins to minimum")
    
    index <- which(unique.bins)
    
    for(i in index){
      interval[which(duplicated(interval[,i])),i] <- NA
    }
    
  }
  
  # df.area <- expand.grid(bin = 1:bins,
  #                        layer = 1:length(rasters)) # make a storage data frame
  # df.area$area <- NA # add the outcome 
  # total.size <- nrow(predictions)
  # for(i in 1:length(rasters)){
  #   for(j in 2:(bins + 1)){
  #     df.area[j-1,i] <- sum(predictions[,i] <= interval[j,i] & predictions[,i] > interval[j-1,i])/total.size
  #   }
  # }
  
  df <- expand.grid(bin = 1:bins,
                    name = 1:length(rasters),
                    bootstrap = 1:bootstrap) # make a storage data frame
  df$intensity <- NA # add the outcome 
  df$bin.numeric <- NA # add the outcome 
  
  pb <- progress_bar$new(format = "(:spin) [:bar] :percent [Elapsed time: :elapsedfull || Estimated time remaining: :eta]", total = bootstrap, complete = "=", incomplete = "-", current = ">", clear = FALSE, width = 100)
  
  for (i in 1:bootstrap) { # for each iteration
    sampled <- intersected %>% 
          sample_n(nrow(intersected), replace = T) # resample with replacement to the full dataset size
    for (j in 2:(bins + 1)){ # step through centiles
      for (k in 1:length(rasters)){ # step through predicted surfaces
        true <- sampled[,rasters[k]] <= interval[j,k] & sampled[,rasters[k]] > interval[j-1,k] # check whether each point is in the focal centile
        df[which(
          df$bin == (j-1) &
            df$name == k &
            df$bootstrap == i
        ),"intensity"] <- sum(true, na.rm = T)/(nrow(intersected)) # store the percentage of bootstrapped points in interval, and the proportion of the landscape in that interval
        
        df[which(
          df$bin == (j-1) &
            df$name == k &
            df$bootstrap == i
        ),"bin.numeric"]  <- interval[j,k]
        }
    }
    pb$tick()
  }
  
  df <- df %>% 
      filter(!is.na(bin.numeric)) %>% 
      group_by(name, bootstrap) %>%  
      summarize(measure = cor(bin, intensity, method = "spearman")) %>% 
      mutate(name = factor(name))
  return (df) # return dataframe
}


compare_mean <- function(data = ssf1.train, rstack = deer1.rstack, bootstrap = 100){
  rasters <- names(rstack) # layers of predicted surface
  
  intersected <- data %>% 
    extract_covariates(rstack) %>% 
    drop_na() 
  
  store <- rep(list(NA), length = bootstrap)
  for(i in 1:bootstrap){
    store[[i]] <- intersected %>% 
      sample_n(nrow(intersected), replace = T) %>%
      pivot_longer(all_of(rasters)) %>% 
      mutate(bootstrap = i) %>% 
      group_by(name, bootstrap) %>% 
      summarize(measure = exp(mean(log(value))), .groups = "keep")%>% 
      mutate(name = factor(name))
  }
  
  return (rbindlist(store)) # return dataframe
}

sqrt.both.sides <- function(x) {
  new <- ifelse(x == 0, 0, sqrt(abs(x)) * x/abs(x))
  }
rev.sqrt.both.sides <- function(x) {
  new <- ifelse(x == 0, 0, x^2 * x/abs(x))
  }
transform_both.sqrt.trans_trans <- function() trans_new("both.sqrt.trans", sqrt.both.sides, rev.sqrt.both.sides)
```

### 2 Simulation

#### 2.1 What’s under the hood

Generate 4 rasters.

```
set.seed(100)
grid <- expand.grid(x = 1:100, y = 1:100) # 100 x 100 landscape
grid$z <- 1
r <- raster::rasterFromXYZ(grid)
sim <- simulate_data(r, n = 4, scale = 0.6, sd = 0.1) # spatial scale of 1 for autocorrelation and sd of 0.1
```

```
## Warning: Support for raster package will be deprecated.
## ℹ Use `terra::rast()` to convert data for future compatibility.
```

```
plot(sim)
```

```
index <- setValues(sim$z.1, 1:ncell(sim$z.1)) # store raster index as an alternate raster
names(index) <- "index"

sim <- stack(sim, index) # stack the values and the index

sims <- lapply(c(0.001, 0.01, 0.05, 0.1, 0.2, 0.3, 0.4, 0.6, 0.8, 0.95), function(x){
  simulated <- simulate_data(terra::rast(r), n = 1, scale = x, sd = 0.1)
  simulated
})
stacked <- stack(terra::rast(sims))
names(stacked) <- c(0.001, 0.01, 0.05, 0.1, 0.2, 0.3, 0.4, 0.6, 0.8, 0.95)
levelplot(stacked, layout = c(5, 2))
```

Combine these rasters using some random combo of betas to create our
surface of “good habitat”.

```
set.seed(101)
beta<-runif(4,-1,1) # random betas
sim.values = values(sim) # values from rasters

sur.val <- (beta[1]*sim.values[,1] + beta[2]*sim.values[,2] + beta[3]*sim.values[,3] + beta[4]*sim.values[,4] ) # combine rasters using betas
sim.sur <- setValues(sim[[1]], sur.val) # force values to a raster of "suitability"

#convert suitability to a kernel
sim.sur.kern <- exp(sur.val)
sim.sur.kern <- sim.sur.kern/sum(sim.sur.kern)
sim.sur.kern <- setValues(sim[[1]], sim.sur.kern) # force values to a kernel of "suitability"

plot(sim.sur.kern, main = "Suitability")
```

Use a recent version of a movement model: “How to scale up from
animal movement decisions to spatiotemporal patterns: An approach via
step selection”, J. R. Potts and L. Borger, J Anim Ecol 2023 Vol. 92
Issue 1 Pages 16-29, Accession Number: 36321473 DOI:
10.1111/1365-2656.13832

Here, we have used an animal mover that makes selections for steps
based on the conditions of the cell its moving to and some
autocorrelation based on past movement. Step length (negative), habitat
quality (positive), and turn angle (positive correlation with last step
angle) affect (exponential) probabilty of selection for every cell in
the landscape. By forcing a high penality for large step-lengths
(lambda), we can force our animal mover to be quite local (sampling very
little of the habitat). We add a little noise in step lengths to help
with the issue of perfect raster coordinate selections.

```
set.seed(100)
lambda <- 1 # take shorter steps
kappa <- 1 # use past turn angle more
step_no <- 10000   # number of steps to be simulated

locations <- matrix(NA, nrow = step_no, ncol = 2) # storage matrix

spts <- as.data.frame(rasterToPoints(sim, spatial = TRUE)) # get points from matrix for simulation

# Simulate path
alpha_x = 0 # initial turn angle

for(step in 1:step_no) # go through steps
{
  if(step == 1) {
    newxy<-sample(1:nrow(spts), 1, prob=exp(sur.val)) # if its the first step, derive random start based on suitability
    locations[step,] <- xyFromCell(sim, newxy) # put this cell in storage and jump to the next step
    next
  }
  
  alpha_z <- atan2(spts$y-locations[step-1,2],spts$x-locations[step-1,1]) # calculate difference in turn angle for all cells based on possible steps
  
  unnorm_mk<-exp(-lambda*sqrt((spts$x-locations[step-1,1])^2+(spts$y-locations[step-1,2])^2) + # negatively weighted distance
                   sur.val + # directly weighted suitability surface
                   kappa*cos(alpha_x-alpha_z)) # Positively weighted small differences in turn angle
  
  mk<-unnorm_mk/sum(unnorm_mk) # make the surface a kernel
  
  newxy<-sample(1:nrow(spts), 1, prob=mk) # draw sample
  
  locations[step,] <- xyFromCell(sim, newxy) # store sampled location
  
  alpha_x<-atan2(spts$y-locations[step-1,2],spts$x-locations[step-1,1]) # calculate difference in turn angle based on the step taken
}


# Plot the layer as a raster
prob.kern <- exp(sim.sur)/sum(exp(values(sim.sur)))
plot((prob.kern))

hab50 <- prob.kern>quantile(prob.kern, 0.5) # pull the top 50% of habitat
lines(rasterToPolygons(hab50, dissolve = T), col = "grey50") # highlight the top 10% of habitat visually

lines(locations, col = "black", lwd = 0.25) # plot track
```

Convert movement data to a track object with a little noise for
fitting the step distribution.

```
locations.df <- as.data.frame(locations)
locations.df$t <- as.Date(1:nrow(locations.df), origin = "1970-01-01")
trk <- amt::make_track(locations.df, .x = V1, .y = V2, .t = t)

ssf.dat <- trk %>% 
  mutate(x_ = x_ + runif(nrow(trk), -0.001, 0.001),
         y_ = y_ + runif(nrow(trk), -0.001, 0.001)) %>% 
  steps()
```

We can fit an SSF from our original smaller sample (both in steps and
in spatial extent). Here, we fitting step lengths as a polynomial, and I
am also sampling a lot more andwider the observed step lengths. We do
this with a wider uniform distribution, just taking the min to 0 and max
up by a factor of 1.25.

```
fatter.unif <- original.unif <- fit_distr(ssf.dat$sl_, "unif") # get the original unif distribution
fatter.unif$params$min <- 0 # take the min down
fatter.unif$params$max <- fatter.unif$params$max * 1.25 # take the max up
```

Now we create random steps.

```
set.seed(100)
ssf.dat. <- ssf.dat %>% 
  random_steps(100,
               sl_distr = fatter.unif) %>% # implement the fatter unif distribution for random steps
  extract_covariates(sim) # pull covariates from the initial four rasters
```

Now we can fit the step selection model.

```
issf.fit <- fit_issf(ssf.dat. %>% 
                       filter(!is.na(z.1),
                              !is.na(z.1),
                              !is.na(z.3),
                              !is.na(z.4)) , case_ ~ (z.1 + z.2 + z.3 + z.4) + sl_ + log(sl_) + strata(step_id_), model = T)
summary(issf.fit)
```

```
## Call:
## coxph(formula = Surv(rep(1, 939218L), case_) ~ (z.1 + z.2 + z.3 + 
##     z.4) + sl_ + log(sl_) + strata(step_id_), data = data, model = ..1, 
##     method = "exact")
## 
##   n= 939218, number of events= 9998 
## 
##               coef exp(coef)  se(coef)      z Pr(>|z|)    
## z.1      -0.243406  0.783953  0.024222 -10.05   <2e-16 ***
## z.2      -0.791340  0.453237  0.026104 -30.32   <2e-16 ***
## z.3       0.391724  1.479529  0.024228  16.17   <2e-16 ***
## z.4       0.285778  1.330797  0.020777  13.76   <2e-16 ***
## sl_      -0.284364  0.752493  0.006119 -46.47   <2e-16 ***
## log(sl_) -0.277679  0.757540  0.007447 -37.29   <2e-16 ***
## ---
## Signif. codes:  0 '***' 0.001 '**' 0.01 '*' 0.05 '.' 0.1 ' ' 1
## 
##          exp(coef) exp(-coef) lower .95 upper .95
## z.1         0.7840     1.2756    0.7476    0.8221
## z.2         0.4532     2.2064    0.4306    0.4770
## z.3         1.4795     0.6759    1.4109    1.5515
## z.4         1.3308     0.7514    1.2777    1.3861
## sl_         0.7525     1.3289    0.7435    0.7616
## log(sl_)    0.7575     1.3201    0.7466    0.7687
## 
## Concordance= 0.854  (se = 0.001 )
## Likelihood ratio test= 19648  on 6 df,   p=<2e-16
## Wald test            = 13461  on 6 df,   p=<2e-16
## Score (logrank) test = 33304  on 6 df,   p=<2e-16
```

Now comes our pipeline above. Mock surface -> cells -> cell
data -> fit a standard lRSS. I pause here just to show whats under
the hood. As you will see, there is a raster cell selected, and it is
being compared to all other raster cells. But, as you might see, that
polynomical step distribution makes it sparse! Very few cells beyond
short distances actually matter - most are (presumably) zeros. I have
shown two radii, one that is the 50th centile and one that is the 99th
centile. I have also outlined all cells with a probability > 0.00001.
Sparse, very sparse.

```
set.seed(100)
mock.surface <- create_mock_surface(sim, F, list(x = 1, y = 1))
pred.data <- get_cells(issf.fit, 
                       mock.surface,
                       sim)
cell.data <- get_cell_data(issf.fit, pred.data)

value <- exp(cell.data$lRSS)/sum(exp(cell.data$lRSS))
cell.data$sl_[which(cell.data$sl_ == 0)] <- 0.01

      

selection <- value

prob.kern <- setValues(mock.surface, selection/sum(selection))

# Plot the layer as a raster
plot(prob.kern)
points(cell.data[which(cell.data$sl_ == 0),c("x","y")])

p.0001 <- prob.kern>0.0001
lines(rasterToPolygons(p.0001, dissolve = T), col = "red")

q.50 <- prob.kern>quantile(prob.kern, 0.5)
lines(rasterToPolygons(q.50, dissolve = T))

q.90 <- prob.kern>quantile(prob.kern, 0.9)
lines(rasterToPolygons(q.90, dissolve = T))
```

99% of observed steps occurred less than 6.7 raster units. For this
example, let’s use this as the definition of our neighborhood. Remember
our example above? Here is what this looks like for that specific cell
comparison. It’s not perfect, but it is still includes the parabolic
step pattern! In a real prediction, one might expect this to be a lot
larger.

```
quantile(ssf.dat$sl_, .95)
```

```
##      95% 
## 4.999819
```

```
raster.step <- setValues(mock.surface, cell.data$sl_)
prob.kern <- ifelse(values(raster.step) <= quantile(ssf.dat$sl_, .95), values(prob.kern), NA)
raster.step <- setValues(mock.surface, prob.kern)
raster.step <- trim(raster.step)
plot(raster.step, useRaster = F)
```

We find the neighbors (neighbor\_lookup), then the neighbors for each
cell (neighbor\_finder), then generate comparisons dataframes
(compile\_ssf\_comparisons). I figured out how to make progress bars -
cool! 100x100 landscape in 1 min on a M1 Mac - not bad!

```
neighbors.found <- neighbor_lookup(mock.surface, cell.data) # generates call-up table
sparse.neighbors <- neighbor_finder(issf.fit, cell.data, neighbors.found, distance.override = quantile(ssf.dat$sl_, .95)) # finds neighbors within the 99th percentile of movement
```

```
## [1] "Creating neighbor comparisons"
## [1] "Finding valid comparisons"
## [1] "Splitting cell.data into list"
## [1] "Running comparisons"
```

```
ssf.comparisons <- compile_ssf_comparisons(sparse.neighbors, cell.data) # grabs predictions datasets

ssf.comparisons <- lapply(ssf.comparisons, function(x) {
  list(.for = x$.for, .given = x$.given) # generates for and given comparisons per cell
})
surface <- predict_ssf_comparisons(issf.fit, ssf.comparisons) # makes predictions to generate the A matrix
```

```
## [1] "Estimating probability surface"
## [1] "Compiling probability surface"
## [1] "Making sparse matrix for transitions"
```

We can extract the probability matrix and convert it into a column
stochastic matrix. From that matrix, we can get an eigenvector, take the
real components, and turn it into a probability distribution kernel.
Because we built a sparse matrix, we can use tools that capitalize on
sparse calculations. This retrieves an eigenvector from a 10,000 by
10,000 sparse matrix in less than a second.

```
prob.matrix <- surface$prob.matrix #the A matrix
A <- prob.matrix
d <- eigs(t(A), 1) # sparse matrix eigen decomposition
d.1 <- Re(d$vectors[,1]) # first eigenvector
prob.d <- d.1/sum(d.1) # kernel - though this should sum to one anyway - the decomp just adds a scale that is unhelpful
ssd.prob.raster <- setValues(mock.surface, prob.d) # make raster
par(mfrow=c(1,2),oma=c(0,0,0,2))

plot((sim.sur.kern), main = "Suitability Kernel", zlim=c(0,max(values(ssd.prob.raster))))
plot((ssd.prob.raster), main = "Stable-State Kernel", zlim=c(0,max(values(ssd.prob.raster))))
```

We can also do the traditional but flawed application of an SSF to an
entire surface (not respecting mechanisms, just selection
coefficients)

```
par(mfrow=c(1,2),oma=c(0,0,0,2))

a.data <- pred.data #grab the prediction data we made
a.data$sl_ <- mean(ssf.dat.$sl_) #use the mean step-length for our model
b.data <- a.data[1,]
log.rss <- amt::log_rss(issf.fit, # the model
                          a.data, # the raster data (including missing values)
                          b.data,  # a row of the raster data (excluding missing values)
                          ci = NA)
ssf. <-  exp(log.rss$df$log_rss)/sum(exp(log.rss$df$log_rss)) # take out of link-scale and turn into kernel
ssf.prob.raster <- setValues(mock.surface, ssf.) # make raster
par(mfrow=c(1,2))
plot((sim.sur.kern), main = "Suitability Kernel", zlim=c(0,max(values(ssd.prob.raster))))
plot((ssf.prob.raster), main = "SSF Kernel", zlim=c(0,max(values(ssd.prob.raster))))
```

We can also create a resource selection function for these data

```
par(mfrow=c(1,2),oma=c(0,0,0,2))

set.seed(100)
rsf <- trk %>% 
  random_points() %>% # generic rsf from amt
  extract_covariates(sim) # extract covariates
rsf %>% 
  arrange(case_) %>% 
  ggplot(aes(x = x_, y = y_, col = case_)) + 
  geom_point(alpha = 1, size = 0.5)+
  ylim(0,100) +
  xlim(0,100) + #plot the RSF so we can see the spatial bias implicit in patterns 
  theme_classic() +
  scale_color_scico_d() +
  labs(x = "x", y = "y", color = "Used")
```

```
rsf <- rsf %>% 
  fit_rsf(case_ ~ (z.1 + z.2 + z.3 + z.4), model = T)  # fit the true suitability surface as the model

probabilities <- exp(predict(rsf$model, newdata = pred.data))/(1+exp(predict(rsf$model, newdata = pred.data))) # take the rsf output into probabilities

probabilities.kern <- probabilities/sum(probabilities) # turn into kernel
rsf.prob.raster <- setValues(mock.surface, probabilities.kern) # make matrix
par(mfrow=c(1,2))
plot((sim.sur.kern), main = "Suitability Kernel", zlim=c(0,max(values(ssd.prob.raster))))
plot((rsf.prob.raster), main = "RSF Kernel", zlim=c(0,max(values(ssd.prob.raster))))
```

#### 2.2 Scaling up

##### 2.2.1 10 animals

```
id1 <- c(0.05, -.05, .05, -.05) #generalist
id2 <- id1*2
id3 <- id1*3
id4 <- id1*4
id5 <- id1*5
id6 <- id1*6
id7 <- id1*7
id8 <- id1*8
id9 <- id1*9
id10 <- id1*10

animals <- list(id1, id2, id3, id4, id5, id6, id7, id8, id9, id10)
```

##### 2.2.2 10 landscapes

```
set.seed(100)
grid <- expand.grid(x = 1:100, y = 1:100) # 100 x 100 landscape
grid$z <- 1
r <- raster::rasterFromXYZ(grid)

sims <- lapply(rep(c(0.001, 0.01, 0.05, 0.1, 0.2, 0.3, 0.4, 0.6, 0.8, 0.95), each=5), function(x){
  simulated <- simulate_data(terra::rast(r), n = 4, scale = x, sd = 0.1)
  names(simulated) <- c("z.1", "z.2", "z.3", "z.4")
  simulated
})

plot(terra::rast(sims[c(11,21,31)]))
```

```
index <- setValues(sims[[1]]$z.1, 1:ncell(sims[[1]]$z.1)) # store raster index as an alternate raster
names(index) <- "index"

sim.index <- index
```

##### 2.2.3 Converting rasters to suitability

```
sur.val <- lapply(animals, function(x){
  out <- lapply(sims, function(y){
    sim.values = values(y)
    sur.val <- (x[1]*sim.values[,1] + x[2]*sim.values[,2] + x[3]*sim.values[,3] + x[4]*sim.values[,4] )
    sur.val
  })
  out
})


sur.val. <- lapply(sur.val, function(x) do.call("cbind", x[1:50]))
sur.val. <- do.call("cbind", sur.val.[1:10])
sur.val. <- data.frame(sur.val.)

sur.val <- lapply(1:500, function(x){
  setValues(sims[[1]][[1]], sur.val.[,x])
})

names(sur.val) <- paste0(rep(paste0("ID",1:10," scale = "), each = 50), rep(rep(c(0.001, 0.01, 0.05, 0.1, 0.2, 0.3, 0.4, 0.6, 0.8, 0.95), each = 5), 10))

suitability <- lapply(1:500, function(x){
  sim.sur.kern <- exp(sur.val.[,x])/sum(exp(sur.val.[,x]))
  setValues(sims[[1]][[1]], sim.sur.kern)
})

names(suitability) <- paste0(rep(paste0("ID",1:10," scale = "), each = 50), rep(rep(c(0.001, 0.01, 0.05, 0.1, 0.2, 0.3, 0.4, 0.6, 0.8, 0.95), each = 5), 10))

plot(terra::rast(suitability[c(1,21,46,201,221,246,451,471,496)]))
```

##### 2.2.4 Simulate function based on params and surfaces

```
# simulate_movements <- function(x, lambda = 1, kappa = 1, step_no = 1E4, teleport = NULL){
# 
#   locations <- matrix(NA, nrow = step_no, ncol = 2) # storage matrix
# 
#   spts <- as.data.frame(rasterToPoints(x, spatial = TRUE)) # get points from matrix for simulation
# 
#   # Simulate path
#   alpha_x = 0 # initial turn angle
# 
#   grid <- expand.grid(x = 1:100, y = 1:100) # 100 x 100 landscape
#   grid$z <- 1
#   sim <- raster::rasterFromXYZ(grid)
# 
#   if(is.null(teleport)) {
#     for(step in 1:step_no) # go through steps
#       {
#       if(step == 1) {
#         newxy<-sample(1:nrow(spts), 1, prob=exp(spts$z.1)) # if its the first step, derive random start based on suitability
#         locations[step,] <- xyFromCell(sim, newxy) # put this cell in storage and jump to the next step
#         next
#       }
# 
#       alpha_z <- atan2(spts$y-locations[step-1,2],spts$x-locations[step-1,1]) # calculate difference in turn angle for all cells based on possible steps
# 
#       unnorm_mk<-exp(-lambda*sqrt((spts$x-locations[step-1,1])^2+(spts$y-locations[step-1,2])^2) + # negatively weighted distance
#                          spts$z.1 + # directly weighted suitability surface
#                          kappa*cos(alpha_x-alpha_z)) # Positively weighted small differences in turn angle
# 
#       mk<-unnorm_mk/sum(unnorm_mk) # make the surface a kernel
# 
#       newxy<-sample(1:nrow(spts), 1, prob=mk) # draw sample
# 
#       locations[step,] <- xyFromCell(sim, newxy) # store sampled location
# 
#       alpha_x<-atan2(spts$y-locations[step-1,2],spts$x-locations[step-1,1]) # calculate difference in turn angle based on the step taken
#     }
#   } else {
#     for(step in 1:step_no) # go through steps
#       {
#       if(step %in% seq(1,step_no, by = teleport)) { # randomly restart the track every 1000 steps
#         newxy<-sample(1:nrow(spts), 1, prob=exp(spts$z.1)) # if its the first or restart steps, derive random starts based on suitability
#         locations[step,] <- xyFromCell(sim, newxy) # put this cell in storage and jump to the next step
#         alpha_x <- 0 # reset turn angle to 0
#         next
#       }
# 
#       alpha_z <- atan2(spts$y-locations[step-1,2],spts$x-locations[step-1,1]) # calculate difference in turn angle for all cells based on possible steps
# 
#       unnorm_mk<-exp(-lambda*sqrt((spts$x-locations[step-1,1])^2+(spts$y-locations[step-1,2])^2) + # negatively weighted distance
#                          spts$z.1 + # directly weighted suitability surface
#                          kappa*cos(alpha_x-alpha_z)) # Positively weighted small differences in turn angle
# 
#       mk<-unnorm_mk/sum(unnorm_mk) # make the surface a kernel
# 
#       newxy<-sample(1:nrow(spts), 1, prob=mk) # draw sample
# 
#       locations[step,] <- xyFromCell(sim, newxy) # store sampled location
# 
#       alpha_x<-atan2(spts$y-locations[step-1,2],spts$x-locations[step-1,1]) # calculate difference in turn angle based on the step taken
#     }
#   }
# 
#   return(locations)
# }
```

##### 2.2.5 Simulate in parallel

```
# library(parallel)
# detectCores()
# 
# cl <- makeCluster(24)
# clusterExport(cl, "simulate_movements")
# clusterExport(cl, "sur.val")
# clusterEvalQ(cl, {
#   library(raster)
#   set.seed(12345)
# })
# locations.sim <- parLapply(cl, 1:500, function(x) {simulate_movements(sur.val[[x]])} )
# 
# stopCluster(cl)
# 
# 
# cl <- makeCluster(24)
# clusterExport(cl, "simulate_movements")
# clusterExport(cl, "sur.val")
# clusterEvalQ(cl, {
#   library(raster)
#   set.seed(12345)
# })
# locations.sim.samplea <- parLapply(cl, 1:24, function(x) {simulate_movements(sur.val[[x]], step_no = 5E5, teleport = 1E3)} )
# locations.sim.sampleb <- parLapply(cl, 25:48, function(x) {simulate_movements(sur.val[[x]], step_no = 5E5, teleport = 1E3)} )
# locations.sim.samplec <- parLapply(cl, 49:72, function(x) {simulate_movements(sur.val[[x]], step_no = 5E5, teleport = 1E3)} )
# locations.sim.sampled <- parLapply(cl, 73:96, function(x) {simulate_movements(sur.val[[x]], step_no = 5E5, teleport = 1E3)} )
# locations.sim.samplee <- parLapply(cl, 97:120, function(x) {simulate_movements(sur.val[[x]], step_no = 5E5, teleport = 1E3)} )
# locations.sim.samplef <- parLapply(cl, 121:144, function(x) {simulate_movements(sur.val[[x]], step_no = 5E5, teleport = 1E3)} )
# locations.sim.sampleg <- parLapply(cl, 145:168, function(x) {simulate_movements(sur.val[[x]], step_no = 5E5, teleport = 1E3)} )
# locations.sim.sampleh <- parLapply(cl, 169:192, function(x) {simulate_movements(sur.val[[x]], step_no = 5E5, teleport = 1E3)} )
# locations.sim.samplei <- parLapply(cl, 193:216, function(x) {simulate_movements(sur.val[[x]], step_no = 5E5, teleport = 1E3)} )
# locations.sim.samplej <- parLapply(cl, 217:240, function(x) {simulate_movements(sur.val[[x]], step_no = 5E5, teleport = 1E3)} )
# locations.sim.samplek <- parLapply(cl, 241:264, function(x) {simulate_movements(sur.val[[x]], step_no = 5E5, teleport = 1E3)} )
# locations.sim.samplel <- parLapply(cl, 265:288, function(x) {simulate_movements(sur.val[[x]], step_no = 5E5, teleport = 1E3)} )
# locations.sim.samplem <- parLapply(cl, 289:312, function(x) {simulate_movements(sur.val[[x]], step_no = 5E5, teleport = 1E3)} )
# locations.sim.samplen <- parLapply(cl, 313:336, function(x) {simulate_movements(sur.val[[x]], step_no = 5E5, teleport = 1E3)} )
# locations.sim.sampleo <- parLapply(cl, 337:360, function(x) {simulate_movements(sur.val[[x]], step_no = 5E5, teleport = 1E3)} )
# locations.sim.samplep <- parLapply(cl, 361:384, function(x) {simulate_movements(sur.val[[x]], step_no = 5E5, teleport = 1E3)} )
# locations.sim.sampleq <- parLapply(cl, 385:408, function(x) {simulate_movements(sur.val[[x]], step_no = 5E5, teleport = 1E3)} )
# locations.sim.sampler <- parLapply(cl, 409:432, function(x) {simulate_movements(sur.val[[x]], step_no = 5E5, teleport = 1E3)} )
# locations.sim.samples <- parLapply(cl, 433:456, function(x) {simulate_movements(sur.val[[x]], step_no = 5E5, teleport = 1E3)} )
# locations.sim.samplet <- parLapply(cl, 457:480, function(x) {simulate_movements(sur.val[[x]], step_no = 5E5, teleport = 1E3)} )
# locations.sim.sampleu <- parLapply(cl, 481:500, function(x) {simulate_movements(sur.val[[x]], step_no = 5E5, teleport = 1E3)} )
# 
# stopCluster(cl)
# 
# 
# 
# locations.sim.sample <- rep(list(NA), 500)
# 
# locations.sim.sample[1:24] <-locations.sim.samplea
# locations.sim.sample[25:48] <-locations.sim.sampleb
# locations.sim.sample[49:72] <-locations.sim.samplec
# locations.sim.sample[73:96] <-locations.sim.sampled
# locations.sim.sample[97:120] <-locations.sim.samplee
# locations.sim.sample[121:144] <-locations.sim.samplef
# locations.sim.sample[145:168] <-locations.sim.sampleg
# locations.sim.sample[169:192] <-locations.sim.sampleh
# locations.sim.sample[193:216] <-locations.sim.samplei
# locations.sim.sample[217:240] <-locations.sim.samplej
# locations.sim.sample[241:264] <-locations.sim.samplek
# locations.sim.sample[265:288] <-locations.sim.samplel
# locations.sim.sample[289:312] <-locations.sim.samplem
# locations.sim.sample[313:336] <-locations.sim.samplen
# locations.sim.sample[337:360] <-locations.sim.sampleo
# locations.sim.sample[361:384] <-locations.sim.samplep
# locations.sim.sample[385:408] <-locations.sim.sampleq
# locations.sim.sample[409:432] <-locations.sim.sampler
# locations.sim.sample[433:456] <-locations.sim.samples
# locations.sim.sample[457:480] <-locations.sim.samplet
# locations.sim.sample[481:500] <-locations.sim.sampleu
# 
# names(locations.sim) <- paste0(rep(paste0("ID",1:10,"."), each = 50), rep(rep(1:10, each = 5), 10), rep(paste0("It",1:5,"."),100))
# names(locations.sim.sample) <- paste0(rep(paste0("ID",1:10,"."), each = 50), rep(rep(1:10, each = 5), 10), rep(paste0("It",1:5,"."),100))

# saveRDS(locations.sim, "Data and Intermediate RDS files/locations.sim.1.RDS")
# saveRDS(locations.sim.sample, "Data and Intermediate RDS files/locations.sim.sample.1.RDS")
locations.sim <- readRDS("Data and Intermediate RDS files/locations.sim.1.RDS")

# locations.sim.sample <- readRDS("Data and Intermediate RDS files/locations.sim.sample.1.RDS")

# locations.sim.sample.a <- locations.sim.sample[1:100]
# locations.sim.sample.b <- locations.sim.sample[101:200]
# locations.sim.sample.c <- locations.sim.sample[201:300]
# locations.sim.sample.d <- locations.sim.sample[301:400]
# locations.sim.sample.e <- locations.sim.sample[401:500]
# 
# saveRDS(locations.sim.sample.a, "Data and Intermediate RDS files/locations.sim.sample.a.RDS")
# saveRDS(locations.sim.sample.b, "Data and Intermediate RDS files/locations.sim.sample.b.RDS")
# saveRDS(locations.sim.sample.c, "Data and Intermediate RDS files/locations.sim.sample.c.RDS")
# saveRDS(locations.sim.sample.d, "Data and Intermediate RDS files/locations.sim.sample.d.RDS")
# saveRDS(locations.sim.sample.e, "Data and Intermediate RDS files/locations.sim.sample.e.RDS")

# this is just to get around git size restrictions
locations.sim.sample.a <- readRDS("Data and Intermediate RDS files/locations.sim.sample.a.RDS")
locations.sim.sample.b <- readRDS("Data and Intermediate RDS files/locations.sim.sample.b.RDS")
locations.sim.sample.c <- readRDS("Data and Intermediate RDS files/locations.sim.sample.c.RDS")
locations.sim.sample.d <- readRDS("Data and Intermediate RDS files/locations.sim.sample.d.RDS")
locations.sim.sample.e <- readRDS("Data and Intermediate RDS files/locations.sim.sample.e.RDS")

locations.sim.sample <- c(locations.sim.sample.a, locations.sim.sample.b, locations.sim.sample.c, locations.sim.sample.d, locations.sim.sample.e)
```

##### 2.2.6 SSF fitting

```
# set.seed(12345)
# ssf.fit <- rep(list(NA), 500)
# for(i in 1:500){
#   
#   locations.df <- as.data.frame(locations.sim[[i]])
#   locations.df$t <- as.Date(1:nrow(locations.df), origin = "1970-01-01")
#   trk <- amt::make_track(locations.df, .x = V1, .y = V2, .t = t)
# 
#   ssf.dat <- trk %>%
#     mutate(x_ = x_ + runif(nrow(trk), -0.001, 0.001),
#            y_ = y_ + runif(nrow(trk), -0.001, 0.001)) %>%
#     steps()
# 
#   fatter.unif <- original.unif <- fit_distr(ssf.dat$sl_, "unif") # get the original unif distribution
#   fatter.unif$params$min <- 0 # take the min to 0
#   fatter.unif$params$max <-  fatter.unif$params$max*1.25 # bump up the max
# 
# 
# 
#   landscape.to.use <- i %% 50
#   landscape.to.use <- ifelse(landscape.to.use == 0, 50, landscape.to.use)
# 
#   sim <- sims[[landscape.to.use]]
# 
#   ssf.dat. <- ssf.dat %>%
#     random_steps(100,
#                  sl_distr = fatter.unif) %>% # implement the fatter uniform distribution for random steps
#     extract_covariates(sim)
# 
#   ssf.fit[[i]] <- fit_issf(ssf.dat. %>%
#                        filter(!is.na(z.1),
#                               !is.na(z.1),
#                               !is.na(z.3),
#                               !is.na(z.4)) , case_ ~ (z.1 + z.2 + z.3 + z.4) + (sl_) + log(sl_) + strata(step_id_), model = T)
#   print(i)
# }
```

##### 2.2.7 SSD Predictions

```
# ssd <- lapply(1:500, function(x){
# 
#   landscape.to.use <- x %% 50
#   landscape.to.use <- ifelse(landscape.to.use == 0, 50, landscape.to.use)
#   sim <- sims[[landscape.to.use]]
# 
#   issf.fit <- ssf.fit[[x]]
#   mock.surface <- create_mock_surface(sim, F, list(x = 1, y = 1))
#   pred.data <- get_cells(issf.fit,
#                          mock.surface,
#                          sim)
#   cell.data <- get_cell_data(issf.fit, pred.data)
#   neighbors.found <- neighbor_lookup(mock.surface, cell.data) # generates call-up table
#   
#   locations.df <- as.data.frame(locations.sim[[x]])
#   locations.df$t <- as.Date(1:nrow(locations.df), origin = "1970-01-01")
#   trk <- amt::make_track(locations.df, .x = V1, .y = V2, .t = t)
# 
#   ssf.dat <- trk %>%
#     mutate(x_ = x_ + runif(nrow(trk), -0.001, 0.001),
#            y_ = y_ + runif(nrow(trk), -0.001, 0.001)) %>%
#     steps()
#   
#   sparse.neighbors <- neighbor_finder(issf.fit, cell.data, neighbors.found, quantile = 0.95, distance.override = quantile(ssf.dat$sl_, 0.95)) # finds neighbors within the 99th percentile of movement
#   ssf.comparisons <- compile_ssf_comparisons(sparse.neighbors, cell.data) # grabs predictions datasets
# 
#   ssf.comparisons <- lapply(ssf.comparisons, function(x) {
#     list(.for = x$.for, .given = x$.given) # generates for and given comparisons per cell
#   })
#   surface <- predict_ssf_comparisons(issf.fit, ssf.comparisons) # makes predictions to generate the A matrix
#   prob.matrix <- surface$prob.matrix #the A matrix
#   A <- prob.matrix
#   d <- eigs(t(A), 1) # sparse matrix eigen decomposition
#   d.1 <- Re(d$vectors[,1]) # first eigenvector
#   prob.d <- d.1/sum(d.1)
#   prob.raster <- setValues(mock.surface, prob.d)
#   prob.raster
# })
```

##### 2.2.8 SSF predictions

```
# set.seed(12345)
# ssf <- lapply(1:500, function(x){
# 
#   landscape.to.use <- x %% 50
#   landscape.to.use <- ifelse(landscape.to.use == 0, 50, landscape.to.use)
#   sim <- sims[[landscape.to.use]]
# 
#   issf.fit <- ssf.fit[[x]]
#   mock.surface <- create_mock_surface(sim, F, list(x = 1, y = 1))
#   pred.data <- get_cells(issf.fit,
#                          mock.surface,
#                          sim)
#   a.data <- pred.data #grab the prediction data we made
# 
#   locations.df <- as.data.frame(locations.sim[[x]])
#   locations.df$t <- as.Date(1:nrow(locations.df), origin = "1970-01-01")
#   trk <- amt::make_track(locations.df, .x = V1, .y = V2, .t = t)
# 
#   ssf.dat <- trk %>%
#     mutate(x_ = x_ + runif(nrow(trk), -0.001, 0.001),
#            y_ = y_ + runif(nrow(trk), -0.001, 0.001)) %>%
#     steps()
# 
#   a.data$sl_ <- mean(ssf.dat$sl_) #use the mean step-length for our model
#   b.data <- a.data[1,]
#   log.rss <- amt::log_rss(issf.fit, # the model
#                             a.data, # the raster data (including missing values)
#                             b.data,  # a row of the raster data (excluding missing values)
#                             ci = NA)
#   ssf. <-  exp(log.rss$df$log_rss)/sum(exp(log.rss$df$log_rss)) # take out of link-scale and turn into kernel
#   prob.raster <- setValues(mock.surface, ssf.)
# 
#   return(prob.raster)
# })
```

##### 2.2.9 RSF predictions

```
# set.seed(12345)
# rsf.model <- lapply(1:500, function(x){
# 
#   landscape.to.use <- x %% 50
#   landscape.to.use <- ifelse(landscape.to.use == 0, 50, landscape.to.use)
#   sim <- sims[[landscape.to.use]]
# 
#   locations.df <- as.data.frame(locations.sim[[x]])
#   locations.df$t <- as.Date(1:nrow(locations.df), origin = "1970-01-01")
#   trk <- amt::make_track(locations.df, .x = V1, .y = V2, .t = t)
# 
#   rsf <- trk %>%
#     random_points() %>% # generic rsf from amt
#     extract_covariates(sim) # extract covariates
# 
#   rsf <- rsf %>%
#     fit_rsf(case_ ~ (z.1 + z.2 + z.3 + z.4), model = T)
# 
#   return(rsf)
# })
# 
# rsf <- lapply(1:500, function(x){
# 
#   landscape.to.use <- x %% 50
#   landscape.to.use <- ifelse(landscape.to.use == 0, 50, landscape.to.use)
#   sim <- sims[[landscape.to.use]]
# 
#   locations.df <- as.data.frame(locations.sim[[x]])
#   locations.df$t <- as.Date(1:nrow(locations.df), origin = "1970-01-01")
#   trk <- amt::make_track(locations.df, .x = V1, .y = V2, .t = t)
# 
#   issf.fit <- ssf.fit[[x]]
#   mock.surface <- create_mock_surface(sim, F, list(x = 1, y = 1))
#   pred.data <- get_cells(issf.fit,
#                          mock.surface,
#                          sim)
# 
#   probabilities <- exp(predict(rsf.model[[x]]$model, newdata = pred.data))/(1+exp(predict(rsf.model[[x]]$model, newdata = pred.data)))
# 
#   probabilities.kern <- probabilities/sum(probabilities)
#   prob.raster <- setValues(mock.surface, probabilities.kern)
# 
#   return(prob.raster)
# })
```

```
# saveRDS(locations.sim, "locations.sim.multi.rds")
# saveRDS(locations.sim.sample, "locations.sim.sample.multi.rds")
# 
# saveRDS(ssf.fit, "ssf.fit.multi.rds")
# saveRDS(rsf.model, "rsf.model.multi.rds")
# 
# saveRDS(ssd, "ssd.pred.rds")
# saveRDS(ssf, "ssf.pred.rds")
# saveRDS(rsf, "rsf.pred.rds")

# locations.sim <- readRDS("locations.sim.multi.rds")
# locations.sim.sample <- readRDS("locations.sim.sample.multi.rds")
# 
# ssf.fit <- readRDS("ssf.fit.multi.rds")
# rsf.model <- readRDS("rsf.model.multi.rds")
# 
ssd <- readRDS("Data and Intermediate RDS files/ssd.pred.rds") # landscape prediction kernels
ssf <- readRDS("Data and Intermediate RDS files/ssf.pred.rds") # landscape prediction kernels
rsf <- readRDS("Data and Intermediate RDS files/rsf.pred.rds") # landscape prediction kernels
```

##### 2.2.10 Merging Estimates

```
# ssf.est <- lapply(1:500, function(x){
#   a <- summary(ssf.fit[[x]]$model)$coefficients
#   
#   df <- data.frame(term = c("z.1","z.2","z.3","z.4"),
#                    coef = a[1:4,1],
#                    se = a[1:4,3],
#                    group = x)
#   df
# })
# 
# rsf.est <- lapply(1:500, function(x){
#   a <- summary(rsf.model[[x]]$model)$coefficients
#   
#   df <- data.frame(term = c("z.1","z.2","z.3","z.4"),
#                    coef = a[2:5,1],
#                    se = a[2:5,2],
#                    group = x)
#   df
# })
# 
# 
# ssf.est <- rbindlist(ssf.est)
# rsf.est <- rbindlist(rsf.est)
# 
# ssf.est$model = "SSF"
# rsf.est$model = "RSF"
# 
# saveRDS(ssf.est, "ssf.est.rds")
# saveRDS(rsf.est, "rsf.est.rds")

ssf.est <- readRDS("Data and Intermediate RDS files/ssf.est.rds") # model coefficients
rsf.est <- readRDS("Data and Intermediate RDS files/rsf.est.rds") # model coefficients

estimates <- rbind(ssf.est, rsf.est)

estimates <- estimates %>% mutate(
  animal = case_when(
    group %in% 1:50 ~ 1,
    group %in% 51:100 ~ 2,
    group %in% 101:150 ~ 3,
    group %in% 151:200 ~ 4,
    group %in% 201:250 ~ 5,
    group %in% 251:300 ~ 6,
    group %in% 301:350 ~ 7,
    group %in% 351:400 ~ 8,
    group %in% 401:450 ~ 9,
    group %in% 451:500 ~ 10,
  ),
  het = case_when(
    group %% 50 %in% 1:5 ~ 0.001,
    group %% 50 %in% 6:10 ~ 0.01,
    group %% 50 %in% 11:15 ~ 0.05,
    group %% 50 %in% 16:20 ~ 0.1,
    group %% 50 %in% 21:25 ~ 0.2,
    group %% 50 %in% 26:30 ~ 0.3,
    group %% 50 %in% 31:35 ~ 0.4,
    group %% 50 %in% 36:40 ~ 0.6,
    group %% 50 %in% 41:45 ~ 0.8,
    group %% 50 %in% c(46:49,0) ~ 0.95
  ),
  iter = case_when(
    group %% 50 %in% seq(1,500, by = 5) ~ 1,
    group %% 50 %in% seq(2,500, by = 5) ~ 2,
    group %% 50 %in% seq(3,500, by = 5) ~ 3,
    group %% 50 %in% seq(4,500, by = 5) ~ 4,
    group %% 50 %in% seq(5,500, by = 5) ~ 5,
  ),
  true.effect = case_when(
    term == "z.1" ~ 0.05 * animal,
    term == "z.2" ~ -0.05 * animal,
    term == "z.3" ~ 0.05 * animal,
    term == "z.4" ~ -0.05 * animal
  )
)
```

###### 2.2.10.1 Estimate error

```
estimate.plot <- estimates %>% 
  mutate(lower = (coef - 1.96*se)* true.effect/abs(true.effect),
         upper = (coef + 1.96*se)* true.effect/abs(true.effect),
         coef = coef * true.effect/abs(true.effect)) %>% 
  arrange(sample(1:n())) %>% 
  mutate(animal = factor(animal, levels = 1:10, labels = paste0("ID",1:10), ordered = T)) %>%
  ggplot(aes(x = factor(het), y = (coef-abs(true.effect)), ymin = (lower-abs(true.effect)), ymax = (upper-abs(true.effect)), color = animal, group = animal)) +
  geom_hline(yintercept = 0) +
  geom_pointrange(alpha = 0.25, position = position_dodge(0.5)) +
  geom_smooth(method = "gam", se = F, alpha = 0.5) +
  facet_grid(.~model) +
  scale_color_viridis_d() +
  scale_shape_manual(values = c(15:18)) +
  theme_classic() +
  labs(x = "Gaussian Process Scale\n(Heterogeneity)",
       y = "Coefficient Estimate Error \n(Absolute Estimate and 95% CI - Truth)",
       color = "Generalist (1) to\nSpecialist (10)") +
  theme(axis.text.x = element_text(angle = 45, vjust = 1, hjust = 1))

estimate.plot.reduced <- estimates %>% 
  mutate(lower = (coef - 1.96*se)* true.effect/abs(true.effect),
         upper = (coef + 1.96*se)* true.effect/abs(true.effect),
         coef = coef * true.effect/abs(true.effect)) %>% 
  arrange(sample(1:n())) %>% 
  # mutate(animal = factor(animal, levels = 1:10, labels = paste0("ID",1:10), ordered = T)) %>%
  mutate(gen.spec = case_when(
    animal == 2 ~ "Strong Generalist",
    animal == 4 ~ "Slight Generalist",
    animal == 6 ~ "Slight Specialist",
    animal == 8 ~ "Strong Specialist"
  )) %>%  
  filter(!is.na(gen.spec)) %>% 
  mutate(gen.spec = factor(gen.spec, levels = c("Strong Generalist","Slight Generalist","Slight Specialist","Strong Specialist"), labels = c("Strong Generalist (2)","Slight Generalist (4)","Slight Specialist (6)","Strong Specialist (8)"), ordered = T))%>%  
  ggplot(aes(x = factor(het), y = (coef-abs(true.effect)), ymin = (lower-abs(true.effect)), ymax = (upper-abs(true.effect)), color = gen.spec, group = gen.spec)) +
  geom_hline(yintercept = 0) +
  geom_pointrange(position = position_dodge(0.9), alpha = 0.75, size = 0.25) +
  geom_smooth(method = "gam", se = F, position = position_dodge(0.9)) +
  facet_grid(.~model) +
  scale_color_scico_d(palette = "glasgow") +
  scale_shape_manual(values = c(15:18)) +
  theme_classic() +
  labs(x = "Gaussian Process Scale\n(Heterogeneity)",
       y = "Coefficient Estimate Error \n(Absolute Estimate and 95% CI - Truth)",
       color = "") +
  theme(axis.text.x = element_text(angle = 45, vjust = 1, hjust = 1),
        legend.position = "inside",
        legend.position.inside = c(0.75, 0.6),
        legend.background = element_blank())

estimate.plot.reduced
```

```
## `geom_smooth()` using formula = 'y ~ s(x, bs = "cs")'
```

```
## Warning: The following aesthetics were dropped during statistical transformation: ymin
## and ymax.
## ℹ This can happen when ggplot fails to infer the correct grouping structure in
##   the data.
## ℹ Did you forget to specify a `group` aesthetic or to convert a numerical
##   variable into a factor?
## The following aesthetics were dropped during statistical transformation: ymin
## and ymax.
## ℹ This can happen when ggplot fails to infer the correct grouping structure in
##   the data.
## ℹ Did you forget to specify a `group` aesthetic or to convert a numerical
##   variable into a factor?
```

```
## Warning: `position_dodge()` requires non-overlapping x intervals.
## `position_dodge()` requires non-overlapping x intervals.
```

```
# ggsave("Plots/Full.Sim.Est.Error.png", estimate.plot, dpi = 600, width = 8, height = 4)
# ggsave("Plots/Text.Sim.Est.Error.svg", estimate.plot.reduced, dpi = 600, width = 6, height = 4)
```

###### 2.2.10.2 Truth contained in estimate

```
estimate.contained <- estimates %>% 
  mutate(lower = coef - 1.96*se,
         upper = coef + 1.96*se) %>% 
  mutate(inside = true.effect >= lower & true.effect <= upper) %>% 
  group_by(animal, het, model, iter) %>% 
  summarize(prop.within = sum(inside)/(4)) %>% 
  group_by(animal, het, model) %>% 
  summarize(prop.within.mu = mean(prop.within),
            prop.within.max = max(prop.within),
            prop.within.min = min(prop.within)) %>% 
  ggplot(aes(x= factor(het), y = prop.within.mu, ymin = prop.within.min, ymax = prop.within.max, fill = model,color = model, group = model)) +
  geom_bar(position = position_dodge(1), stat = "identity", alpha = 0.5, color = NA) +
  geom_errorbar(position = position_dodge(1), stat = "identity", width = 0.25) +
  facet_grid(animal~.) +
  scale_fill_scico_d(begin = 0.75, direction = -1) +
  scale_color_scico_d(begin = 0.75, direction = -1) +
  theme_classic() +
  labs(x = "Gaussian Process Scale (Heterogeneity)",
       y = "Mean (Min and Max) Proportion of Selection Coefficient\nEstimates contained within 95% CI in 5 Replicates",
       fill = "Inferential\nModel",
       color = "Inferential\nModel")
```

```
## `summarise()` has grouped output by 'animal', 'het', 'model'. You can override
## using the `.groups` argument.
## `summarise()` has grouped output by 'animal', 'het'. You can override using the
## `.groups` argument.
```

```
estimate.contained
```

```
estimate.contained.reduced <- estimates %>% 
  mutate(lower = coef - 1.96*se,
         upper = coef + 1.96*se) %>% 
  mutate(inside = true.effect >= lower & true.effect <= upper) %>% 
  group_by(animal, het, model, iter) %>% 
  summarize(prop.within = sum(inside)/(4)) %>% 
  group_by(animal, het, model) %>% 
  summarize(prop.within.mu = mean(prop.within),
            prop.within.max = max(prop.within),
            prop.within.min = min(prop.within)) %>% 
  mutate(gen.spec = case_when(
    animal == 2 ~ "Strong Generalist",
    animal == 4 ~ "Slight Generalist",
    animal == 6 ~ "Slight Specialist",
    animal == 8 ~ "Strong Specialist"
  )) %>%  
  filter(!is.na(gen.spec)) %>% 
  mutate(gen.spec = factor(gen.spec, levels = c("Strong Generalist","Slight Generalist","Slight Specialist","Strong Specialist"), labels = c("Strong Generalist (2)","Slight Generalist (4)","Slight Specialist (6)","Strong Specialist (8)"), ordered = T))%>% 
  ggplot(aes(x= factor(het), y = prop.within.mu, ymin = prop.within.min, ymax = prop.within.max, fill = model,color = model, group = model)) +
  geom_bar(position = position_dodge(1), stat = "identity", alpha = 0.5, color = NA) +
  geom_errorbar(position = position_dodge(1), stat = "identity", width = 0.25) +
  facet_grid(gen.spec~.) +
  scale_fill_scico_d(begin = 0.75, direction = -1) +
  scale_color_scico_d(begin = 0.75, direction = -1) +
  theme_classic() +
  labs(x = "Gaussian Process Scale (Heterogeneity)",
       y = "Mean (Min and Max) Proportion of Selection Coefficient\nEstimates contained within 95% CI in 5 Replicates",
       fill = "Inferential\nModel",
       color = "Inferential\nModel")
```

```
## `summarise()` has grouped output by 'animal', 'het', 'model'. You can override
## using the `.groups` argument.
## `summarise()` has grouped output by 'animal', 'het'. You can override using the
## `.groups` argument.
```

```
estimate.contained.reduced
```

```
# ggsave("Plots/Full.Sim.Est.Cont.png", estimate.contained, dpi = 600, width = 8, height = 8)
```

##### 2.2.11 Merging Predictions

```
spts <- as.data.frame(sim.index, xy = TRUE) # get points from matrix for simulation

# data.frames <- lapply(1:500, function(x){
#   merged <- merge(spts, as.data.frame(locations.sim.sample[[x]]), by.x = c("x", "y"), by.y = c("V1","V2")) # merge map with use
# 
#   tallied <- merged %>%
#     group_by(index) %>%
#     summarize(n = n()) # observed selected steps per cell
#   not.used <- spts$index[which(!spts$index %in% tallied$index)] # cells are not used, meaning zero counts
#   not.used.df <- tibble(index = not.used, n = 0) # we can make a mock df of these unused points
# 
#   tallied <- rbind(tallied, not.used.df) # now we need to add these used and unused locations together for a full landscape depiction
# 
#   tallied <- tallied %>%
#     mutate(value.true.kernel = values(suitability[[x]])[tallied$index],
#            value.true.original = (sur.val.[,x])[tallied$index],
#            prop.obs = n/(5e5),
#            ssd = values(ssd[[x]])[tallied$index],
#            ssf = values(ssf[[x]])[tallied$index],
#            rsf = values(rsf[[x]])[tallied$index],
#            group = x)
#   print(x)
#   tallied
# })
# 
# g <- rbindlist(data.frames)
# g <- g %>% mutate(
#   animal = case_when(
#     group %in% 1:50 ~ "ID1",
#     group %in% 51:100 ~ "ID2",
#     group %in% 101:150 ~ "ID3",
#     group %in% 151:200 ~ "ID4",
#     group %in% 201:250 ~ "ID5",
#     group %in% 251:300 ~ "ID6",
#     group %in% 301:350 ~ "ID7",
#     group %in% 351:400 ~ "ID8",
#     group %in% 401:450 ~ "ID9",
#     group %in% 451:500 ~ "ID10",
#   ),
#   het = case_when(
#     group %% 50 %in% 1:5 ~ 0.001,
#     group %% 50 %in% 6:10 ~ 0.01,
#     group %% 50 %in% 11:15 ~ 0.05,
#     group %% 50 %in% 16:20 ~ 0.1,
#     group %% 50 %in% 21:25 ~ 0.2,
#     group %% 50 %in% 26:30 ~ 0.3,
#     group %% 50 %in% 31:35 ~ 0.4,
#     group %% 50 %in% 36:40 ~ 0.6,
#     group %% 50 %in% 41:45 ~ 0.8,
#     group %% 50 %in% c(46:49,0) ~ 0.95
#   ),
#   iter = case_when(
#     group %% 50 %in% seq(1,500, by = 5) ~ 1,
#     group %% 50 %in% seq(2,500, by = 5) ~ 2,
#     group %% 50 %in% seq(3,500, by = 5) ~ 3,
#     group %% 50 %in% seq(4,500, by = 5) ~ 4,
#     group %% 50 %in% seq(0,500, by = 5) ~ 5,
#   )
# )

# saveRDS(g, "predictions.G.RDS")
# g <- readRDS("Data and Intermediate RDS files/predictions.G.RDS")

# g.a <- g[1:1000000,]
# g.b <- g[1000001:2000000,]
# g.c <- g[2000001:3000000,]
# g.d <- g[3000001:4000000,]
# g.e <- g[4000001:5000000,]
# 
# saveRDS(g.a, "Data and Intermediate RDS files/g.a.RDS")
# saveRDS(g.b, "Data and Intermediate RDS files/g.b.RDS")
# saveRDS(g.c, "Data and Intermediate RDS files/g.c.RDS")
# saveRDS(g.d, "Data and Intermediate RDS files/g.d.RDS")
# saveRDS(g.e, "Data and Intermediate RDS files/g.e.RDS")

#to get around git restrictions
g.a <- readRDS("Data and Intermediate RDS files/g.a.RDS")
g.b <- readRDS("Data and Intermediate RDS files/g.b.RDS")
g.c <- readRDS("Data and Intermediate RDS files/g.c.RDS")
g.d <- readRDS("Data and Intermediate RDS files/g.d.RDS")
g.e <- readRDS("Data and Intermediate RDS files/g.e.RDS")

g <- rbind(g.a, g.b, g.c, g.d, g.e)


g <- merge(g, spts, by = "index")
```

###### 2.2.11.1 Mapping Simulations

```
ha <- g %>% 
  mutate(gen.spec = case_when(
    animal == "ID2" ~ "Strong Generalist",
    animal == "ID4" ~ "Slight Generalist",
    animal == "ID6" ~ "Slight Specialist",
    animal == "ID8" ~ "Strong Specialist"
  )) %>%  
  filter(!is.na(gen.spec)) %>% 
  mutate(gen.spec = factor(gen.spec, levels = c("Strong Generalist","Slight Generalist","Slight Specialist","Strong Specialist"), labels = c("Strong Generalist (2)","Slight Generalist (4)","Slight Specialist (6)","Strong Specialist (8)"), ordered = T))%>% 
  filter(animal %in% c("ID2","ID4","ID6", "ID8") & het == 0.05) %>% 
  pivot_longer(c(prop.obs, value.true.kernel, ssd, ssf, rsf)) %>% 
  group_by(animal, name, gen.spec) %>% 
  arrange(-value) %>% 
  filter(iter == 1) %>% 
  mutate(animal = factor(animal, levels = paste0("ID",1:10), ordered = T)) %>% 
  mutate(name = factor(name, 
                       levels = c("prop.obs", "value.true.kernel", "ssd","ssf","rsf"),
                       labels = c("Simulated Use", "Selection", "SSD","SSF","RSF"))) %>% 
  ggplot(aes(x,y,fill = value)) +
  geom_raster() +
  facet_grid(name~gen.spec) +
  scale_fill_scico(palette = "vikO", name = "P(Use) Kernel", trans = "sqrt") + 
  labs(title = "Scale = 0.05")

hb <- g %>% 
  mutate(gen.spec = case_when(
    animal == "ID2" ~ "Strong Generalist",
    animal == "ID4" ~ "Slight Generalist",
    animal == "ID6" ~ "Slight Specialist",
    animal == "ID8" ~ "Strong Specialist"
  )) %>%  
  filter(!is.na(gen.spec)) %>% 
  mutate(gen.spec = factor(gen.spec, levels = c("Strong Generalist","Slight Generalist","Slight Specialist","Strong Specialist"), labels = c("Strong Generalist (2)","Slight Generalist (4)","Slight Specialist (6)","Strong Specialist (8)"), ordered = T))%>% 
  filter(animal %in% c("ID2","ID4","ID6", "ID8") & het == 0.2) %>% 
  pivot_longer(c(prop.obs, value.true.kernel, ssd, ssf, rsf)) %>% 
  group_by(animal, name, gen.spec) %>% 
  arrange(-value) %>% 
  filter(iter == 1) %>% 
  mutate(animal = factor(animal, levels = paste0("ID",1:10), ordered = T)) %>% 
  mutate(name = factor(name, 
                       levels = c("prop.obs", "value.true.kernel", "ssd","ssf","rsf"),
                       labels = c("Simulated Use", "Selection", "SSD","SSF","RSF"))) %>% 
  ggplot(aes(x,y,fill = value)) +
  geom_raster() +
  facet_grid(name~gen.spec) +
  scale_fill_scico(palette = "vikO", name = "P(Use) Kernel", trans = "sqrt")+ 
  labs(title = "Scale = 0.2")

hc <- g %>% 
  mutate(gen.spec = case_when(
    animal == "ID2" ~ "Strong Generalist",
    animal == "ID4" ~ "Slight Generalist",
    animal == "ID6" ~ "Slight Specialist",
    animal == "ID8" ~ "Strong Specialist"
  )) %>%  
  filter(!is.na(gen.spec)) %>% 
  mutate(gen.spec = factor(gen.spec, levels = c("Strong Generalist","Slight Generalist","Slight Specialist","Strong Specialist"), labels = c("Strong Generalist (2)","Slight Generalist (4)","Slight Specialist (6)","Strong Specialist (8)"), ordered = T)) %>% 
  filter(animal %in% c("ID2","ID4","ID6", "ID8") & het == 0.6) %>% 
  pivot_longer(c(prop.obs, value.true.kernel, ssd, ssf, rsf)) %>% 
  group_by(animal, name, gen.spec) %>% 
  arrange(-value) %>% 
  filter(iter == 1) %>% 
  mutate(animal = factor(animal, levels = paste0("ID",1:10), ordered = T)) %>% 
  mutate(name = factor(name, 
                       levels = c("prop.obs", "value.true.kernel", "ssd","ssf","rsf"),
                       labels = c("Simulated Use", "Selection", "SSD","SSF","RSF"))) %>% 
  ggplot(aes(x,y,fill = value)) +
  geom_raster() +
  facet_grid(name~gen.spec) +
  scale_fill_scico(palette = "vikO", name = "P(Use) Kernel", trans = "sqrt")+ 
  labs(title = "Scale = 0.6")

ggarrange(ha + theme_classic(), hb + theme_classic(), hc + theme_classic(), legend = "bottom", nrow = 1, labels = c("A", "B", "C"))
```

```
# ggsave("Plots/simulation.maps.png", dpi = 600, width = 18, height = 7)
```

```
g %>% 
  filter(animal == "ID6" &
           iter == 5 &
           het == 0.6) %>%  
  pivot_longer(c(prop.obs, value.true.kernel, ssd, ssf, rsf)) %>% 
  mutate(animal = factor(animal, levels = paste0("ID",1:10), ordered = T)) %>% 
  mutate(name = factor(name, 
                       levels = c("prop.obs", "value.true.kernel", "ssd","ssf","rsf"),
                       labels = c("Simulated Use", "Selection", "SSD","SSF","RSF"))) %>% 
  ggplot(aes(x,y,fill = value)) +
  geom_raster() +
  facet_wrap(~name, nrow = 2) +
  coord_equal()+
  scale_fill_viridis_c(option = "H") +
  theme_void()
```

```
# ggsave("Plots/simulation.maps.exp.svg", dpi = 600, width = 12, height = 7)
```

###### 2.2.11.2 Mapping 50% of “Habitat”

```
ha <- g %>% 
  mutate(gen.spec = case_when(
    animal == "ID2" ~ "Strong Generalist",
    animal == "ID4" ~ "Slight Generalist",
    animal == "ID6" ~ "Slight Specialist",
    animal == "ID8" ~ "Strong Specialist"
  )) %>%  
  filter(!is.na(gen.spec)) %>% 
  mutate(gen.spec = factor(gen.spec, levels = c("Strong Generalist","Slight Generalist","Slight Specialist","Strong Specialist"), labels = c("Strong Generalist (2)","Slight Generalist (4)","Slight Specialist (6)","Strong Specialist (8)"), ordered = T))%>% 
  filter(animal %in% c("ID2","ID4","ID6", "ID8") & het == 0.05) %>% 
  pivot_longer(c(prop.obs, value.true.kernel, ssd, ssf, rsf)) %>% 
  group_by(animal, name, gen.spec) %>% 
  arrange(-value) %>% 
  filter(iter == 1) %>% 
  mutate(animal = factor(animal, levels = paste0("ID",1:10), ordered = T)) %>% 
  mutate(cdf = cumsum(value)) %>% 
  mutate(name = factor(name, 
                       levels = c("prop.obs", "value.true.kernel", "ssd","ssf","rsf"),
                       labels = c("Simulated Use", "Selection", "SSD","SSF","RSF"))) %>% 
  ggplot(aes(x,y,fill = cdf < 0.5)) +
  geom_raster() +
  facet_grid(name~gen.spec) +
  scale_fill_scico_d(name = "In top 50% of\nCDF of P(Use)")+ 
  labs(title = "Scale = 0.05")

hb <- g %>% 
  mutate(gen.spec = case_when(
    animal == "ID2" ~ "Strong Generalist",
    animal == "ID4" ~ "Slight Generalist",
    animal == "ID6" ~ "Slight Specialist",
    animal == "ID8" ~ "Strong Specialist"
  )) %>%  
  filter(!is.na(gen.spec)) %>% 
  mutate(gen.spec = factor(gen.spec, levels = c("Strong Generalist","Slight Generalist","Slight Specialist","Strong Specialist"), labels = c("Strong Generalist (2)","Slight Generalist (4)","Slight Specialist (6)","Strong Specialist (8)"), ordered = T))%>% 
  filter(animal %in% c("ID2","ID4","ID6", "ID8") & het == 0.2) %>% 
  pivot_longer(c(prop.obs, value.true.kernel, ssd, ssf, rsf)) %>% 
  group_by(animal, name, gen.spec) %>% 
  arrange(-value) %>% 
  filter(iter == 1) %>% 
  mutate(animal = factor(animal, levels = paste0("ID",1:10), ordered = T)) %>% 
  mutate(cdf = cumsum(value)) %>% 
  mutate(name = factor(name, 
                       levels = c("prop.obs", "value.true.kernel", "ssd","ssf","rsf"),
                       labels = c("Simulated Use", "Selection", "SSD","SSF","RSF"))) %>% 
  ggplot(aes(x,y,fill = cdf < 0.5)) +
  geom_raster() +
  facet_grid(name~gen.spec)+
  scale_fill_scico_d(name = "In top 50% of\nCDF of P(Use)")+ 
  labs(title = "Scale = 0.2")

hc <- g %>% 
  mutate(gen.spec = case_when(
    animal == "ID2" ~ "Strong Generalist",
    animal == "ID4" ~ "Slight Generalist",
    animal == "ID6" ~ "Slight Specialist",
    animal == "ID8" ~ "Strong Specialist"
  )) %>%  
  filter(!is.na(gen.spec)) %>% 
  mutate(gen.spec = factor(gen.spec, levels = c("Strong Generalist","Slight Generalist","Slight Specialist","Strong Specialist"), labels = c("Strong Generalist (2)","Slight Generalist (4)","Slight Specialist (6)","Strong Specialist (8)"), ordered = T)) %>% 
  filter(animal %in% c("ID2","ID4","ID6", "ID8") & het == 0.6) %>% 
  pivot_longer(c(prop.obs, value.true.kernel, ssd, ssf, rsf)) %>% 
  group_by(animal, name, gen.spec) %>% 
  arrange(-value) %>% 
  filter(iter == 1) %>% 
  mutate(animal = factor(animal, levels = paste0("ID",1:10), ordered = T)) %>% 
  mutate(cdf = cumsum(value)) %>% 
  mutate(name = factor(name, 
                       levels = c("prop.obs", "value.true.kernel", "ssd","ssf","rsf"),
                       labels = c("Simulated Use", "Selection", "SSD","SSF","RSF"))) %>% 
  ggplot(aes(x,y,fill = cdf < 0.5)) +
  geom_raster() +
  facet_grid(name~gen.spec)+
  scale_fill_scico_d(name = "In top 50% of\nCDF of P(Use)")+ 
  labs(title = "Scale = 0.6")

ggarrange(ha, hb, hc, legend = "bottom", nrow = 1, labels = c("A", "B", "C"))
```

```
# ggsave("Plots/discrete.maps.png", dpi = 600, width = 18, height = 7)
```

###### 2.2.11.3 Proportion Habitat (>= 0.5 CDF of P(Use))

```
atta<-g %>% 
  pivot_longer(c(prop.obs, value.true.kernel, ssd, ssf, rsf)) %>% 
  group_by(animal, name, het,iter) %>% 
  arrange(-value) %>% 
  mutate(animal = factor(animal, levels = paste0("ID",1:10), ordered = T)) %>% 
  mutate(cdf = cumsum(value)) %>% 
  summarize(total.50 = sum(cdf <= 0.5)) %>% 
  pivot_wider(names_from = name, values_from = total.50) %>% 
  pivot_longer(c(value.true.kernel, ssd, ssf, rsf))
```

```
## `summarise()` has grouped output by 'animal', 'name', 'het'. You can override
## using the `.groups` argument.
```

```
atta %>% 
  # mutate(name = factor(name, 
  #                      levels = c("prop.obs", "value.true.kernel", "ssd","ssf","rsf"),
  #                      labels = c("Simulated Use", "Selection", "SSD","SSF","RSF"))) %>% 
  # pivot_wider(names_from = name, values_from = total.50) %>%  
  # pivot_longer(c("Simulated Use", "Selection", "SSD","SSF","RSF")) %>% 
  group_by(het, animal, name) %>% 
  mutate(mean = mean((value/prop.obs)-1),
         min = min((value/prop.obs)-1),
         max = max((value/prop.obs)-1)) %>% 
  mutate(name = factor(name, 
                       levels = c("prop.obs", "value.true.kernel", "ssd","ssf","rsf"),
                       labels = c("Simulated Use", "Selection", "SSD","SSF","RSF"), ordered = T)) %>%
  ggplot(aes(x = factor(het), y = mean, ymin = min, ymax = max, color = name, group = paste(name))) +
  geom_hline(yintercept=0)+
  geom_path(size = 0.5, alpha = 0.5, position = position_dodge(0.01))+
  geom_pointrange(position= position_dodge(0.01)) +
  theme(axis.ticks = element_blank(),
        axis.text.x = element_text(angle = 0.45, vjust = 1, hjust = 1)) +
  facet_wrap(~animal, ncol = 5) +
  theme_classic()+
  scale_color_scico_d(begin = 0.25) +
  labs(x = "Gaussian Process Scale (Heterogeneity)",
       y = "Predicted 50% P(U) Area Realtive to Observed Use\n(Mean and Range in 5 ",
       color = "Prediction") +
  theme(axis.text.x = element_text(angle = 45, vjust = 1, hjust = 1)) +
  scale_y_continuous(labels = scales::percent, breaks = c(-.5, -.25, 0, .25, .50, .75, 1, 1.25, 1.5))
```

```
## Warning: Using `size` aesthetic for lines was deprecated in ggplot2 3.4.0.
## ℹ Please use `linewidth` instead.
## This warning is displayed once every 8 hours.
## Call `lifecycle::last_lifecycle_warnings()` to see where this warning was
## generated.
```

```
# ggsave("Plots/simulation.area.full.png", dpi = 600, width = 10, height = 7)

atta %>% 
  mutate(gen.spec = case_when(
    animal == "ID2" ~ "Strong Generalist",
    animal == "ID4" ~ "Slight Generalist",
    animal == "ID6" ~ "Slight Specialist",
    animal == "ID8" ~ "Strong Specialist"
  )) %>%  
  filter(!is.na(gen.spec)) %>% 
  mutate(gen.spec = factor(gen.spec, levels = c("Strong Generalist","Slight Generalist","Slight Specialist","Strong Specialist"), labels = c("Strong Generalist (2)","Slight Generalist (4)","Slight Specialist (6)","Strong Specialist (8)"), ordered = T))  %>% 
  group_by(het, gen.spec, name) %>% 
  mutate(mean = mean((value/prop.obs)-1),
         min = min((value/prop.obs)-1),
         max = max((value/prop.obs)-1)) %>% 
  mutate(name = factor(name, 
                       levels = c("prop.obs", "value.true.kernel", "ssd","ssf","rsf"),
                       labels = c("Simulated Use", "Selection", "SSD","SSF","RSF"), ordered = T)) %>%
  ggplot(aes(x = factor(het), y = mean, ymin = min, ymax = max, color = name, group = paste(name))) +
  geom_hline(yintercept=0)+
  geom_path(size = 0.5, alpha = 0.5,position= position_dodge(0.1))+
  geom_pointrange(position= position_dodge(0.1)) +
  theme(axis.ticks = element_blank(),
        axis.text.x = element_text(angle = 0.45, vjust = 1, hjust = 1)) +
  facet_grid(~gen.spec, scales = "free_y") +
  theme_classic()+
  scale_color_scico_d(begin = 0.25) +
  labs(x = "Gaussian Process Scale (Heterogeneity)",
       y = "Minimum Predicted 50% P(U) Area Relative to Observed Use\n(Mean and Range in 5 Replicates",
       color = "Prediction Kernel") +
  theme(axis.text.x = element_text(angle = 45, vjust = 1, hjust = 1)) +
  scale_y_continuous(labels = scales::percent)
```

```
# ggsave("Plots/simulation.area.svg", dpi = 600, width = 10, height = 4)
```

###### 2.2.11.4 Case Matching of Habitat

```
wide.hab <- g %>% 
  pivot_longer(c(prop.obs, value.true.kernel, ssd, ssf, rsf)) %>% 
  group_by(animal, name, het,iter) %>% 
  arrange(-value) %>% 
  mutate(animal = factor(animal, levels = paste0("ID",1:10), ordered = T)) %>% 
  mutate(cdf = cumsum(value)) %>% 
  mutate(hab = cdf <= 0.5) %>% 
  dplyr::select(index, group, animal, het, x, y, name, hab,iter) %>% 
  pivot_wider(names_from = name,
              values_from = hab) %>% 
  pivot_longer(c(ssd, ssf, rsf)) %>% 
  pivot_longer(c(prop.obs, value.true.kernel), names_to = "truth", values_to = "values.truth") %>% 
  mutate(case = case_when(
    values.truth == T & value == T ~ "True Positive",
    values.truth == F & value == F ~ "True Negative",
    values.truth == F & value == T ~ "False Positive",
    values.truth == T & value == F ~ "False Negative")) %>% 
  mutate(gen.spec = case_when(
    animal == "ID2" ~ "Strong Generalist",
    animal == "ID4" ~ "Slight Generalist",
    animal == "ID6" ~ "Slight Specialist",
    animal == "ID8" ~ "Strong Specialist"
  )) %>%  
  # filter(!is.na(gen.spec)) %>% 
  mutate(gen.spec = factor(gen.spec, levels = c("Strong Generalist","Slight Generalist","Slight Specialist","Strong Specialist"), labels = c("Strong Generalist (2)","Slight Generalist (4)","Slight Specialist (6)","Strong Specialist (8)"), ordered = T)) %>% 
  mutate(name = factor(name, 
                       levels = c("prop.obs", "value.true.kernel", "ssd","ssf","rsf"),
                       labels = c("Simulated Use", "Selection", "SSD","SSF","RSF")))
  


accuracy <- wide.hab %>% 
  filter(truth == "prop.obs") %>% 
  group_by(animal, name, het, iter, gen.spec) %>% 
  summarize(accuracy = sum(values.truth == T & value == T | values.truth == F & value == F)/n()) %>% 
  group_by(animal, name, het, gen.spec) %>% 
  summarize(mean = mean(accuracy),
            lower =min(accuracy),
            upper =max(accuracy))
```

```
## `summarise()` has grouped output by 'animal', 'name', 'het', 'iter'. You can
## override using the `.groups` argument.
## `summarise()` has grouped output by 'animal', 'name', 'het'. You can override
## using the `.groups` argument.
```

```
accuracy %>% 
  # filter(!is.na(gen.spec)) %>% 
  ggplot(aes(x = factor(het), y = mean, ymin = lower, ymax = upper, color = name, group = name)) +
  geom_path() +
  geom_pointrange() +
  facet_wrap(.~animal, nrow = 2)+
  scale_color_scico_d(begin = 0.5) + 
  theme_classic() +
  labs(x = "Gaussian Process Scale (Heterogeneity)",
       y = "Accuracy of Top 50% Cumulative P(U) Classification",
       color = "Prediction Kernel") +
  theme(axis.text.x = element_text(angle = 45, vjust = 1, hjust = 1))
```

```
# ggsave("Plots/accuracy_classification.png", dpi = 600, width = 9, height = 5)

wide.hab <- wide.hab %>% 
  mutate(case = factor(case, levels = c("True Positive", "True Negative", "False Positive", "False Negative"), ordered = T))
ha <- wide.hab %>% 
  filter(animal %in% c("ID2","ID4", "ID6", "ID8") & truth == "prop.obs", iter == 1) %>% 
  filter(het == 0.1) %>% 
  ggplot(aes(x, y, fill = case)) +
  geom_raster() +
  facet_grid(name~gen.spec)+
  scale_fill_scico_d(name = "Classification Case", palette = "lipari") +
  theme_classic() +
  labs(title = "Scale = 0.1")

hb <- wide.hab %>% 
  filter(animal %in% c("ID2","ID4", "ID6", "ID8") & truth == "prop.obs", iter == 1) %>% 
  filter(het == 0.6) %>% 
  ggplot(aes(x, y, fill = case)) +
  geom_raster() +
  facet_grid(name~gen.spec)+
  scale_fill_scico_d(name = "Classification Case", palette = "lipari") +
  theme_classic()+
  labs(title = "Scale = 0.6")

ggarrange(ha, hb, legend = "bottom", nrow = 1, labels = c("A", "B"), common.legend = T)
```

```
# ggsave("Plots/discrete.maps.case.png", dpi = 600, width = 12, height = 5)
```

```
wide.hab %>% 
  filter(animal %in% c("ID2", "ID8") &
           iter == 5 &
           het == 0.6) %>%  
  filter(truth== "prop.obs") %>% 
  mutate(case = factor(case, levels = c("True Positive", "True Negative", "False Positive", "False Negative"), ordered = T)) %>% 
   mutate(gen.spec = case_when(
    animal == "ID2" ~ "Strong Generalist",
    animal == "ID4" ~ "Slight Generalist",
    animal == "ID6" ~ "Slight Specialist",
    animal == "ID8" ~ "Strong Specialist"
  )) %>%  
  # filter(!is.na(gen.spec)) %>% 
  mutate(gen.spec = factor(gen.spec, levels = c("Strong Generalist","Slight Generalist","Slight Specialist","Strong Specialist"), labels = c("Strong Generalist (2)","Slight Generalist (4)","Slight Specialist (6)","Strong Specialist (8)"), ordered = T)) %>% 
  ggplot(aes(x,y,fill = case)) +
  geom_raster() +
  scale_fill_scico_d(name = "Classification Case", palette = "lipari") +
  theme_classic() +
  facet_grid(name~gen.spec) +
  coord_equal() +
  theme_void()
```

```
# ggsave("Plots/simulation.maps.exp1.svg", dpi = 600, width = 6, height = 6)
```

###### 2.2.11.5 Kernel prediction heterogeneity

```
kernel.hetero.all <- g %>% 
  mutate(animal = factor(animal, levels = paste0("ID",1:10), ordered = T)) %>%
  mutate(specialism = factor(animal, levels = paste0("ID",1:10), labels = 1:10, ordered = T)) %>% 
  filter(iter == 1) %>% 
  dplyr::select(index, animal, specialism, het, group, prop.obs, value.true.kernel) %>% 
  pivot_longer(c(prop.obs, value.true.kernel)) %>% 
  mutate(name = factor(name, levels = c("prop.obs", "value.true.kernel"), labels = c("Simulated Use", "Selection Kernel"))) %>% 
  group_by(name, animal, specialism, het) %>% 
  summarize(median = median(value),
            lower = quantile(value,0.025),
            upper = quantile(value,0.975),
            low.50 = quantile(value,0.25),
            upr.50 = quantile(value,0.75)) %>%
  ggplot(aes(x = factor(het), y = median, color = name)) +
  geom_linerange(aes(ymin = lower, ymax = upper), position = position_dodge(0.5)) +
  geom_linerange(aes(ymin = low.50, ymax = upr.50), position = position_dodge(0.5), linewidth = 1) +
  geom_point(, position = position_dodge(0.5)) +
  scale_color_scico_d() + 
  scale_y_log10() +
  facet_grid(.~factor(specialism)) +
  theme_classic() +
  labs(color = "Kernel Type", x = "Gaussian Process Scale (Heterogeneity)",
       y = "Median Probability of Use (50% & 95% Quantiles)")+
  theme(axis.text.x = element_text(angle = 45, vjust = 1, hjust = 1))
```

```
## `summarise()` has grouped output by 'name', 'animal', 'specialism'. You can
## override using the `.groups` argument.
```

```
kernel.hetero.all
```

```
## Warning in scale_y_log10(): log-10 transformation introduced infinite values.
```

```
# ggsave("Plots/KernelHetall.png", kernel.hetero.all, dpi = 600, width = 12, height = 6)


kernel.hetero <- g %>% 
  filter(iter == 1) %>% 
  filter(animal %in% c("ID2","ID4","ID6","ID8"),
         het %in% c(0.01,0.2,0.6)) %>% 
  mutate(gen.spec = case_when(
    animal == "ID2" ~ "Strong Generalist",
    animal == "ID4" ~ "Slight Generalist",
    animal == "ID6" ~ "Slight Specialist",
    animal == "ID8" ~ "Strong Specialist"
  )) %>%    
  dplyr::select(index, gen.spec, het, group, prop.obs, value.true.kernel) %>% 
  pivot_longer(c(prop.obs, value.true.kernel)) %>% 
  mutate(gen.spec = factor(gen.spec, levels = c("Strong Generalist","Slight Generalist","Slight Specialist","Strong Specialist"), labels = c("Strong Generalist (2)","Slight Generalist (4)","Slight Specialist (6)","Strong Specialist (8)"), ordered = T))%>% 
  mutate(name = factor(name, levels = c("prop.obs", "value.true.kernel"), labels = c("Simulated Use", "Selection Kernel"))) %>% 
  group_by(name, gen.spec, het) %>% 
  summarize(median = median(value),
            lower = quantile(value,0.025),
            upper = quantile(value,0.975),
            low.50 = quantile(value,0.25),
            upr.50 = quantile(value,0.75)) %>%
  ggplot(aes(x = factor(het), y = median, color = name)) +
  geom_linerange(aes(ymin = lower, ymax = upper), position = position_dodge(0.5)) +
  geom_linerange(aes(ymin = low.50, ymax = upr.50), position = position_dodge(0.5), linewidth = 1) +
  geom_point(, position = position_dodge(0.5)) +
  scale_color_scico_d() + 
  scale_y_log10() +
  facet_grid(.~factor(gen.spec)) +
  theme_classic() +
  labs(color = "Kernel Type", x = "Gaussian Process Scale (Heterogeneity)",
       y = "Median Probability of Use (50% & 95% Quantiles)")+
  theme(axis.text.x = element_text(angle = 45, vjust = 1, hjust = 1))
```

```
## `summarise()` has grouped output by 'name', 'gen.spec'. You can override using
## the `.groups` argument.
```

```
kernel.hetero
```

```
# ggsave("Plots/KernelHet.png", kernel.hetero, dpi = 600, width = 8, height = 5)
```

```
kernel.hetero <- g %>% 
  dplyr::select(index, animal, het, group, prop.obs, value.true.kernel, iter) %>% 
  pivot_longer(c(prop.obs, value.true.kernel)) %>% 
  mutate(name = factor(name, levels = c("prop.obs", "value.true.kernel"), labels = c("Simulated Use", "Selection Kernel"))) %>% 
  group_by(name, animal, het, iter) %>% 
  summarize(shannon = -sum(ifelse(value == 0, 0, (value)*log((value))))) %>% 
  group_by(name, animal, het) %>% 
  summarize(mean = mean(shannon))
```

```
## `summarise()` has grouped output by 'name', 'animal', 'het'. You can override
## using the `.groups` argument.
## `summarise()` has grouped output by 'name', 'animal'. You can override using
## the `.groups` argument.
```

```
kernel.hetero %>% 
  mutate(specialism = factor(animal, levels = paste0("ID",1:10), labels = 1:10, ordered = T)) %>%   
  mutate(name = factor(name, levels = c("Selection Kernel", "Simulated Use"), ordered = T)) %>%  
  ggplot(aes(x = factor(het), y = specialism, fill = mean)) +
  geom_raster() +
  scale_fill_viridis_c(option = "H", direction = -1) +
  facet_grid(.~name) +
  theme_classic() +
  labs(y = "Animal Selection Strength (Multiplier)",
       x = "Gaussian Process Scale (Heterogeneity)",
       fill = "Mean\nShannon\nEntropy") +
  coord_equal()+
  theme(axis.text.x = element_text(angle = 45, vjust = 1, hjust = 1))
```

```
# ggsave("Plots/simulation.het.heat.svg", dpi = 600, width = 6.5, height = 3)

g %>% 
  dplyr::select(index, animal, het, group, prop.obs, value.true.kernel, iter) %>% 
  pivot_longer(c(prop.obs, value.true.kernel)) %>% 
  mutate(name = factor(name, levels = c("prop.obs", "value.true.kernel"), labels = c("Simulated Use", "Selection Kernel"))) %>% 
  group_by(name, animal, het, iter) %>% 
  filter(iter == 5, animal == "ID8" & het == 0.6 | animal == "ID2" & het == 0.01) %>% 
  ggplot(aes(x = value, fill = name)) +
  geom_density(alpha = 0.2, color = NA) +
  scale_x_sqrt() +
  scale_y_sqrt() +
  facet_grid(animal~.) +
  theme_classic() +
  scale_color_scico_d(labels = c("Simulated Use","Selection Kernel"), palette = "vik") +
  scale_fill_scico_d(labels = c("Simulated Use","Selection Kernel"), palette = "vik") +
  labs(x = "Relative Probability of Use",
       y = "Count",
       color = "Prob.\nType",
       fill = "Prob.\nType")
```

```
# ggsave("Plots/simulation.het.density.svg", dpi = 600, width = 5, height = 3)
```

```
g %>% 
  dplyr::select(index, animal, het, group, prop.obs, value.true.kernel, iter, x, y) %>% 
  mutate(error = (value.true.kernel - prop.obs)/value.true.kernel) %>% 
  mutate(gen.spec = factor(animal, levels = c("ID2", "ID8"), labels = c("Strong Generalist (2)","Strong Specialist (8)"), ordered = T)) %>% 
  mutate(scale = factor(het, levels = c(0.01, 0.1, 0.3, 0.6), labels = paste("Scale =", c(0.01, 0.1, 0.3, 0.6)), ordered = T)) %>% 
  # mutate(name = factor(name, levels = c("Selection Kernel", "Simulated Use"), ordered = T)) %>%  
  filter(het %in% c(0.01, 0.1, 0.3, 0.6) &
           animal %in% paste0("ID",c(2, 8)) &
           iter == 5) %>% 
  ggplot(aes(x = x, y = y, fill = error, z = error)) +
  geom_raster() +
  coord_equal() +
  geom_contour(breaks = c(0), h = c(0.001,0.001), color = "black", size = 0.1) +
  facet_grid(gen.spec~scale) +
  scale_fill_scico(palette = "vik", midpoint = 0, #breaks = c(-0.0012, -0.0008, -0.0004, -0.0001 , 0.000001, 0.0001), 
                   name = expression(frac("Selection - Use","Selection")),labels = scales::label_percent()) +
  theme_void()
```

```
## Warning in geom_contour(breaks = c(0), h = c(0.001, 0.001), color = "black", :
## Ignoring unknown parameters: `h`
```

```
## Warning: The following aesthetics were dropped during statistical transformation: fill.
## ℹ This can happen when ggplot fails to infer the correct grouping structure in
##   the data.
## ℹ Did you forget to specify a `group` aesthetic or to convert a numerical
##   variable into a factor?
## The following aesthetics were dropped during statistical transformation: fill.
## ℹ This can happen when ggplot fails to infer the correct grouping structure in
##   the data.
## ℹ Did you forget to specify a `group` aesthetic or to convert a numerical
##   variable into a factor?
## The following aesthetics were dropped during statistical transformation: fill.
## ℹ This can happen when ggplot fails to infer the correct grouping structure in
##   the data.
## ℹ Did you forget to specify a `group` aesthetic or to convert a numerical
##   variable into a factor?
## The following aesthetics were dropped during statistical transformation: fill.
## ℹ This can happen when ggplot fails to infer the correct grouping structure in
##   the data.
## ℹ Did you forget to specify a `group` aesthetic or to convert a numerical
##   variable into a factor?
## The following aesthetics were dropped during statistical transformation: fill.
## ℹ This can happen when ggplot fails to infer the correct grouping structure in
##   the data.
## ℹ Did you forget to specify a `group` aesthetic or to convert a numerical
##   variable into a factor?
## The following aesthetics were dropped during statistical transformation: fill.
## ℹ This can happen when ggplot fails to infer the correct grouping structure in
##   the data.
## ℹ Did you forget to specify a `group` aesthetic or to convert a numerical
##   variable into a factor?
## The following aesthetics were dropped during statistical transformation: fill.
## ℹ This can happen when ggplot fails to infer the correct grouping structure in
##   the data.
## ℹ Did you forget to specify a `group` aesthetic or to convert a numerical
##   variable into a factor?
## The following aesthetics were dropped during statistical transformation: fill.
## ℹ This can happen when ggplot fails to infer the correct grouping structure in
##   the data.
## ℹ Did you forget to specify a `group` aesthetic or to convert a numerical
##   variable into a factor?
```

```
# ggsave("Plots/simulation.diff.per.svg", dpi = 600, width = 10, height = 5)
```

###### 2.2.11.6 Kernel prediction entropy (Shannon Entropy)

```
shannon <- g %>% 
  mutate(animal = factor(animal, levels = paste0("ID",1:10), ordered = T)) %>%
  mutate(specialism = factor(animal, levels = paste0("ID",1:10), labels = 1:10, ordered = T)) %>%
  pivot_longer(c(prop.obs,ssd, value.true.kernel, ssf, rsf)) %>% 
  group_by(name,specialism, het,iter) %>% 
  summarize(entropy = sum(ifelse(value == 0, 0, (value)*log(1/(value))))) %>% 
  group_by(name,specialism, het) %>% 
  summarize(mean = mean(entropy),
            min = min(entropy),
            max = max(entropy)) %>% 
  mutate(name = factor(name, 
                       levels = c("prop.obs", "value.true.kernel", "ssd","ssf","rsf"),
                       labels = c("Simulated Use", "True Selection", "SSD","SSF","RSF"))) %>% 
  ggplot(aes(x = factor(het), y = mean, ymin = min, ymax = max, color = name, group = paste(name))) +
  geom_path(position = position_dodge(0.2)) +
  geom_pointrange(position = position_dodge(0.2)) +
  facet_wrap(~specialism, ncol = 5) +
  labs(x = "Gaussian Process Scale (Heterogeneity)",
       y = "Shannon Entropy\n(Mean and Range in 5 Replicates)",
       color = "Probability Kernel") +
  scale_color_scico_d() +
  theme_classic() +
  theme(axis.text.x = element_text(angle = 45, vjust = 1, hjust = 1))
```

```
## `summarise()` has grouped output by 'name', 'specialism', 'het'. You can
## override using the `.groups` argument.
## `summarise()` has grouped output by 'name', 'specialism'. You can override
## using the `.groups` argument.
```

```
shannon
```

```
# ggsave("Plots/shannonentropyall.png", shannon, dpi = 600, width = 12, height = 6)

shannon.reduced <- g %>% 
  mutate(gen.spec = case_when(
    animal == "ID2" ~ "Strong Generalist",
    animal == "ID4" ~ "Slight Generalist",
    animal == "ID6" ~ "Slight Specialist",
    animal == "ID8" ~ "Strong Specialist"
  )) %>%  
  filter(!is.na(gen.spec)) %>% 
  mutate(gen.spec = factor(gen.spec, levels = c("Strong Generalist","Slight Generalist","Slight Specialist","Strong Specialist"), labels = c("Strong Generalist (2)","Slight Generalist (4)","Slight Specialist (6)","Strong Specialist (8)"), ordered = T)) %>% 
  pivot_longer(c(prop.obs,ssd, value.true.kernel, ssf, rsf)) %>% 
  group_by(name,gen.spec, het,iter) %>% 
  summarize(entropy = sum(ifelse(value == 0, 0, (value)*log(1/(value))))) %>% 
  group_by(name,gen.spec, het) %>% 
  summarize(mean = mean(entropy),
            min = min(entropy),
            max = max(entropy)) %>% 
  mutate(name = factor(name, 
                       levels = c("prop.obs", "value.true.kernel", "ssd","ssf","rsf"),
                       labels = c("Simulated Use", "True Selection", "SSD","SSF","RSF"))) %>% 
  ggplot(aes(x = factor(het), y = mean, ymin = min, ymax = max, color = name, group = paste(name))) +
  geom_path(position = position_dodge(0.2)) +
  geom_pointrange(position = position_dodge(0.2)) +
  facet_wrap(~gen.spec, nrow = 1) +
  labs(x = "Gaussian Process Scale (Heterogeneity)",
       y = "Shannon Entropy\n(Mean and Range in 5 Replicates)",
       color = "Probability Kernel") +
  scale_color_scico_d(guide = "none") +
  theme_classic() +
  theme(axis.text.x = element_text(angle = 45, vjust = 1, hjust = 1))
```

```
## `summarise()` has grouped output by 'name', 'gen.spec', 'het'. You can override
## using the `.groups` argument.
## `summarise()` has grouped output by 'name', 'gen.spec'. You can override using
## the `.groups` argument.
```

```
shannon.reduced
```

```
# ggsave("Plots/shannonentropyreduced.svg", shannon.reduced, dpi = 600, width = 10, height = 3)
```

###### 2.2.11.7 Kernel prediction relative entropy (Kullback–Leibler Divergence)

```
relative.entropy <- g %>% 
  mutate(animal = factor(animal, levels = paste0("ID",1:10), ordered = T)) %>%
  mutate(specialism = factor(animal, levels = paste0("ID",1:10), labels = 1:10, ordered = T)) %>%
  pivot_longer(c(ssd, value.true.kernel, ssf, rsf)) %>% 
  group_by(name,specialism, het,iter) %>% 
  summarize(entropy = sum(value*log((value + 1E-6)/(prop.obs + 1E-6)), na.rm = T)) %>% 
  group_by(name,specialism, het) %>% 
  summarize(entropy_median = mean(entropy),
            entropy_min = min(entropy),
            entropy_max = max(entropy)) %>% 
  mutate(name = factor(name, 
                       levels = c("prop.obs", "value.true.kernel", "ssd","ssf","rsf"),
                       labels = c("Simulated Use", "True Selection", "SSD","SSF","RSF"))) %>% 
  ggplot(aes(x = factor(het), y = entropy_median, ymin = entropy_min, ymax = entropy_max, color = name, group = paste(name))) +
  geom_path(position = position_dodge(0.2)) +
  geom_pointrange(position = position_dodge(0.2)) +
  facet_wrap(~specialism, ncol = 5, scales = "free_y") +
  labs(x = "Gaussian Process Scale (Heterogeneity)",
       y = "Relative Entropy Compared to Observed Use\n(Mean and Range in 5 Replicates)",
       color = "Probability Kernel") +
  scale_color_scico_d(begin = 0.25) +
  theme_classic() +
  theme(axis.text.x = element_text(angle = 45, vjust = 1, hjust = 1))
```

```
## `summarise()` has grouped output by 'name', 'specialism', 'het'. You can
## override using the `.groups` argument.
## `summarise()` has grouped output by 'name', 'specialism'. You can override
## using the `.groups` argument.
```

```
relative.entropy
```

```
# ggsave("Plots/relativeentropyall.png", relative.entropy, dpi = 600, width = 12, height = 6)


relative.entropy.reduced <- g %>% 
  mutate(gen.spec = case_when(
    animal == "ID2" ~ "Strong Generalist",
    animal == "ID4" ~ "Slight Generalist",
    animal == "ID6" ~ "Slight Specialist",
    animal == "ID8" ~ "Strong Specialist"
  )) %>%  
  filter(!is.na(gen.spec)) %>% 
  mutate(gen.spec = factor(gen.spec, levels = c("Strong Generalist","Slight Generalist","Slight Specialist","Strong Specialist"), labels = c("Strong Generalist (2)","Slight Generalist (4)","Slight Specialist (6)","Strong Specialist (8)"), ordered = T)) %>% 
  pivot_longer(c(ssd, value.true.kernel, ssf, rsf)) %>% 
  group_by(name,gen.spec, het,iter) %>% 
  summarize(entropy = sum(prop.obs*log((prop.obs + 1E-10)/(value + 1E-10)), na.rm = T)) %>% 
  group_by(name,gen.spec, het) %>% 
  summarize(entropy_median = mean(entropy),
            entropy_min = min(entropy),
            entropy_max = max(entropy)) %>% 
  mutate(name = factor(name, 
                       levels = c("prop.obs", "value.true.kernel", "ssd","ssf","rsf"),
                       labels = c("Simulated Use", "True Selection", "SSD","SSF","RSF"))) %>% 
  ggplot(aes(x = factor(het), y = entropy_median, ymin = entropy_min, ymax = entropy_max, color = name, group = paste(name))) +
  geom_path(position = position_dodge(0.2)) +
  geom_pointrange(position = position_dodge(0.2)) +
  facet_wrap(~gen.spec, nrow = 1) +
  labs(x = "Gaussian Process Scale (Heterogeneity)",
       y = "Relative Entropy (vs. Observed Use)\n(Mean and Range in 5 Replicates)",
       color = "Probability Kernel") +
  scale_color_scico_d(begin = 0.25) +
  theme_classic() + 
  theme(axis.text.x = element_text(angle = 45, vjust = 1, hjust = 1))
```

```
## `summarise()` has grouped output by 'name', 'gen.spec', 'het'. You can override
## using the `.groups` argument.
## `summarise()` has grouped output by 'name', 'gen.spec'. You can override using
## the `.groups` argument.
```

```
relative.entropy.reduced
```

```
# ggsave("Plots/relativeentropy.svg", relative.entropy.reduced, dpi = 600, width = 10, height = 4)
```

###### 2.2.11.8 Kernel prediction rootmean square error

```
rmse <- g %>% 
  mutate(animal = factor(animal, levels = paste0("ID",1:10), ordered = T)) %>%
  mutate(specialism = factor(animal, levels = paste0("ID",1:10), labels = 1:10, ordered = T)) %>%
  pivot_longer(c(ssd, value.true.kernel, ssf, rsf)) %>% 
  group_by(name,specialism, het,iter) %>% 
  summarize(RMSE = sqrt(mean((value-prop.obs)^2))) %>% 
  group_by(name,specialism, het) %>% 
  summarize(RMSE_mean = mean(RMSE),
            RMSE_min = min(RMSE),
            RMSE_max = max(RMSE)) %>% 
  mutate(name = factor(name, 
                       levels = c("prop.obs", "value.true.kernel", "ssd","ssf","rsf"),
                       labels = c("Simulated Use", "True Selection", "SSD","SSF","RSF"))) %>% 
  ggplot(aes(x = factor(het), y = RMSE_mean, ymin = RMSE_min, ymax = RMSE_max, color = name, group = paste(name))) +
  geom_path(position = position_dodge(0.2)) +
  geom_pointrange(position = position_dodge(0.2)) +
  facet_wrap(~specialism, scales = "free_y", ncol = 5) +
  labs(x = "Gaussian Process Scale (Heterogeneity)",
       y = "Root Mean Squared Error to Observed Use\n(Mean and Range in 5 Replicates)",
       color = "Probability Kernel") +
  scale_color_scico_d(begin = 0.25) +
  theme_classic() +
  theme(axis.text.x = element_text(angle = 45, vjust = 1, hjust = 1))
```

```
## `summarise()` has grouped output by 'name', 'specialism', 'het'. You can
## override using the `.groups` argument.
## `summarise()` has grouped output by 'name', 'specialism'. You can override
## using the `.groups` argument.
```

```
rmse
```

```
# ggsave("Plots/rmseall.png", rmse, dpi = 600, width = 12, height = 6)

rmse.reduced <- g %>% 
  mutate(gen.spec = case_when(
    animal == "ID2" ~ "Strong Generalist",
    animal == "ID4" ~ "Slight Generalist",
    animal == "ID6" ~ "Slight Specialist",
    animal == "ID8" ~ "Strong Specialist"
  )) %>%  
  filter(!is.na(gen.spec)) %>% 
  mutate(gen.spec = factor(gen.spec, levels = c("Strong Generalist","Slight Generalist","Slight Specialist","Strong Specialist"), labels = c("Strong Generalist (2)","Slight Generalist (4)","Slight Specialist (6)","Strong Specialist (8)"), ordered = T)) %>% 
  pivot_longer(c(ssd, value.true.kernel, ssf, rsf)) %>% 
  group_by(name,gen.spec, het,iter) %>% 
  summarize(RMSE = sqrt(mean((value-prop.obs)^2))) %>% 
  group_by(name,gen.spec, het) %>% 
  summarize(RMSE_mean = mean(RMSE),
            RMSE_min = min(RMSE),
            RMSE_max = max(RMSE)) %>% 
  mutate(name = factor(name, 
                       levels = c("prop.obs", "value.true.kernel", "ssd","ssf","rsf"),
                       labels = c("Simulated Use", "True Selection", "SSD","SSF","RSF"))) %>% 
  ggplot(aes(x = factor(het), y = RMSE_mean, ymin = RMSE_min, ymax = RMSE_max, color = name, group = paste(name))) +
  geom_path(position = position_dodge(0.2)) +
  geom_pointrange(position = position_dodge(0.2)) +
  facet_grid(~gen.spec) +
  labs(x = "Gaussian Process Scale (Heterogeneity)",
       y = "Root Mean Squared Error to Observed Use\n(Mean and Range in 5 Replicates)",
       color = "Probability Kernel") +
  scale_color_scico_d(begin = 0.25) +
  theme_classic() +
  theme(axis.text.x = element_text(angle = 45, vjust = 1, hjust = 1),
        legend.position = "inside",
        legend.position.inside = c(0.2, 0.8))
```

```
## `summarise()` has grouped output by 'name', 'gen.spec', 'het'. You can override
## using the `.groups` argument.
## `summarise()` has grouped output by 'name', 'gen.spec'. You can override using
## the `.groups` argument.
```

```
rmse.reduced
```

```
# ggsave("Plots/rmse.reduced.png", rmse.reduced, dpi = 600, width = 8, height = 5)
```

##### 2.2.12 Validating comparison tools

###### 2.2.12.1 Spearman rho comparisons are not effective at discriminating best models, but Pearson is

```
g.cor <- g %>%
  pivot_longer(c(ssd, ssf, rsf)) %>%
  group_by(name, het, animal, iter) %>%
  mutate(quantile = as.numeric(cut(value, c(-Inf,quantile(value, seq(0.01,0.99,by= 0.01)),Inf)))) %>%
  mutate(bin = as.numeric(quantile)) %>%
  group_by(name, het, animal, iter, bin) %>%
  summarize(sum = sum(prop.obs)) %>%
  group_by(name, het, animal, iter) %>%
  summarize(error.obs = cor(bin, sum, method = "spearman"))
```

```
## `summarise()` has grouped output by 'name', 'het', 'animal', 'iter'. You can
## override using the `.groups` argument.
## `summarise()` has grouped output by 'name', 'het', 'animal'. You can override
## using the `.groups` argument.
```

```
g.cor.s <- g %>% 
  pivot_longer(c(ssd, ssf, rsf)) %>% 
  group_by(name, het, animal, iter) %>% 
  summarize(error.obs = cor(prop.obs, value, method = "spearman"),
            error.exp = cor(value.true.kernel, value, method = "spearman")) %>% 
  pivot_longer(c("error.obs", "error.exp"), names_to = "truth")
```

```
## `summarise()` has grouped output by 'name', 'het', 'animal'. You can override
## using the `.groups` argument.
```

```
g.cor.s %>% 
  filter(truth == "error.obs") %>% 
  mutate(het = factor(het)) %>% 
  mutate(truth = factor(truth, levels = c("error.obs", "error.exp"), labels = c("Observed Use", "Theoretical Selection"), ordered = T)) %>%
  mutate(name = factor(name, levels = c("ssd", "ssf", "rsf"), labels = c("SSD","SSF","RSF"), ordered = T)) %>%
  mutate(animal = factor(animal, levels = paste0("ID",1:10), ordered = T)) %>% 
  group_by(name, het, animal, truth) %>% 
  summarize(mean = mean(value),
            min = min(value),
            max = max(value)) %>% 
  ggplot(aes(x = factor(het), color = name, y = mean, ymin = min, ymax = max)) +
  geom_pointrange(position = position_dodge(0.5)) +
  facet_wrap(.~animal, scales = "free_y", ncol = 5) +
  scale_color_scico_d(begin = 0.5) +
  theme_classic() +
  labs(y = "Spearman Correlation between Prediction and\nSimulated Use (Mean and Range in 5 Replicates)",
       color = "Probability Kernel",
       x = "Gaussian Process Scale (Heterogeneity)")  +
  theme(axis.text.x = element_text(angle = 45, vjust = 1, hjust = 1))
```

```
## `summarise()` has grouped output by 'name', 'het', 'animal'. You can override
## using the `.groups` argument.
```

```
# ggsave("Plots/spearman.simulation.png", dpi = 600, width = 9, height = 5)

g.cor.p <- g %>% 
  pivot_longer(c(ssd, ssf, rsf)) %>% 
  group_by(name, het, animal, iter) %>% 
  summarize(error.obs = cor(prop.obs, value, method = "pearson"),
            error.exp = cor(value.true.kernel, value, method = "pearson")) %>% 
  pivot_longer(c("error.obs", "error.exp"), names_to = "truth")
```

```
## `summarise()` has grouped output by 'name', 'het', 'animal'. You can override
## using the `.groups` argument.
```

```
g.cor.p %>% 
  filter(truth == "error.obs") %>% 
  mutate(het = factor(het)) %>% 
  mutate(truth = factor(truth, levels = c("error.obs", "error.exp"), labels = c("Observed Use", "Theoretical Selection"), ordered = T)) %>%
  mutate(name = factor(name, levels = c("ssd", "ssf", "rsf"), labels = c("SSD","SSF","RSF"), ordered = T)) %>%
  mutate(animal = factor(animal, levels = paste0("ID",1:10), ordered = T)) %>% 
  group_by(name, het, animal, truth) %>% 
  summarize(mean = mean(value),
            min = min(value),
            max = max(value)) %>% 
  ggplot(aes(x = factor(het), color = name, y = mean, ymin = min, ymax = max)) +
  geom_pointrange(position = position_dodge(0.5)) +
  facet_wrap(~animal, scales = "free_y", ncol = 5) +
  scale_color_scico_d(begin = 0.5) +
  theme_classic() +
  labs(y = "Pearson Correlation between Prediction and\nSimulated Use (Mean and Range in 5 Replicates)",
       color = "Probability Kernel",
       x = "Gaussian Process Scale (Heterogeneity)") +
  theme(axis.text.x = element_text(angle = 45, vjust = 1, hjust = 1))
```

```
## `summarise()` has grouped output by 'name', 'het', 'animal'. You can override
## using the `.groups` argument.
```

```
# ggsave("Plots/pearson.simulation.png", dpi = 600, width = 9, height = 5)
```

###### 2.2.12.2 Geometric mean predicted use of out of sample points may provide a more accurate representation, but is influenced by heterogeneity in SSD - this is similar in some ways to shannon entropy - the log just dampens the bias in taking means of skewed distributions

```
d <- g %>% 
  filter(iter == 1, het == 0.6, animal == "ID8")
mean(d$prop.obs*d$ssd)
```

```
## [1] 5.321433e-08
```

```
g.mu <- g %>% 
  pivot_longer(c(ssd, ssf, rsf)) %>% 
  group_by(name, het, animal, iter) %>% 
  summarize(obs.w.mean = exp(weighted.mean(log(value), prop.obs)),
            exp.w.mean = exp(weighted.mean(log(value), value.true.kernel))) %>% 
  pivot_longer(c("obs.w.mean", "exp.w.mean"), names_to = "truth")
```

```
## `summarise()` has grouped output by 'name', 'het', 'animal'. You can override
## using the `.groups` argument.
```

```
g.mu %>% 
  filter(truth == "obs.w.mean") %>% 
  mutate(truth = factor(truth, levels = c("obs.w.mean", "exp.w.mean"), labels = c("Observed Use", "Theoretical Selection"), ordered = T)) %>%
  mutate(het = factor(het)) %>% 
  mutate(name = factor(name, levels = c("ssd", "ssf", "rsf"), labels = c("SSD","SSF","RSF"), ordered = T)) %>%
  mutate(animal = factor(animal, levels = paste0("ID",1:10), ordered = T)) %>% 
  group_by(truth, name, het, animal) %>% 
  summarize(mean = mean(value),
            min = min(value),
            max = max(value)) %>% 
  ggplot(aes(x = factor(het), color = name, y = mean, ymin = min, ymax = max)) +
  geom_pointrange(position = position_dodge(0.5)) +
  facet_wrap(~animal, scales = "free_y", ncol = 5) +
  scale_color_scico_d(begin = 0.5) +
  theme_classic() +
  labs(y = "Geometric Mean Predicted Use of Out-of-Sample\nPoints (Mean and Range in 5 Replicates)",
       color = "Probability Kernel",
       x = "Gaussian Process Scale (Heterogeneity)") +
  theme(axis.text.x = element_text(angle = 45, vjust = 1, hjust = 1))
```

```
## `summarise()` has grouped output by 'truth', 'name', 'het'. You can override
## using the `.groups` argument.
```

```
# ggsave("Plots/geomean.simulation.png", dpi = 600, width = 12, height = 5)
```

##### 2.2.13 Total Linear Comparisons

```
linear.comparisons.data <-  g %>% 
  # filter(animal %in%  paste0("ID",c(2,4,6,8))) %>% 
  # sample_frac(0.01) %>% 
  # filter(het %in% c(0.001, 0.1, 0.6)) %>% 
  pivot_longer(c(ssd, ssf, rsf)) %>%
  # mutate(difference = prop.obs - value.true.kernel) %>% 
  mutate(het = factor(het)) %>% 
  mutate(name = factor(name, levels = c("ssd", "ssf", "rsf"), labels = c("SSD","SSF","RSF"), ordered = T)) %>%
  mutate(animal = factor(animal, levels = paste0("ID",1:10), ordered = T)) 

linear.comparisons <- linear.comparisons.data %>% 
  filter(het %in% c(0.05, 0.2, 0.6) &
           animal %in% paste0("ID",c(2,4,6,8))) %>% 
  mutate(gen.spec = case_when(
    animal == "ID2" ~ "Strong Generalist",
    animal == "ID4" ~ "Slight Generalist",
    animal == "ID6" ~ "Slight Specialist",
    animal == "ID8" ~ "Strong Specialist"
  )) %>% 
  mutate(gen.spec = factor(gen.spec, levels = c("Strong Generalist","Slight Generalist","Slight Specialist","Strong Specialist"), labels = c("Strong Generalist (2)","Slight Generalist (4)","Slight Specialist (6)","Strong Specialist (8)"), ordered = T)) %>% 
  mutate(scale = paste("Scale = ", het)) %>%
  ggplot(aes(x = prop.obs, y = value, color = name, group = paste(name,iter))) +
  geom_abline(intercept = 0, slope = 1, size = 0.1) +
  geom_point(alpha = 0.1) +
  stat_smooth(geom="line", method ="lm", alpha = 0.75) +  
  scale_color_scico_d(begin = 0.5) +
  theme_classic()+
  facet_grid(scale~gen.spec) +
  scale_y_log10() +
  scale_x_log10() +
  labs(x = "Observed Probability of Use",
       y = "Predicted Probability of Use", 
       color = "Probability Kernel")

linear.comparisons
```

```
## Warning in scale_x_log10(): log-10 transformation introduced infinite values.
## log-10 transformation introduced infinite values.
```

```
## `geom_smooth()` using formula = 'y ~ x'
```

```
## Warning: Removed 3696 rows containing non-finite outside the scale range
## (`stat_smooth()`).
```

```
# ggsave("Plots/linear.comparison.png", linear.comparisons, dpi = 300, width = 10, height = 8)
```

### 3 Application

#### 3.1 Deer case study

We can scale this idea to predict *future movements* of an
individual

```
set.seed(100)
data("deer") # read in the deer data included as part of the amt package
# data("sh_forest") # read in the spatial included as part of the amt package
# sh_forest <- raster(terra::unwrap(sh_forest))
# polygon.forest <- rasterToPolygons(sh_forest, fun = function(x){x == 1}, n = 16, dissolve = T) # convert the forest layer to dissolved polygons
# polygon..not.forest <- rasterToPolygons(sh_forest, fun = function(x){x == 0}, n = 16, dissolve = T) # convert the forest layer to dissolved polygons
# spts <- rasterToPoints(sh_forest, spatial = TRUE) # draw the point cloud of raster
# 
# require(sf)
# 
# dd <- sf::st_distance(st_as_sf(as(sh_forest,"SpatialPoints")), st_as_sf(polygon.forest))
# dd1 <- sf::st_distance(st_as_sf(as(sh_forest,"SpatialPoints")), st_as_sf(polygon..not.forest))
# 
# dd <- units::drop_units(dd[,1])
# dd1 <- units::drop_units(dd1[,1])
# 
# forest_distance <- setValues(sh_forest, ifelse(dd == 0, dd1, dd)) # force shortest distance to a new matrix
# names(forest_distance) <- "dist.forest" # name it
# rstack <- stack(sh_forest, forest_distance) # stack the data
# saveRDS(rstack, "deerrasters.RDS") # save the data to save time for compilation
rstack <- readRDS("Data and Intermediate RDS files/deerrasters.RDS") # call the data
```

Replicate preprocessing as above

```
sh_forest <- aggregate(rstack, 5, fun=mean) # aggregate the data for the sake of demonstration

ssf1 <- deer %>% arrange(t_) %>% mutate(row = 1:n())
ssf1.train <- ssf1 %>% filter(row %in% 1:round(n()*0.75))
ssf1.test <- ssf1 %>% filter(!row %in% ssf1.train$row)

ssf1 <- ssf1.train %>% 
  steps_by_burst() # the data are already cleaned to 6hr steps, make the step-by-burst file

fatter.unif <- original.unif <- fit_distr(ssf1$sl_, "unif") # get the original unif distribution
fatter.unif$params$min <- 0 # take the min to 0 
fatter.unif$params$max <- fatter.unif$params$max * 1.25 # take the max up 

ssf1.deer <- ssf1 %>% 
  random_steps(n_control = 100, sl_distr = fatter.unif) %>% # as above, 40 random steps
  extract_covariates(sh_forest) # extract covariates
```

```
## Warning in random_steps.bursted_steps_xyt(., n_control = 100, sl_distr =
## fatter.unif): Some bursts contain < 3 steps and will be removed
```

```
plot(sh_forest[[2]]) # distance to forest
lines(ssf1.train$x_, ssf1.train$y_, col = "black") # the train data
lines(ssf1.test$x_, ssf1.test$y_, col = "red") # the test data
```

```
plot(sh_forest[[1]])
```

We can fit a basic movement model just using a polynomial of step
distance and the distance to forest. This deer enjoys the forest and
tends to move a bit every six hours.

```
model.deer <- ssf1.deer %>%
  fit_issf(case_ ~ dist.forest * forest + sl_ + log(sl_) + strata(step_id_), model = T)
summary(model.deer)
```

```
## Call:
## coxph(formula = Surv(rep(1, 57368L), case_) ~ dist.forest * forest + 
##     sl_ + log(sl_) + strata(step_id_), data = data, model = ..1, 
##     method = "exact")
## 
##   n= 57245, number of events= 568 
##    (123 observations deleted due to missingness)
## 
##                          coef  exp(coef)   se(coef)       z Pr(>|z|)    
## dist.forest        -0.0065280  0.9934932  0.0006087 -10.725  < 2e-16 ***
## forest             -0.4945428  0.6098496  0.2604859  -1.899 0.057625 .  
## sl_                -0.0020736  0.9979286  0.0001737 -11.941  < 2e-16 ***
## log(sl_)           -0.1842541  0.8317245  0.0485654  -3.794 0.000148 ***
## dist.forest:forest  0.0063524  1.0063726  0.0018477   3.438 0.000586 ***
## ---
## Signif. codes:  0 '***' 0.001 '**' 0.01 '*' 0.05 '.' 0.1 ' ' 1
## 
##                    exp(coef) exp(-coef) lower .95 upper .95
## dist.forest           0.9935     1.0065    0.9923    0.9947
## forest                0.6098     1.6397    0.3660    1.0161
## sl_                   0.9979     1.0021    0.9976    0.9983
## log(sl_)              0.8317     1.2023    0.7562    0.9148
## dist.forest:forest    1.0064     0.9937    1.0027    1.0100
## 
## Concordance= 0.952  (se = 0.003 )
## Likelihood ratio test= 2327  on 5 df,   p=<2e-16
## Wald test            = 701.5  on 5 df,   p=<2e-16
## Score (logrank) test = 4820  on 5 df,   p=<2e-16
```

We can make our surface for predictions. The units of each cell - the
resolution - is 125m x 125m. Other than that, the pipeline as above is
repeated here.

```
mock.surface <- create_mock_surface(sh_forest, F, list(x = xres(sh_forest), y = yres(sh_forest))) # generates prediction surface
pred.data <- get_cells(model.deer, 
                       mock.surface,
                       sh_forest) # gets the contexts of each cell
cell.data <- get_cell_data(model.deer, pred.data) # checks that the ssf model works for cell contexts
neighbors.found <- neighbor_lookup(mock.surface, cell.data) # finds the neighbor call-up matrix
sparse.neighbors <- neighbor_finder(model.deer, cell.data, neighbors.found, quantile = 0.95, distance.override = quantile(ssf1$sl_, 0.95)) # finds neighbors within the 90th percentils
```

```
## [1] "Creating neighbor comparisons"
## [1] "Finding valid comparisons"
## [1] "Splitting cell.data into list"
## [1] "Running comparisons"
```

```
ssf.comparisons <- compile_ssf_comparisons(sparse.neighbors, cell.data) # grabs predictions datasets
ssf.comparisons <- lapply(ssf.comparisons, function(x) {
  list(.for = x$.for, .given = x$.given) # generates for and given comparisons per cell
})
surface <- predict_ssf_comparisons(model.deer, ssf.comparisons) # makes predictions to generate the A matrix
```

```
## [1] "Estimating probability surface"
## [1] "Compiling probability surface"
## [1] "Making sparse matrix for transitions"
```

We generate the stable state distribution as above.

```
A <- surface$prob.matrix # matrix A
d <- eigs(t(A), 1) # eigen decomp
d.1 <- Re(d$vectors[,1]) # first vector
prob.d.deer <- d.1/sum(d.1) # make sure it sums to 1
ssd.raster.deer <- setValues(mock.surface, prob.d.deer) # make raster
par(oma = c(1,1,1,1))
plot((ssd.raster.deer))
```

```
# points(deer, pch = ".", col = alpha("black", 0.25))


raster.df.deer.ssd <- as.data.frame(rasterToPoints(ssd.raster.deer)) # save raster as a df for plotting later
```

Application of an SSF to an entired surface (not respecting
mechanisms, just selection coefficients).

```
a.data <- pred.data # grab prior matrix data 
a.data$sl_ <- mean(ssf1.deer$sl_) # use mean step length
a.data$x2_ <- a.data$x
a.data$y2_ <- a.data$y
b.data <- a.data[1,] # set the first cell as the baseline

log.rss <- amt::log_rss(model.deer, # the model
                          a.data, # the raster data (including missing values)
                          b.data,  # a row of the raster data (excluding missing values)
                          ci = NA)
ssf <-  exp(log.rss$df$log_rss)/sum(exp(log.rss$df$log_rss)) # turn into kernel after making values positive
ssf.prob.raster.deer <- setValues(mock.surface, ssf) # make raster

par(oma = c(1,1,1,1))
plot(ssf.prob.raster.deer)
```

```
raster.df.deer.ssf <- as.data.frame(rasterToPoints(ssf.prob.raster.deer)) # save raster as a df for plotting later
# points(deer, pch = ".", col = alpha("black", 0.25))
```

We replicate an rsf with same underlying model.

```
rsf <- ssf1.train %>% 
  random_points() %>% # defaults
  extract_covariates(sh_forest) # grab distance

rsf <- rsf %>% 
  mutate(x2_ = x_, y2_ = y_) %>% 
  fit_rsf(case_ ~ dist.forest * forest , model = T) # fit model

pred.data$x2_ <- pred.data$x
pred.data$y2_ <- pred.data$y

probabilities <- exp(predict(rsf$model, newdata = pred.data))/(1+exp(predict(rsf$model, newdata = pred.data))) # get to probabilities
probabilities.kern <- probabilities/sum(probabilities)
rsf.prob.raster.deer <- setValues(mock.surface, probabilities.kern)
par(oma = c(1,1,1,1))
plot(rsf.prob.raster.deer)
```

```
raster.df.deer.rsf <- as.data.frame(rasterToPoints(rsf.prob.raster.deer)) # save raster as a df for plotting later
# points(deer, pch = ".", col = alpha("black", 0.25))
```

##### 3.1.1 Comparisons

Merging the predicted surfaces and generating outputs.

```
deer1.rstack <- stack(ssd.raster.deer, ssf.prob.raster.deer, rsf.prob.raster.deer)

require(rasterVis)
levelplot(deer1.rstack)
```

```
pred.values <- as.data.frame(values(deer1.rstack))
colnames(pred.values) <- c("SSD","SSF","RSF")

shannon.deer1 <- pred.values %>% 
  pivot_longer(all_of(colnames(pred.values))) %>% 
  group_by(name) %>% 
  mutate(name = factor(name, c("SSD","SSF","RSF"))) %>% 
  summarize(shannon = -sum(value*log(value))) 

shannon.deer1 %>% 
  ggplot(aes(x = name, y = shannon, group = "A")) +
  geom_point() +
  geom_path() +
  scale_color_scico_d(begin = 0.5)+
  theme_classic()
```

```
pred.values %>% 
  pivot_longer(all_of(colnames(pred.values))) %>% 
  group_by(name) %>% 
  arrange(-value) %>% 
  mutate(name = factor(name, c("SSD","SSF","RSF"))) %>% 
  mutate(cdf = cumsum(value)) %>% 
  summarize(total.50 = sum(cdf <= 0.5))
```

```
## # A tibble: 3 × 2
##   name  total.50
##   <fct>    <int>
## 1 SSD       3329
## 2 SSF       4408
## 3 RSF       4385
```

```
use.area.dee1 <- pred.values %>% 
  pivot_longer(all_of(colnames(pred.values))) %>% 
  group_by(name) %>% 
  mutate(name = factor(name, c("SSD","SSF","RSF"))) %>% 
  arrange(-value) %>% 
  mutate(cdf = cumsum(value)) %>% 
  summarize(sum = sum(cdf <= 0.5))
```

```
# spearman correlation
comparison.in.spearman <- compare_spearman(ssf1.train, deer1.rstack, bootstrap = 100, bins = 10)
```

```
## `summarise()` has grouped output by 'name'. You can override using the
## `.groups` argument.
```

```
comparison.id.spearman <- compare_spearman(ssf1.test, deer1.rstack, bootstrap = 100, bins = 10)
```

```
## `summarise()` has grouped output by 'name'. You can override using the
## `.groups` argument.
```

```
# mean log predicted use
comparison.in.mean <- compare_mean(ssf1.train, deer1.rstack, bootstrap = 100)
comparison.id.mean <- compare_mean(ssf1.test, deer1.rstack, bootstrap = 100)


comparison.in.spearman$in.out <- "In-Sample"
comparison.id.spearman$in.out <- "Within Individual"
comparison.in.mean$in.out <- "In-Sample"
comparison.id.mean$in.out <- "Within Individual"

comparison.in.spearman$type <- "Spearman"
comparison.id.spearman$type <- "Spearman"
comparison.in.mean$type <- "Geometric Mean"
comparison.id.mean$type <- "Geometric Mean"

bound.comparisons.deer <- rbind(comparison.in.spearman, comparison.id.spearman, comparison.in.mean, comparison.id.mean)


deer.data.1 <- bound.comparisons.deer %>% 
  mutate(bootstrap = factor(bootstrap),
         model = case_when(
           name %in% factor(1:3) ~ name,
           name == "layer.1" ~ factor(1),
           name == "layer.2" ~ factor(2),
           name == "layer.3" ~ factor(3)
         )) %>% 
  mutate(model = factor(model, levels = 1:3, labels = c("SSD", "SSF", "RSF"), ordered = T)) %>% 
  mutate(in.out = factor(in.out, levels = c("In-Sample", "Within Individual", "Between Individuals", "Between Contexts"), ordered = T)) %>% 
  ungroup() %>% 
  dplyr::select(-name) %>% 
  group_by(in.out, model, type) %>%
  pivot_wider(names_from = c(model), values_from = measure) %>% 
  pivot_longer(c(RSF, SSF)) %>% 
  group_by(in.out, name, type) %>%
  mutate(delta = SSD-value) %>%
  group_by(in.out, name, type) %>%
  summarize(median = median(delta),
            lower = quantile(delta, 0.025),
            upper = quantile(delta, 0.975)) %>%
  # pivot_wider(names_from = type, values_from = c(median, lower, upper))
  mutate(model = factor(name, levels = c("SSF","RSF"), labels = c("SSD-SSF","SSD-RSF"), ordered = T))
```

```
## `summarise()` has grouped output by 'in.out', 'name'. You can override using
## the `.groups` argument.
```

```
deer.data.1 %>% 
  arrange(model) %>% 
  ggplot(aes(x = model, y = median, ymax = upper, ymin = lower, shape = type, group = type)) +
  geom_hline(yintercept = 0) +
  geom_path(position = position_dodge(0.2)) +
  geom_pointrange(position = position_dodge(0.2)) +
  facet_grid(type~in.out, scales = "free_y") +
  labs(x = "Prediction Comparison",
       y = "Difference from SSD Prediction\n(Median and 95% Quantile)",
       shape = "Comparison Heuristic") + 
  theme_classic() +
  scale_alpha_discrete(guide = F, range = c(0.5, 1))+
  scale_y_continuous(trans = "both.sqrt.trans_trans")
```

```
## Warning: Using alpha for a discrete variable is not advised.
```

```
## Warning: The `guide` argument in `scale_*()` cannot be `FALSE`. This was deprecated in
## ggplot2 3.3.4.
## ℹ Please use "none" instead.
## This warning is displayed once every 8 hours.
## Call `lifecycle::last_lifecycle_warnings()` to see where this warning was
## generated.
```

```
# simulated.test <- conover.test(simple.deer.comparison$rho, simple.deer.comparison$model, method="bonferroni")
# simulated.test$P.adjusted
```

##### 3.1.2 Storing some output and plots for later

```
ssd.raster.deer.df <- as.data.frame(rasterToPoints(ssd.raster.deer))
ssf.prob.raster.deer.df <- as.data.frame(rasterToPoints(ssf.prob.raster.deer))
rsf.prob.raster.deer.df <- as.data.frame(rasterToPoints(rsf.prob.raster.deer))

min.max.deer <- rbind(ssd.raster.deer.df, ssf.prob.raster.deer.df, rsf.prob.raster.deer.df)
min.deer <- min(min.max.deer$layer)
max.deer <- max(min.max.deer$layer)

theme.current <- theme(
        panel.background = element_blank(),
        panel.grid = element_blank(),
        legend.position = "bottom",
        plot.title = element_text(hjust = 0.5))

ssd.deer.plot <- ssd.raster.deer.df %>% 
  ggplot() +
  geom_raster(aes(x = (x-min(x))/1000, y = (y-min(y))/1000, fill = layer), color = NA) +
  coord_equal() +
  scale_fill_viridis_c(option = "H", name = "Kernel Probability",
                       limits = c(min.deer,max.deer)) +
  # geom_point(data = deer, mapping = aes(x = x_, y = y_), color = "green", size = size.point, alpha = alpha.point) +
  # geom_path(data = ssf1, mapping = aes(x = x1_, y = y1_), color = "white", size = 0.1, alpha = 0.1) +
  ggtitle("SSD") +
  theme.current +
  labs(x = "Easting (km)", y = "Northing (km)")
```

```
## Warning in geom_raster(aes(x = (x - min(x))/1000, y = (y - min(y))/1000, :
## Ignoring unknown parameters: `colour`
```

```
ssf.deer.plot <- ssf.prob.raster.deer.df %>% 
  ggplot() +
  geom_raster(aes(x = (x-min(x))/1000, y = (y-min(y))/1000, fill = layer), color = NA) +
  coord_equal() +
  scale_fill_viridis_c(option = "H", name = "Kernel Probability",
                       limits = c(min.deer,max.deer)) +
  # geom_point(data = deer, mapping = aes(x = x_, y = y_), color = "green", size = size.point, alpha = alpha.point) +
  # geom_path(data = ssf1, mapping = aes(x = x1_, y = y1_), color = "white", size = 0.1, alpha = 0.1) +
  ggtitle("SSF") +
  theme.current+
  labs(x = "Easting (km)", y = "Northing (km)")
```

```
## Warning in geom_raster(aes(x = (x - min(x))/1000, y = (y - min(y))/1000, :
## Ignoring unknown parameters: `colour`
```

```
rsf.deer.plot <- rsf.prob.raster.deer.df %>% 
  ggplot() +
  geom_raster(aes(x = (x-min(x))/1000, y = (y-min(y))/1000, fill = layer), color = NA) +
  coord_equal() +
  scale_fill_viridis_c(option = "H", name = "Kernel Probability",
                       limits = c(min.deer,max.deer)) +
  # geom_point(data = deer, mapping = aes(x = x_, y = y_), color = "green", size = size.point, alpha = alpha.point) +
  # geom_path(data = ssf1, mapping = aes(x = x1_, y = y1_), color = "white", size = 0.1, alpha = 0.1) +
  ggtitle("RSF") +
  theme.current+
  labs(x = "Easting (km)", y = "Northing (km)")
```

```
## Warning in geom_raster(aes(x = (x - min(x))/1000, y = (y - min(y))/1000, :
## Ignoring unknown parameters: `colour`
```

```
simple.deer <- ggarrange(rsf.deer.plot, ssf.deer.plot, ssd.deer.plot, common.legend = T, nrow = 1, legend = "left")
simple.deer
```

#### 3.2 Italy Deer Project

We can scale up our projections to think about *other*
individuals.

```
# Deer <- read_csv("EuroDeer_ Roe deer in Italy 2005-2008.csv")
# 
# Deer$t_ <- as.POSIXct(Deer$timestamp)
# Deer$id <- Deer$`individual-local-identifier`
# Deer$x_ <- Deer$`utm-easting`
# Deer$y_ <- Deer$`utm-northing`
# Deer$day <- yday(Deer$t_)
# Deer$season <- ifelse(Deer$day %in% c(1:120, 290:366), "Winter", NA)
# Deer$season <- ifelse(Deer$day %in% c(120:290), "Summer", Deer$season)
# 
# Deer <- Deer %>% 
#   dplyr::select(c("id", "x_", "y_", "t_", "season", "day"))  %>% 
#   filter(season == "Summer")
# 
# saveRDS(Deer, "italy.deer.RDS")
Deer <- readRDS("Data and Intermediate RDS files/italy.deer.RDS")

Deer %>% 
  group_by(id) %>% 
  summarize(fix = median(diff(t_)/3600,))
```

```
## # A tibble: 5 × 2
##   id               fix          
##   <chr>            <drtn>       
## 1 Agostino (M03)   3.994722 secs
## 2 Alessandra (F10) 3.999722 secs
## 3 Daniela (F09)    4.000000 secs
## 4 Decimo (M10)     4.000000 secs
## 5 Sandro (M06)     3.998889 secs
```

```
Deer %>% 
  filter(!is.na(x_)) %>% 
  group_by(id) %>% 
  arrange(t_) %>% 
  mutate(x.dist = x_ - median(Deer$x_, na.rm = T),
         y.dist = y_ - median(Deer$y_, na.rm = T),
         dist = sqrt(x.dist^2 + y.dist^2),
         prop.d = dist/max(dist, na.rm = T)) %>% 
  ggplot(aes(x = x_, y = y_, color = id)) +
  geom_path() +
  coord_equal() +
  theme_linedraw()
```

```
Focal <- Deer %>% group_by(id) %>% arrange(t_) %>% mutate(row = 1:n()) %>% ungroup() %>% mutate(unique = 1:n())
train.italy <- Focal %>% group_by(id) %>% filter(row %in% 1:round(n()*.75))
test.italy <- Focal %>% filter(!unique %in% train.italy$unique)
```

```
# sr <- "+proj=utm +zone=32 +ellps=WGS84 +datum=WGS84 +units=m +no_defs"
# Landclass <- raster("/Users/willrogers/Documents/GitHub/SSurFace/ItalyLandcover.tif")
# Elevation <- raster("/Users/willrogers/Documents/GitHub/SSurFace/ItalyElevation.tif")
# Landclass <- raster::projectRaster(Landclass, crs = sr, method = "ngb")
# Elevation <- raster::projectRaster(Elevation, crs = sr, method = "bilinear")
# 
# LC <- crop(Landclass, extent(c(range(Deer$x_, na.rm = T) + 3200*c(-1,1),range(Deer$y_, na.rm = T)+ 3200*c(-1,1))))
# spts <- coordinates(LC)
# spdf <- SpatialPoints(coords = spts, proj4string = CRS(sr))
# forest <- (LC %in% c(23:25))
# forest <- rasterToPolygons(forest, fun = function(x){x == 1}, dissolve = T)
# # forest.l <- as(forest, "SpatialLinesDataFrame")
# forest_distance <- rgeos::gDistance(forest, spdf, byid=TRUE)
# 
# open <- (LC %in% c(18,29))
# open <- rasterToPolygons(open, fun = function(x){x == 1}, dissolve = T)
# # forest.l <- as(forest, "SpatialLinesDataFrame")
# open_distance <- rgeos::gDistance(open, spdf, byid=TRUE)
# 
# farm <- (LC %in% c(12:17,19:22))
# farm <- rasterToPolygons(farm, fun = function(x){x == 1}, dissolve = T)
# # forest.l <- as(forest, "SpatialLinesDataFrame")
# farm_distance <- rgeos::gDistance(farm, spdf, byid=TRUE)
# 
# # nforest <- (!LC %in% c(23:25))
# # nforest <- rasterToPolygons(nforest, fun = function(x){x == 1}, dissolve = T)
# # # nforest.l <- as(nforest, "SpatialLinesDataFrame")
# # nforest_distance <- rgeos::gDistance(nforest, spdf, byid=TRUE)
# #
# # ForestDistance <- setValues(LC, ifelse(forest_distance[,1] == 0, nforest_distance[,1], forest_distance[,1]))
# 
# ForestDistance <- setValues(LC, forest_distance)
# OpenDistance <- setValues(LC, open_distance)
# FarmDistance <- setValues(LC, farm_distance)
# 
# Elevation <- aggregate(Elevation, 2)
# 
# Landclass <- projectRaster(from = Landclass, to = Elevation,
#                            method = "ngb",
#                            format = "raster",
#                            overwrite = TRUE) # interpolate forest cover
# ForestDistance <- projectRaster(from = ForestDistance, to = Elevation,
#                                 method = "bilinear",
#                                 format = "raster",
#                                 overwrite = TRUE) # interpolate forest cover
# OpenDistance <- projectRaster(from = OpenDistance, to = Elevation,
#                                 method = "bilinear",
#                                 format = "raster",
#                                 overwrite = TRUE) # interpolate forest cover
# FarmDistance <- projectRaster(from = FarmDistance, to = Elevation,
#                                 method = "bilinear",
#                                 format = "raster",
#                                 overwrite = TRUE) # interpolate forest cover
# 
# rstack <- stack(Landclass, Elevation, ForestDistance, OpenDistance, FarmDistance)
# rstack$TRI <- terrain(rstack$ItalyElevation, opt = "TRI")
# rstack$Slope <- terrain(rstack$ItalyElevation, opt = "slope")
# rstack$Aspect <- cos((terrain(rstack$ItalyElevation, opt = "aspect", unit = "degrees")-35)/360)
# 
# names(rstack) <- c("Land", "Elev", "ForestD", "OpenD", "FarmD", "TRI", "Slope", "Aspect")
# 
# rstack <- crop(rstack, extent(c(range(Deer$x_, na.rm = T) + 1500*c(-1,1),range(Deer$y_, na.rm = T)+ 1500*c(-1,1))))
# 
# rstack$Artificial <- (rstack$Land %in% c(1:11))
# rstack$Farm <- (rstack$Land %in% c(12:17,19:22))
# rstack$Open <- (rstack$Land %in% c(18,29))
# rstack$Forest <- (rstack$Land %in% c(23:25))
# 
# plot(sqrt(rstack$OpenD))
# lines(Deer$x_, Deer$y_)
# saveRDS(rstack, "ItalyCov.RDS")
rstack.italy<- readRDS("Data and Intermediate RDS files/ItalyCov.RDS")
par(mfrow = c(2,2))
plot(sqrt(rstack.italy$ForestD))
plot((rstack.italy$Slope))
plot(sqrt(rstack.italy$ForestD))
points(train.italy$x_, train.italy$y_, col = "black", pch = ".")
points(test.italy$x_, test.italy$y_, col = "red", pch = ".")
plot((rstack.italy$Slope))
points(train.italy$x_, train.italy$y_, col = "black", pch = ".")
points(test.italy$x_, test.italy$y_, col = "red", pch = ".")
```

This follows the exact same processes as above, but does so within
map() calls. This is just a vectorized approach for multiple individuals
that helps avoid errors in copy+paste code chunks and is a bit cleaner
than a for-loop, though they all arrive at the same output.

```
set.seed(100)
ind <- train.italy %>% 
  group_by(id) %>% 
  ungroup() %>% 
  nest(data = -c("id")) %>% 
  mutate(trk = lapply(data,
                      function(d) {
                        make_track(d, x_, y_, t_, 
                                   season = season) %>%
                          track_resample(rate = hours(4), tolerance = minutes(10)) %>% 
                          steps_by_burst(keep_cols = "start")
                      })) %>% 
  mutate(dist = map(trk, function(x) {
    fatter.unif <- fit_distr(x$sl_, "unif")
    fatter.unif$params$min <- 0
    fatter.unif$params$max <- fatter.unif$params$max*1.25
    fatter.unif
  })) %>% 
  mutate(steps = map2(trk,dist, function(x,y) {
    x %>%
      random_steps(100, 
                   sl_distr = y) %>%
      extract_covariates(rstack.italy) 
  }))
```

```
## Warning: There were 10 warnings in `mutate()`.
## The first warning was:
## ℹ In argument: `dist = map(...)`.
## Caused by warning in `dunif()`:
## ! NaNs produced
## ℹ Run `dplyr::last_dplyr_warnings()` to see the 9 remaining warnings.
```

```
m <- ind %>%
  mutate(model = map(steps, function(x) {
    x %>% 
      filter(!is.na(sl_),
             !is.na(Slope)) %>% 
      fit_clogit(case_ ~ (sqrt(ForestD) + Slope) + (sl_) + log(sl_) + strata(step_id_), model = T)
  }))

mock.surface <- create_mock_surface(rstack.italy, F, list(x = xres(rstack.italy), y = yres(rstack.italy)))

pred.data <- get_cells(m$model[[1]], 
                       mock.surface,
                       rstack.italy)
pred.data <- pred.data %>% 
  arrange(cellnr)

cell.data <- get_cell_data(m$model[[1]], pred.data)
cell.data.list <- pbmclapply(as.list(1:dim(cell.data)[1]), function(x) cell.data[x[1],])
neighbors.found <- neighbor_lookup(mock.surface, cell.data, cell.data.list)

sparse.neighbors <- neighbor_finder(NA, 
                                    cell.data, 
                                    neighbors.found, 
                                    cell.data.list, 
                                    distance.override = max(sapply(m$trk, function(x) quantile(x$sl_, 0.95, na.rm = T))))
```

```
## [1] "Creating neighbor comparisons"
## [1] "Finding valid comparisons"
## [1] "Splitting cell.data into list"
## [1] "Running comparisons"
```

```
ssf.comparisons <- compile_ssf_comparisons(sparse.neighbors, cell.data)
predicted.surfaces <- m %>% 
  mutate(surfaces = map(model, function(x) {
    predict_ssf_comparisons(x, ssf.comparisons)
  }))
```

```
## [1] "Estimating probability surface"
## [1] "Compiling probability surface"
## [1] "Making sparse matrix for transitions"
## [1] "Estimating probability surface"
## [1] "Compiling probability surface"
## [1] "Making sparse matrix for transitions"
## [1] "Estimating probability surface"
## [1] "Compiling probability surface"
## [1] "Making sparse matrix for transitions"
## [1] "Estimating probability surface"
## [1] "Compiling probability surface"
## [1] "Making sparse matrix for transitions"
## [1] "Estimating probability surface"
## [1] "Compiling probability surface"
## [1] "Making sparse matrix for transitions"
```

```
graphs <- predicted.surfaces %>% 
  mutate(graph = map(surfaces, function(x) {
    A <- x$prob.matrix # matrix A
    d <- eigs(t(A), 1) # eigen decomp
    d.1 <- Re(d$vectors[,1]) # first vector
    prob.d.deer <- d.1/sum(d.1) # make sure it sums to 1
    ssd.raster.deer <- setValues(mock.surface, prob.d.deer)
    ssd.raster.deer
  }))

ssdpredictions.deer <- stack(graphs$graph)
names(ssdpredictions.deer) <- graphs$id
levelplot(((ssdpredictions.deer)))
```

```
par(oma = c(1,1,1,2))
plot(ssdpredictions.deer)
```

```
ssf.surfaces <- m %>% 
  mutate(surfaces = map2(model, trk, function(x,y) {
    a.data <- pred.data # grab prior matrix data 
    a.data$sl_ <- mean(y$sl_, na.rm = T) # use mean step length
    b.data <- a.data[1,] # set the first cell as the baseline
    
    log.rss <- amt::log_rss(x, # the model
                            a.data, # the raster data (including missing values)
                            b.data,  # a row of the raster data (excluding missing values)
                            ci = NA)
    ssf <-  exp(log.rss$df$log_rss)/sum(exp(log.rss$df$log_rss)) # turn into kernel after making values positive
    setValues(mock.surface, ssf)
  }))

ssfpredictions.deer <- stack(ssf.surfaces$surfaces)
names(ssfpredictions.deer) <- graphs$id
levelplot((ssfpredictions.deer))
```

```
par(oma = c(1,1,1,2))
plot(ssfpredictions.deer)
```

```
rsf.surfaces <- m %>% 
  mutate(surfaces = map2(data, trk, function(x,y) {
    rsf.data <- x %>% filter(!is.na(x_) & !(is.na(y_))) %>% 
      make_track(x_, y_, t_) %>% 
      random_points() %>% # defaults
      extract_covariates(rstack.italy) # grab distance
    
    rsf.m <- rsf.data %>% 
      fit_rsf(case_ ~  sqrt(ForestD) + Slope , model = T) # fit model
    probabilities <- exp(predict(rsf.m$model, newdata = pred.data))/(1+exp(predict(rsf.m$model, newdata = pred.data))) # get to probabilities
    probabilities <- ifelse(probabilities == "NaN", NA, probabilities)
    probabilities.kern <- probabilities/sum(probabilities, na.rm = T)
    setValues(mock.surface, probabilities.kern)
  }))

rsfpredictions.deer <- stack(rsf.surfaces$surfaces)
names(rsfpredictions.deer) <- graphs$id
levelplot((rsfpredictions.deer))
```

```
par(oma = c(1,1,1,2))
plot(rsfpredictions.deer)
```

##### 3.2.1 Comparisons

Here, we are comparing the ability of deer to predict their future
movements and to predict the movements of other deer in the same spatial
extent (i.e. same mountain)

```
full.deer <- stack(ssdpredictions.deer, ssfpredictions.deer, rsfpredictions.deer)
names(full.deer) <- paste(
  rep(c("SSD", "SSF", "RSF"), each = 5),
  rep(c("M03", "F09", "F10", "M06", "M10"), 3))

iter <- 100
bins <- 10
levelplot((full.deer))
```

```
pred.values <- as.data.frame(values(full.deer))

shannon.deer2 <- pred.values %>% 
  pivot_longer(all_of(colnames(pred.values))) %>% 
  mutate(model = str_split(name, "\\.",simplify = T)[,1],
         animal = str_split(name, "\\.",simplify = T)[,2]) %>% 
  group_by(model, animal) %>% 
  mutate(model = factor(model, c("SSD","SSF","RSF"))) %>% 
  summarize(shannon = -sum(value*log(value)))
```

```
## `summarise()` has grouped output by 'model'. You can override using the
## `.groups` argument.
```

```
shannon.deer2 %>% 
  ggplot(aes(x = model, y = shannon, color = animal, group = animal)) +
  geom_path() +
  geom_point() +
  scale_color_scico_d(begin = 0.5) +
  theme_classic()
```

```
pred.values %>% 
  pivot_longer(all_of(colnames(pred.values))) %>% 
  mutate(model = str_split(name, "\\.",simplify = T)[,1],
         animal = str_split(name, "\\.",simplify = T)[,2]) %>% 
  group_by(model, animal) %>% 
  mutate(model = factor(model, c("SSD","SSF","RSF"))) %>% 
  arrange(-value) %>% 
  mutate(cdf = cumsum(value)) %>% 
  summarize(total.50 = sum(cdf <= 0.5)) %>% 
  pivot_wider(names_from = model, values_from = total.50)
```

```
## `summarise()` has grouped output by 'model'. You can override using the
## `.groups` argument.
```

```
## # A tibble: 5 × 4
##   animal   SSD   SSF   RSF
##   <chr>  <int> <int> <int>
## 1 F09     1658  1969  1240
## 2 F10      309   592   565
## 3 M03      722  1204   619
## 4 M06      552  1005  1135
## 5 M10      985  1411  1168
```

```
use.area.deer2 <- pred.values %>% 
  pivot_longer(all_of(colnames(pred.values))) %>% 
  mutate(model = str_split(name, "\\.",simplify = T)[,1],
         animal = str_split(name, "\\.",simplify = T)[,2]) %>% 
  group_by(model, animal) %>% 
  mutate(model = factor(model, c("SSD","SSF","RSF"))) %>% 
  arrange(-value) %>% 
  mutate(cdf = cumsum(value)) %>% 
  summarize(sum = sum(cdf <= 0.5))
```

```
## `summarise()` has grouped output by 'model'. You can override using the
## `.groups` argument.
```

```
M03.in <- compare_spearman(train.italy %>% filter(!is.na(x_), id == "Agostino (M03)") %>% make_track(x_,y_) %>% ungroup(), 
                       subset(full.deer, seq(1, 15, by = 5)), iter, bins)
```

```
## Adding missing grouping variables: `id`
## `summarise()` has grouped output by 'name'. You can override using the
## `.groups` argument.
```

```
M03.id <- compare_spearman(test.italy %>% filter(!is.na(x_), id == "Agostino (M03)") %>% make_track(x_,y_) %>% ungroup(), 
                       subset(full.deer, seq(1, 15, by = 5)), iter, bins)
```

```
## `summarise()` has grouped output by 'name'. You can override using the
## `.groups` argument.
```

```
M03.out <- compare_spearman(Deer %>% filter(!is.na(x_), id != "Agostino (M03)") %>% make_track(x_,y_), 
                        subset(full.deer, seq(1, 15, by = 5)), iter, bins)
```

```
## `summarise()` has grouped output by 'name'. You can override using the
## `.groups` argument.
```

```
F09.in <- compare_spearman(train.italy %>% filter(!is.na(x_), id == "Daniela (F09)") %>% make_track(x_,y_) %>% ungroup(), 
                       subset(full.deer, seq(2, 15, by = 5)), iter, bins)
```

```
## Adding missing grouping variables: `id`
## `summarise()` has grouped output by 'name'. You can override using the
## `.groups` argument.
```

```
F09.id <- compare_spearman(test.italy %>% filter(!is.na(x_), id == "Daniela (F09)") %>% make_track(x_,y_) %>% ungroup(), 
                       subset(full.deer, seq(2, 15, by = 5)), iter, bins)
```

```
## `summarise()` has grouped output by 'name'. You can override using the
## `.groups` argument.
```

```
F09.out <- compare_spearman(Deer %>% filter(!is.na(x_), id != "Daniela (F09)") %>% make_track(x_,y_), 
                        subset(full.deer, seq(2, 15, by = 5)), iter, bins)
```

```
## `summarise()` has grouped output by 'name'. You can override using the
## `.groups` argument.
```

```
F10.in <- compare_spearman(train.italy %>% filter(!is.na(x_), id == "Alessandra (F10)") %>% make_track(x_,y_) %>% ungroup(), 
                       subset(full.deer, seq(3, 15, by = 5)), iter, bins)
```

```
## Adding missing grouping variables: `id`
## `summarise()` has grouped output by 'name'. You can override using the
## `.groups` argument.
```

```
F10.id <- compare_spearman(test.italy %>% filter(!is.na(x_), id == "Alessandra (F10)") %>% make_track(x_,y_) %>% ungroup(), 
                       subset(full.deer, seq(3, 15, by = 5)), iter, bins)
```

```
## `summarise()` has grouped output by 'name'. You can override using the
## `.groups` argument.
```

```
F10.out <- compare_spearman(Deer %>% filter(!is.na(x_), id != "Alessandra (F10)") %>% make_track(x_,y_), 
                        subset(full.deer, seq(3, 15, by = 5)), iter, bins)
```

```
## `summarise()` has grouped output by 'name'. You can override using the
## `.groups` argument.
```

```
M06.in <- compare_spearman(train.italy %>% filter(!is.na(x_), id == "Sandro (M06)") %>% make_track(x_,y_) %>% ungroup(), 
                       subset(full.deer, seq(4, 15, by = 5)), iter, bins)
```

```
## Adding missing grouping variables: `id`
## `summarise()` has grouped output by 'name'. You can override using the
## `.groups` argument.
```

```
M06.id <- compare_spearman(test.italy %>% filter(!is.na(x_), id == "Sandro (M06)") %>% make_track(x_,y_) %>% ungroup(), 
                       subset(full.deer, seq(4, 15, by = 5)), iter, bins)
```

```
## `summarise()` has grouped output by 'name'. You can override using the
## `.groups` argument.
```

```
M06.out <- compare_spearman(Deer %>% filter(!is.na(x_), id != "Sandro (M06)") %>% make_track(x_,y_), 
                        subset(full.deer, seq(4, 15, by = 5)), iter, bins)
```

```
## `summarise()` has grouped output by 'name'. You can override using the
## `.groups` argument.
```

```
M10.in <- compare_spearman(train.italy %>% filter(!is.na(x_), id == "Decimo (M10)") %>% make_track(x_,y_) %>% ungroup(), 
                       subset(full.deer, seq(5, 15, by = 5)), iter, bins)
```

```
## Adding missing grouping variables: `id`
## `summarise()` has grouped output by 'name'. You can override using the
## `.groups` argument.
```

```
M10.id <- compare_spearman(test.italy %>% filter(!is.na(x_), id == "Decimo (M10)") %>% make_track(x_,y_) %>% ungroup(), 
                       subset(full.deer, seq(5, 15, by = 5)), iter, bins)
```

```
## `summarise()` has grouped output by 'name'. You can override using the
## `.groups` argument.
```

```
M10.out <- compare_spearman(Deer %>% filter(!is.na(x_), id != "Decimo (M10)") %>% make_track(x_,y_), 
                        subset(full.deer, seq(5, 15, by = 5)), iter, bins)
```

```
## `summarise()` has grouped output by 'name'. You can override using the
## `.groups` argument.
```

```
correlations.deer <- rbind(M03.in,M03.id,M03.out,F09.in,F09.id,F09.out,F10.in,F10.id,F10.out,M06.in,M06.id,M06.out,M10.in,M10.id,M10.out) %>% 
  ungroup() %>% 
  mutate(id = rep(c("M03", "F09", "F10", "M06", "M10"), each = 3*3*100), 
         bootstrap = factor(bootstrap),
         in.out = rep(rep(c("In-Sample","Within Individual", "Between Individuals"), each = 3*100), 5), 
         model = factor(name, levels = c(1:3), labels = c("SSD", "SSF", "RSF"), ordered = T)) %>% 
  mutate(in.out = factor(in.out, levels = c("In-Sample", "Within Individual", "Between Individuals", "Between Contexts"), ordered = T)) 

# Mean log-use
M03.in <- compare_mean(train.italy %>% filter(!is.na(x_), id == "Agostino (M03)") %>% make_track(x_,y_) %>% ungroup(), 
                       subset(full.deer, seq(1, 15, by = 5)), iter)
```

```
## Adding missing grouping variables: `id`
```

```
M03.id <- compare_mean(test.italy %>% filter(!is.na(x_), id == "Agostino (M03)") %>% make_track(x_,y_) %>% ungroup(), 
                       subset(full.deer, seq(1, 15, by = 5)), iter)
M03.out <- compare_mean(Deer %>% filter(!is.na(x_), id != "Agostino (M03)") %>% make_track(x_,y_), 
                        subset(full.deer, seq(1, 15, by = 5)), iter)

F09.in <- compare_mean(train.italy %>% filter(!is.na(x_), id == "Daniela (F09)") %>% make_track(x_,y_) %>% ungroup(), 
                       subset(full.deer, seq(2, 15, by = 5)), iter)
```

```
## Adding missing grouping variables: `id`
```

```
F09.id <- compare_mean(test.italy %>% filter(!is.na(x_), id == "Daniela (F09)") %>% make_track(x_,y_) %>% ungroup(), 
                       subset(full.deer, seq(2, 15, by = 5)), iter)
F09.out <- compare_mean(Deer %>% filter(!is.na(x_), id != "Daniela (F09)") %>% make_track(x_,y_), 
                        subset(full.deer, seq(2, 15, by = 5)), iter)

F10.in <- compare_mean(train.italy %>% filter(!is.na(x_), id == "Alessandra (F10)") %>% make_track(x_,y_) %>% ungroup(), 
                       subset(full.deer, seq(3, 15, by = 5)), iter)
```

```
## Adding missing grouping variables: `id`
```

```
F10.id <- compare_mean(test.italy %>% filter(!is.na(x_), id == "Alessandra (F10)") %>% make_track(x_,y_) %>% ungroup(), 
                       subset(full.deer, seq(3, 15, by = 5)), iter)
F10.out <- compare_mean(Deer %>% filter(!is.na(x_), id != "Alessandra (F10)") %>% make_track(x_,y_), 
                        subset(full.deer, seq(3, 15, by = 5)), iter)

M06.in <- compare_mean(train.italy %>% filter(!is.na(x_), id == "Sandro (M06)") %>% make_track(x_,y_) %>% ungroup(), 
                       subset(full.deer, seq(4, 15, by = 5)), iter)
```

```
## Adding missing grouping variables: `id`
```

```
M06.id <- compare_mean(test.italy %>% filter(!is.na(x_), id == "Sandro (M06)") %>% make_track(x_,y_) %>% ungroup(), 
                       subset(full.deer, seq(4, 15, by = 5)), iter)
M06.out <- compare_mean(Deer %>% filter(!is.na(x_), id != "Sandro (M06)") %>% make_track(x_,y_), 
                        subset(full.deer, seq(4, 15, by = 5)), iter)

M10.in <- compare_mean(train.italy %>% filter(!is.na(x_), id == "Decimo (M10)") %>% make_track(x_,y_) %>% ungroup(), 
                       subset(full.deer, seq(5, 15, by = 5)), iter)
```

```
## Adding missing grouping variables: `id`
```

```
M10.id <- compare_mean(test.italy %>% filter(!is.na(x_), id == "Decimo (M10)") %>% make_track(x_,y_) %>% ungroup(), 
                       subset(full.deer, seq(5, 15, by = 5)), iter)
M10.out <- compare_mean(Deer %>% filter(!is.na(x_), id != "Decimo (M10)") %>% make_track(x_,y_), 
                        subset(full.deer, seq(5, 15, by = 5)), iter)

means.deer <- rbind(M03.in,M03.id,M03.out,F09.in,F09.id,F09.out,F10.in,F10.id,F10.out,M06.in,M06.id,M06.out,M10.in,M10.id,M10.out) %>% 
  mutate(id = rep(c("M03", "F09", "F10", "M06", "M10"), each = 3*3*100), 
         bootstrap = factor(bootstrap),
         in.out = rep(rep(c("In-Sample","Within Individual", "Between Individuals"), each = 3*100), 5), 
         model = factor(str_split(name,"\\.",simplify = T)[,1], levels = c("SSD", "SSF", "RSF"), ordered = T)) %>% 
  mutate(in.out = factor(in.out, levels = c("In-Sample", "Within Individual", "Between Individuals", "Between Contexts"), ordered = T)) 


correlations.deer$type <- "Spearman"
means.deer$type <- "Geometric Mean"

bound.comparisons.deer.2 <- rbind(correlations.deer, means.deer)

deer.data.2 <- bound.comparisons.deer.2 %>% 
  mutate(model = factor(model, levels = c("SSD", "SSF", "RSF"), ordered = T)) %>% 
  mutate(in.out = factor(in.out, levels = c("In-Sample", "Within Individual", "Between Individuals", "Between Contexts"), ordered = T)) %>% 
  ungroup() %>% 
  dplyr::select(-name) %>% 
  group_by(id, in.out, model, type) %>%
  pivot_wider(names_from = c(model), values_from = measure) %>% 
  pivot_longer(c(RSF, SSF)) %>% 
  mutate(delta = SSD-value) %>%
  group_by(id, in.out, name, type) %>%
  summarize(median = median(delta),
            lower = quantile(delta, 0.025),
            upper = quantile(delta, 0.975)) %>%
  # pivot_wider(names_from = type, values_from = c(median, lower, upper))
  mutate(model = factor(name, levels = c("SSF","RSF"), labels = c("SSD-SSF","SSD-RSF"), ordered = T)) %>% 
  group_by(in.out, id, type) %>% 
  mutate(top = median == max(median))
```

```
## `summarise()` has grouped output by 'id', 'in.out', 'name'. You can override
## using the `.groups` argument.
```

```
deer.data.2 %>% 
  arrange(model) %>% 
  ggplot(aes(x = model, y = median, ymax = upper, ymin = lower, shape = type, group = paste(id), color = id)) +
  geom_hline(yintercept = 0) +
  geom_path(position = position_dodge(0.2)) +
  geom_pointrange(position = position_dodge(0.2)) +
  facet_grid(type~in.out, scales = "free_y") +
  labs(x = "Prediction Comparison",
       y = "Difference from SSD Prediction\n(Median and 95% Quantile)",
       shape = "Comparison Heuristic") + 
  theme_classic() +
  scale_alpha_discrete(guide = F, range = c(0.5, 1))
```

```
## Warning: Using alpha for a discrete variable is not advised.
```

##### 3.2.2 Storing some output and plots for later

```
ssd.raster.deer.complex.df <- as.data.frame(rasterToPoints(full.deer$SSD.M10))
ssf.prob.raster.deer.complex.df <- as.data.frame(rasterToPoints(full.deer$SSF.M10))
rsf.prob.raster.deer.complex.df <- as.data.frame(rasterToPoints(full.deer$RSF.M10))

colnames(ssd.raster.deer.complex.df)[3] <- "layer"
colnames(ssf.prob.raster.deer.complex.df)[3] <- "layer"
colnames(rsf.prob.raster.deer.complex.df)[3] <- "layer"

min.max.deer.complex <- rbind(ssd.raster.deer.complex.df, ssf.prob.raster.deer.complex.df, rsf.prob.raster.deer.complex.df)
min.deer.complex <- min(min.max.deer.complex$layer)
max.deer.complex <- max(min.max.deer.complex$layer)

ssd.deer.complex.plot <- ssd.raster.deer.complex.df %>% 
  ggplot() +
  geom_raster(aes(x = (x-min(x))/1000, y = (y-min(y))/1000, fill = layer), color = NA) +
  coord_equal() +
  scale_fill_viridis_c(option = "H", name = "Kernel Probability",
                       limits = c(min.deer.complex,max.deer.complex)) +
  # geom_point(data = deer, mapping = aes(x = x_, y = y_), color = "green", size = size.point, alpha = alpha.point) +
  # geom_path(data = ssf1, mapping = aes(x = x1_, y = y1_), color = "white", size = 0.1, alpha = 0.1) +
  theme.current+
  labs(x = "Easting (km)", y = "Northing (km)")
```

```
## Warning in geom_raster(aes(x = (x - min(x))/1000, y = (y - min(y))/1000, :
## Ignoring unknown parameters: `colour`
```

```
ssf.deer.complex.plot <- ssf.prob.raster.deer.complex.df %>% 
  ggplot() +
  geom_raster(aes(x = (x-min(x))/1000, y = (y-min(y))/1000, fill = layer), color = NA) +
  coord_equal() +
  scale_fill_viridis_c(option = "H", name = "Kernel Probability",
                       limits = c(min.deer.complex,max.deer.complex)) +
  # geom_point(data = deer, mapping = aes(x = x_, y = y_), color = "green", size = size.point, alpha = alpha.point) +
  # geom_path(data = ssf1, mapping = aes(x = x1_, y = y1_), color = "white", size = 0.1, alpha = 0.1) +
  theme.current+
  labs(x = "Easting (km)", y = "Northing (km)")
```

```
## Warning in geom_raster(aes(x = (x - min(x))/1000, y = (y - min(y))/1000, :
## Ignoring unknown parameters: `colour`
```

```
rsf.deer.complex.plot <- rsf.prob.raster.deer.complex.df %>% 
  ggplot() +
  geom_raster(aes(x = (x-min(x))/1000, y = (y-min(y))/1000, fill = layer), color = NA) +
  coord_equal() +
  scale_fill_viridis_c(option = "H", name = "Kernel Probability",
                       limits = c(min.deer.complex,max.deer.complex)) +
  # geom_point(data = deer, mapping = aes(x = x_, y = y_), color = "green", size = size.point, alpha = alpha.point) +
  # geom_path(data = ssf1, mapping = aes(x = x1_, y = y1_), color = "white", size = 0.1, alpha = 0.1) +
  theme.current+
  labs(x = "Easting (km)", y = "Northing (km)")
```

```
## Warning in geom_raster(aes(x = (x - min(x))/1000, y = (y - min(y))/1000, :
## Ignoring unknown parameters: `colour`
```

```
complex.deer <- ggarrange(rsf.deer.complex.plot, ssf.deer.complex.plot, ssd.deer.complex.plot, common.legend = T, nrow = 1, legend = "left")
complex.deer
```

```
# rm(ssdpredictions.deer, ssfpredictions.deer, rsfpredictions.deer, rsf.surfaces, ssf.surfaces, mock.surface, pred.data, cell.data, cell.data.list, neighbors.found, sparse.neighbors, ssf.comparisons, predicted.surfaces, graphs, ssdpredictions.deer, m, ind, rstack, Deer, Focal, train, test)
```

#### 3.3 Fisher

We can scale up our projections to think about other individuals in
*different* places.

```
# fisher <- read_csv("Martes pennanti LaPoint New York.csv")
# fisher$t_ <- as.POSIXct(fisher$timestamp)
# fisher$id <- fisher$`individual-local-identifier`
# fisher$x_ <- fisher$`utm-easting`
# fisher$y_ <- fisher$`utm-northing`
# fisher$group <- ifelse(fisher$id == "M5", "Non-focal", "Focal")
# 
# saveRDS(fisher, "fisher.data.RDS")
fisher <- readRDS("Data and Intermediate RDS files/fisher.data.RDS")

fisher <- fisher %>% dplyr::select(c("id", "group", "x_", "y_", "t_"))
fisher.focal <- fisher %>% filter(id != "M5")
fisher.nonfocal <- fisher %>% filter(id == "M5")

fisher %>%
  arrange(t_) %>%
  ggplot(aes(x = x_, y = y_, color = id)) +
  geom_path() +
  coord_equal() +
  facet_grid(group~.) +
  theme_linedraw()
```

```
## Warning: Removed 208 rows containing missing values or values outside the scale range
## (`geom_path()`).
```

Filtering out historic movements from the future movements we want to
predict

```
Focal <- fisher %>% group_by(id) %>% filter(!is.na(x_)) %>% arrange(t_) %>% mutate(row = 1:n()) %>% ungroup() %>% mutate(unique = 1:n()) 
train.fisher <- Focal %>% group_by(id) %>% filter(row %in% 1:round(n()*.75))
test.fisher <- Focal %>% filter(!unique %in% train.fisher$unique)

train.fisher %>%
  group_by(id) %>%
  tally()
```

```
## # A tibble: 8 × 2
##   id        n
##   <chr> <int>
## 1 F1     1012
## 2 F2     2253
## 3 F3     1126
## 4 M1      689
## 5 M2     1228
## 6 M3     1827
## 7 M4     6718
## 8 M5     9824
```

```
test.fisher %>%
  group_by(id) %>%
  tally()
```

```
## # A tibble: 8 × 2
##   id        n
##   <chr> <int>
## 1 F1      337
## 2 F2      751
## 3 F3      375
## 4 M1      230
## 5 M2      410
## 6 M3      609
## 7 M4     2240
## 8 M5     3275
```

We can also read in geospatial info, and alter it to suite reasonable
info for species movements.

```
# Elevation <- raster("/Users/willrogers/Documents/GitHub/SSurFace/n42_w074_1arc_v2.tif")
# Popden <- raster("/Users/willrogers/Documents/GitHub/SSurFace/Popden.tif")
# Landcover <- raster("/Users/willrogers/Documents/GitHub/SSurFace/Landcover.tif")
# HumanDistance <- raster("/Users/willrogers/Documents/GitHub/SSurFace/FisherHumanDistance.tif",
#                         crs = crs("+proj=utm +zone=18 +datum=NAD83 +units=m +no_defs"))
# WetlandDistance <- raster("/Users/willrogers/Documents/GitHub/SSurFace/FisherWetlandDistance.tif",
#                           crs = crs("+proj=utm +zone=18 +datum=NAD83 +units=m +no_defs"))
# ForestDistance <- raster("/Users/willrogers/Documents/GitHub/SSurFace/FisherForestDistance.tif",
#                          crs = crs("+proj=utm +zone=18 +datum=NAD83 +units=m +no_defs"))
# sr <- "+proj=utm +zone=18 +ellps=GRS80 +datum=NAD83 +units=m +no_defs"
# Elevation <- raster::projectRaster(Elevation, crs = sr, method = "bilinear")
# Popden <- raster::projectRaster(Popden, crs = sr, method = "bilinear")
# Landcover <- raster::projectRaster(Landcover, crs = sr, method = "ngb")
# HumanDistance <- raster::projectRaster(HumanDistance, crs = sr, method = "bilinear")
# WetlandDistance <- raster::projectRaster(WetlandDistance, crs = sr, method = "bilinear")
# ForestDistance <- raster::projectRaster(ForestDistance, crs = sr, method = "bilinear")
# 
# focal.extent <- extent(c(range(fisher.focal$x_, na.rm = T) + 4000*c(-1,1),range(fisher.focal$y_, na.rm = T)+ 4000*c(-1,1)))
# non.focal.extent <- extent(c(range(fisher.nonfocal$x_, na.rm = T)+ 4000*c(-1,1),range(fisher.nonfocal$y_, na.rm = T)+ 4000*c(-1,1)))
# Elev.focal <- crop(Elevation, focal.extent)
# PopD.focal <- crop(Popden, focal.extent)
# Land.focal <- crop(Landcover, focal.extent)
# HumanDistance.focal <- crop(HumanDistance, focal.extent)
# WetlandDistance.focal <- crop(WetlandDistance, focal.extent)
# ForestDistance.focal <- crop(ForestDistance, focal.extent)
# Elev.nfocal <- crop(Elevation, non.focal.extent)
# PopD.nfocal <- crop(Popden, non.focal.extent)
# Land.nfocal <- crop(Landcover, non.focal.extent)
# HumanDistance.nfocal <- crop(HumanDistance, non.focal.extent)
# WetlandDistance.nfocal <- crop(WetlandDistance, non.focal.extent)
# ForestDistance.nfocal <- crop(ForestDistance, non.focal.extent)
# 
# Elev.focal. <- aggregate(Elev.focal,4)
# Elev.nfocal. <- aggregate(Elev.nfocal,4)
# 
# pop.repro <- projectRaster(from = PopD.focal, to = Elev.focal.,
#                            method = "bilinear",
#                            format = "raster",
#                            overwrite = TRUE) # interpolate population density
# landuse.repro <- projectRaster(from = Land.focal, to = Elev.focal.,
#                                method = "ngb",
#                                format = "raster",
#                                overwrite = TRUE) # interpolate forest cover
# HumanDistance.focal. <- projectRaster(from = HumanDistance.focal, to = Elev.focal.,
#                                       method = "bilinear",
#                                       format = "raster",
#                                       overwrite = TRUE) # interpolate forest cover
# WetlandDistance.focal. <- projectRaster(from = WetlandDistance.focal, to = Elev.focal.,
#                                         method = "bilinear",
#                                         format = "raster",
#                                         overwrite = TRUE) # interpolate forest cover
# ForestDistance.focal. <- projectRaster(from = ForestDistance.focal, to = Elev.focal.,
#                                        method = "bilinear",
#                                        format = "raster",
#                                        overwrite = TRUE) # interpolate forest cover
# 
# rstack.focal <- stack(Elev.focal., pop.repro,landuse.repro, HumanDistance.focal., WetlandDistance.focal., ForestDistance.focal.)
# 
# popn.repro <- projectRaster(from = PopD.nfocal, to = Elev.nfocal.,
#                             method = "bilinear",
#                             format = "raster",
#                             overwrite = TRUE) # interpolate population density
# landusen.repro <- projectRaster(from = Land.nfocal, to = Elev.nfocal.,
#                                 method = "ngb",
#                                 format = "raster",
#                                 overwrite = TRUE) # interpolate forest cover
# HumanDistance.nfocal. <- projectRaster(from = HumanDistance.nfocal, to = Elev.nfocal.,
#                                        method = "bilinear",
#                                        format = "raster",
#                                        overwrite = TRUE) # interpolate forest cover
# WetlandDistance.nfocal. <- projectRaster(from = WetlandDistance.nfocal, to = Elev.nfocal.,
#                                          method = "bilinear",
#                                          format = "raster",
#                                          overwrite = TRUE) # interpolate forest cover
# ForestDistance.nfocal. <- projectRaster(from = ForestDistance.nfocal, to = Elev.nfocal.,
#                                         method = "bilinear",
#                                         format = "raster",
#                                         overwrite = TRUE) # interpolate forest cover
# 
# rstack.nonfocal <- stack(Elev.nfocal., popn.repro, landusen.repro, HumanDistance.nfocal., WetlandDistance.nfocal., ForestDistance.nfocal.)
# 
# rstack.focal$Slope <- terrain(rstack.focal$n42_w074_1arc_v2, opt = "Slope")
# rstack.nonfocal$Slope <- terrain(rstack.nonfocal$n42_w074_1arc_v2, opt = "Slope")
# 
# names(rstack.focal) <- c("Elevation", "PopDen", "Landcover", "DistHuman", "DistWetland", "DistForest", "Slope")
# names(rstack.nonfocal) <- c("Elevation", "PopDen", "Landcover", "DistHuman", "DistWetland", "DistForest", "Slope")
# 
# saveRDS(rstack.focal, "Focal.Landscape.RDS")
# saveRDS(rstack.nonfocal, "Nonfocal.Landscape.RDS")

rstack.focal <- readRDS("Data and Intermediate RDS files/Focal.Landscape.RDS")
rstack.nonfocal <- readRDS("Data and Intermediate RDS files/Nonfocal.Landscape.RDS")
```

Defining the prediction landscapes and some preliminary plotting to
verify.

```
focal.extent <- extent(c(range(fisher.focal$x_, na.rm = T) + 1000*c(-1,1),range(fisher.focal$y_, na.rm = T)+ 1000*c(-1,1)))
non.focal.extent <- extent(c(range(fisher.nonfocal$x_, na.rm = T)+ 1000*c(-1,1),range(fisher.nonfocal$y_, na.rm = T)+ 1000*c(-1,1)))

focal <- rstack.focal
focal <- crop(focal, focal.extent)

nonfocal <- rstack.nonfocal

nonfocal <- crop(nonfocal, non.focal.extent)

dist.nat <- ifelse(values(focal$DistWetland) <= values(focal$DistForest), values(focal$DistWetland), values(focal$DistForest))
focal$DistForWet <- setValues(focal$DistWetland, dist.nat)

dist.nat <- ifelse(values(nonfocal$DistWetland) <= values(nonfocal$DistForest), values(nonfocal$DistWetland), values(nonfocal$DistForest))
nonfocal$DistForWet <- setValues(nonfocal$DistWetland, dist.nat)

focal <- crop(focal, focal.extent)
nonfocal <- crop(nonfocal, non.focal.extent)

focal$ForWet <- focal$DistForWet == 0
nonfocal$ForWet <- nonfocal$DistForWet == 0

par(mfrow = c(1,2))
plot(sqrt(focal$DistForest))
plot(sqrt(focal$DistForest))
points(train.fisher$x_, train.fisher$y_, col = "black", pch = ".")
points(test.fisher$x_, test.fisher$y_, col = "red", pch = ".")
```

```
plot(focal)
```

```
par(mfrow = c(1,2), oma = c(0,0,0,1))
plot(sqrt(nonfocal$DistForest))
plot(sqrt(nonfocal$DistForest))
points(test.fisher$x_, test.fisher$y_, col = "red", pch = ".")
```

```
plot(nonfocal)
```

Again, we are using map to contain the same pipeline as above.

```
ind <- train.fisher %>%
  nest(data = -c("id")) %>%
  mutate(trk = lapply(data,
                      function(d) {
                        difference <- round(as.numeric(median(diff(d$t_), na.rm = T), units = "mins"))
                        make_track(d, x_, y_, t_, group) %>%
                          track_resample(rate = minutes(difference), tolerance = minutes(2)) %>%
                          steps_by_burst(keep_cols = "start")
                      })) %>%
  mutate(dist = map(trk, function(x) {
    fatter.unif <- fit_distr(x$sl_, "unif")
    fatter.unif$params$min <- 0
    fatter.unif$params$max <- fatter.unif$params$max*1.25
    fatter.unif
  })) %>% 
  filter(id != "M5") %>%
  mutate(steps = map2(trk, dist, function(x,y) {
    if("Focal" %in% x$group) {
      data <- x %>%
        random_steps(100,
                     sl_distr = y) %>%
        extract_covariates(focal)
    }
    if("Non-focal" %in% x$group) {
      data <- x %>%
        random_steps(100,
                     sl_distr = y) %>%
        extract_covariates(nonfocal)
    }
    data
  }))
```

```
## Warning: There were 16 warnings in `mutate()`.
## The first warning was:
## ℹ In argument: `dist = map(...)`.
## ℹ In group 1: `id = "F1"`.
## Caused by warning in `dunif()`:
## ! NaNs produced
## ℹ Run `dplyr::last_dplyr_warnings()` to see the 15 remaining warnings.
```

```
## Warning: There were 7 warnings in `mutate()`.
## The first warning was:
## ℹ In argument: `steps = map2(...)`.
## ℹ In group 1: `id = "F1"`.
## Caused by warning in `random_steps.bursted_steps_xyt()`:
## ! Some bursts contain < 3 steps and will be removed
## ℹ Run `dplyr::last_dplyr_warnings()` to see the 6 remaining warnings.
```

```
m <- ind %>% 
  filter(id != "M5") %>%
  mutate(model = map(steps, function(x) {
    x %>% drop_na() %>%
      fit_clogit(case_ ~ sqrt(DistForest) + sl_ + log(sl_) + strata(step_id_), model = T)
  }))

mock.surface <- create_mock_surface(focal, F, list(x = xres(focal), y = yres(focal)))

pred.data <- get_cells(m$model[[1]],
                       mock.surface,
                       focal)
pred.data <- pred.data %>%
  arrange(cellnr)
cell.data <- get_cell_data(m$model[[1]], pred.data)
cell.data.list <- pbmclapply(as.list(1:dim(cell.data)[1]), function(x) cell.data[x[1],])
neighbors.found <- neighbor_lookup(mock.surface, cell.data, cell.data.list = cell.data.list)

sparse.neighbors <- neighbor_finder(NA, cell.data, neighbors.found, cell.data.list = cell.data.list, distance.override = max(sapply(m$trk, function(x) quantile(x$sl_, 0.95))))
```

```
## [1] "Creating neighbor comparisons"
## [1] "Finding valid comparisons"
## [1] "Using inputted list of cell data"
## [1] "Running comparisons"
```

```
ssf.comparisons <- compile_ssf_comparisons(sparse.neighbors, cell.data)

predicted.surfaces <- m %>%
  filter(id != "M5") %>%
  mutate(surfaces = map(model, function(x) {
    predict_ssf_comparisons(x, ssf.comparisons)
  }))
```

```
## [1] "Estimating probability surface"
## [1] "Compiling probability surface"
## [1] "Making sparse matrix for transitions"
## [1] "Estimating probability surface"
## [1] "Compiling probability surface"
## [1] "Making sparse matrix for transitions"
## [1] "Estimating probability surface"
## [1] "Compiling probability surface"
## [1] "Making sparse matrix for transitions"
## [1] "Estimating probability surface"
## [1] "Compiling probability surface"
## [1] "Making sparse matrix for transitions"
## [1] "Estimating probability surface"
## [1] "Compiling probability surface"
## [1] "Making sparse matrix for transitions"
## [1] "Estimating probability surface"
## [1] "Compiling probability surface"
## [1] "Making sparse matrix for transitions"
## [1] "Estimating probability surface"
## [1] "Compiling probability surface"
## [1] "Making sparse matrix for transitions"
```

```
graphs <- predicted.surfaces %>%
  filter(id != "M5") %>%
  mutate(graph = map(surfaces, function(x) {
    A <- x$prob.matrix # matrix A
    d <- eigs(t(A), 1) # eigen decomp
    d.1 <- Re(d$vectors[,1]) # first vector
    prob.d.deer <- d.1/sum(d.1) # make sure it sums to 1
    ssd.raster.deer <- setValues(mock.surface, prob.d.deer)
    ssd.raster.deer
  }))
ssdpredictions <- stack(graphs$graph)
names(ssdpredictions) <- graphs$id
par(oma = c(0,0,0,2))
plot((ssdpredictions))
```

```
levelplot((ssdpredictions))
```

```
ssf.surfaces <- m %>%
  filter(id != "M5") %>%
  mutate(surfaces = map2(model, trk, function(x,y) {
    a.data <- pred.data # grab prior matrix data
    a.data$sl_ <- mean(y$sl_, na.rm = T) # use mean step length
    b.data <- a.data[1,] # set the first cell as the baseline

    log.rss <- amt::log_rss(x, # the model
                            a.data, # the raster data (including missing values)
                            b.data,  # a row of the raster data (excluding missing values)
                            ci = NA)
    ssf <-  exp(log.rss$df$log_rss)/sum(exp(log.rss$df$log_rss)) # turn into kernel after making values positive
    setValues(mock.surface, ssf)
  }))

ssfpredictions <- stack(ssf.surfaces$surfaces)
names(ssfpredictions) <- graphs$id
par(oma = c(0,0,0,2))
plot((ssfpredictions))
```

```
levelplot((ssfpredictions))
```

```
rsf.surfaces <- m %>%
  filter(id != "M5") %>%
  mutate(surfaces = map2(data, trk, function(x,y) {
    if("Focal" %in% x$group) {
      rsf.data <- x %>% filter(!is.na(x_) & !(is.na(y_))) %>%
        make_track(x_, y_, t_) %>%
        random_points() %>% # defaults
        extract_covariates(focal) # grab distance
    }
    if("Non-focal" %in% x$group) {
      rsf.data <- x %>% filter(!is.na(x_) & !(is.na(y_))) %>%
        make_track(x_, y_, t_) %>%
        random_points() %>% # defaults
        extract_covariates(nonfocal) # grab distance
    }

    rsf.m <- rsf.data %>%
      fit_rsf(case_ ~  sqrt(DistForest), model = T) # fit model
    probabilities <- exp(predict(rsf.m$model, newdata = pred.data))/(1+exp(predict(rsf.m$model, newdata = pred.data))) # get to probabilities
    probabilities <- ifelse(probabilities == "NaN", NA, probabilities)
    probabilities.kern <- probabilities/sum(probabilities, na.rm = T)
    setValues(mock.surface, probabilities.kern)
  }))


rsfpredictions <- stack(rsf.surfaces$surfaces)
names(rsfpredictions) <- graphs$id
par(oma = c(0,0,0,2))
plot((rsfpredictions))
```

```
levelplot((rsfpredictions))
```

```
levelplot(stack((ssdpredictions)[[1]],(ssfpredictions)[[1]],(rsfpredictions)[[1]]))
```

Now that we have the focal predictions (a shared landscape), let’s
look at a nonfocal landscape (a different landscape)

```
mock.surface. <- create_mock_surface(nonfocal, F, list(x = xres(nonfocal), y = yres(nonfocal)))

pred.data. <- get_cells(m$model[[1]],
                       mock.surface.,
                       nonfocal)
pred.data. <- pred.data. %>%
  arrange(cellnr)
cell.data. <- get_cell_data(m$model[[1]], pred.data.)
cell.data.list. <- pbmclapply(as.list(1:dim(cell.data.)[1]), function(x) cell.data.[x[1],])
neighbors.found. <- neighbor_lookup(mock.surface., cell.data., cell.data.)
sparse.neighbors. <- neighbor_finder(NA, cell.data., neighbors.found., quantile = 0.95, cell.data.list = cell.data.list., distance.override = max(sapply(m$trk, function(x) quantile(x$sl_, 0.95))))
```

```
## [1] "Creating neighbor comparisons"
## [1] "Finding valid comparisons"
## [1] "Using inputted list of cell data"
## [1] "Running comparisons"
```

```
ssf.comparisons. <- compile_ssf_comparisons(sparse.neighbors., cell.data.)

predicted.surfaces. <- m %>%
  filter(id != "M5") %>%
  mutate(surfaces = map(model, function(x) {
    predict_ssf_comparisons(x, ssf.comparisons.)
  }))
```

```
## [1] "Estimating probability surface"
## [1] "Compiling probability surface"
## [1] "Making sparse matrix for transitions"
## [1] "Estimating probability surface"
## [1] "Compiling probability surface"
## [1] "Making sparse matrix for transitions"
## [1] "Estimating probability surface"
## [1] "Compiling probability surface"
## [1] "Making sparse matrix for transitions"
## [1] "Estimating probability surface"
## [1] "Compiling probability surface"
## [1] "Making sparse matrix for transitions"
## [1] "Estimating probability surface"
## [1] "Compiling probability surface"
## [1] "Making sparse matrix for transitions"
## [1] "Estimating probability surface"
## [1] "Compiling probability surface"
## [1] "Making sparse matrix for transitions"
## [1] "Estimating probability surface"
## [1] "Compiling probability surface"
## [1] "Making sparse matrix for transitions"
```

```
graphs. <- predicted.surfaces. %>%
  filter(id != "M5") %>%
  mutate(graph = map(surfaces, function(x) {
    A <- x$prob.matrix # matrix A
    d <- eigs(t(A), 1) # eigen decomp
    d.1 <- Re(d$vectors[,1]) # first vector
    prob.d.deer <- d.1/sum(d.1) # make sure it sums to 1
    ssd.raster.deer <- setValues(mock.surface., prob.d.deer)
    ssd.raster.deer
  }))

ssdpredictions. <- stack(graphs.$graph)
names(ssdpredictions.) <- graphs.$id
par(oma = c(0,0,0,2))
plot((ssdpredictions.))
```

```
ssf.surfaces. <- m %>%
  filter(id != "M5") %>%
  mutate(surfaces = map2(model, trk, function(x,y) {
    a.data <- pred.data. # grab prior matrix data
    a.data$sl_ <- mean(y$sl_, na.rm = T) # use mean step length
    b.data <- a.data[1,] # set the first cell as the baseline

    log.rss <- amt::log_rss(x, # the model
                            a.data, # the raster data (including missing values)
                            b.data,  # a row of the raster data (excluding missing values)
                            ci = NA)
    ssf <-  exp(log.rss$df$log_rss)/sum(exp(log.rss$df$log_rss)) # turn into kernel after making values positive
    setValues(mock.surface., ssf)
  }))

ssfpredictions. <- stack(ssf.surfaces.$surfaces)
names(ssfpredictions.) <- graphs$id
par(oma = c(0,0,0,2))
plot((ssfpredictions.))
```

```
rsf.surfaces. <- m %>% 
  filter(id != "M5") %>%
  mutate(surfaces = map2(data, trk, function(x,y) {
    if("Focal" %in% x$group) {
      rsf.data <- x %>% filter(!is.na(x_) & !(is.na(y_))) %>% 
        make_track(x_, y_, t_) %>% 
        random_points() %>% # defaults
        extract_covariates(focal) # grab distance
    }
    if("Non-focal" %in% x$group) {
      rsf.data <- x %>% filter(!is.na(x_) & !(is.na(y_))) %>% 
        make_track(x_, y_, t_) %>% 
        random_points() %>% # defaults
        extract_covariates(nonfocal) # grab distance
    }
    
    rsf.m <- rsf.data %>% 
      fit_rsf(case_ ~ sqrt(DistForest), model = T) # fit model
    probabilities <- exp(predict(rsf.m$model, newdata = pred.data.))/(1+exp(predict(rsf.m$model, newdata = pred.data.))) # get to probabilities
    probabilities <- ifelse(probabilities == "NaN", NA, probabilities)
    probabilities.kern <- probabilities/sum(probabilities, na.rm = T)
    setValues(mock.surface., probabilities.kern)
  }))


rsfpredictions. <- stack(rsf.surfaces.$surfaces)
names(rsfpredictions.) <- graphs.$id
par(oma = c(0,0,0,2))
plot((rsfpredictions.))
```

##### 3.3.1 Comparisons

Now we can compare all of the abilities of the surfaces to the
observed intensity of use for individuals in the future (in), for other
individuals in the same place (out), and for new environments (new).

```
focal.pred <- stack(ssdpredictions, ssfpredictions, rsfpredictions)
non.focal.pred <- stack(ssdpredictions., ssfpredictions., rsfpredictions.)
names(focal.pred) <- paste(rep(c("SSD", "SSF", "RSF"), each = 7), rep(graphs$id, 3), "Focal")
names(non.focal.pred) <- paste(rep(c("SSD", "SSF", "RSF"), each = 7), rep(graphs$id, 3), "Nonocal")

pred.values <- as.data.frame(values(focal.pred))

shannon.fisher <- pred.values %>% 
  pivot_longer(all_of(colnames(pred.values))) %>% 
  mutate(model = str_split(name, "\\.",simplify = T)[,1],
         animal = str_split(name, "\\.",simplify = T)[,2]) %>% 
  group_by(model, animal) %>% 
  mutate(model = factor(model, c("SSD","SSF","RSF"))) %>% 
  summarize(shannon = -sum(value*log(value))) %>% 
  group_by(animal)
```

```
## `summarise()` has grouped output by 'model'. You can override using the
## `.groups` argument.
```

```
shannon.fisher %>% 
  ggplot(aes(x = model, y = shannon, color = animal, group = animal)) +
  geom_path() +
  geom_point() +
  scale_color_scico_d(begin = 0.5) +
  theme_classic()
```

```
non.focal.pred.values <- as.data.frame(values(non.focal.pred))

non.focal.pred.values %>% 
  pivot_longer(all_of(colnames(non.focal.pred.values))) %>% 
  mutate(model = str_split(name, "\\.",simplify = T)[,1],
         animal = str_split(name, "\\.",simplify = T)[,2]) %>% 
  group_by(model, animal) %>% 
  mutate(model = factor(model, c("SSD","SSF","RSF"))) %>% 
  summarize(shannon = -sum(value*log(value))) %>% 
  group_by(animal) %>% 
  ggplot(aes(x = model, y = shannon, color = animal, group = animal)) +
  geom_path() +
  geom_point() +
  scale_color_scico_d(begin = 0.5) +
  theme_linedraw()
```

```
## `summarise()` has grouped output by 'model'. You can override using the
## `.groups` argument.
```

```
use.area.fisher <- pred.values %>% 
  pivot_longer(all_of(colnames(pred.values))) %>% 
  mutate(model = str_split(name, "\\.",simplify = T)[,1],
         animal = str_split(name, "\\.",simplify = T)[,2]) %>% 
  group_by(model, animal) %>% 
  mutate(model = factor(model, c("SSD","SSF","RSF"))) %>% 
  arrange(-value) %>% 
  mutate(cdf = cumsum(value)) %>% 
  summarize(sum = sum(cdf <= 0.5))
```

```
## `summarise()` has grouped output by 'model'. You can override using the
## `.groups` argument.
```

```
use.area.fisher %>% 
  ggplot(aes(x = model, y = sum, color = animal, group = animal)) +
  geom_point() +
  geom_path()
```

```
M1.in <- compare_spearman(train.fisher %>% filter(!is.na(x_), id == "M1", id != "M5") %>% make_track(x_,y_) %>% ungroup(),
                       subset(focal.pred, seq(1, 21, by = 7)), iter)
```

```
## Adding missing grouping variables: `id`
```

```
## [1] "unique bins less than requested bins, decreasing bins to minimum"
```

```
## `summarise()` has grouped output by 'name'. You can override using the
## `.groups` argument.
```

```
M1.id <- compare_spearman(test.fisher %>% filter(!is.na(x_), id == "M1", id != "M5") %>% make_track(x_,y_),
                       subset(focal.pred, seq(1, 21, by = 7)), iter)
```

```
## [1] "unique bins less than requested bins, decreasing bins to minimum"
```

```
## `summarise()` has grouped output by 'name'. You can override using the
## `.groups` argument.
```

```
M1.out <- compare_spearman(Focal %>% filter(!is.na(x_), id != "M1", id != "M5") %>% make_track(x_,y_),
                        subset(focal.pred, seq(1, 21, by = 7)), iter)
```

```
## [1] "unique bins less than requested bins, decreasing bins to minimum"
```

```
## `summarise()` has grouped output by 'name'. You can override using the
## `.groups` argument.
```

```
M4.in <- compare_spearman(train.fisher %>% filter(!is.na(x_), id == "M4", id != "M5") %>% make_track(x_,y_) %>% ungroup(),
                       subset(focal.pred, seq(2, 21, by = 7)), iter)
```

```
## Adding missing grouping variables: `id`
```

```
## [1] "unique bins less than requested bins, decreasing bins to minimum"
```

```
## `summarise()` has grouped output by 'name'. You can override using the
## `.groups` argument.
```

```
M4.id <- compare_spearman(test.fisher %>% filter(!is.na(x_), id == "M4", id != "M5") %>% make_track(x_,y_),
                       subset(focal.pred, seq(2, 21, by = 7)), iter)
```

```
## [1] "unique bins less than requested bins, decreasing bins to minimum"
```

```
## `summarise()` has grouped output by 'name'. You can override using the
## `.groups` argument.
```

```
M4.out <- compare_spearman(Focal %>% filter(!is.na(x_), id != "M4", id != "M5") %>% make_track(x_,y_),
                        subset(focal.pred, seq(2, 21, by = 7)), iter)
```

```
## [1] "unique bins less than requested bins, decreasing bins to minimum"
```

```
## `summarise()` has grouped output by 'name'. You can override using the
## `.groups` argument.
```

```
F2.in <- compare_spearman(train.fisher %>% filter(!is.na(x_), id == "F2", id != "M5") %>% make_track(x_,y_) %>% ungroup(),
                       subset(focal.pred, seq(3, 21, by = 7)), iter)
```

```
## Adding missing grouping variables: `id`
```

```
## [1] "unique bins less than requested bins, decreasing bins to minimum"
```

```
## `summarise()` has grouped output by 'name'. You can override using the
## `.groups` argument.
```

```
F2.id <- compare_spearman(test.fisher %>% filter(!is.na(x_), id == "F2", id != "M5") %>% make_track(x_,y_),
                       subset(focal.pred, seq(3, 21, by = 7)), iter)
```

```
## [1] "unique bins less than requested bins, decreasing bins to minimum"
```

```
## `summarise()` has grouped output by 'name'. You can override using the
## `.groups` argument.
```

```
F2.out <- compare_spearman(Focal %>% filter(!is.na(x_), id != "F2", id != "M5") %>% make_track(x_,y_),
                        subset(focal.pred, seq(3, 21, by = 7)), iter)
```

```
## [1] "unique bins less than requested bins, decreasing bins to minimum"
```

```
## `summarise()` has grouped output by 'name'. You can override using the
## `.groups` argument.
```

```
M3.in <- compare_spearman(train.fisher %>% filter(!is.na(x_), id == "M3", id != "M5") %>% make_track(x_,y_) %>% ungroup(),
                       subset(focal.pred, seq(4, 21, by = 7)), iter)
```

```
## Adding missing grouping variables: `id`
```

```
## [1] "unique bins less than requested bins, decreasing bins to minimum"
```

```
## `summarise()` has grouped output by 'name'. You can override using the
## `.groups` argument.
```

```
M3.id <- compare_spearman(test.fisher %>% filter(!is.na(x_), id == "M3", id != "M5") %>% make_track(x_,y_),
                       subset(focal.pred, seq(4, 21, by = 7)), iter)
```

```
## [1] "unique bins less than requested bins, decreasing bins to minimum"
```

```
## `summarise()` has grouped output by 'name'. You can override using the
## `.groups` argument.
```

```
M3.out <- compare_spearman(Focal %>% filter(!is.na(x_), id != "M3", id != "M5") %>% make_track(x_,y_),
                        subset(focal.pred, seq(4, 21, by = 7)), iter)
```

```
## [1] "unique bins less than requested bins, decreasing bins to minimum"
```

```
## `summarise()` has grouped output by 'name'. You can override using the
## `.groups` argument.
```

```
F3.in <- compare_spearman(train.fisher %>% filter(!is.na(x_), id == "F3", id != "M5") %>% make_track(x_,y_) %>% ungroup(),
                       subset(focal.pred, seq(5, 21, by = 7)), iter)
```

```
## Adding missing grouping variables: `id`
```

```
## [1] "unique bins less than requested bins, decreasing bins to minimum"
```

```
## `summarise()` has grouped output by 'name'. You can override using the
## `.groups` argument.
```

```
F3.id <- compare_spearman(test.fisher %>% filter(!is.na(x_), id == "F3", id != "M5") %>% make_track(x_,y_),
                       subset(focal.pred, seq(5, 21, by = 7)), iter)
```

```
## [1] "unique bins less than requested bins, decreasing bins to minimum"
```

```
## `summarise()` has grouped output by 'name'. You can override using the
## `.groups` argument.
```

```
F3.out <- compare_spearman(Focal %>% filter(!is.na(x_), id != "F3", id != "M5") %>% make_track(x_,y_),
                        subset(focal.pred, seq(5, 21, by = 7)), iter)
```

```
## [1] "unique bins less than requested bins, decreasing bins to minimum"
```

```
## `summarise()` has grouped output by 'name'. You can override using the
## `.groups` argument.
```

```
M2.in <- compare_spearman(train.fisher %>% filter(!is.na(x_), id == "M2", id != "M5") %>% make_track(x_,y_) %>% ungroup(),
                      subset(focal.pred, seq(6, 21, by = 7)), iter)
```

```
## Adding missing grouping variables: `id`
```

```
## [1] "unique bins less than requested bins, decreasing bins to minimum"
```

```
## `summarise()` has grouped output by 'name'. You can override using the
## `.groups` argument.
```

```
M2.id <- compare_spearman(test.fisher %>% filter(!is.na(x_), id == "M2", id != "M5") %>% make_track(x_,y_),
                      subset(focal.pred, seq(6, 21, by = 7)), iter)
```

```
## [1] "unique bins less than requested bins, decreasing bins to minimum"
```

```
## `summarise()` has grouped output by 'name'. You can override using the
## `.groups` argument.
```

```
M2.out <- compare_spearman(Focal %>% filter(!is.na(x_), id != "M2", id != "M5") %>% make_track(x_,y_),
                       subset(focal.pred, seq(6, 21, by = 7)), iter)
```

```
## [1] "unique bins less than requested bins, decreasing bins to minimum"
```

```
## `summarise()` has grouped output by 'name'. You can override using the
## `.groups` argument.
```

```
F1.in <- compare_spearman(train.fisher %>% filter(!is.na(x_), id == "F1", id != "M5") %>% make_track(x_,y_) %>% ungroup,
                      subset(focal.pred, seq(7, 21, by = 7)), iter)
```

```
## Adding missing grouping variables: `id`
```

```
## [1] "unique bins less than requested bins, decreasing bins to minimum"
```

```
## `summarise()` has grouped output by 'name'. You can override using the
## `.groups` argument.
```

```
F1.id <- compare_spearman(test.fisher %>% filter(!is.na(x_), id == "F1", id != "M5") %>% make_track(x_,y_),
                      subset(focal.pred, seq(7, 21, by = 7)), iter)
```

```
## [1] "unique bins less than requested bins, decreasing bins to minimum"
```

```
## `summarise()` has grouped output by 'name'. You can override using the
## `.groups` argument.
```

```
F1.out <- compare_spearman(Focal %>% filter(!is.na(x_), id != "F1", id != "M5") %>% make_track(x_,y_),
                       subset(focal.pred, seq(7, 21, by = 7)), iter)
```

```
## [1] "unique bins less than requested bins, decreasing bins to minimum"
```

```
## `summarise()` has grouped output by 'name'. You can override using the
## `.groups` argument.
```

```
M1.new <- compare_spearman(test.fisher %>% filter(!is.na(x_), id == "M5") %>% make_track(x_,y_),
                       subset(non.focal.pred, seq(1, 21, by = 7)), iter)
```

```
## [1] "unique bins less than requested bins, decreasing bins to minimum"
```

```
## `summarise()` has grouped output by 'name'. You can override using the
## `.groups` argument.
```

```
M4.new <- compare_spearman(test.fisher %>% filter(!is.na(x_), id == "M5") %>% make_track(x_,y_),
                       subset(non.focal.pred, seq(2, 21, by = 7)), iter)
```

```
## [1] "unique bins less than requested bins, decreasing bins to minimum"
```

```
## `summarise()` has grouped output by 'name'. You can override using the
## `.groups` argument.
```

```
F2.new <- compare_spearman(test.fisher %>% filter(!is.na(x_), id == "M5") %>% make_track(x_,y_),
                       subset(non.focal.pred, seq(3, 21, by = 7)), iter)
```

```
## [1] "unique bins less than requested bins, decreasing bins to minimum"
```

```
## `summarise()` has grouped output by 'name'. You can override using the
## `.groups` argument.
```

```
M3.new <- compare_spearman(test.fisher %>% filter(!is.na(x_), id == "M5") %>% make_track(x_,y_),
                       subset(non.focal.pred, seq(4, 21, by = 7)), iter)
```

```
## [1] "unique bins less than requested bins, decreasing bins to minimum"
```

```
## `summarise()` has grouped output by 'name'. You can override using the
## `.groups` argument.
```

```
F3.new <- compare_spearman(test.fisher %>% filter(!is.na(x_), id == "M5") %>% make_track(x_,y_),
                       subset(non.focal.pred, seq(5, 21, by = 7)), iter)
```

```
## [1] "unique bins less than requested bins, decreasing bins to minimum"
```

```
## `summarise()` has grouped output by 'name'. You can override using the
## `.groups` argument.
```

```
M2.new <- compare_spearman(test.fisher %>% filter(!is.na(x_), id == "M5") %>% make_track(x_,y_),
                      subset(non.focal.pred, seq(6, 21, by = 7)), iter)
```

```
## [1] "unique bins less than requested bins, decreasing bins to minimum"
```

```
## `summarise()` has grouped output by 'name'. You can override using the
## `.groups` argument.
```

```
F1.new <- compare_spearman(test.fisher %>% filter(!is.na(x_), id == "M5") %>% make_track(x_,y_),
                      subset(non.focal.pred, seq(7, 21, by = 7)), iter)
```

```
## [1] "unique bins less than requested bins, decreasing bins to minimum"
```

```
## `summarise()` has grouped output by 'name'. You can override using the
## `.groups` argument.
```

```
id.names <- c(rep(c("M1", "M4", "F2", "M3", "F3", "M2", "F1"), each = 3*100*3),
              rep(c("M1", "M4", "F2", "M3", "F3", "M2", "F1"), each = 1*100*3))

type.names <- c(rep(rep(c("In-Sample","Within Individual", "Between Individuals"), each = 1*100*3), 7),
              rep(c("Between Contexts", "Between Contexts", "Between Contexts", "Between Contexts", "Between Contexts", "Between Contexts", "Between Contexts"), each = 1*100*3))

correlations.fisher.spearman <- rbind(M1.in,M1.id,M1.out,M4.in,M4.id,M4.out,F2.in,F2.id,F2.out,M3.in,M3.id,M3.out,F3.in,F3.id,F3.out,M2.in,M2.id,M2.out,F1.in,F1.id,F1.out,M1.new,M4.new,F2.new,M3.new,F3.new,M2.new,F1.new) %>%
  ungroup() %>% 
  mutate(id = id.names,
         bootstrap = factor(bootstrap),
         in.out = type.names,
         model = factor(name, levels = 1:3, labels = c("SSD", "SSF", "RSF"), ordered = T)) %>% 
  mutate(in.out = factor(in.out, levels = c("In-Sample","Within Individual", "Between Individuals", "Between Contexts"), ordered = T))


M1.in <- compare_mean(train.fisher %>% filter(!is.na(x_), id == "M1", id != "M5") %>% make_track(x_,y_) %>% ungroup(),
                       subset(focal.pred, seq(1, 21, by = 7)), iter)
```

```
## Adding missing grouping variables: `id`
```

```
M1.id <- compare_mean(test.fisher %>% filter(!is.na(x_), id == "M1", id != "M5") %>% make_track(x_,y_),
                       subset(focal.pred, seq(1, 21, by = 7)), iter)
M1.out <- compare_mean(Focal %>% filter(!is.na(x_), id != "M1", id != "M5") %>% make_track(x_,y_),
                        subset(focal.pred, seq(1, 21, by = 7)), iter)

M4.in <- compare_mean(train.fisher %>% filter(!is.na(x_), id == "M4", id != "M5") %>% make_track(x_,y_) %>% ungroup(),
                       subset(focal.pred, seq(2, 21, by = 7)), iter)
```

```
## Adding missing grouping variables: `id`
```

```
M4.id <- compare_mean(test.fisher %>% filter(!is.na(x_), id == "M4", id != "M5") %>% make_track(x_,y_),
                       subset(focal.pred, seq(2, 21, by = 7)), iter)
M4.out <- compare_mean(Focal %>% filter(!is.na(x_), id != "M4", id != "M5") %>% make_track(x_,y_),
                        subset(focal.pred, seq(2, 21, by = 7)), iter)

F2.in <- compare_mean(train.fisher %>% filter(!is.na(x_), id == "F2", id != "M5") %>% make_track(x_,y_) %>% ungroup(),
                       subset(focal.pred, seq(3, 21, by = 7)), iter)
```

```
## Adding missing grouping variables: `id`
```

```
F2.id <- compare_mean(test.fisher %>% filter(!is.na(x_), id == "F2", id != "M5") %>% make_track(x_,y_),
                       subset(focal.pred, seq(3, 21, by = 7)), iter)
F2.out <- compare_mean(Focal %>% filter(!is.na(x_), id != "F2", id != "M5") %>% make_track(x_,y_),
                        subset(focal.pred, seq(3, 21, by = 7)), iter)

M3.in <- compare_mean(train.fisher %>% filter(!is.na(x_), id == "M3", id != "M5") %>% make_track(x_,y_) %>% ungroup(),
                       subset(focal.pred, seq(4, 21, by = 7)), iter)
```

```
## Adding missing grouping variables: `id`
```

```
M3.id <- compare_mean(test.fisher %>% filter(!is.na(x_), id == "M3", id != "M5") %>% make_track(x_,y_),
                       subset(focal.pred, seq(4, 21, by = 7)), iter)
M3.out <- compare_mean(Focal %>% filter(!is.na(x_), id != "M3", id != "M5") %>% make_track(x_,y_),
                        subset(focal.pred, seq(4, 21, by = 7)), iter)

F3.in <- compare_mean(train.fisher %>% filter(!is.na(x_), id == "F3", id != "M5") %>% make_track(x_,y_) %>% ungroup(),
                       subset(focal.pred, seq(5, 21, by = 7)), iter)
```

```
## Adding missing grouping variables: `id`
```

```
F3.id <- compare_mean(test.fisher %>% filter(!is.na(x_), id == "F3", id != "M5") %>% make_track(x_,y_),
                       subset(focal.pred, seq(5, 21, by = 7)), iter)
F3.out <- compare_mean(Focal %>% filter(!is.na(x_), id != "F3", id != "M5") %>% make_track(x_,y_),
                        subset(focal.pred, seq(5, 21, by = 7)), iter)

M2.in <- compare_mean(train.fisher %>% filter(!is.na(x_), id == "M2", id != "M5") %>% make_track(x_,y_) %>% ungroup(),
                      subset(focal.pred, seq(6, 21, by = 7)), iter)
```

```
## Adding missing grouping variables: `id`
```

```
M2.id <- compare_mean(test.fisher %>% filter(!is.na(x_), id == "M2", id != "M5") %>% make_track(x_,y_),
                      subset(focal.pred, seq(6, 21, by = 7)), iter)
M2.out <- compare_mean(Focal %>% filter(!is.na(x_), id != "M2", id != "M5") %>% make_track(x_,y_),
                       subset(focal.pred, seq(6, 21, by = 7)), iter)

F1.in <- compare_mean(train.fisher %>% filter(!is.na(x_), id == "F1", id != "M5") %>% make_track(x_,y_) %>% ungroup,
                      subset(focal.pred, seq(7, 21, by = 7)), iter)
```

```
## Adding missing grouping variables: `id`
```

```
F1.id <- compare_mean(test.fisher %>% filter(!is.na(x_), id == "F1", id != "M5") %>% make_track(x_,y_),
                      subset(focal.pred, seq(7, 21, by = 7)), iter)
F1.out <- compare_mean(Focal %>% filter(!is.na(x_), id != "F1", id != "M5") %>% make_track(x_,y_),
                       subset(focal.pred, seq(7, 21, by = 7)), iter)

M1.new <- compare_mean(test.fisher %>% filter(!is.na(x_), id == "M5") %>% make_track(x_,y_),
                       subset(non.focal.pred, seq(1, 21, by = 7)), iter)

M4.new <- compare_mean(test.fisher %>% filter(!is.na(x_), id == "M5") %>% make_track(x_,y_),
                       subset(non.focal.pred, seq(2, 21, by = 7)), iter)

F2.new <- compare_mean(test.fisher %>% filter(!is.na(x_), id == "M5") %>% make_track(x_,y_),
                       subset(non.focal.pred, seq(3, 21, by = 7)), iter)

M3.new <- compare_mean(test.fisher %>% filter(!is.na(x_), id == "M5") %>% make_track(x_,y_),
                       subset(non.focal.pred, seq(4, 21, by = 7)), iter)

F3.new <- compare_mean(test.fisher %>% filter(!is.na(x_), id == "M5") %>% make_track(x_,y_),
                       subset(non.focal.pred, seq(5, 21, by = 7)), iter)

M2.new <- compare_mean(test.fisher %>% filter(!is.na(x_), id == "M5") %>% make_track(x_,y_),
                      subset(non.focal.pred, seq(6, 21, by = 7)), iter)

F1.new <- compare_mean(test.fisher %>% filter(!is.na(x_), id == "M5") %>% make_track(x_,y_),
                      subset(non.focal.pred, seq(7, 21, by = 7)), iter)

id.names <- c(rep(c("M1", "M4", "F2", "M3", "F3", "M2", "F1"), each = 3*100*3),
              rep(c("M1", "M4", "F2", "M3", "F3", "M2", "F1"), each = 1*100*3))

type.names <- c(rep(rep(c("In-Sample","Within Individual", "Between Individuals"), each = 1*100*3), 7),
              rep(c("Between Contexts", "Between Contexts", "Between Contexts", "Between Contexts", "Between Contexts", "Between Contexts", "Between Contexts"), each = 1*100*3))

length(type.names)
```

```
## [1] 8400
```

```
correlations.fisher.mean <- rbind(M1.in,M1.id,M1.out,M4.in,M4.id,M4.out,F2.in,F2.id,F2.out,M3.in,M3.id,M3.out,F3.in,F3.id,F3.out,M2.in,M2.id,M2.out,F1.in,F1.id,F1.out,M1.new,M4.new,F2.new,M3.new,F3.new,M2.new,F1.new) %>%
  ungroup() %>% 
  mutate(id = id.names,
         bootstrap = factor(bootstrap),
         in.out = type.names,
         model = case_when(str_detect(name, "RSF") ~ "RSF", str_detect(name, "SSF") ~ "SSF", str_detect(name, "SSD") ~ "SSD"),
         model = factor(model, levels = c("SSD", "SSF", "RSF"), ordered = T)) %>% 
  mutate(in.out = factor(in.out, levels = c("In-Sample","Within Individual", "Between Individuals", "Between Contexts"), ordered = T))


correlations.fisher.spearman$type <- "Spearman"
correlations.fisher.mean$type <- "Geometric Mean"

bound.comparisons.fisher <- rbind(correlations.fisher.spearman, correlations.fisher.mean)

fisher.data <- bound.comparisons.fisher %>% 
  mutate(model = factor(model, levels = c("SSD", "SSF", "RSF"), ordered = T)) %>% 
  mutate(in.out = factor(in.out, levels = c("In-Sample", "Within Individual", "Between Individuals", "Between Contexts"), ordered = T)) %>% 
  ungroup() %>% 
  dplyr::select(-name) %>% 
  group_by(id, in.out, model, type) %>%
  pivot_wider(names_from = c(model), values_from = measure) %>% 
  pivot_longer(c(RSF, SSF)) %>% 
  mutate(delta = SSD-value) %>%
  group_by(id, in.out, name, type) %>%
  summarize(median = median(delta),
            lower = quantile(delta, 0.025),
            upper = quantile(delta, 0.975)) %>%
  # pivot_wider(names_from = type, values_from = c(median, lower, upper))
  mutate(model = factor(name, levels = c("SSF","RSF"), labels = c("SSD-SSF","SSD-RSF"), ordered = T)) %>% 
  group_by(in.out, id, type) %>% 
  mutate(top = median == max(median))
```

```
## `summarise()` has grouped output by 'id', 'in.out', 'name'. You can override
## using the `.groups` argument.
```

```
fisher.data %>% 
  arrange(model) %>% 
  ggplot(aes(x = model, y = median, ymax = upper, ymin = lower, shape = type, group = paste(id), color = id)) +
  geom_hline(yintercept = 0) +
  geom_path(position = position_dodge(0.3)) +
  geom_pointrange(position = position_dodge(0.3)) +
  facet_grid(type~in.out, scales = "free_y") +  
  labs(x = "Prediction Comparison",
       y = "Difference from SSD Prediction\n(Median and 95% Quantile)",
       shape = "Comparison Heuristic") + 
  theme_classic() +
  scale_alpha_discrete(guide = F, range = c(0.5, 1)) +
  scale_y_continuous(trans = "both.sqrt.trans_trans")
```

```
## Warning: Using alpha for a discrete variable is not advised.
```

##### 3.3.2 Storing some output and plots for later

```
ssd.raster.fisher.df <- as.data.frame(rasterToPoints(focal.pred$SSD.M2.Focal))
ssf.prob.raster.fisher.df <- as.data.frame(rasterToPoints(focal.pred$SSF.M2.Focal))
rsf.prob.raster.fisher.df <- as.data.frame(rasterToPoints(focal.pred$RSF.M2.Focal))

colnames(ssd.raster.fisher.df)[3] <- "layer"
colnames(ssf.prob.raster.fisher.df)[3] <- "layer"
colnames(rsf.prob.raster.fisher.df)[3] <- "layer"

min.max.fisher <- rbind(ssd.raster.fisher.df, ssf.prob.raster.fisher.df, rsf.prob.raster.fisher.df)
min.fisher <- min(min.max.fisher$layer)
max.fisher <- max(min.max.fisher$layer)

ssd.fisher.plot <- ssd.raster.fisher.df %>% 
  ggplot() +
  geom_raster(aes(x = (x-min(x))/1000, y = (y - min(y))/1000, fill = layer), color = NA) +
  coord_equal() +
  scale_fill_viridis_c(option = "H", name = "Kernel Probability",
                       limits = c(min.fisher,max.fisher)) +
  # geom_point(data = deer, mapping = aes(x = x_, y = y_), color = "green", size = size.point, alpha = alpha.point) +
  # geom_path(data = ssf1, mapping = aes(x = x1_, y = y1_), color = "white", size = 0.1, alpha = 0.1) +
  theme.current+
  labs(x = "Easting (km)", y = "Northing (km)")
```

```
## Warning in geom_raster(aes(x = (x - min(x))/1000, y = (y - min(y))/1000, :
## Ignoring unknown parameters: `colour`
```

```
ssf.fisher.plot <- ssf.prob.raster.fisher.df %>% 
  ggplot() +
  geom_raster(aes(x = (x-min(x))/1000, y = (y - min(y))/1000, fill = layer), color = NA) +
  coord_equal() +
  scale_fill_viridis_c(option = "H", name = "Kernel Probability",
                       limits = c(min.fisher,max.fisher)) +
  # geom_point(data = deer, mapping = aes(x = x_, y = y_), color = "green", size = size.point, alpha = alpha.point) +
  # geom_path(data = ssf1, mapping = aes(x = x1_, y = y1_), color = "white", size = 0.1, alpha = 0.1) +
  theme.current+
  labs(x = "Easting (km)", y = "Northing (km)")
```

```
## Warning in geom_raster(aes(x = (x - min(x))/1000, y = (y - min(y))/1000, :
## Ignoring unknown parameters: `colour`
```

```
rsf.fisher.plot <- rsf.prob.raster.fisher.df %>% 
  ggplot() +
  geom_raster(aes(x = (x-min(x))/1000, y = (y - min(y))/1000, fill = layer), color = NA) +
  coord_equal() +
  scale_fill_viridis_c(option = "H", name = "Kernel Probability",
                       limits = c(min.fisher,max.fisher)) +
  # geom_point(data = deer, mapping = aes(x = x_, y = y_), color = "green", size = size.point, alpha = alpha.point) +
  # geom_path(data = ssf1, mapping = aes(x = x1_, y = y1_), color = "white", size = 0.1, alpha = 0.1) +
  theme.current+
  labs(x = "Easting (km)", y = "Northing (km)")
```

```
## Warning in geom_raster(aes(x = (x - min(x))/1000, y = (y - min(y))/1000, :
## Ignoring unknown parameters: `colour`
```

```
fisher <- ggarrange(rsf.fisher.plot, ssf.fisher.plot, ssd.fisher.plot, common.legend = T, nrow = 1, legend = "left")
fisher
```

#### 3.4 Plotting for the main results figures

#### 3.5 Maps

```
colnames(use.area.dee1)[1] <- "model"
use.area.dee1$animal <- ""
use.area.dee1 <- use.area.dee1[,c(1,3,2)]
use.area.dee1$data <- "Red Deer"
use.area.deer2$data <- "Roe Deer"
use.area.fisher$data <- "Fisher"
use.pred <- rbind(use.area.dee1, use.area.deer2, use.area.fisher)

entropy <- use.pred %>% 
  group_by(animal) %>% 
  mutate(data = factor(data, levels = c("Red Deer", "Roe Deer", "Fisher"))) %>% 
  mutate(model = factor(model, levels = c("RSF", "SSF", "SSD"))) %>% 
  pivot_wider(names_from = "model", values_from = "sum") %>% 
  pivot_longer(cols = c("RSF", "SSF")) %>% 
  mutate(change = (SSD - value)/value) %>% 
  group_by(data, name) %>% 
  mutate(min = min(change),
         max = max(change),
         mean = mean(change)) %>% 
  ggplot(aes(y = mean, x = name, ymin = min, ymax = max)) +
  geom_hline(yintercept = 0) +
  geom_pointrange(width = 0.25) +
  geom_point(mapping = aes(y=change), position = position_jitter(0.1), alpha = 0.25) +
  scale_color_scico_d() +
  scale_y_continuous(labels = scales::percent) + 
  labs(y = "50% Use Area of SSD Relative to RSF and SSF",
       x = "Prediction\nKernel",
       color = "Prediction Type") +
  theme_classic() +
  facet_grid(data~.) +
  scale_alpha_discrete(guide = "none", range = c(0.25, 1))
```

```
## Warning in geom_pointrange(width = 0.25): Ignoring unknown parameters: `width`
```

```
## Warning: Using alpha for a discrete variable is not advised.
```

```
prediction.maps <- ggarrange(ggarrange(simple.deer, complex.deer, fisher, ncol = 1, nrow = 3, legend = "left"), entropy, nrow = 1, widths = c(0.4, 0.15), align = "h")
```

```
## Warning: Graphs cannot be horizontally aligned unless the axis parameter is
## set. Placing graphs unaligned.
```

```
# ggsave("Plots/PredictionRasters2.svg", prediction.maps, width = 10, height = 8)
prediction.maps
```

#### 3.6 Correlations and means

```
a <- deer.data.1 %>% 
  filter(type == "Geometric Mean") %>% 
  ungroup() %>% 
  add_row(median = c(0), 
          model = factor(c("SSF","SSF","SSF","SSF"), levels = c("SSF","RSF"), labels = c("SSD-SSF","SSD-RSF"), ordered = T),
          in.out = factor(c("Between Individuals", "Between Contexts","Between Individuals", "Between Contexts"), levels = c("In-Sample","Within Individual", "Between Individuals", "Between Contexts"), ordered = T),
          type = c("Geometric Mean","Geometric Mean", "Spearman", "Spearman")) %>% 
  group_by(model, type, in.out) %>% 
  mutate(pop.med = median(median)) %>% 
  arrange(model) %>% 
  ggplot(aes(x = model, y = median, ymax = upper, ymin = lower, group = type)) +
  geom_hline(yintercept = 0) +
  # geom_path(position = position_dodge(0.2)) +
  geom_pointrange(position = position_dodge(0.4)) +
  facet_grid(~in.out) +
  labs(x = "Prediction Comparison",
       y = "Difference from SSD Prediction\n(Median and 95% Quantile)",
       shape = "Comparison Heuristic") + 
  theme_classic() +
  scale_alpha_discrete(guide = F, range = c(0.5, 1))+
  scale_y_continuous(trans = "both.sqrt.trans_trans") + 
  scale_x_discrete(labels = c("SSD-SSF" = expression(delta, "SSF"),
                              "SSD-RSF" = expression(delta, "SSF")))
```

```
## Warning: Using alpha for a discrete variable is not advised.
```

```
b <- deer.data.2 %>% 
  filter(type == "Geometric Mean") %>% 
   ungroup() %>% 
    add_row(median = c(0), 
          model = factor(c("SSF","SSF"), levels = c("SSF","RSF"), labels = c("SSD-SSF","SSD-RSF"), ordered = T),
          in.out = factor(c("Between Contexts", "Between Contexts"), levels = c("In-Sample","Within Individual", "Between Individuals", "Between Contexts"), ordered = T),
          type = c("Geometric Mean","Geometric Mean")) %>% 
  group_by(model, type, in.out) %>% 
  mutate(pop.med = median(median)) %>% 
  ggplot(aes(x = model, y = median, ymax = upper, ymin = lower, group = paste(id), color = id)) +
  geom_hline(yintercept = 0) +
  # geom_path(position = position_dodge(0.2)) +
  geom_pointrange(position = position_dodge(0.4)) +
  geom_point(aes(y = pop.med), color = "red") +
  facet_grid(~in.out) +
  labs(x = "Prediction Comparison",
       y = "Difference from SSD Prediction\n(Median and 95% Quantile)",
       shape = "Comparison Heuristic",
       color = "ID") + 
  MetBrewer::scale_color_met_d(name = "Navajo") +
  theme_classic() +
  scale_y_continuous(trans = "both.sqrt.trans_trans")

c <- fisher.data %>% 
  filter(type == "Geometric Mean") %>% 
  group_by(model, type, in.out) %>% 
  mutate(pop.med = median(median)) %>% 
  ggplot(aes(x = model, y = median, ymax = upper, ymin = lower, group = paste(id), color = id)) +
  geom_hline(yintercept = 0) +
  # geom_path(position = position_dodge(0.2)) +
  geom_pointrange(position = position_dodge(0.4)) +
  geom_point(aes(y = pop.med), color = "red") + 
  facet_grid(~in.out) +
  labs(x = "Prediction Comparison",
       y = "Difference from SSD Prediction\n(Median and 95% Quantile)",
       shape = "Comparison Heuristic",
       color = "ID") +
  MetBrewer::scale_color_met_d(name = "Tiepolo", guide = F) +
  theme_classic() +
  scale_y_continuous(trans = "both.sqrt.trans_trans")

d <- deer.data.1 %>% 
  filter(type == "Spearman") %>% 
  ungroup() %>% 
  add_row(median = c(0), 
          model = factor(c("SSF","SSF","SSF","SSF"), levels = c("SSF","RSF"), labels = c("SSD-SSF","SSD-RSF"), ordered = T),
          in.out = factor(c("Between Individuals", "Between Contexts","Between Individuals", "Between Contexts"), levels = c("In-Sample","Within Individual", "Between Individuals", "Between Contexts"), ordered = T),
          type = c("Geometric Mean","Geometric Mean", "Spearman", "Spearman")) %>% 
  group_by(model, type, in.out) %>% 
  mutate(pop.med = median(median)) %>% 
  arrange(model) %>% 
  ggplot(aes(x = model, y = median, ymax = upper, ymin = lower, group = type)) +
  geom_hline(yintercept = 0) +
  # geom_path(position = position_dodge(0.2)) +
  geom_pointrange(position = position_dodge(0.4)) +
  facet_grid(~in.out) +
  labs(x = "Prediction Comparison",
       y = "Difference from SSD Prediction\n(Median and 95% Quantile)",
       shape = "Comparison Heuristic") + 
  theme_classic() +
  scale_alpha_discrete(guide = F, range = c(0.5, 1))+
  scale_y_continuous(trans = "both.sqrt.trans_trans")
```

```
## Warning: Using alpha for a discrete variable is not advised.
```

```
e <- deer.data.2 %>% 
  filter(type == "Spearman") %>% 
   ungroup() %>% 
    add_row(median = c(0), 
          model = factor(c("SSF","SSF","SSF","SSF"), levels = c("SSF","RSF"), labels = c("SSD-SSF","SSD-RSF"), ordered = T),
          in.out = factor(c("Between Individuals", "Between Contexts","Between Individuals", "Between Contexts"), levels = c("In-Sample","Within Individual", "Between Individuals", "Between Contexts"), ordered = T),
          type = c("Geometric Mean","Geometric Mean", "Spearman", "Spearman")) %>% 
  group_by(model, type, in.out) %>% 
  mutate(pop.med = median(median)) %>% 
  group_by(id, in.out) %>% 
  mutate(top = median == max(median)) %>% 
  ggplot(aes(x = model, y = median, ymax = upper, ymin = lower, group = paste(id), color = id)) +
  geom_hline(yintercept = 0) +
  # geom_path(position = position_dodge(0.2)) +
  geom_pointrange(position = position_dodge(0.4)) + 
  geom_point(aes(y = pop.med), color = "red") + 
  facet_grid(~in.out) +
  labs(x = "Prediction Comparison",
       y = "Difference from SSD Prediction\n(Median and 95% Quantile)",
       shape = "Comparison Heuristic",
       color = "ID") + 
  MetBrewer::scale_color_met_d(name = "Navajo") +
  theme_classic() +
  scale_y_continuous(trans = "both.sqrt.trans_trans")

f <- fisher.data %>% 
  filter(type == "Spearman") %>% 
  group_by(id, in.out) %>% 
  mutate(top = median == max(median)) %>% 
  group_by(model, type, in.out) %>% 
  mutate(pop.med = median(median)) %>% 
  ggplot(aes(x = model, y = median, ymax = upper, ymin = lower, group = paste(id), color = id)) +
  geom_hline(yintercept = 0) +
  # geom_path(position = position_dodge(0.2)) +
  geom_pointrange(position = position_dodge(0.4)) +
  geom_point(aes(y = pop.med), color = "red") +
  facet_grid(~in.out) +
  labs(x = "Prediction Comparison",
       y = "Difference from SSD Prediction\n(Median and 95% Quantile)",
       shape = "Comparison Heuristic",
       color = "ID") +
  MetBrewer::scale_color_met_d(name = "Tiepolo") +
  theme_classic() +
  scale_y_continuous(trans = "both.sqrt.trans_trans")

compiled.mlpu <- ggarrange(a,b,c, align = "v", nrow = 3)
```

```
## Warning: Removed 2 rows containing missing values or values outside the scale range
## (`geom_segment()`).
```

```
## Warning: Removed 2 rows containing missing values or values outside the scale range
## (`geom_segment()`).
## Removed 2 rows containing missing values or values outside the scale range
## (`geom_segment()`).
```

```
compiled.scor <- ggarrange(d,e,f, align = "v", nrow = 3)
```

```
## Warning: Removed 2 rows containing missing values or values outside the scale range
## (`geom_segment()`).
## Removed 2 rows containing missing values or values outside the scale range
## (`geom_segment()`).
## Removed 2 rows containing missing values or values outside the scale range
## (`geom_segment()`).
## Removed 2 rows containing missing values or values outside the scale range
## (`geom_segment()`).
```

```
total <-ggarrange(compiled.mlpu, compiled.scor, labels = c("A","B"))
# ggsave("Plots/FigComparisons.svg", total, width = 15, height = 10)

compiled.mlpu
```

```
# ggsave("Plots/Fig3A2.1.svg", width = 15, height = 10)
compiled.scor
```

```
# ggsave("Plots/Fig3A2.2.svg", width = 15, height = 10)
```

```
deer.table1 <- deer.data.1 %>% 
  mutate(id = "") %>% 
  group_by(in.out,id, type,model) %>% 
  arrange(-median) %>% 
  slice(1) %>% 
  mutate(top = case_when(
    median > 0 ~ "SSD",
    median < 0 ~ name,
    median == 0 ~ "Tie"
  )) %>% 
  dplyr::select(in.out,id, top)%>% 
  mutate(data = "Red Deer")
```

```
## Adding missing grouping variables: `type`, `model`
```

```
deer.table2 <- deer.data.2 %>% 
  group_by(in.out,id, type,model) %>% 
  arrange(-median) %>% 
  slice(1) %>% 
  mutate(top = case_when(
    median > 0 ~ "SSD",
    median < 0 ~ name,
    median == 0 ~ "Tie"
  )) %>% 
  dplyr::select(in.out,id, top)%>% 
  mutate(data = "Roe Deer")
```

```
## Adding missing grouping variables: `type`, `model`
```

```
fisher.table <- fisher.data %>% 
  group_by(in.out,id, type,model) %>% 
  arrange(-median) %>% 
  slice(1) %>% 
  mutate(top = case_when(
    median > 0 ~ "SSD",
    median < 0 ~ name,
    median == 0 ~ "Tie"
  )) %>% 
  dplyr::select(in.out,id, top)%>% 
  mutate(data = "Fisher")
```

```
## Adding missing grouping variables: `type`, `model`
```

```
values = rev(c(scico(n=3, begin = 0.5, palette = "batlow"), "grey90"))

rbind(deer.table1, deer.table2, fisher.table) %>% 
  mutate(data = factor(data, levels = (c("Red Deer", "Roe Deer", "Fisher")))) %>% 
  mutate(top = factor(top, levels = rev(c("SSD", "RSF", "SSF", "Tie")))) %>% 
  group_by(type, top, in.out) %>% 
  ggplot(aes(x = type, fill = top)) +
  geom_bar(position = position_fill()) +
  facet_grid(data~model + in.out) +
  theme_classic() +
  scale_fill_manual(values = values)
```

```
bound.comparisons.deer$id <- "1"
bound.comparisons.deer <- bound.comparisons.deer[,c(1,2,3)]
bound.comparisons.deer.2
```

```
## # A tibble: 9,000 × 7
##    name  bootstrap measure id    in.out    model type    
##    <fct> <fct>       <dbl> <chr> <ord>     <ord> <chr>   
##  1 1     1           0.939 M03   In-Sample SSD   Spearman
##  2 1     2           0.936 M03   In-Sample SSD   Spearman
##  3 1     3           0.948 M03   In-Sample SSD   Spearman
##  4 1     4           0.930 M03   In-Sample SSD   Spearman
##  5 1     5           0.903 M03   In-Sample SSD   Spearman
##  6 1     6           0.912 M03   In-Sample SSD   Spearman
##  7 1     7           0.912 M03   In-Sample SSD   Spearman
##  8 1     8           0.954 M03   In-Sample SSD   Spearman
##  9 1     9           0.912 M03   In-Sample SSD   Spearman
## 10 1     10          0.967 M03   In-Sample SSD   Spearman
## # ℹ 8,990 more rows
```

```
bound.comparisons.fisher
```

```
## # A tibble: 16,800 × 7
##    name  bootstrap measure id    in.out    model type    
##    <fct> <fct>       <dbl> <chr> <ord>     <ord> <chr>   
##  1 1     1           0.697 M1    In-Sample SSD   Spearman
##  2 1     2           0.648 M1    In-Sample SSD   Spearman
##  3 1     3           0.661 M1    In-Sample SSD   Spearman
##  4 1     4           0.782 M1    In-Sample SSD   Spearman
##  5 1     5           0.784 M1    In-Sample SSD   Spearman
##  6 1     6           0.648 M1    In-Sample SSD   Spearman
##  7 1     7           0.806 M1    In-Sample SSD   Spearman
##  8 1     8           0.539 M1    In-Sample SSD   Spearman
##  9 1     9           0.766 M1    In-Sample SSD   Spearman
## 10 1     10          0.802 M1    In-Sample SSD   Spearman
## # ℹ 16,790 more rows
```

```
colnames(bound.comparisons.deer)
```

```
## [1] "name"      "bootstrap" "measure"
```

```
colnames(bound.comparisons.deer.2)
```

```
## [1] "name"      "bootstrap" "measure"   "id"        "in.out"    "model"    
## [7] "type"
```
